# Small Extracellular Vesicles by Nucleus Pulposus Cells Maintain Niche and Cell Homeostasis via Receptor Shuffling and Metabolic Enzyme Supplements

**DOI:** 10.1101/2024.12.12.628054

**Authors:** Ankita Samanta, Mi-Jeong Yoo, Jin Koh, Thomas Lufkin, Petra Kraus

## Abstract

**Background:** Small extracellular vesicles (sEV) are a conserved mean of communication across the domains of life and lately gained more interest in mammalian non-cancerous work as non-cellular, biological therapeutic with encouraging results in recent studies of chronic degenerative diseases. The nucleus pulposus (NP) is the avascular and aneural center of an intervertebral disc (IVD), home to unique niche conditions and affected in IVD degeneration. We investigated the proteome of autologous and mesenchymal stem cell (MSC) sEVs for their potential to contribute to cell and tissue homeostasis in this niche.

**Methods:** We aimed to identify characteristics of NP sEVs via mass spectrometric proteome and functional enrichment analysis. We explored the proteome profile of sEVs generated by autologous parent cell lines from bovine coccygeal IVD and adipose tissue, with a focus on NP cells, using adult and fetal donors. We compared these findings to published sEV databases and MSC sEV data.

**Findings:** We propose several mechanisms associated with NP sEVs: 1. Membrane receptor trafficking to modify signal responses promoting niche homeostasis. 2. Cell homeostasis via proteasome delivery; 3. Redox and energy homeostasis via metabolic enzymes delivery. 4. Immunomodulation beyond an association with a serum protein corona.

**Conclusion:** The proteome signature of sEVs generated by NP parent cells is similar to previously published sEV data, yet with a focus on supplementing anaerobic metabolism and redox balance while contributing to the maintenance of an aneural and avascular microniche.

## BACKGROUND

Low back pain (LBP) and neck pain present global health issues as they are significant causes of disability worldwide. Every year, an estimated 200 billion dollars are spent on treating LBP (1). It is the most frequent cause of worker’s compensation, lost working time, and decreased productivity (1,2). Even if the etiology of most cases of back pain is unknown, intervertebral disc degeneration (IVDD) is one of the most common factors (3,4). IVDD principally impacts the intervertebral disc (IVD), resulting in structural modifications and functional impairments within the spinal column (5–7). Current therapy options for IVDD alleviate symptoms but fail to cure the underlying problem (6,8–10). Difficulty of effectively addressing this complex and multidimensional illness with conventional approaches hampers clinical success (11). Regenerative medicine aims to repair the tissues and organs damaged by disease, trauma, or congenital disabilities and efforts have been made to develop novel efficient and safe non-operative strategies (6). A significant study area is the direct injection of active compounds to prevent, slow down, or reverse IVDD (12,13). Since the last 30 years, hundreds of clinical trials have investigated biologic, cell-based, and scaffold-based injectable approaches towards symptomatic IVDD as reviewed in detail elsewhere (6,14–20). Several preclinical animal studies support each clinical trial (15,21–24). Mesenchymal stem cell (MSC) therapy could stimulate proteoglycan (PG) and collagen production to restore disc height (9). Nucleus pulposus (NP)-MSCs have been identified in both degenerative and normal IVD tissues (25). Notably, NP-MSCs have a remarkable resilience to acidic conditions *in vitro* compared to other MSCs, alongside increased expression levels of key extracellular matrix (ECM) components such as PGs and collagen II (25). Transplanting NP-MSCs into the IVD for differentiation into NP cells is a regenerative approach for IVDD (26). However, the harsh and hypoxic microenvironment in the mature NP challenges the viability of MSCs further limiting the density of viable cells and PG production rate, making it uncertain whether therapeutic MSC will be successful long term (27–33).

The IVD is made up of a central NP surrounded by an inner and outer annulus fibrosus (AF), which is present in between the cartilaginous endplates (CEPs) (Figure 1A) (27,34–41). The NP has a relatively loose collagen network, is highly hydrated, and exhibits hydrostatic behavior under load when non-degenerate (6,27,38,42). The main PG of the disc is aggrecan (Acan), which offers the osmotic characteristics required to withstand compression due to its high anionic glycosaminoglycan (GAG) content, specifically chondroitin sulfate and keratan sulfate (43,44). The AF is mostly made up of collagen type I (38,42,43,45). Cells of the outer AF appear elongated, thin, and parallel to the collagen lamellae, while the inner AF looks fibrocartilaginous (27,37,46,47). Proteoglycans, particularly Acan, comprise 5-15 % of the weight of wet tissue. Their charges ensure the swelling pressure necessary to maintain disc height and turgor under compressive load by attracting water molecules into the disc (27,48,49). One of the earliest changes in IVDD is PG-loss and consequentially reduced water retention, resulting in decreased disc height and flexibility (27). NP cells originate from the notochord (NC), a transient embryonic structure (37,50,51). In humans, these NC cells are replaced by chondrocyte-like cells (CLC) around the age of ten years (45,51–61). However, NC cells are retained into mature adulthood in other species like rodents, and pigs (37,62). These cells continue to generate PGs and collagen; however, with time, the NP becomes stiffer and less hydrated (63). The adult NP has a low cell density, with cells occupying less than 0.5% of the tissue volume (32). Unlike other tissues, NP cells are embedded in a vast amount of ECM (38,42). With aging, the density of active NP cells declines further, presumably due to decreased oxygen and nutrient supply in this slightly acidic tissue (64–66). Only recently, more information has become available on the proteome and transcriptome of healthy or degenerated IVDs from various species (25,67–75) and more potential biomarkers defining NP and AF cells join those previously established on the transcriptome level (68,73,76,77). This data allows for a characterization of different individual cell types of the heterogeneous NP cell population and to define hallmarks for regenerative therapies to maintain or restore IVD tissue composition.

**Figure 1:**
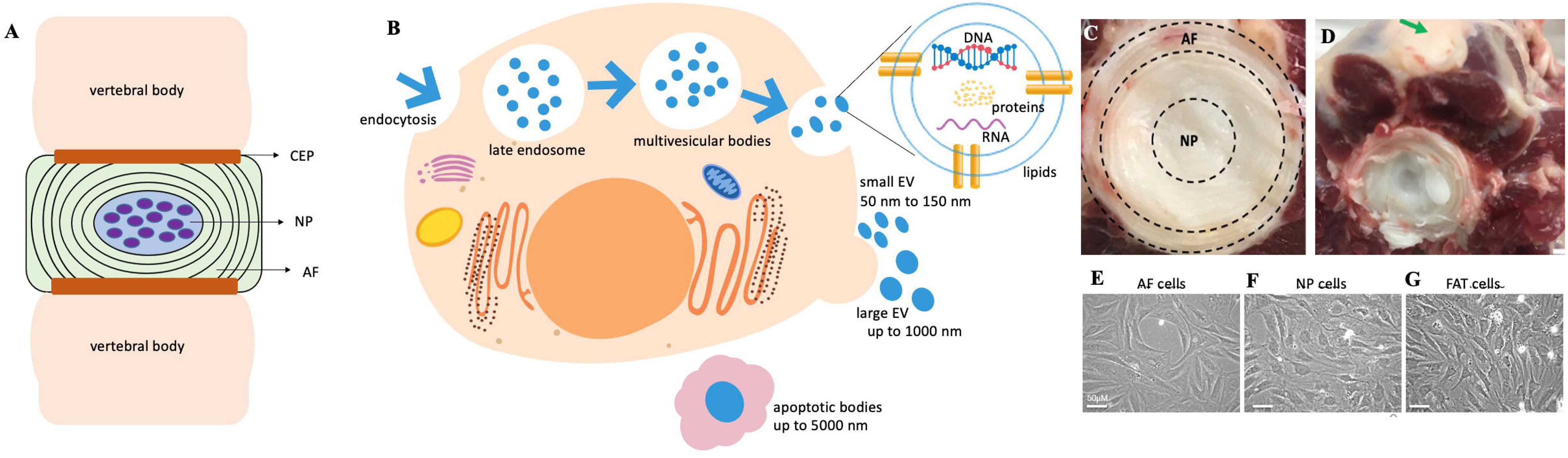
Isolation of bovine sEVs. A) Schematic representation of the IVD containing NP and AF present in between the cartilaginous endplates. This illustration was created on the Wondershare Edraw Max Platform. (https://edrawmax.wondershare.com/). B) Small EV biogenesis and composition. C) A cut through the bovine tail shows the coccygeal IVD. D) Bovine tail, the green arrow indicates adipose tissue used to derive FAT cells. E) AF cells F) NP cells G) FAT cells in 2D culture. The scale bar represents 50 µm. AF: Annulus fibrosus; CEP: Cartilaginous endplate; DNA: Deoxyribonucleic acid; FAT: Adipose tissue; IVD: Intervertebral disc; NP: Nucleus pulposus; RNA: Ribonucleic acid.

The homeostasis of a microenvironment is naturally maintained via effective cell-cell communication (78). The healthy NP is avascular and aneural (32,41,79,80), hence in need for non-standard means of cellular communication to maintain tissue homeostasis. The development of large-scale “-omics” technologies has improved our understanding of the secretome including extracellular vesicles (EVs) with specific cargo of proteins and nucleic acids (78,81). Although the classification of EVs constantly changes (82) they usually fall into one of three categories based on size (Figure 1B): Apoptotic bodies (up to 5000nm), ectosomes (up to 1000nm), and small extracellular vesicles (sEV) (50–150nm), previously referred to as exosomes (83,84). Recent guidelines recommend a nomenclature limiting the term “exosome” to only those sEVs demonstrated as generated via multivesicular bodies (MVB) (82). Most cell types produce sEVs, and their release into body fluids and culture media has sparked interest in the identification of biomarkers (85,86). The size, composition, functional influence on recipient cells, and the biological origin of sEVs all contribute to their heterogeneity (78,83). Small EVs with unique proteins, lipids, and nucleic acids specific to their parent cells are now recognized as intercellular communication methods (87). Small EVs as generated by parent cells may interact with target cells, conveying cargo of bioactive molecules to recipient cells thereby influencing their behavior and phenotypic traits by impacting cellular processes such as cell proliferation, differentiation, immune responses, and gene expression (83,88,89). Small EVs play significant roles in disease progression, including cancer and neurodegenerative disorders (90).

The a-cellular nature of sEVs offers novel opportunities for IVDD therapy unaffected by constrains of stem cell therapies such as tumorigenesis, cell rejection or cell survival (81,91,92). As part of the cell secretome sEVs maintain the therapeutic benefits of their parent cell and protect their cargo via a phospholipid membrane (81,93). Increasing interest also exists in EVs produced by resident cells to identify cell or disease-specific biomarkers, similar to biomarker research in oncology (94–97). Given the socioeconomic impact of IVDD, IVD sEV biomarkers would add to a refined IVD cell phenotype analysis. Previous research has focused on the function and composition of NC cell derived EVs, with a focus on microRNAs (miRNAs). These small non-coding RNA molecules are being studied extensively for their therapeutic potential in other domains (98). NC cell-derived EVs enhanced DNA and GAG content in human NP cell micro-aggregates compared to untreated control conditions, although the underlying mechanism and associated EV content were not determined (99). Limited research has been conducted on IVD-derived EVs in the context of IVD homeostasis, largely focusing on benefits of MSC derived sEVs (92,93,100–107). Investigating NP cell derived sEV cargo can provide insight into cell-cell communication and signaling cascades that maintain IVD homeostasis and enable novel strategies to alleviate IVDD (6,81,100). In a recent study, EVs derived from NP cells promoted the proliferation of degenerated NP cells and reduce senescence *in vitro*, while also ameliorating pain and attenuating IVDD *in vivo* (108). For a degenerative disease like IVDD cell therapies employing MSC for tissue regeneration are considered. Based on IVD location and anatomy, cell sources such as autologous cells of the healthy NP would be difficult to harvest without inflicting organ damage (Figure 1). While peripheral AF cells might be easier to access, adipose tissue would be the most readily accessible source for autologous MSC (109–111). However, dysfunctional fat metabolism and chronic activation of ERK/MAPK signaling has been linked to tissue aging in a number of animal models including humans (112–114).

Here we investigate the proteome of sEVs generated by cells of the bovine IVD and adipose tissues. Bovine coccygeal IVDs are considered similar to healthy human IVDs (115–118). Through gene ontology (GO) annotation and functional enrichment analyses we aimed to 1) investigate similarities and differences in the sEV cargo protein profiles of autologous bovine NP, AF, and adipose (FAT) parent cells, adult and fetal NP parent cells, and MSC parent cells and 2) perform comparative sEV proteomics studies between NP parent cells and their sEVs to find proteins that are actively deposited in or removed from the cargo. We identified sEV proteins and molecular pathways relevant for NP cell homeostasis to serve as targets for future therapeutic IVDD interventions.

## MATERIAL AND METHODS

### Cell Culture and Small EV Isolation from Primary IVD Cells

The work has been reported in line with the ARRIVE guidelines 2.0 and is covered under IACUC protocol/approval number 19-04, approved 10/24/2021. Project title: Analysis of Gene Expression. Institution: Clarkson University. Primary bovine coccygeal cell-lines derived from adult and fetal central NPs (referred to as NP), outer AFs (referred to as AF) and adipose tissues (referred to as FAT) were maintained in Dulbecco’s Modified Eagles Medium (DMEM, GIBCO) with 10% (v/v) heat inactivated fetal bovine serum (HI-FBS, GIBCO) and trypsinized with 0.05% Trypsin/EDTA (GIBCO) once 80-90% confluent as previously described (55,99) (Figure 1). Cells were aliquoted and stored in liquid nitrogen at passage (p) two or three. For sEV harvest cells were expanded to ∼70 % confluency in 20 x 150 mm culture dishes, washed three times with 1 × phosphate-buffered saline (PBS), then cultured in a total of 300 ml DMEM with 10% (v/v) sEV-depleted FBS (ED-FBS) (GIBCO). After 48 h at 37°C under normal oxygen and 5% CO_2_ supernatants were harvested for sEV isolation. A total cell count was estimated from the average of three 150 mm plates. Cells from two plates were lysed in cold 1 × RIPA lysis buffer (50 mM Tris HCl, 150 mM NaCl, 1.0% (v/v) NP40, 0.5% (w/v) sodium deoxycholate, 1.0 mM EDTA, 0.1% (w/v) SDS and 0.01% (w/v) sodium azide, pH 7.4) and stored at -80°C for further protein analysis. One plate from each cell line was subjected to senescence associated (SA)-β-galactosidase staining for quality control (119). Three biological replicates, NP cell lines TT32, TT33, and TT39 from adult donors were used for the quantitative proteome analysis of NP sEVs, while NP, AF and FAT cell lines TT39 and fetal NP cells were used for explorative analysis of sEV proteome cargo.

Small EVs for proteome analysis were generally isolated as illustrated (Figure 2A) using differential ultracentrifugation (DUC) (120,121). Cell debris was eliminated at 300 × *g* in an Eppendorf 5810 centrifuge followed by filtration of the supernatant (0.22 µm cut-off). Larger EVs were removed from the supernatant through DUC at 2000 (2k) × *g* for 30 min at 4°C in a J2-21 Sorvall ultracentrifuge (rotor JA-20), followed by 10k × *g* for 1 h at 4°C and 130 k × *g* for 2 h at 4°C in a Beckman Optima XPN-100 ultracentrifuge (rotor Ti45). Each pellet was resuspended in 1 × PBS and stored at -80°C. Final pellets were washed with 1 × PBS and repelleted at 130 k × *g* for 2 h at 4°C before the final resuspension of the sEV fraction in 1 × PBS and storage at -80°C. Triplicate measurements of protein concentration were conducted with a NanoDrop ND-1000 spectrophotometer (Thermo Fisher Scientific) and Bradford assay. All solutions and consumables were nuclease and protease-free. Water or 1 × PBS was filtered with a 0.22 µm cut-off to eliminate dust particles.

**Figure 2:**
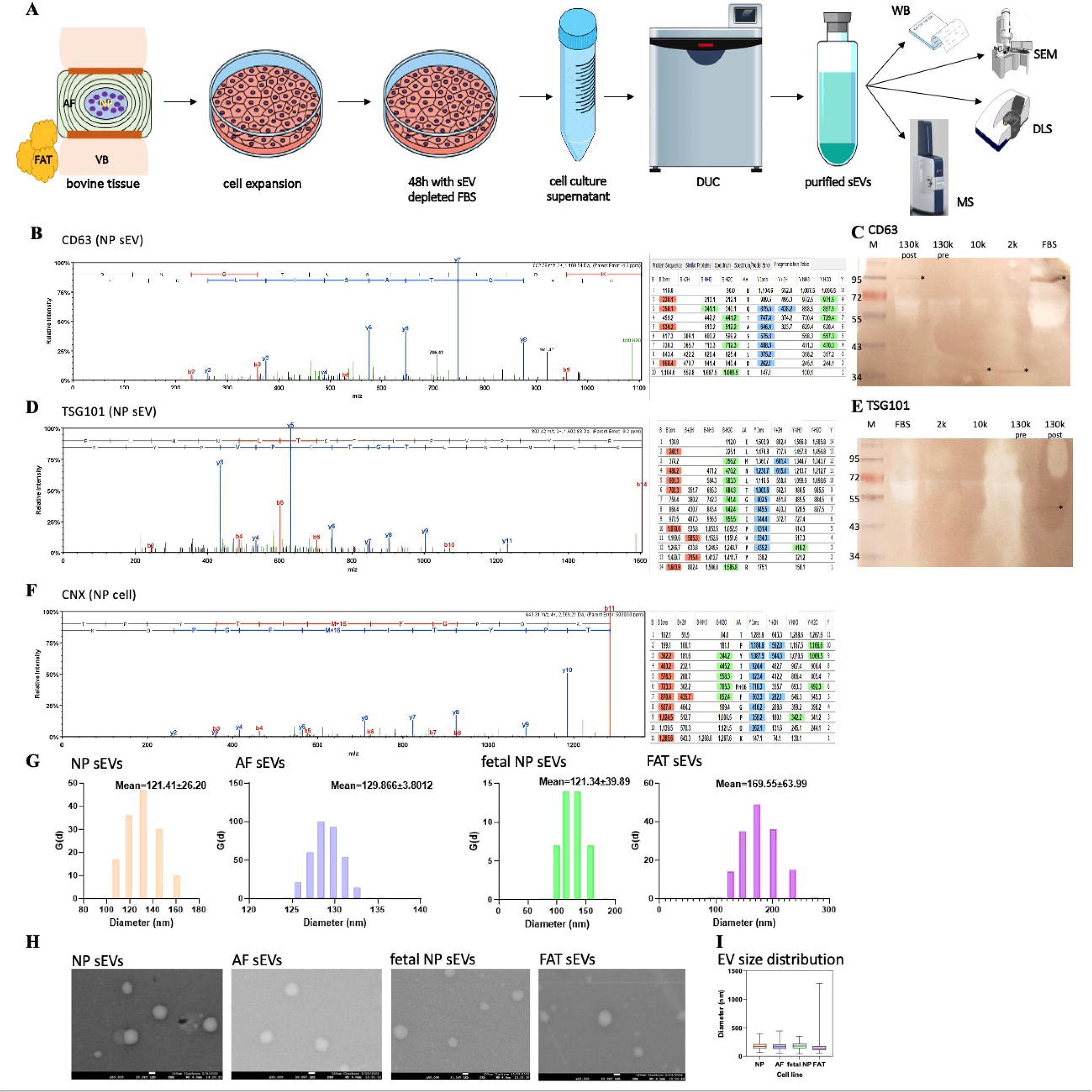
Isolation and characterization of sEVs. A) Simplified illustration of the workflow for sEV harvesting and downstream analysis. B) Spectrum for CD63. C) Cropped Western blot for CD63, signal is indicated by an asterix. Full-length blots are presented in Supplementary figure 1. D) Spectrum for TSG101. E) Western blot for TSG101, signal is indicated by an asterix. Full-length blots are presented in Supplementary figure 1. F) Spectrum for calnexin. G) DLS data showing the relative amount of particles at each size and the cumulative undersize distribution of sEVs from each parent cell line. H) SEM imaging of sEVs from each parent cell line. The scale bar represents 100 nm. HI Size distribution of sEVs present in 130 k fraction after DUC determined using ImageJ (n=150 (NP), n=157 (AF), n=130 (FAT), n=216 (fetal NP)). 2k: DUC fraction containing debris and larger EVs; 10k: DUC fraction containing larger EVs; 130k pre: DUC sEV fraction before PBS wash; 130K post: DUC sEV fraction after PBS wash; AF: Annulus fibrosus; CD63: Cell surface protein of the tetraspanin family; CNX: Calnexin; DLS: Dynamic light scattering; EVs: Extracellular vesicles; FAT: Adipose tissue; FAT: Adipose tissue; FBS: sEV fraction of heat inactivated fetal bovine serum used for cell culture prior to exosome harvest serving as sEV positive control. M: New England Biolabs (NEB) marker; MS: Mass spectrometry; NP: Nucleus pulposus; SEM: Scanning electron microscopy; TSG101: Tumor susceptibility gene 101; VB: Vertebral body.

### Senescence Associated (SA)-β Galactosidase Staining

Quality of parent cells was assessed via SA-β-Galactosidase staining at the time of sEV harvest and only deemed suitable if none or few (<10) senescent cells were detected as identified by blue staining. Staining was carried out at pH 6 with a staining solution of 1 mg/mL 5-bromo-4-chloro-3-indolyl-beta-D-galacto-pyranoside (X-gal), 5 mM potassium ferricyanide, 5 mM potassium ferrocyanide, 150 mM NaCl and 2 mM MgCl_2_ in 40 mM citric acid/sodium phosphate buffer at pH 6 (119,122).

### Western Blot (WB) Analysis

Protein samples were separated on a 12% SDS acrylamide gel (Biorad Laboratories) and transferred to a nitrocellulose membrane (Biorad Laboratories) at 90 volts for 120 min in 1 × transfer buffer. Membrane blocking was done for 2 h at room temperature, followed by overnight incubation with a primary antibody at 4°C. The membrane was washed 3 × with 1 × Tris buffered saline with Tween 20 (TBST), reblocked for 2 h, and incubated for 2 h at 4°C with the goat anti-rabbit IgG, cross-adsorbed, and horseradish peroxidase (HRP) conjugated secondary antibody (1:5000, Thermofisher Scientific). Primary antibodies used in this bovine study included CD63 (rabbit, polyclonal (1:1000, Proteintech)), TSG101 (rabbit, polyclonal (1:1000, Thermofisher Scientific)), and Calnexin (rabbit, polyclonal (1:1000, Thermofisher Scientific). SuperBlock^TM^ PBS Blocking Buffer was used for all blocking and antibody dilutions (Thermofisher Scientific). The chromogenic 1-Step^TM^ TMB-Blotting Substrate (ThermoFisher Scientific) was used to visualize protein bands.

### Dynamic Light Scattering (DLS)

Particle analysis was performed by estimating the hydrodynamic diameter of particles present in the different fractions with a Nanobrook 90Plus (Brookhaven Instruments). Samples for DLS were prepared as a suspension. 25 µl of each sample were resuspended well and diluted 1:80 in 1 × PBS before triplicate measurements. After stabilization of laser and temperature of the device, the following parameters were set for reproducibility and standardization: Laser Wavelength: 659 nm, dust cut-off: 40, angle: 90°, temperature: 25°C. Each DLS measurement consisted of 10 acquisitions for 5 sec each. BIC Particle Sizing software from Brookhaven Instruments was used for data acquisition and analysis. Particle size distribution data were presented graphically and numerically, showing the relative particle amount at each size and the cumulative undersize distribution.

### Scanning Electron Microscopy (SEM)

Small EVs were fixed in 2% paraformaldehyde (PFA), applied to a Formvar coated 400 mesh copper grid (EMS), post fixed with 1% glutaraldehyde, stained for 2 min with UranyLess (EMS) at room temperature, rinsed with deionized water, and imaged on a JEOL JSM 7900F-LV FESEM at 30kV in with a specialty STEM-in-SEM holder. Two technical replicates from one or two biological replicates were generated depending on the cell line.

### Database Generation

An extensive library of bovine sEV proteins was generated by integrating data from UniProt (123) and ExoCarta (124). The ExoCarta proteins were renamed with the UniProt ID mapping tool for subsequent analysis. To identify putative bovine sEV proteins, human sEV protein sequences were searched against bovine protein sequences obtained from UniProt using the Basic Local Alignment Search Tool (BLAST) (125). The BLAST results were merged with the bovine sEV protein sequences, and all sequences were clustered using CD-HIT v4.6.8 with a sequence identity threshold of 0.95 and a minimum length of 100 (126). For the explorative bovine sEV proteome profile study, we also used a comprehensive umbilical cord MSC (UCMSC) sEV proteome database (127).

### Small EVs protein analysis by liquid chromatography-tandem mass spectrometry (LC-MS/MS)

To deplete high-abundance proteins, extracted sEV proteins were processed with the Pierce albumin and IgG removal kit (P/N 89875 ThermoFisher Scientific) according to the manufacturer’s protocol, which allows the analysis of less abundant proteins. In brief, after the gel slurry was washed, the sample was loaded into the gel slurry and incubated with the orbital rotator for 10 min at room temperature. Then, the flow-through was collected at 10,000 × *g* for 1 min using the spin column. The elution was repeated with the binding and elution process using 75 µl of the binding/wash buffer to obtain the non-redundant proteins.

Depleted sEV proteins were dissolved in protein buffer (8 M Urea, 0.1% SDS, 25 mM Triethylammonium Bicarbonate, pH 8.0) and quantified following a previous method (128). Protein assays were performed to quantify purified proteins by the EZQ™ Protein Quantification Kit (Thermo Fisher Scientific, San Jose, CA, USA) with the SoftMax Pro Software v5.3 under the SpectraMax M5 (Molecular Devices, LLC). For each sample, a total of 10 μg of protein was reduced with 40 mM DTT, alkylated with 100 mM 2-chloroiodoacetamide, and trypsin-digested (at an enzyme-to-protein ratio (w/w) of 1:100). Tryptic digested peptides were desalted with C18-solid phase extraction (The Nest Group, Inc., Southborough, MA) for the capture of polar and non-polar peptides. Briefly, after equilibrating the cartilage with 1 ml of acetonitrile and 2 ml of water in 0.1% TFA sequentially. The peptide samples were passed over the columns three times before they were washed with 0.1% TFA. The peptides retained on the Sep-Pak C-18 column were eluted in 1 ml of 80% acetonitrile in 0.1% TFA, and the eluant was in the lyophilization.

The hybrid trapped ion mobility-quadrupole time-of-flight mass spectrometer (timsTOF fleX, Bruker Daltonics, Bremen, Germany) with a modified nano-electrospray ion source was interfaced with an ultra-performance EvoSep One LC system (Evosep Biosystem, Odense, Denmark). Briefly, Evotip was wetted with 100 µl of 100% iso-propanol, rinsed with 20 µl of Solvent B (99.9% acetonitrile and 0.1% formic acid (v/v)), equilibrated with 20 ul of 0.1% formic acid (v/v), loaded with 200 ng of digested peptides, and subsequently washed with 20 ul of 0.1% formic acid (v/v) using centrifugal force at 700 x *g* for 1 min. 100 μl of 0.1% formic acid were added to Evotip to prevent drying. Samples were injected into the Bruker timsTOF fleX MS coupled with the Evosep One instrument (Evosep Biosystems). The standard preset method of 15 SPD was used with the EV1106 Endurance column (Evosep One Biosystem) with 200 ng injection. The spectrum library was produced in the data-dependent mode with Parallel Accumulation Serial Fragmentation (PASEF) to improve ion utilization efficiency and data acquisition speed. The dual TIMS operated the system at 100% duty cycle and recorded the MS/MS mode scanning from 100 to 1700 m/z. The ion mobility was scanned from 0.6 to 1.6□Vs/cm^−2^, and TIMS ion charge control was set to 5e6. The TIMS dimension was calibrated linearly using three selected ions from the Hexakis (1H, 1H, 2H-difluoroethoxy) phosphazene, Hexakis (1H, 1H, 3H-tetrafluoropropoxy) phosphazene Agilent ESI LC/MS tuning mix [m/z, 1/K0: (622.0289, 0.9915□Vs□cm^−2^), (922.0097, 1.1996□Vs□cm^−2^), (1221,9906, 1.3934□Vs□cm^−2^)] in positive mode.

### Data Searching, Identification, and Quantification

The MS/MS data were extracted peak lists under Data Analysis (Bruker Daltonics, Bremen, Germany; version 6.1) and analyzed with Mascot (Matrix Science, London, UK; version 2.7). Mascot was set up to search against the customized protein database described above (customized 5,922 contigs) with a decoy database for false discovery rate (FDR) using digestion enzyme trypsin/Lys-C, parental ion tolerance of 15 ppm and fragment ion mass tolerance of 0.5 Da, respectively. Carbamidomethyl of cysteine (+57.021 Da) was set as the static modification, and oxidation of methionine (+15.995 Da), deamidation of glutamine and asparagine (+0.984 Da), pyro-glutamine formation from N-terminal glutamine (-17.026 Da), as well as phosphorylation of serine, threonine, and tyrosine (+79.966 Da) were specified as the variable modifications. Scaffold Q+S (Proteome Software Inc., Portand, OR, USA; version 5.4.2) was used to validate MS/MS based peptide and protein identifications. Peptide identifications were accepted if they could be established at greater than 95.0% probability by the Peptide Prophet algorithm. Protein quantification was accepted if they could be established at greater than 95.0% probability at significance with at least three peptides with 99.9%. Differences in protein abundance between NP cells and NP sEVs were evaluated using Student’s *t*-test. To be identified as being significantly differentially abundant, proteins should be quantified with at least four unique peptides in both experimental replicates with a *p* < 0.05 and a fold change >1.5 or <0.5.

### Data Availability

The proteome data that were newly generated have been deposited in the ProteomeXchange and MassIVE partner repositories with the data set identifiers PXD056784 and MSV000096081. RNA sequencing data relevant to the cell lines used here was published previously (129) and can be found in NCBI GEO (https://www.ncbi.nlm.nih.gov/geo/), accession number GSE216377. All additional files are included in the manuscript.

### Gene Ontology Analysis, Network and Clustering of Small EV Proteins

The gene ontology (GO) and functional enrichment analysis of differentially abundant NP sEV proteins, IVD sEV proteins (NP, AF, and fetal NP), and FAT sEV proteins was performed using the Bioinformatics Resources 6.8 Database for Annotation, Visualization, and Integrated Discovery (DAVID) and the number of genes associated with each term, and enrichment bubble plots were generated using SRplot (130). Alternatively, ToppFun of the ToppGene Suite was utilized (FDR correction and p-value cut-off 0.05) (131). Using the search tool for the retrieval of interacting genes/proteins (STRING) database (132), a protein network was built for overrepresented and all NP sEV proteins. For medium confidence network construction, default settings were employed. To find clusters in the created protein network, the Markov clustering algorithm (MCL) was utilized and carried out with an inflation parameter of 3. Following clustering, biological processes (BP), molecular functions (MF), and cellular components (CC) clusters were examined in the context of NP sEV protein function. Venny 2.1 (https://bioinfogp.cnb.csic.es/tools/venny/index.html) was used for comparison.

## RESULTS

### 3.1. Small EV Isolation

Small EVs were confirmed in the final DUC fraction for all primary bovine parent cell lines through the presence of established sEV markers CD63 and TSG101 and the absence of the cellular marker Calnexin via spectral counts and by Western blot, if sufficient material was available, as shown exemplarily for NP parent cells (Figures 2B-F, Supplementary figure 1). The expected particle size range for sEVs (50-150 nm) was confirmed by DLS and ImageJ analysis of SEM micrographs (Figures 2G to 2I). The protein concentration in each sEV fraction was calculated per cultured parent cell (Supplementary table 1).

### 3.2 Bovine Small EV Protein Database

A bovine sEV protein database was constructed by integrating sEV proteins from ExoCarta (124) and bovine proteins from UniProt (123). A total of 47,128 bovine protein sequences was downloaded from UniProt, while ExoCarta provided 1,416 bovine and 6,514 human sEV proteins as of October 2, 2023. The BLAST results of human sEV proteins against 47,128 bovine protein sequences resulted in 4,972 bovine proteins, which were merged with bovine sEV proteins obtained from ExoCarta. After sequence clustering with 1,416 bovine sEV proteins and 4,972 bovine proteins homologous to human sEV proteins, the final bovine sEV protein database BOVXCU24 had 5,922 unique entries. This database represents the most extensive bovine sEV protein database to date.

### 3.3 Proteome Profiling of Bovine Small EVs

For profiling purposes sEV proteins were first identified in a non-quantitative manner. Small EVs harvested from different tissues of fetal or adult primary bovine parent cells were compared to existing sEV and MSC sEV data.

#### 3.3.1. Bovine Small EV Proteins in the Context of Existing Data

Our bovine sEV data was essentially consistent with findings by Kugeratski *et al.* identifying Syntenin 1 alongside GTPases and membrane proteins such as an integrin subunits and the “classic” tetraspanin sEV markers CD9, CD63 and CD81 (133). Additionally, established sEV markers such as the programmed cell death 6 interacting protein (PDCD6IP or ALIX) (134) and the tumor susceptibility marker TSG101 known for its role in vacuolar sorting and sEV biogenesis (135) were also detected. ExoCarta (124) and Vesiclepedia (136) are evolving databases for (s)EV research. Of the 486 bovine sEV proteins identified in this study seven proteins were not listed in either database at the time (Figure 3, Table 1, Supplementary table 2) despite cross-referencing available information with GeneCards (https://www.genecards.org) and other search engines to identify aliases. Those seven proteins were ATPase H+ Transporting V0 Subunit E2 (ATP6V0E2), H2A Histone Family Member B2 (H2AB2), Mannose binding lectin 2 (MBL), Asparaginyl-tRNA synthetase 1 (NARS1), the actin binding lymphocyte cytosolic protein 1 (PLS2), Arginyl-tRNA synthetase 1 (RARS1) and Stomatin-like 1 (STOML1), however these might be variants or isoforms of listed proteins or added recently. The 479 bovine sEV proteins were associated with a range of biological pathways through functional enrichment analysis in DAVID (Figure 3B) and ToppFun (Figure 3C). Among those, based on fold enrichment over background, the top five highest ranking pathways by fold change (FC) in the KEGG database were proteasome (bta03050), pentose phosphate pathway (PPP) (bta00030), glycolysis/gluconeogenesis (bta00010), complement and coagulation cascades (bta04610), and ECM receptor interaction (bta04512) (Figure 3B). Analysis in ToppFun identified the (innate) immune system (Reactome MM14661), neutrophil degeneration (Reactome M27620), hemostasis (Reactome M8395), nervous system development (M29853), integrin1 (Pathway Interaction Database (PID) M18), and ephrin signaling (Reactome, M27201) amongst the top 200 pathways (Figure 3C and Supplementary table 3).

**Figure 3:**
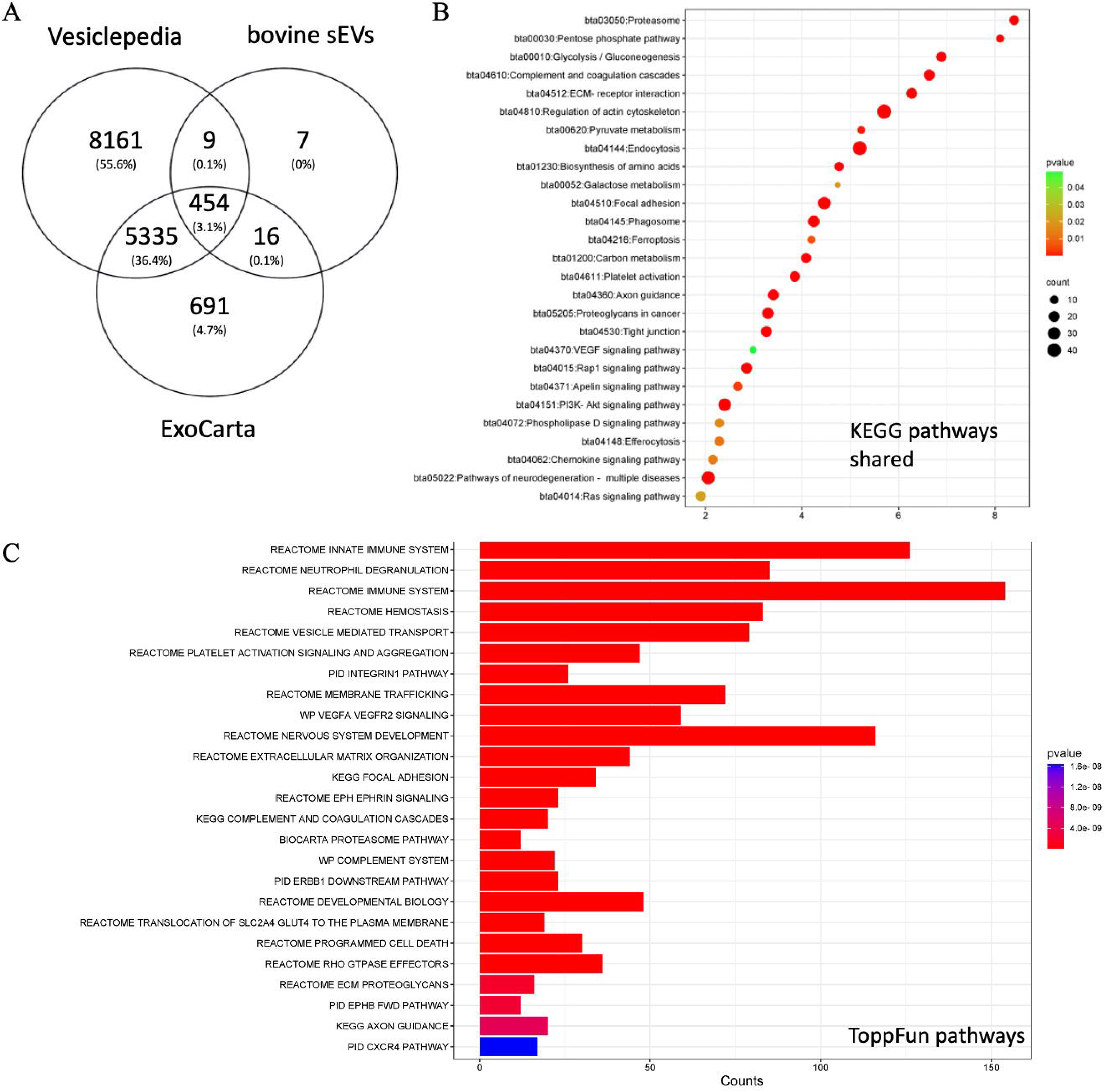
Bovine sEVs and existing data. A) Venn diagram illustrating the comparison of bovine sEV proteins with the ExoCarta and Vesiclepedia databases. B) KEGG pathway analysis in DAVID and (C) ToppFun pathway analysis of 479 shared proteins. DAVID: Database for annotation, visualization, and integrated discovery; KEGG: Kyoto encyclopedia of genes and genomes.

**Table 1:** DAVID based functional enrichment analysis comparing bovine sEV proteins with the ExoCarta database. DAVID: Database for annotation, visualization, and integrated discovery GO: Gene ontology; bta: Bos taurus; ECM: Extracellular matrix; KEGG: Kyoto encyclopedia of genes and genomes.

| Data type | Category | KEGG Pathway | Pathway Identifier | Gene Symbol |
| --- | --- | --- | --- | --- |
| 470 shared proteins of bovine sEV derived primary IVD and FAT parent cells and the ExoCarta database | Metabolism | Glycolysis / Gluconeogenesis<br>Carbon metabolism<br>Biosynthesis of amino acids<br>Pentose phosphate pathway | bta00010<br>bta01200<br>bta01230<br>bta00030 | ACO1, ADH5, AKR1A1, ALDOA, ALDOC, ALDH3B1, ALDH9A1, G6PD, GAPDH, GPI, IDH1, LDHA, LDHB, PGD, PFKL, PHGDH, PKLR, PKM, PGM1, PSAT1, TKT, TPI1 |
|  | Genetic Information Processing | Proteasome | bta03050 | PSMA1, PSMA2, PSMA3, PSMA4, PSMA5, PSMB1, PSMB2, PSMB3, PSMB4, PSMB5, PSMB6, PSMC6, PSMD8, PSMD12 |
|  | Environmental Information Processing | PI3K-Akt signaling pathway<br>Rap1 signaling pathway<br>ECM-receptor interaction | bta04151<br>bta04015<br>bta04512 | ABI1, CDC42, COL1A1, COL2A1, COL6A1, COL6A2, COL6A3, EPHA2, FN1, GNAI1, GNAI2, GNAI3, GNAQ, GNB1, GNG5, GNG12, HRAS, HSPG2, ITGA1, ITGA3, ITGA5, ITGA6, ITGAV, ITGB1, ITGB2, ITGB5, ITGB7, KRAS, LAMA5, LAMB1, LAMC1, NRAS, PFN1, PFN2, RAC1, RALA, RAP1B, RHEB, RHOA, RRAS, RRAS2, THBS1, TLN1, VASP, YWHAQ, YWHAZ |
|  | Cellular Processes | Endocytosis<br>Focal adhesion<br>Phagosome<br>Tight junction<br>Regulation of actin cytoskeleton<br>Cytoskeleton in muscle cells | bta04144<br>bta04510<br>bta04145<br>bta04530<br>bta04810<br>bta04820 | ACTN1, ACTR2, ACTR3, AP2M1, ARF1, ARPC2, ARPC4, ARPC5, ARPC5L, ATP1B1, ATP6AP1, ATP6V0A1, ATP6V1B2, ATP6V1D, BGN, C3, |
|  |  |  |  | C5, C6, C7, C8A, C8G,<br>CAPN2, CAPZA2,<br>CAPZB, CDC42,<br>CHMP1B, CHMP2A,<br>CHMP4B, CHMP6,<br>COL1A1, COL2A1,<br>COL3A1, COL5A2,<br>COL6A1, COL6A2,<br>COL6A3, CSRP1,<br>C8A, DNM2, DSC2,<br>EEA1, EHD2, EHD4,<br>F2, FLNA, FLNB,<br>FMNL3, FN1, GNG12,<br>GSN, HRAS, HSPG2,<br>ILK, IQGAP1,<br>IQGAP2, IST1, ITGA1,<br>ITGA3, ITGA5,<br>ITGA6, ITGAV,<br>ITGB1, ITGB2,<br>ITGB5, ITGB7, IGF2R,<br>KRAS, LAMA5,<br>LAMC1, LAMP1,<br>LAMP2, MSN, M6PR,<br>MYH9, MYH14,<br>NCKAP1, NID1, NID2,<br>NRAS, PDCD6IP,<br>PFN1, PFN2, RAB5B,<br>RAB5C, RAB7A,<br>RAB8A, RAB8B,<br>RAB10, RAB13,<br>RAB22A, RAB35,<br>RAC1, RAP1B, RDX,<br>RRAS, RRAS2,<br>SMURF1, SNX3,<br>SNX12, SPTBN1,<br>TSG101, THBS1,<br>TLN1, TTN, TUBA1A,<br>TUBB2A, TUBB6,<br>VAMP3, VCL, VPS28,<br>VPS35, VPS36,<br>VPS37B, VPS4A,<br>VPS4B, VTA1, VASP |
|  | Human Diseases | Parkinson disease<br>Amyotrophic lateral<br>sclerosis<br>Prion disease<br>Bacterial invasion of<br>epithelial cells<br>Proteoglycans in cancer<br>Salmonella infection<br>Amoebiasis<br>Human papillomavirus<br>infection | bta05012<br>bta05014<br>bta05020<br>bta05100<br>bta05205<br>bta05132<br>bta05146<br>bta05165 | ACTN1, ACTR1A,<br>ACTR2, ACTR3,<br>ANXA7, ANXA11,<br>ARPC2, ARPC4,<br>ARPC5, ARPC5L,<br>ATP6AP1,<br>ATP6V0A1,<br>ATP6V1B2,<br>ATP6V1D, C1QA,<br>C1QC, C5, C6, C7,<br>C8A, C8G, C9,<br>CDC42, COL1A1,<br>COL2A1, COL3A1,<br>COL6A1, COL6A2, |
|  |  |  |  | COL6A3, DCTN2,<br>DDX5, DNM2, FN1,<br>GNAI1, GNAI2,<br>GNAI3, GNA11,<br>GNAQ, HRAS,<br>HSPA5, HSPA8,<br>HSPB1, HSPG2, ILK,<br>IQGAP1, ITGA1,<br>ITGA3, ITGA5,<br>ITGA6, ITGAV,<br>ITGB1, ITGB2,<br>ITGB5, ITGB7, IST1,<br>KRAS, LAMA5,<br>LAMB1, LAMC1,<br>LAMP1, LAMP2,<br>MSN, MMP2, MYH9,<br>NCKAP1, NRAS,<br>NOTCH2, NOTCH3,<br>PARK7, PDCD6IP,<br>PFN1, PFN2, PKM,<br>PLAU, PSMA1,<br>PSMA2, PSMA3,<br>PSMA4, PSMA5,<br>PSMC6, PSMD8,<br>PSMD12, PSMB1,<br>PSMB2, PSMB3,<br>PSMB4, PSMB5,<br>PSMB6, RAB5B,<br>RAB5C, RAB7A,<br>RAB8A, RAC1, RDX,<br>RHOG, RHEB, RRAS,<br>RRAS2, SOD1,<br>SEC13, SMURF1,<br>SNX3, SPTBN1,<br>THBS1, TUBA1A,<br>TUBB2A, TUBB6,<br>UBA1, VASP, VCL,<br>VDAC1, VCP, VPS28,<br>VPS35, VPS36,<br>VPS37B |
|  | Organismal Systems | Axon guidance<br>Complement and<br>coagulation cascades<br>Platelet activation | bta04360<br>bta04610<br>bta04611 | C1QA, C1QC, C3, C5,<br>C6, C7, C8A, C8G, C9,<br>CDC42, CDK5, CLU,<br>COL1A1, COL3A1,<br>EPHA2, EPHA3,<br>EPHA5, EPHB1,<br>EPHB2, EPHB3,<br>EPHB4, F10, F2, F5,<br>FGA, FGB, FGG,<br>GNAI1, GNAI2,<br>GNAI3, GNAQ,<br>HRAS, ILK, ITGB1,<br>ITGB2, KRAS,<br>MASP1, NRAS,<br>PLAU, PLXNB2,<br>RAC1, RAP1B,<br>RHOA, RRAS,<br>SERPIND1,<br>SERPINF2, TLN1,<br>VAMP8, VASP |
| 16 bovine<br>exosomes<br>derived<br>primary IVD<br>and FAT<br>parent cells | No specific<br>association |  |  |  |

#### 3.3.2. Small EV Proteins from Autologous Parent Cells of Different Tissues

First, proteome profiling of autologous sEVs isolated from bovine IVD (NP and AF) and adipose (FAT) cells was conducted in a non-quantitative manner to identify NP parent sEV unique protein signatures. Of the 335 sEV proteins identified for AF, NP and FAT parent cells, a core of 102 (30%) were shared. Functional enrichment analysis in DAVID associated those proteins with a range of important pathways (Figure 4). An additional 156 sEV proteins were only identified for NP parent cells. Based on their pathway association in DAVID a metabolic theme stood out, especially the PPP (bta00030) showed a high FC and significance. Associations relevant to NP cell function such as L-ascorbic acid binding (GO:0031418), angiostatin binding (0043532), and general protein stabilization (GO:0050821) were also noted amongst many others (Table 2). Functional enrichment analysis in DAVID of the additional 24 sEV proteins shared only between NP and AF parent cells were involved in regulation of the cytoskeleton (bta04810), focal adhesion (bta04510) and endocytosis (bta04144), while additional 43 sEV proteins common only to NP and adipose (FAT) parent cells associated with functions and pathways of the 20S proteasome (KEGG, bta03050). Other sEV proteins in this group were involved with the citrate cycle (bta00020) or calcium ion binding (Figure 4, Table2 Supplementary figure 2). For analysis in ToppFun we first identified the pathways associated with sEV proteins for each of the three autologous parent cell lines and then determined overlaps and unique contribution by sEVs from NP parent cells. A total of 39 pathways were common to all three sources, including but not limited to the (innate) immune system, nervous system development, and programmed cell death. Additional 24 were shared only between sEV proteins of NP and AF parent cells and 75 between NP and FAT parent cells (Figure 4D, Supplementary table 4). Among the 23 pathways uniquely associated with sEV proteins from NP parent cells were glycolysis and gluconeogenesis (KEGG, WP) and the PPP (KEGG) represented through glyeraldehyde-3-phosphate dehydrogenase (GAPDH), aldo-keto reductase family member A1 (AKR1A1), phosphofructokinase (PFKL), phosphoglucomutase 1 (PGM1), alcoholdehydrogenase 5 (ADH5), pyruvate kinase (PKLR, PKM), aldehydedehydrogenase 3 and 9 family members (ALDH3B1, ALDH9A1), fructose-bi-phosphate aldolases (ALDOA, ALDOC) glucose-6-phosphate isomerase (GPI), lactate dehydrogenases (LDHA, LDHB), triosephosphate isomerase 1 (TPI1), transketolase (TKT), and glucose-6-phosphate dehydrogenase (G6PD). Also identified was signaling by the Roundabout (ROBO) family (Reactome), transmembrane receptors involved in axon guidance and cell migration through the association with several proteasome 20S subunits, two ribosomal proteins, profilins (PFN1, PFN2), vasodilator stimulated phosphoprotein (VASP), cyclase associated actin cytoskeleton regulatory protein1 (CAP1), RAC family small GTPase 1 (RAC1), protein kinase cAMP-dependent type II regulatory subunit alpha (PRKAR2A), and cell division cycle 42 (CDC42) (Supplementary table 4).

**Figure 4:**
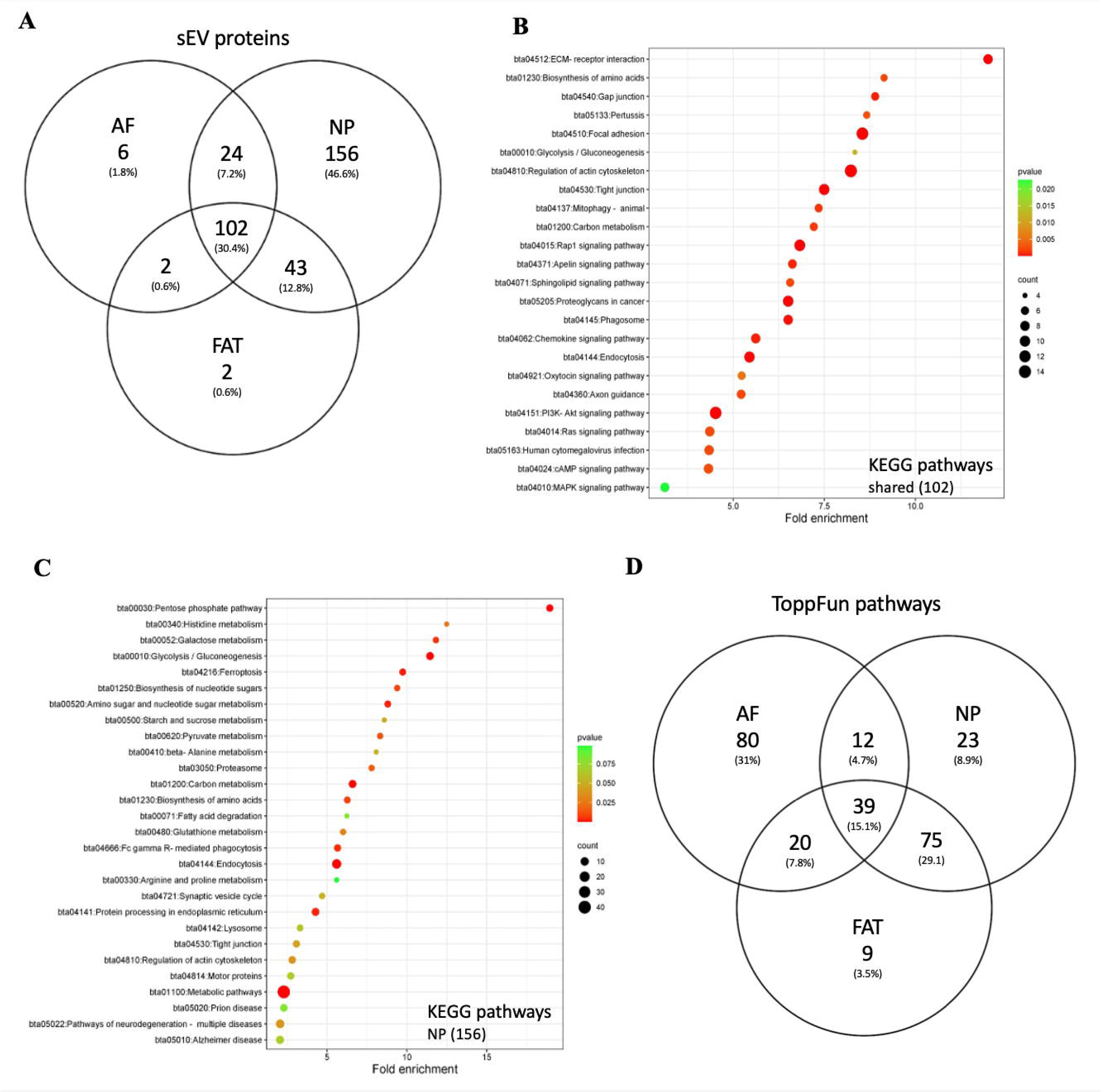
Autologous sEV profiling: A) Venn diagram comparing exosome proteins from autologous NP, AF, and FAT parent cells. B) KEGG pathway analysis in DAVID of 102 shared proteins and (C) KEGG pathway analysis in DAVID of 156 proteins unique to NP sEVs. D) ToppFun pathway analysis of top 200 shared pathways. AF: Annulus fibrosus; DAVID: Database for annotation, visualization, and integrated discovery; FAT: Adipose tissue; KEGG: Kyoto encyclopedia of genes and genomes; NP: Nucleus pulposus.

**Table 2:**
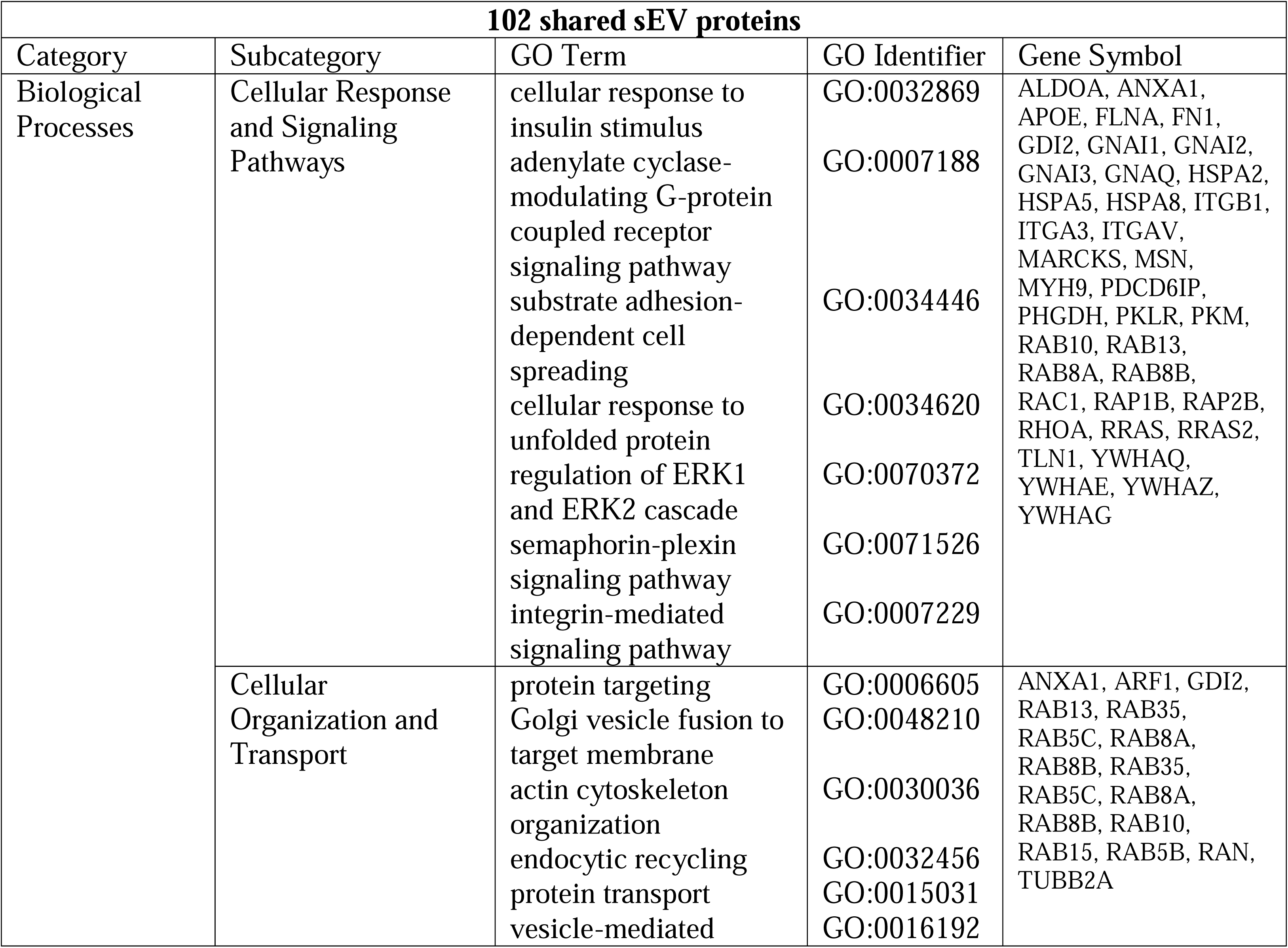
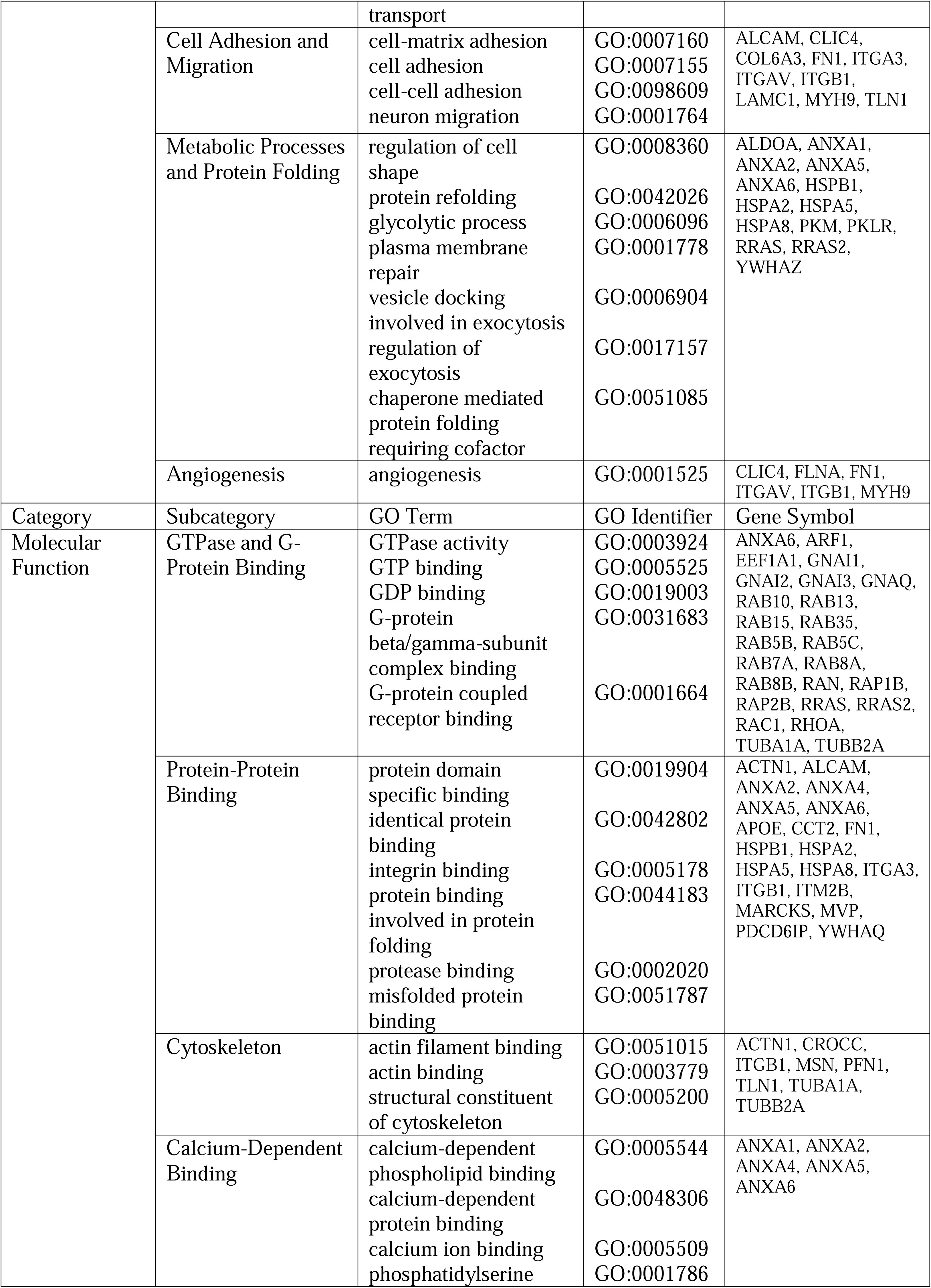
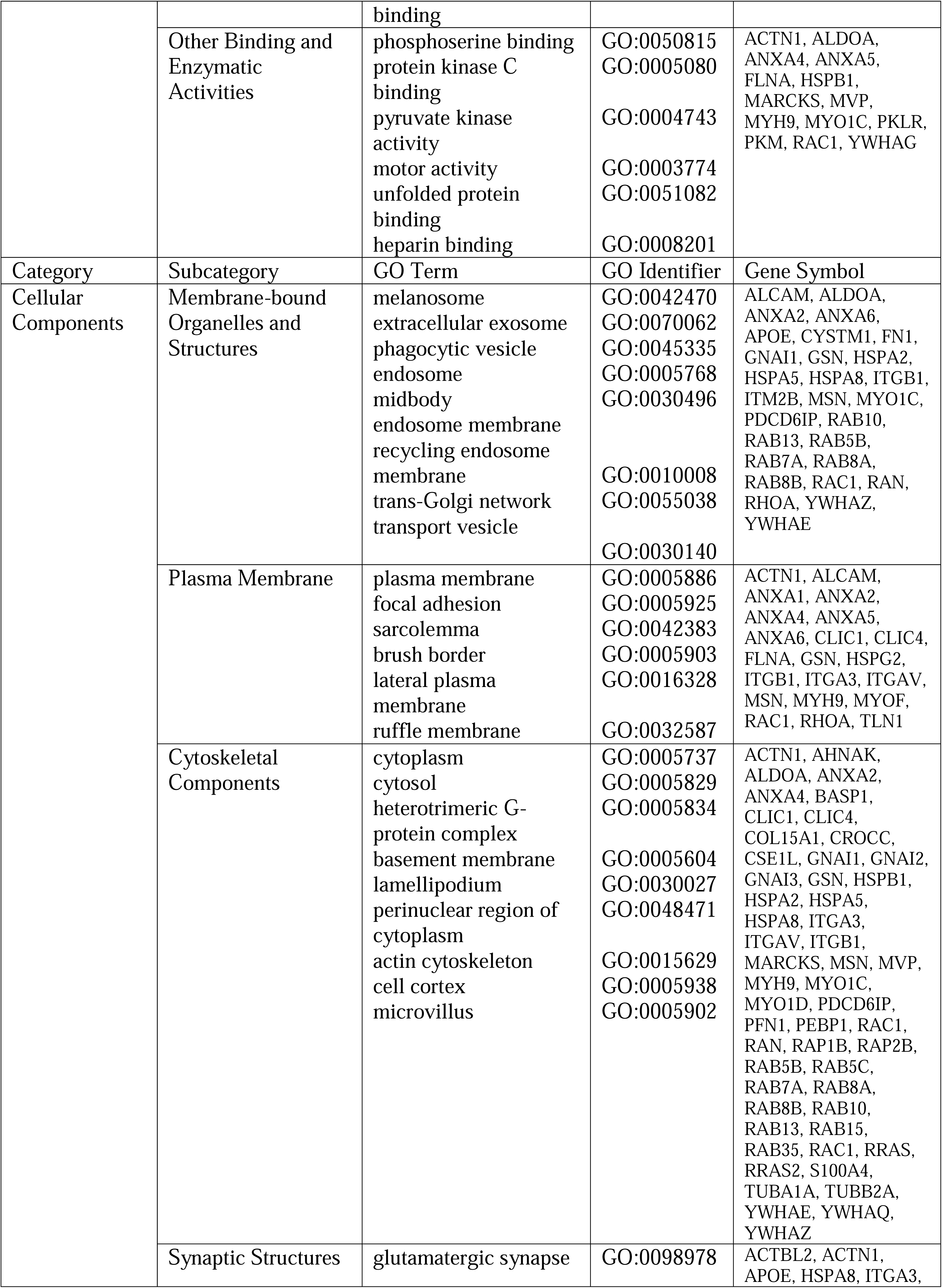
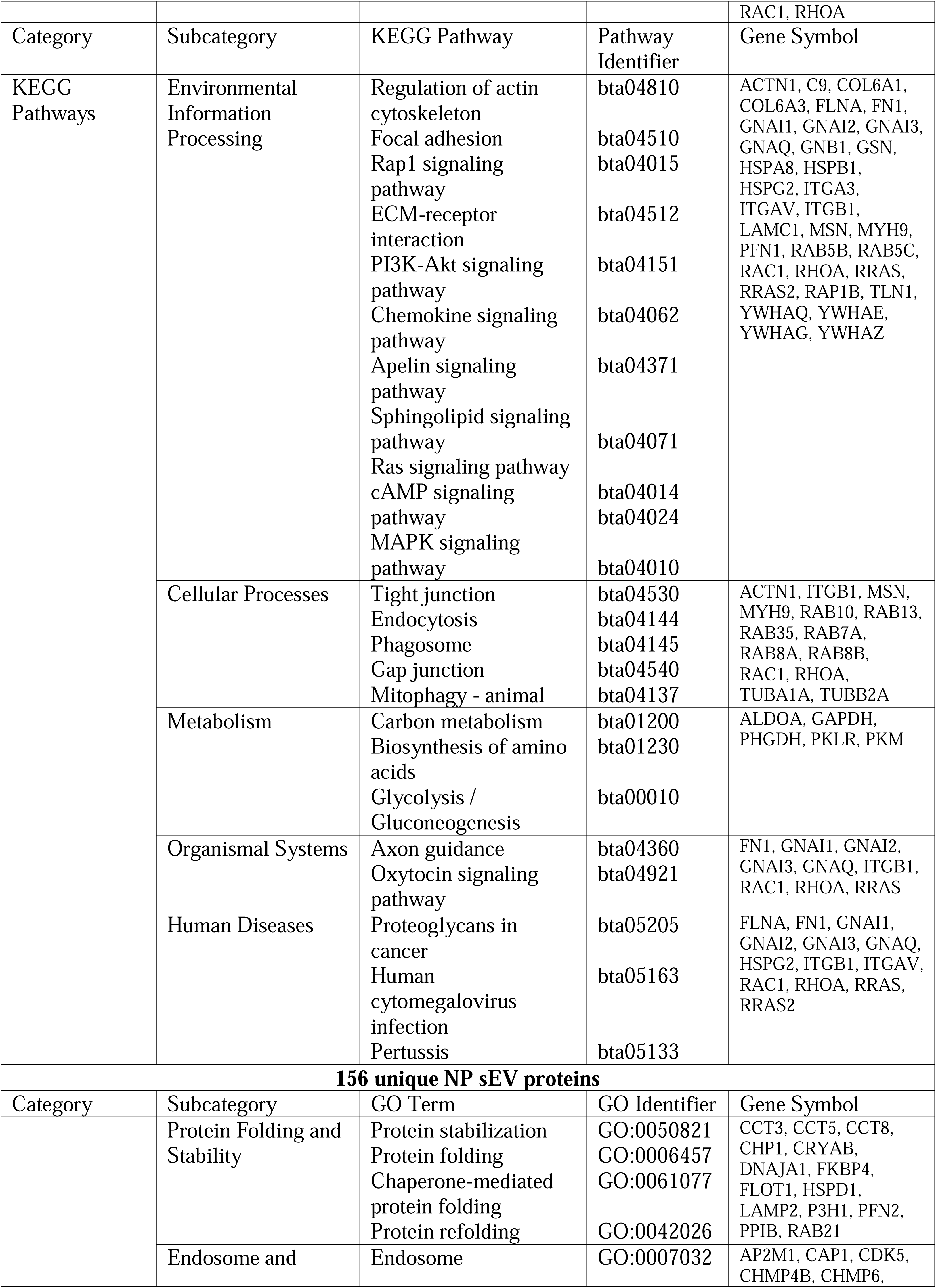
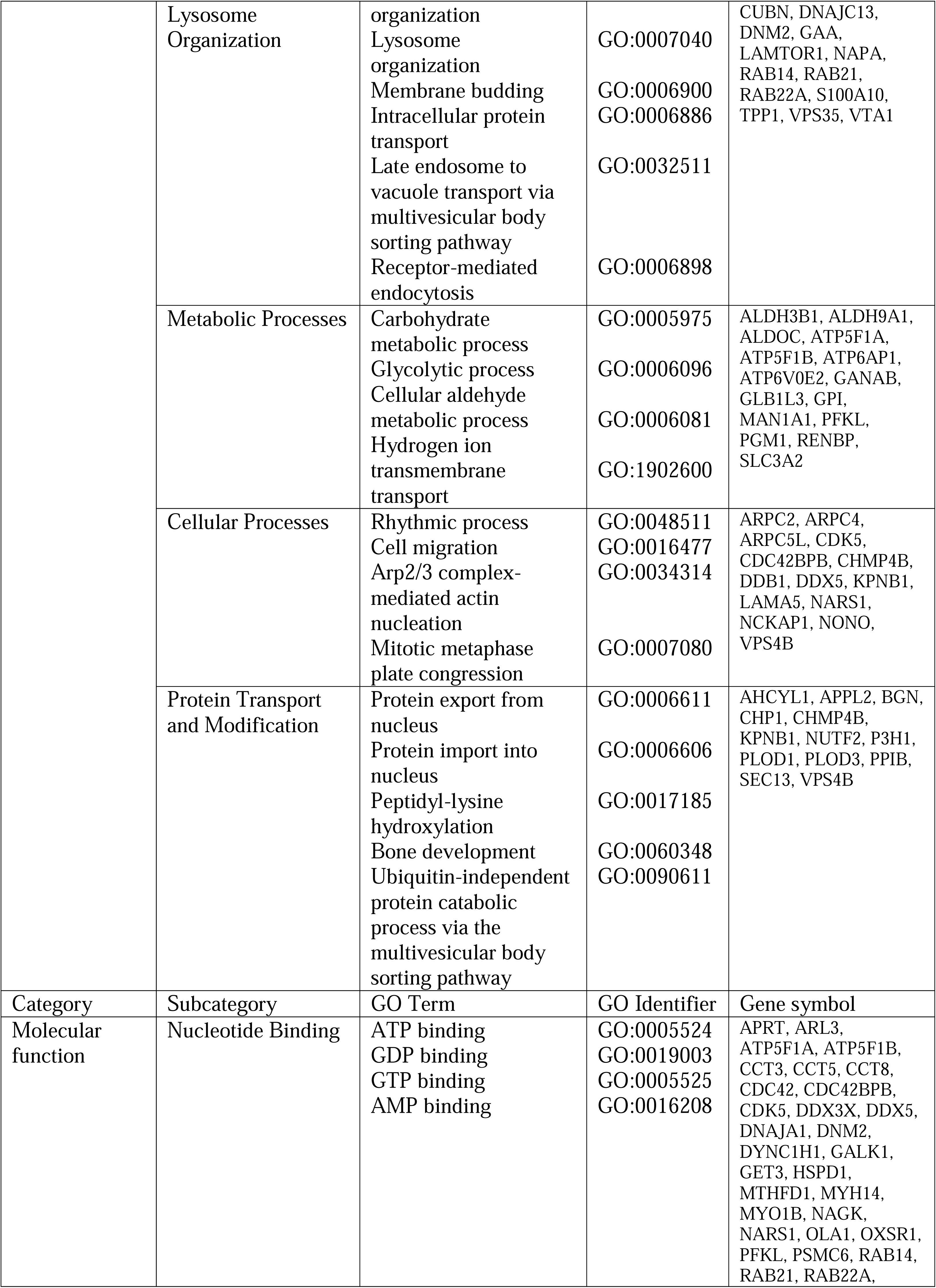
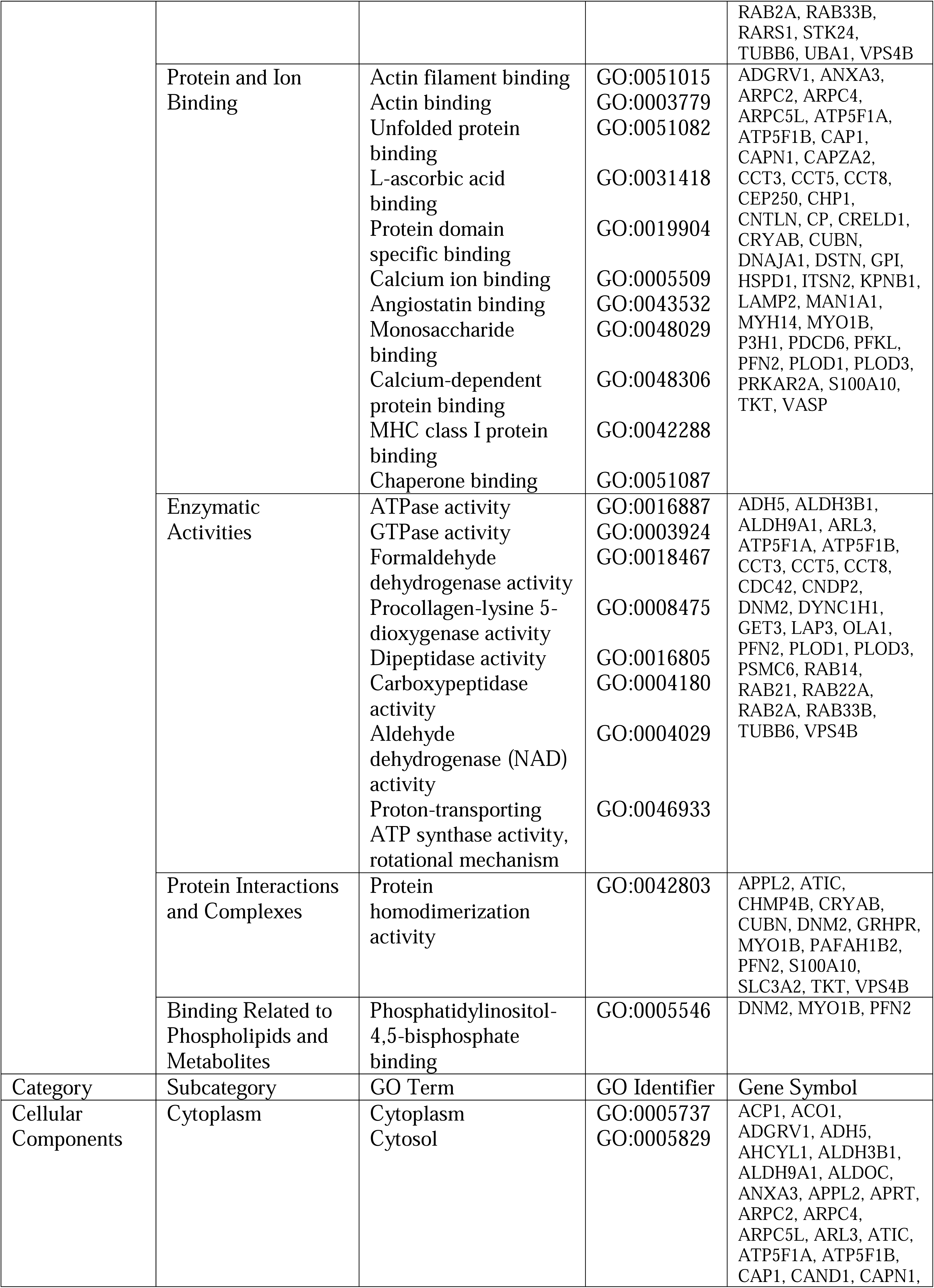
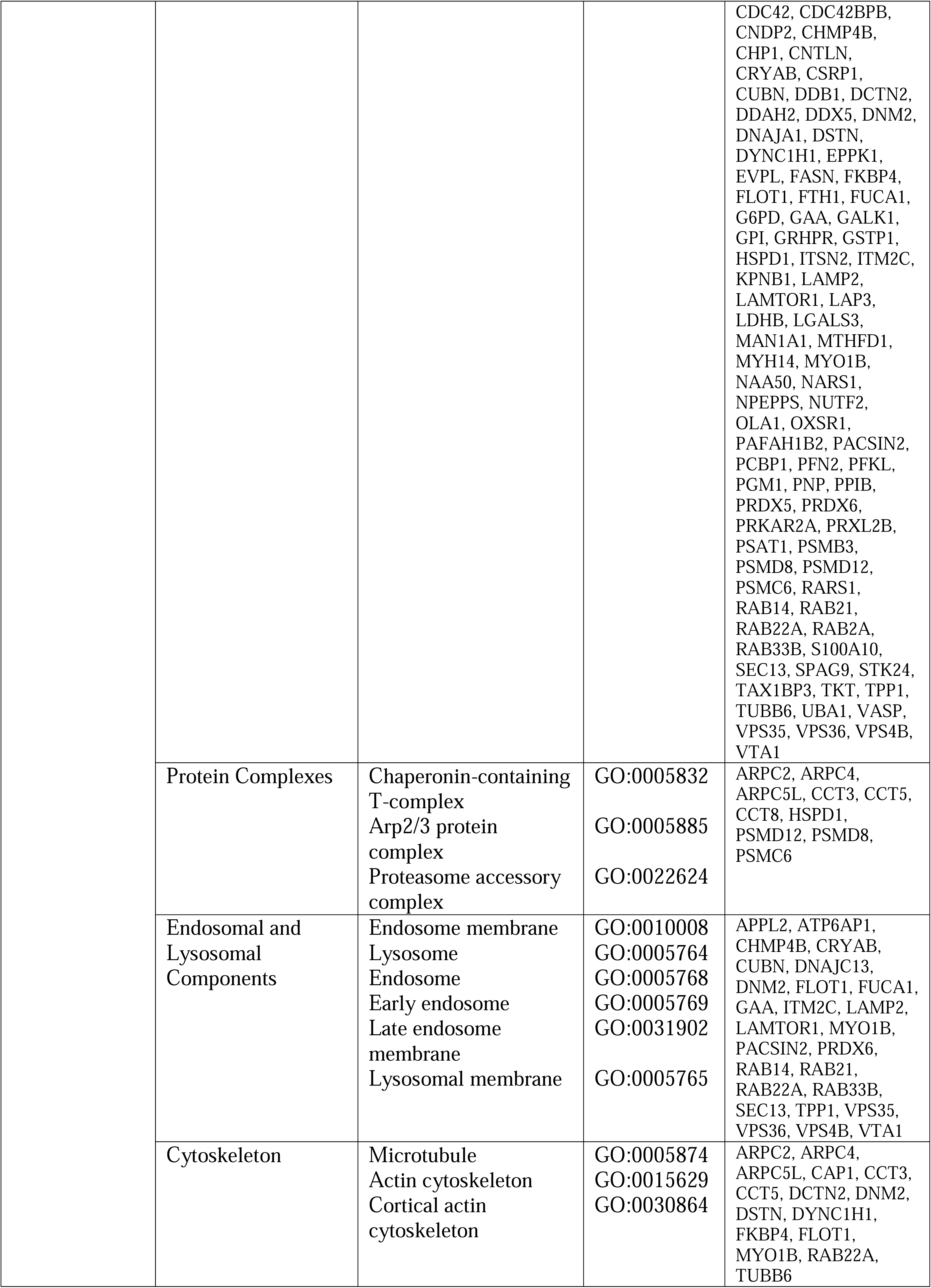
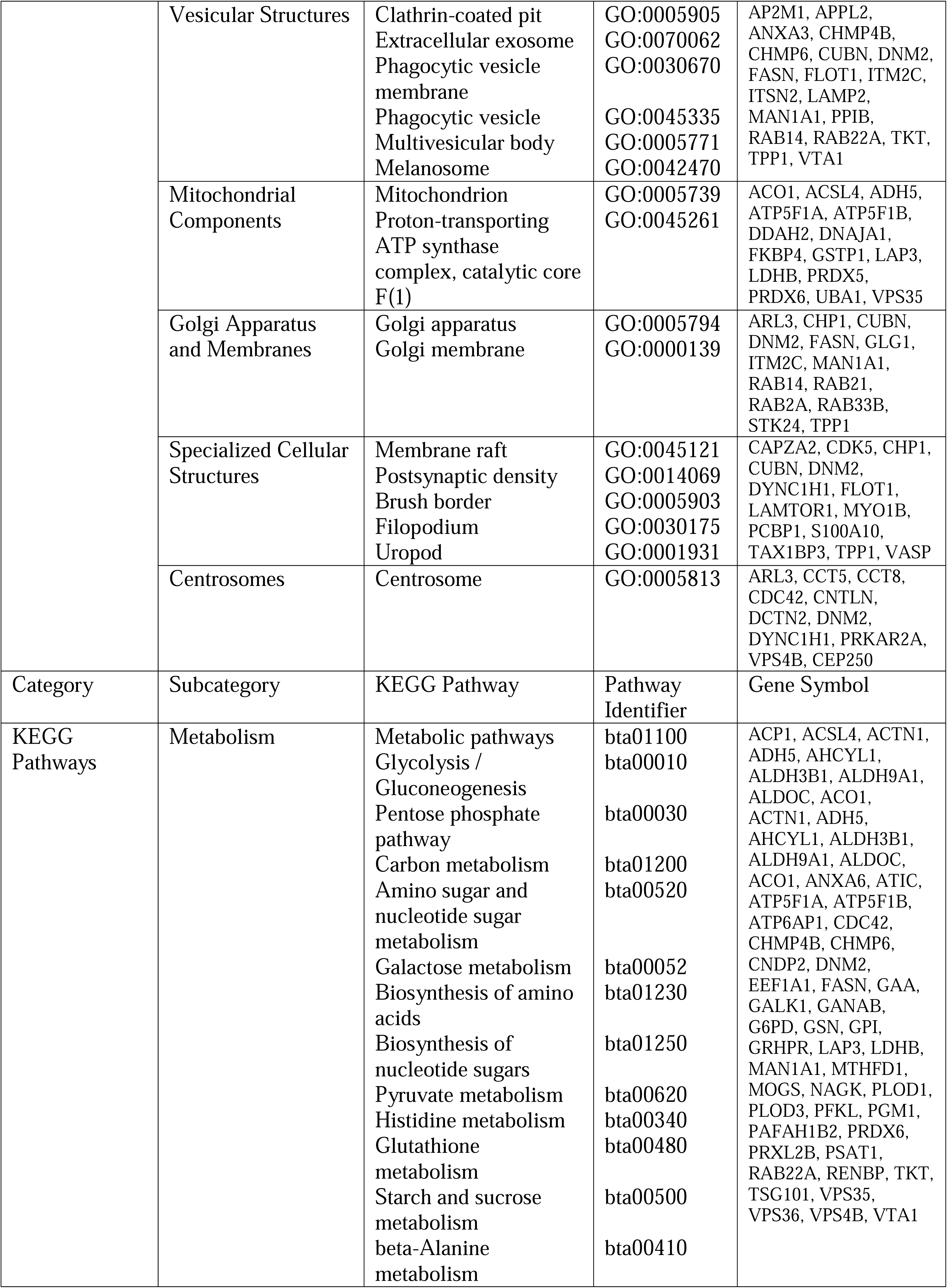
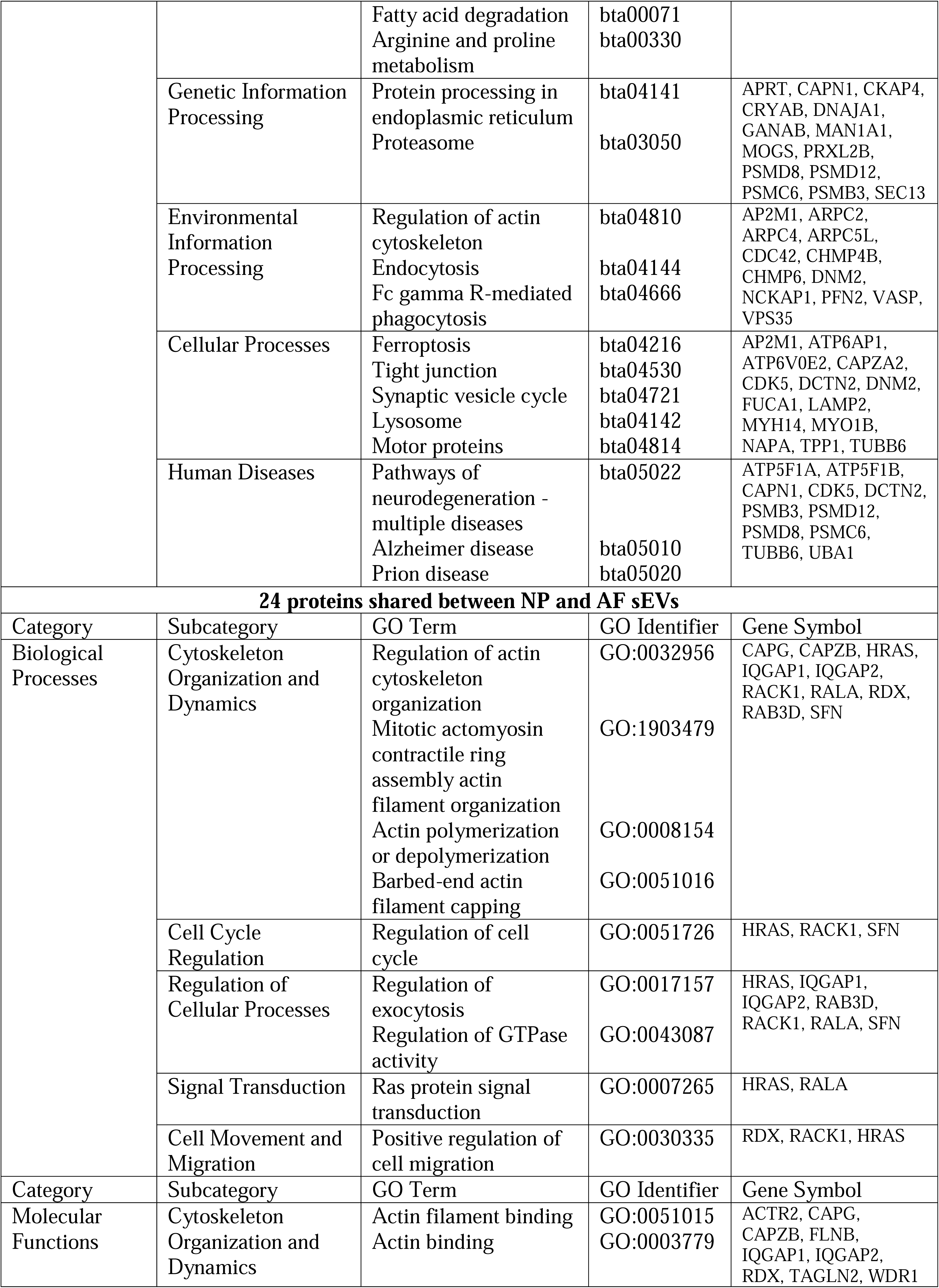
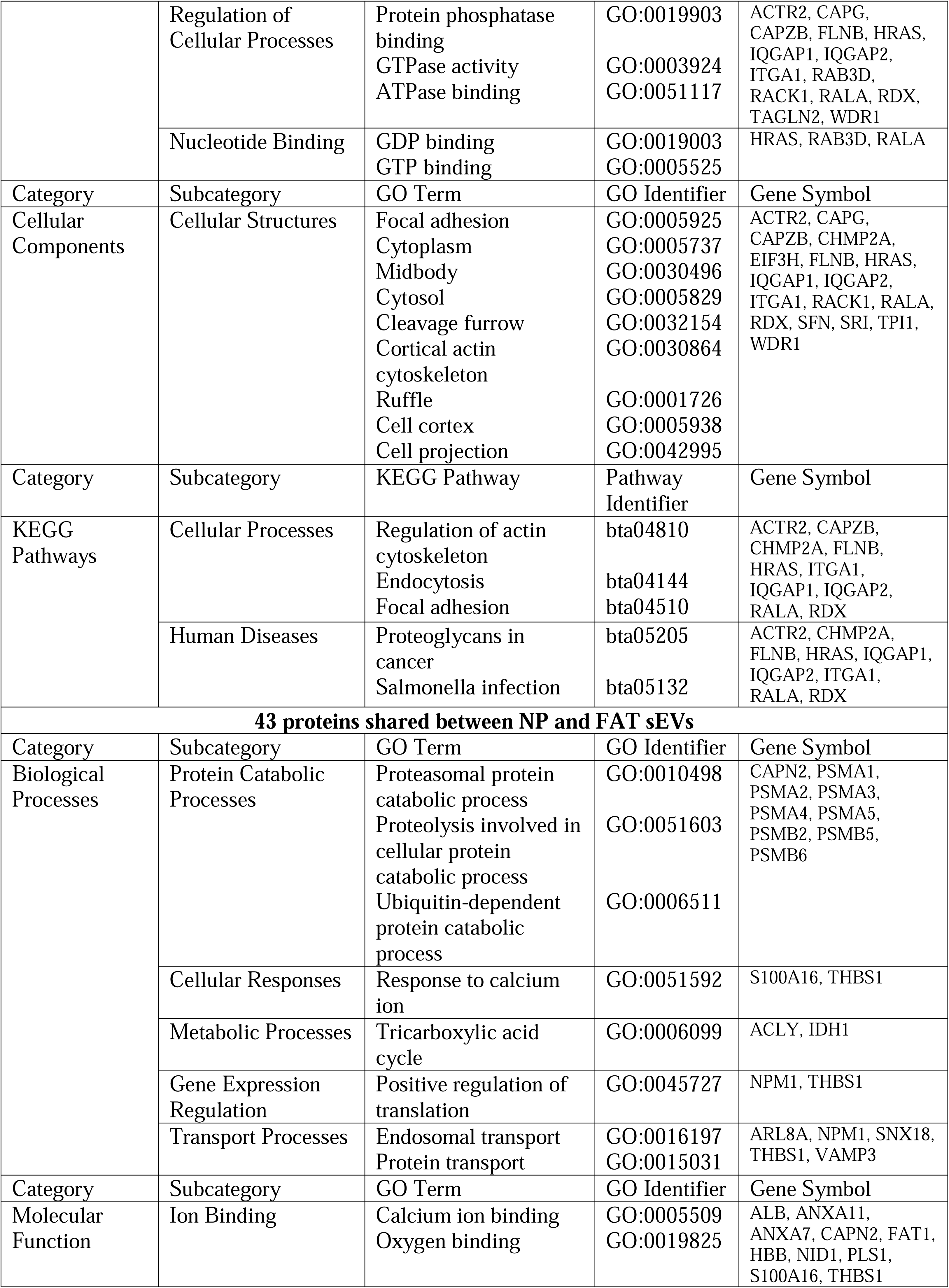
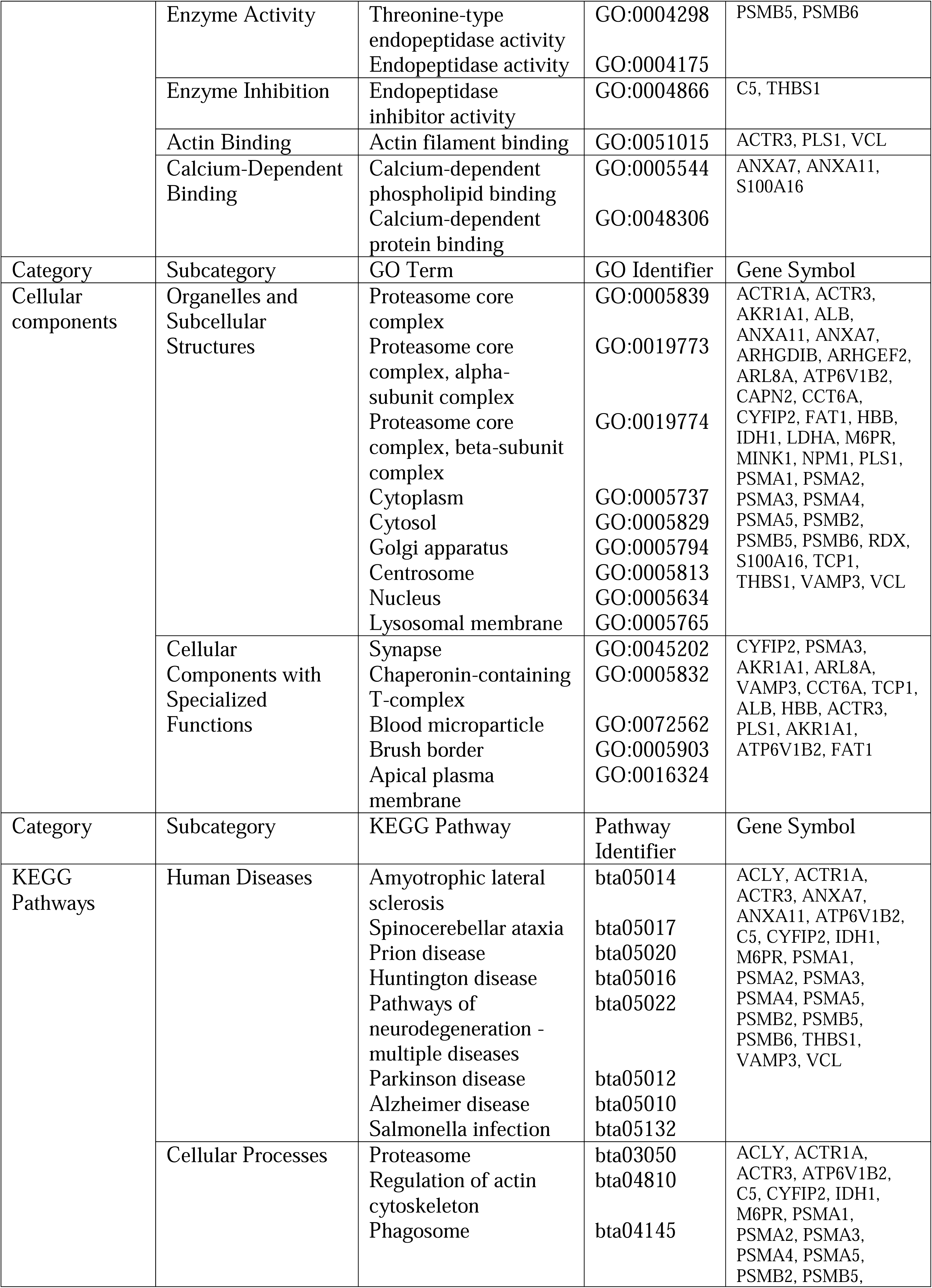
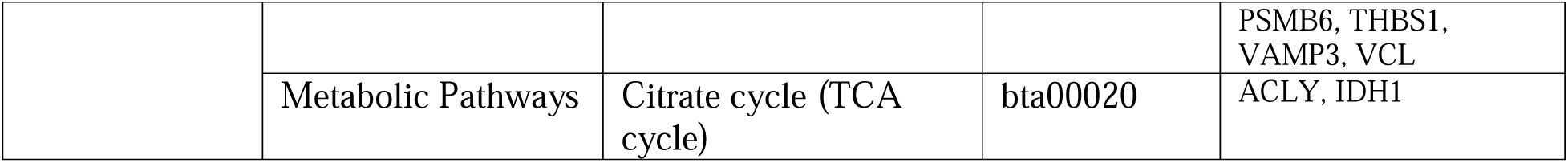
DAVID based functional enrichment analysis comparing sEV proteins of autologous bovine NP, AF, and FAT parent cells. AF: Annulus fibrosus; bta: Bos taurus; cAMP: Cyclic adenosine monophosphate; DAVID: Database for annotation, visualization, and integrated discovery; ECM: Extracellular matrix; FAT: Subcutaneous adipose tissue; GO: Gene ontology; KEGG: Kyoto encyclopedia of genes and genomes; FAT: Subcutaneous adipose tissue; MAPK: Mitogen-activated protein kinase; NP: Nucleus pulposus.

#### 3.3.3 Small EV Proteins from Adult or Fetal NP and MSC Parent Cells

A comparison of the sEV protein profile of adult and fetal NP parent cells with that of MSC sEVs was conducted to investigate if NP sEV proteins can positively impact on cell homeostasis or progenitor cell mobilization. In this study, data from human umbilical cord mesenchymal stem cells (UCMSCs) was chosen for comparison as it was the most comprehensive MSC sEV protein dataset at the time of the analysis (127). Of the sEV proteins identified, 206 were shared between fetal NP, adult NP, and UCMSC parent cells; additional 37 only between fetal and adult NP parent cells; 67 only between adult NP and UCMSC parent cells; and 37 only between fetal NP and UCMSC parent cells. Another 13 and 15 were only identified in sEVs of fetal or adult NP cells, respectively (Figure 5A). Functional enrichment analysis by DAVID of the 206 shared sEV proteins identified proteasome (bta03050), PPP (bta00030) and glycolysis/gluconeogenesis (bta00010) as pathways with high significance and highest FC (Figure 5B, Table 3, Supplementary figure 3). DAVID analysis associated the 37 sEV proteins identified for fetal NP and UCMSC parent cells essentially with endocytosis and vesicle formation, while sEV proteins shared between UCMSC and adult NP parent cells related to metabolism, though the FC was low (Figure 5C to E; Table 3). ToppFun pathway analysis for each of the three parent cell lines identified a total of 74 pathways common to all three sources, including the platelet-derived growth factor receptor β (PDGFRB) pathway (PID), and β-catenin independent WNT signaling (Reactome) (Figure 5F, Supplementary table 5). Among pathways linked to sEV proteins from UCMSC and adult NP parent cells were signaling by ROBO receptors (Reactome) and for UCMSC and fetal NP parent cells signal transduction by growth factor receptors and second messengers (Reactome). Many of the 57 pathways associated with sEV proteins shared between adult and fetal NP parent cells affected various signaling pathways including Notch signaling, but often through their proteasome association. Glycolysis/gluconeogenesis (KEGG) and aerobic glycolysis (WP) was identified, essentially via most proteins mentioned for glycolysis above (Table 3, Supplementary table 5). Hematopoietic stem cells are amongst the best studied progenitor cells and can be mobilized through the G-protein-coupled chemokine receptor CXCR4 and its ligand CXCL12 (137). These proteins were not detected here, however similar mechanisms might exist and association with the CXCR4 pathway (PID, M124) was suggested for sEV proteins shared between adult and fetal NP parent cells based on the identification of 14 proteins containing several integrin and G-protein subunits, receptor for activated C kinase (RACK1), RAC1, vacuolar protein sorting 4 homolog B (VPS4B), RAS homolog family member A (RHOA) and CDC42.

**Figure 5:**
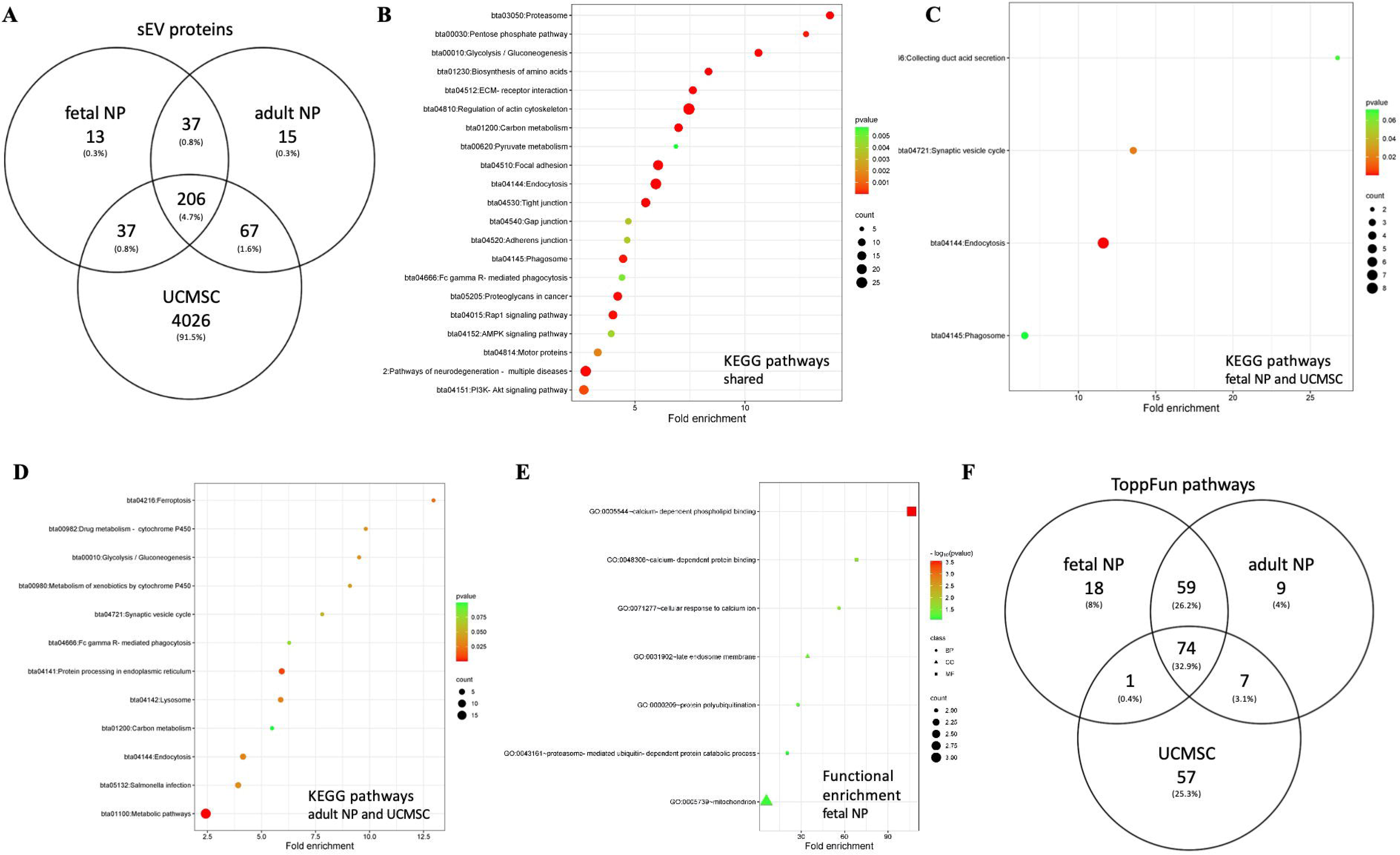
Nucleus pulposus and MSC sEV profiling: A) Comparison of sEV proteins from fetal and adult NP parent cells to those of UCMSCs (128). B) KEGG pathway analysis in DAVID of 206 shared proteins (C) KEGG pathway analysis in DAVID of sEV proteins shared by fetal NP and UCMSC parent cells. D) KEGG pathways analysis in DAVID of sEV proteins shared between adult NP and UCMSC parent cells. E) Functional enrichment analysis in DAVID of 13 proteins only identified for fetal NP sEVs. F) ToppFun pathway analysis of top 200 shared pathways. DAVID: Database for annotation, visualization, and integrated discovery; KEGG: Kyoto encyclopedia of genes and genomes; NP: Nucleus pulposus; UCMSCs: Umbilical cord mesenchymal stem cells.

**Table 3:**
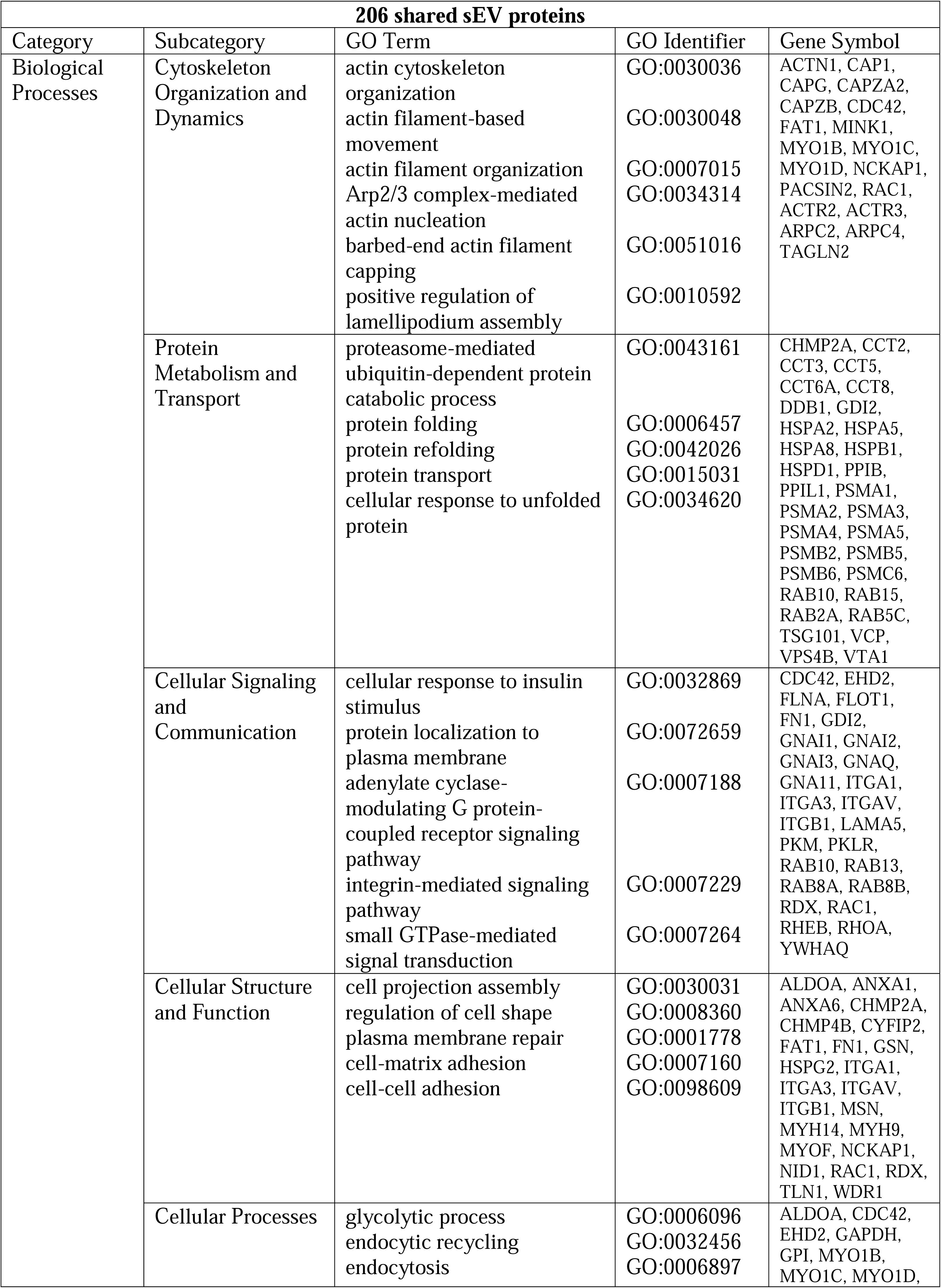
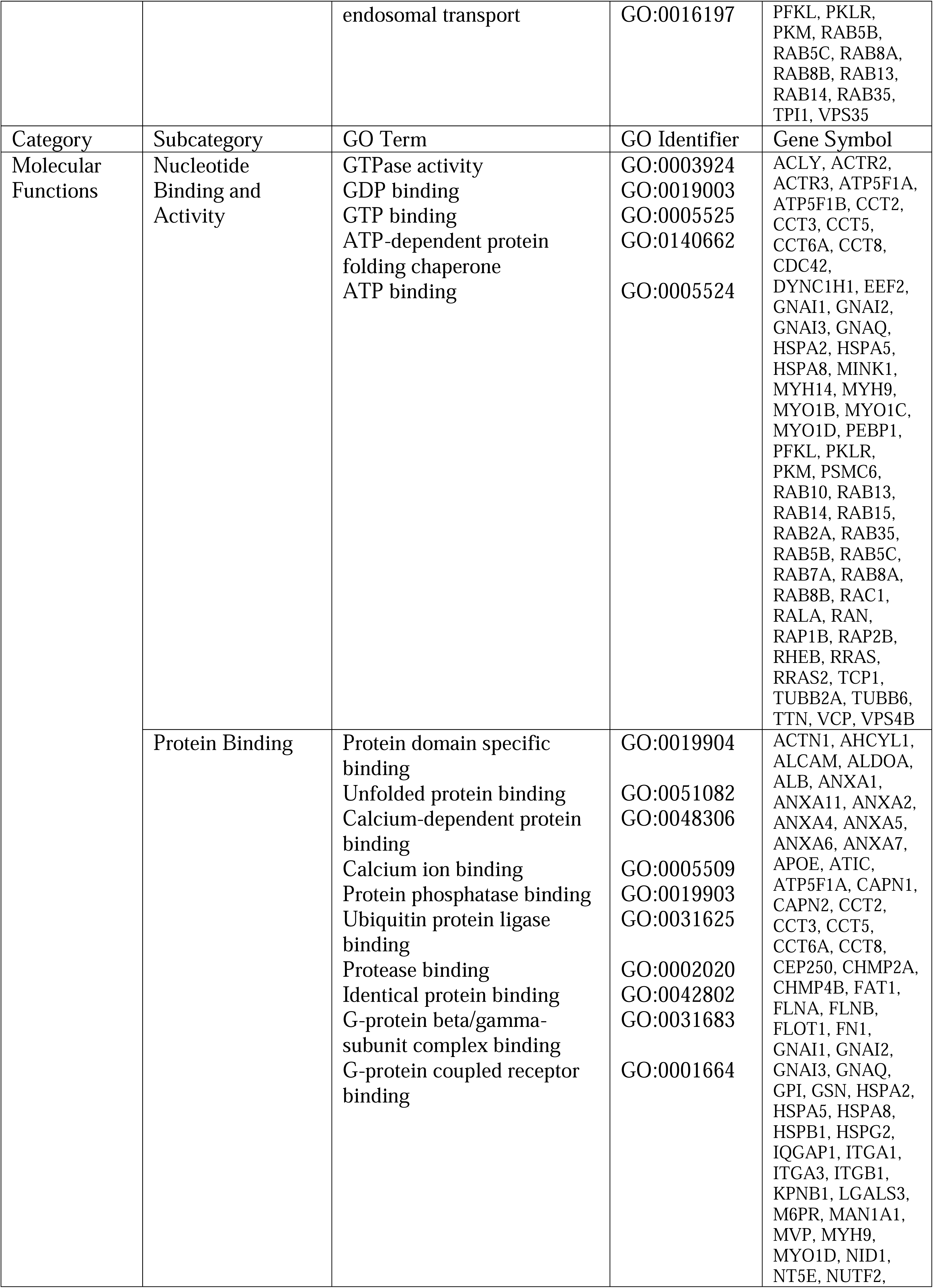
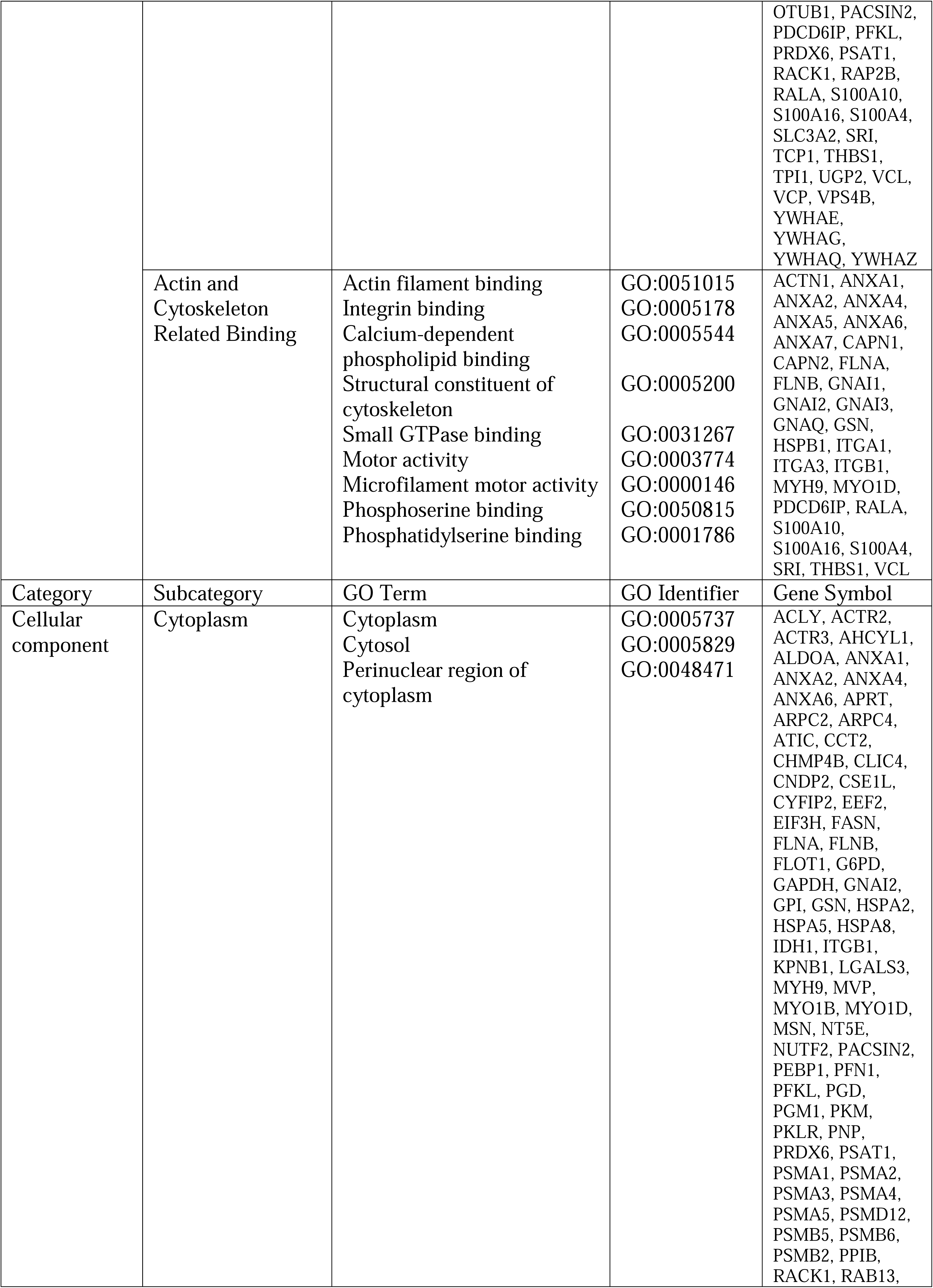
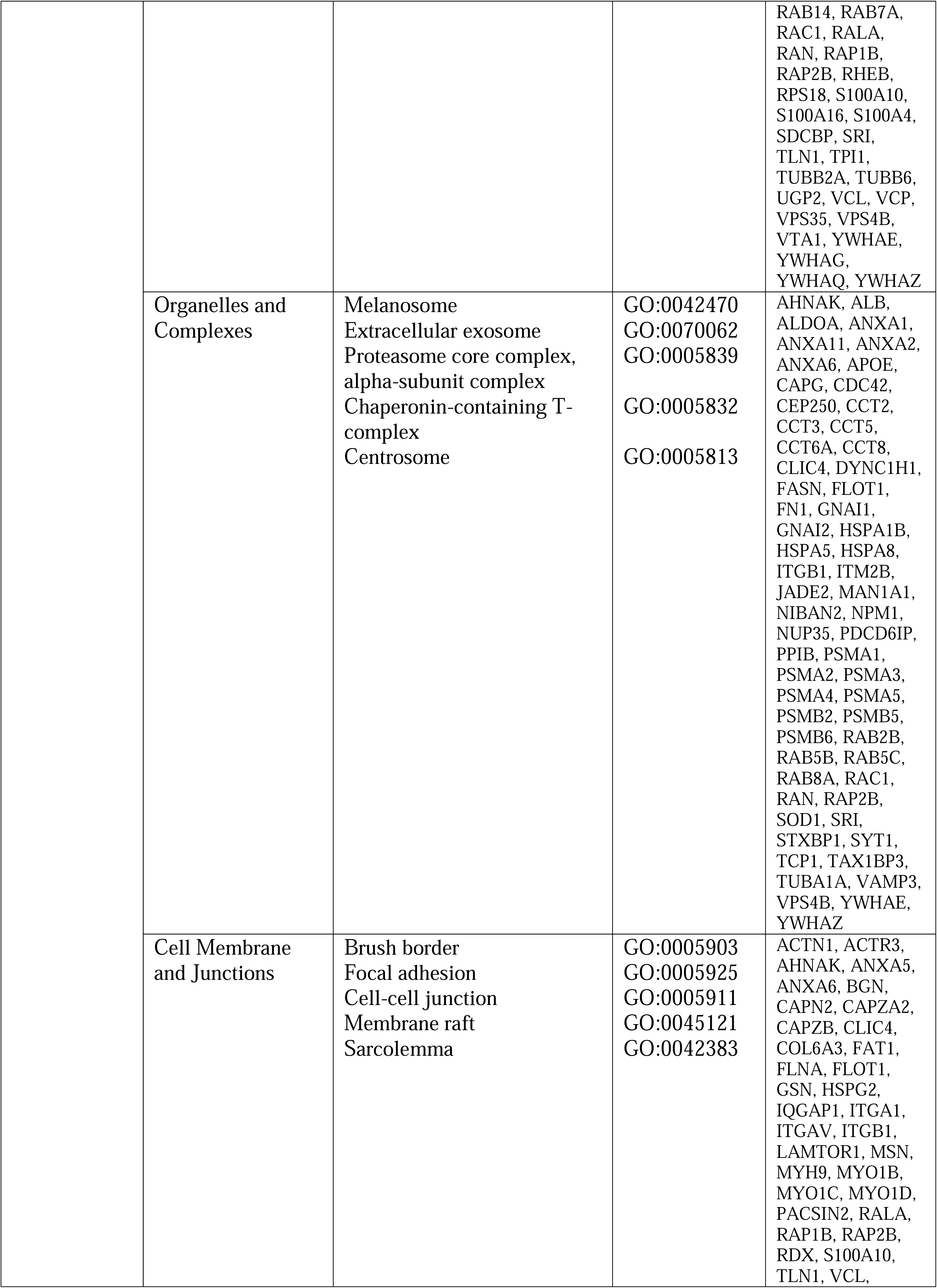
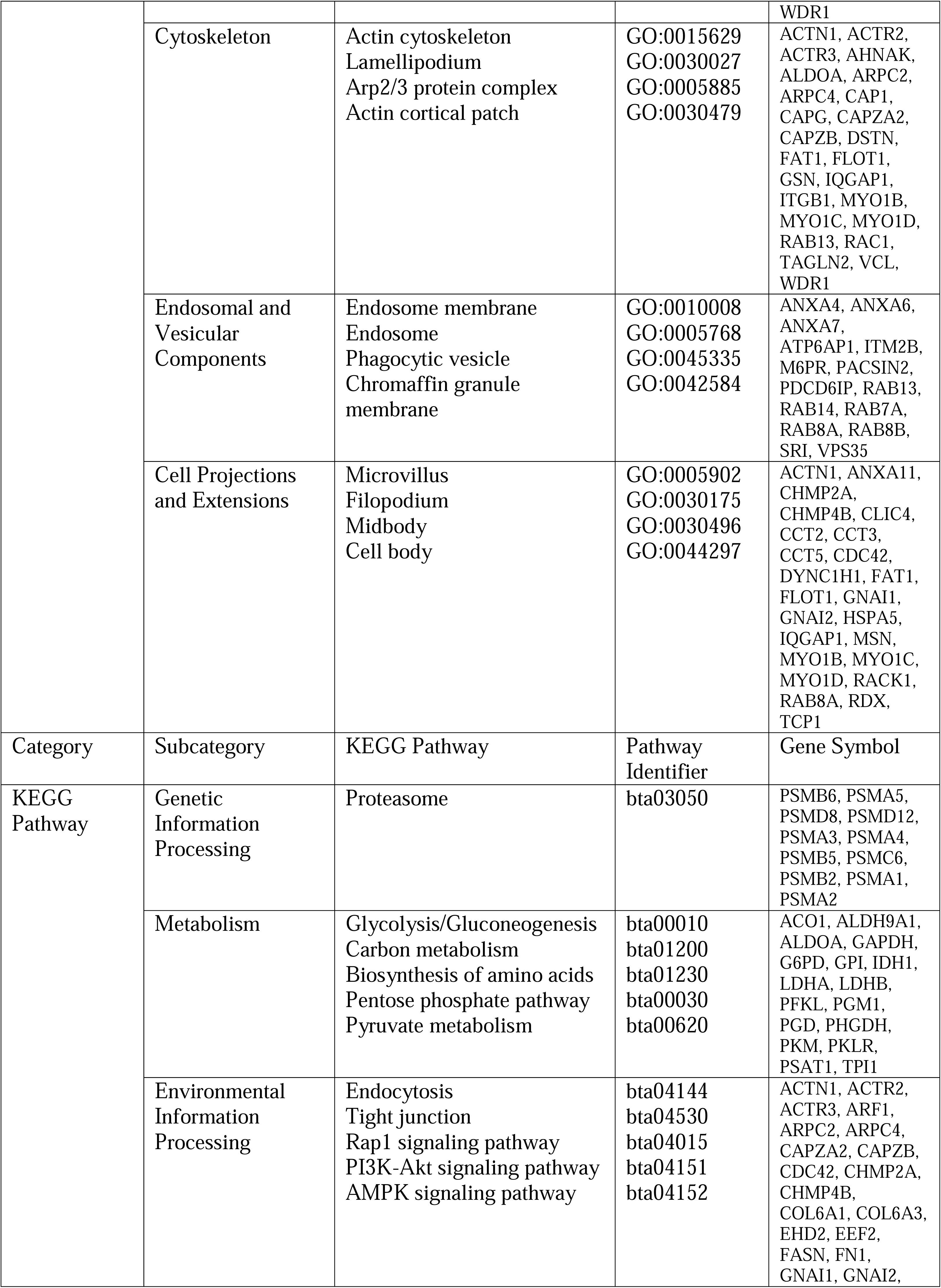
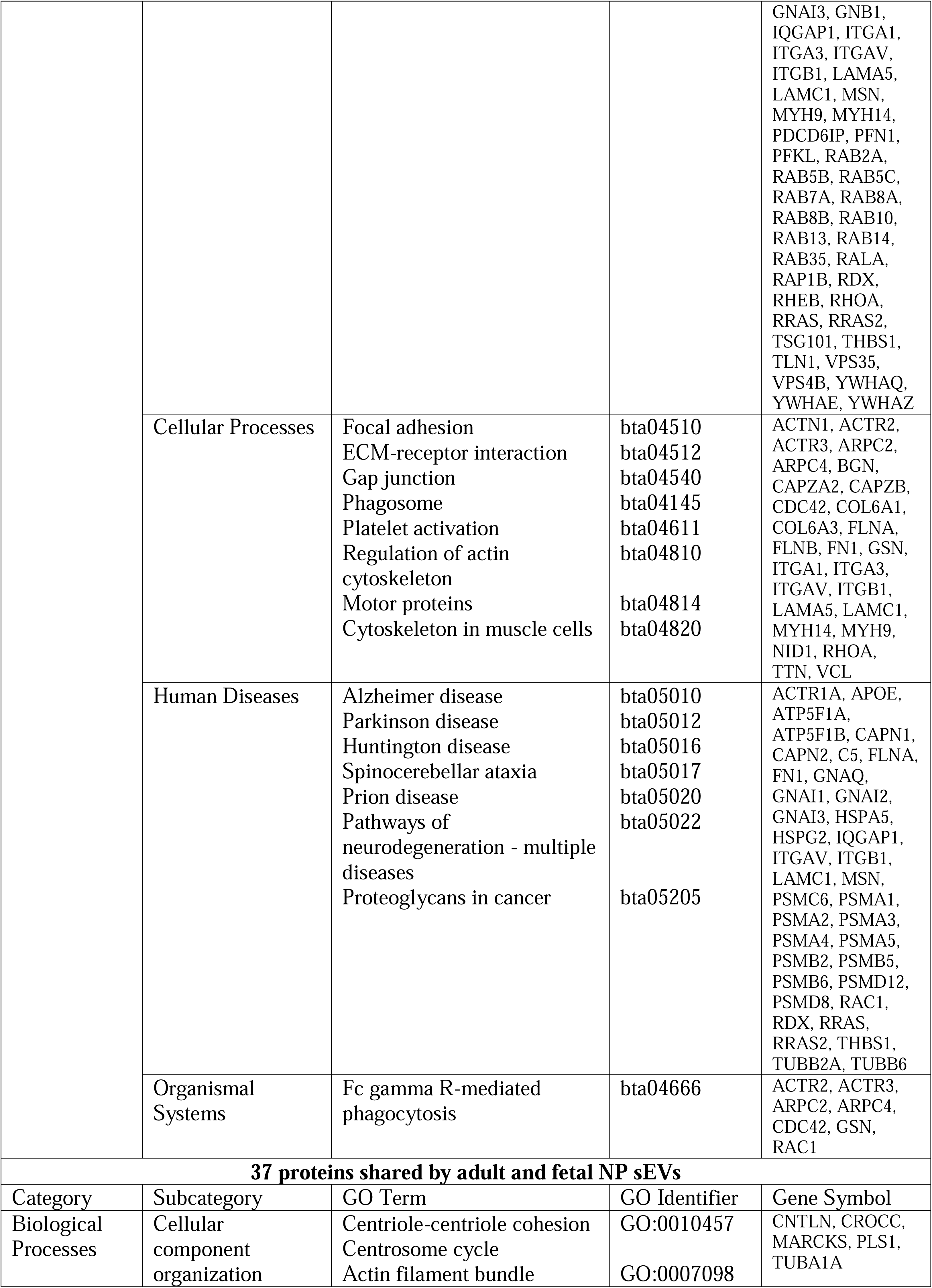
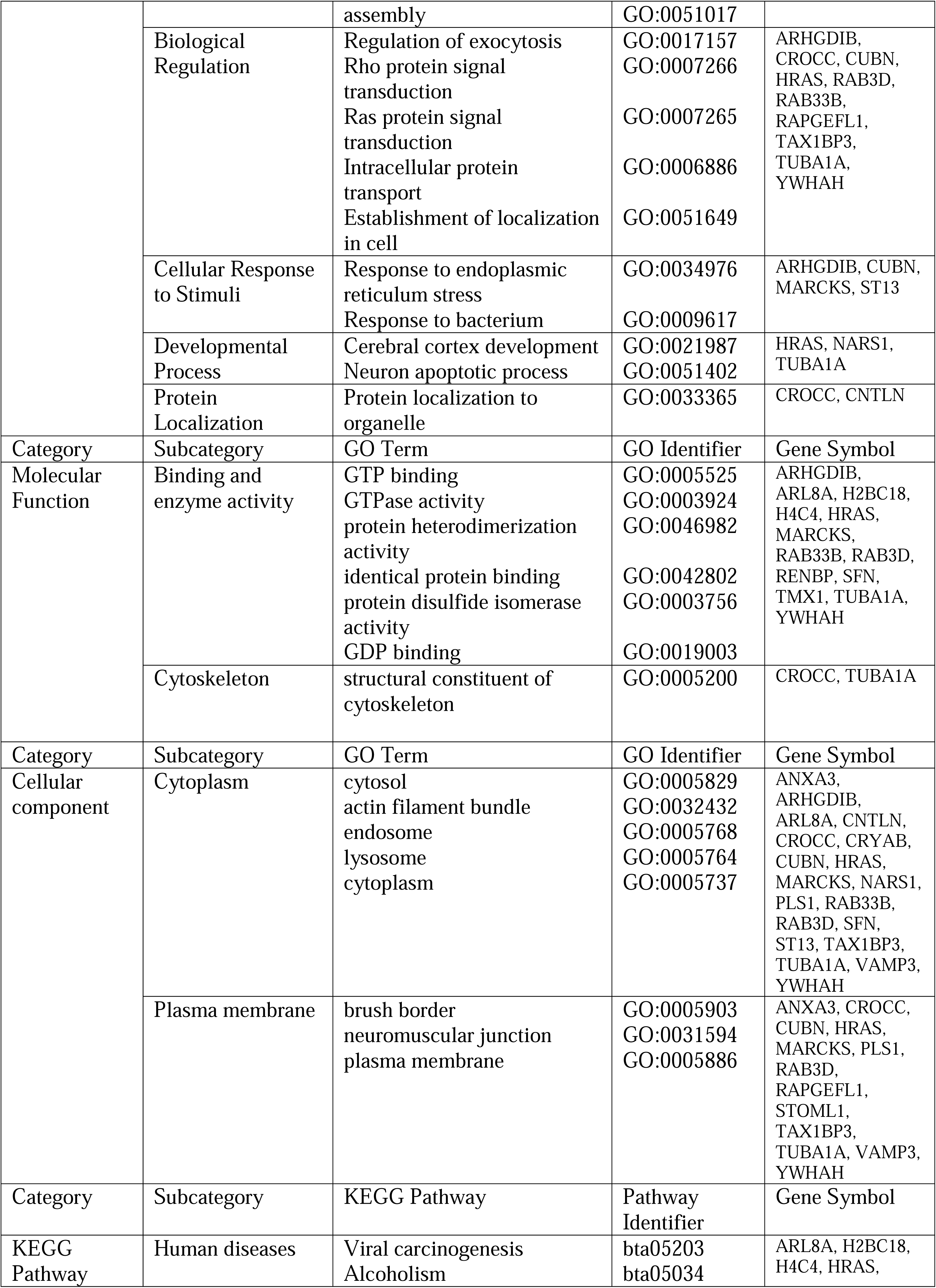
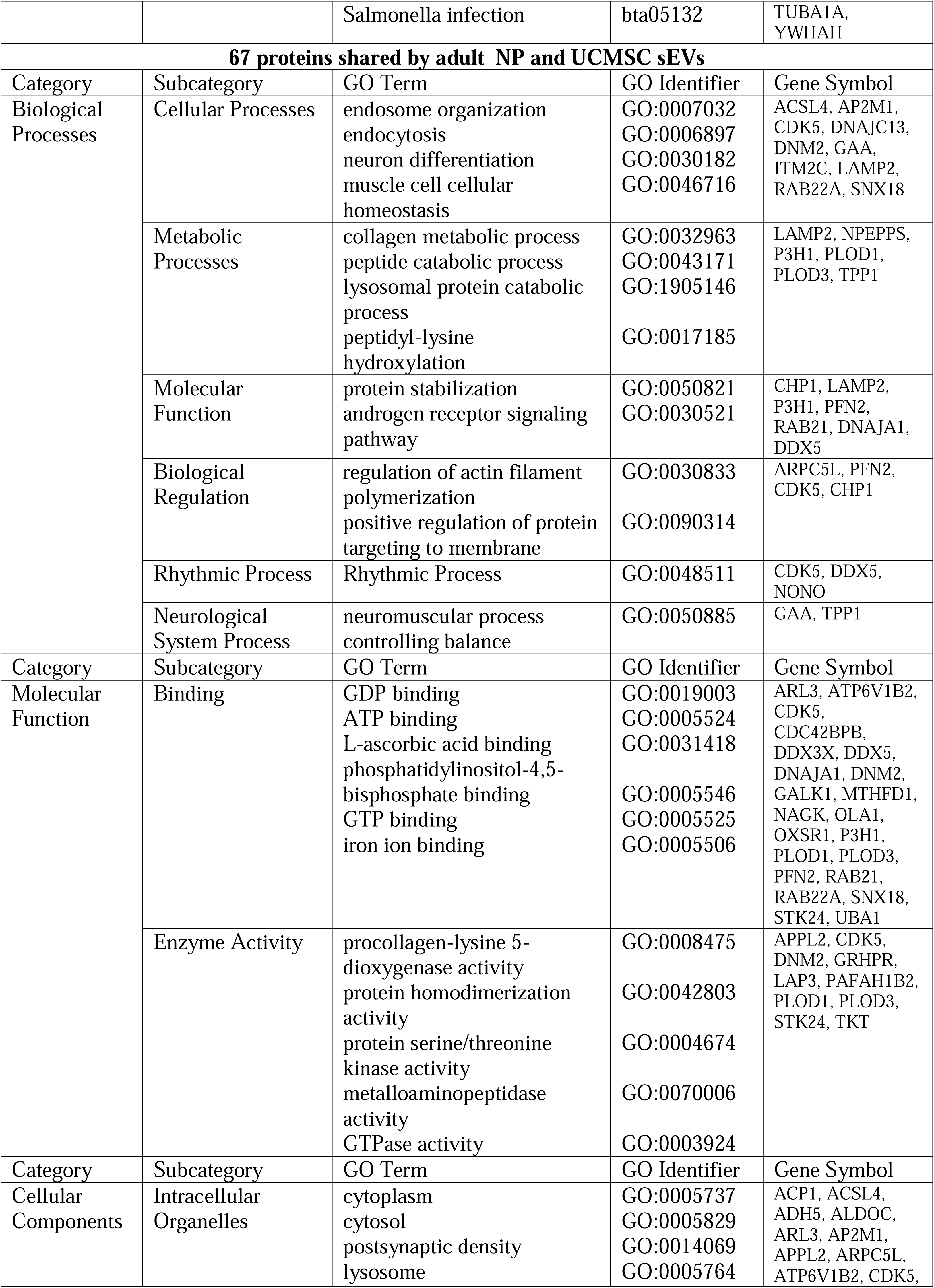
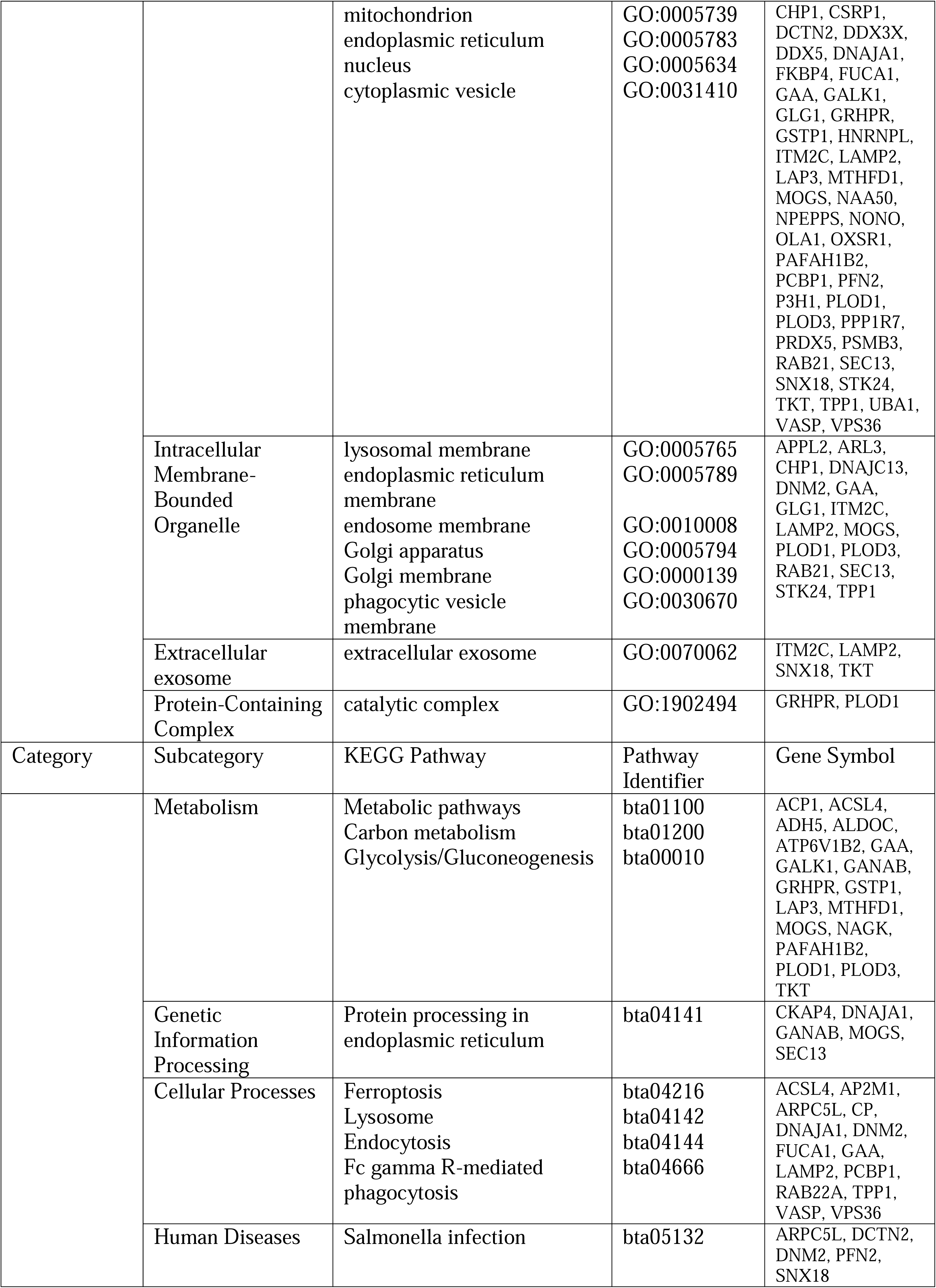
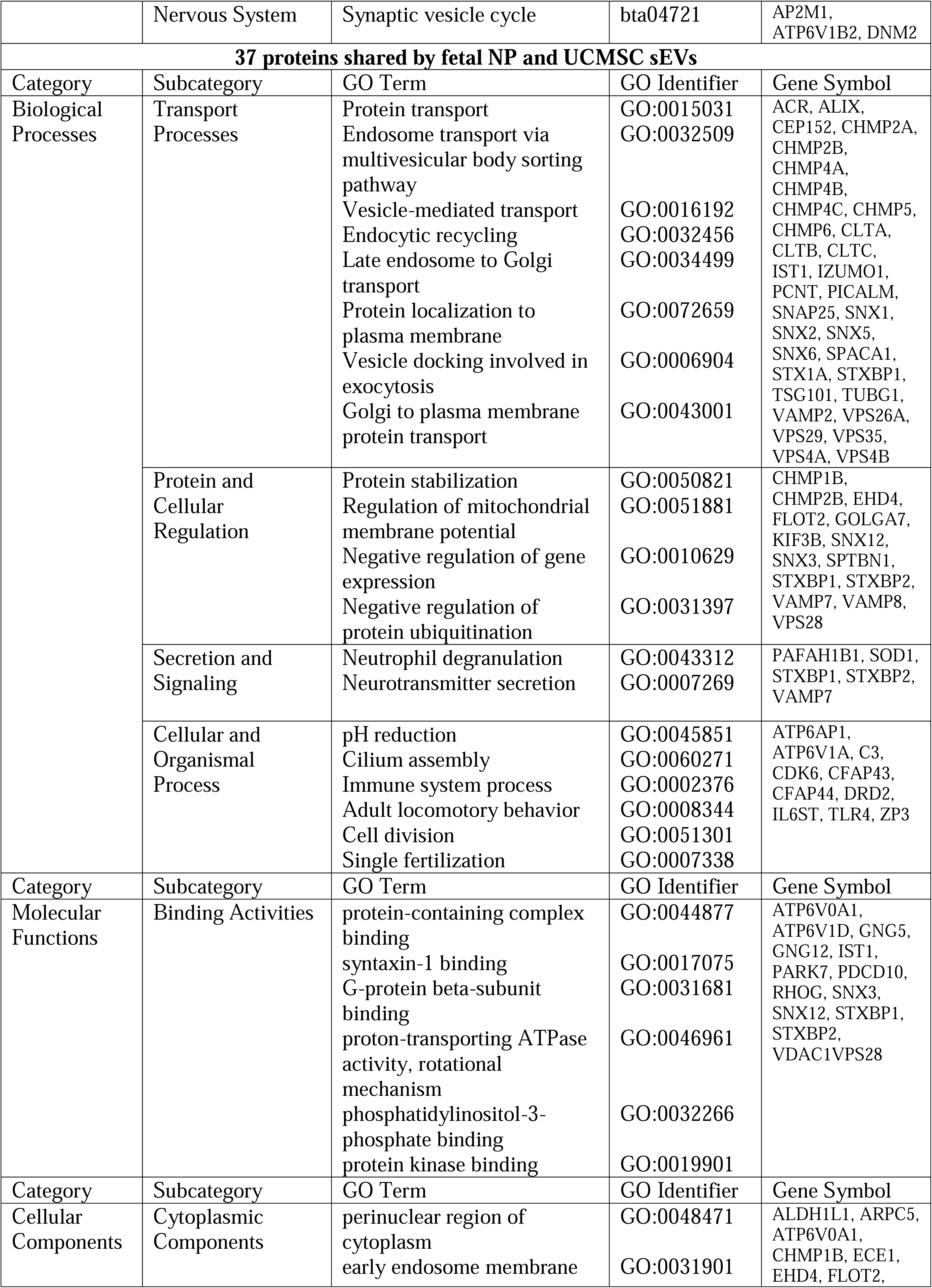
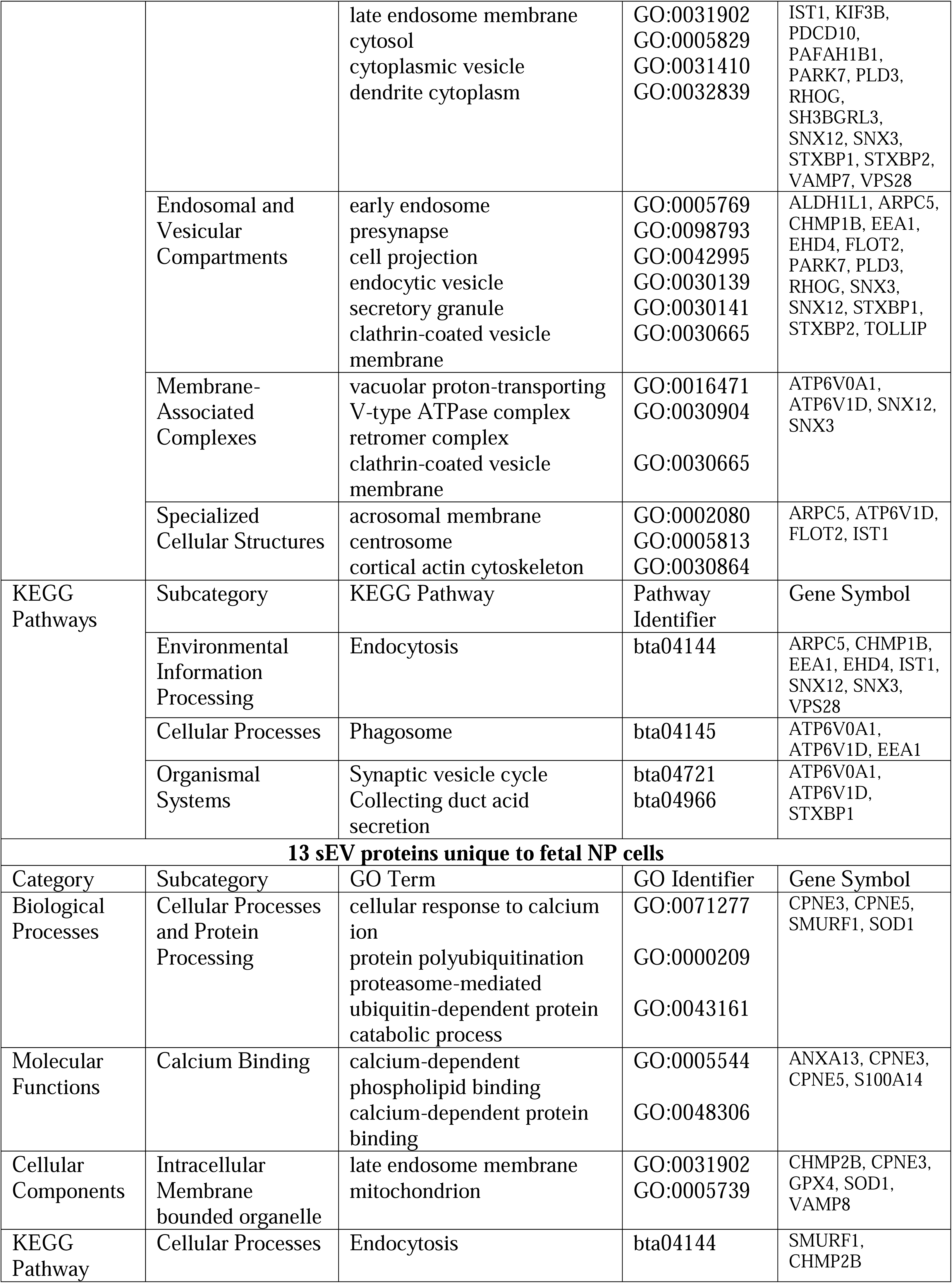
DAVID based functional enrichment analysis comparing sEV proteins of adult and fetal NP cells with UCMSC cells DAVID functional enrichment analysis for KEGG pathways of 207 shared exosome proteins of NP, fetal NP, and UCMSC cells. bta: Bos taurus; DAVID: Database for annotation, visualization, and integrated discovery; FAT: Adipose tissue; hsa: Homo sapiens; KEGG: Kyoto encyclopedia of genes and genomes; NP: Nucleus pulposus; UCMSC: Umbilical cord mesenchymal stem cells.

### 3.4 Quantitative Analysis of the NP Small EV Proteome

Bioactive molecules in the membrane or lumen of sEVs might end up there by chance, reflecting their abundance in the parent cell, or they could be actively deposited or removed during sEV biogenesis. To identify over- or underrepresented sEV proteins compared to their parent cell, quantitative proteomics of NP sEVs and parent cells was conducted. We identified a total of 151 differentially represented proteins, which were subjected to functional enrichment analysis using DAVID and ToppFun (Figure 6, Table 4, Supplementary figure 4, Supplementary table 6), then protein clusters were visualized in STRING (Figure 7).

**Figure 6:**
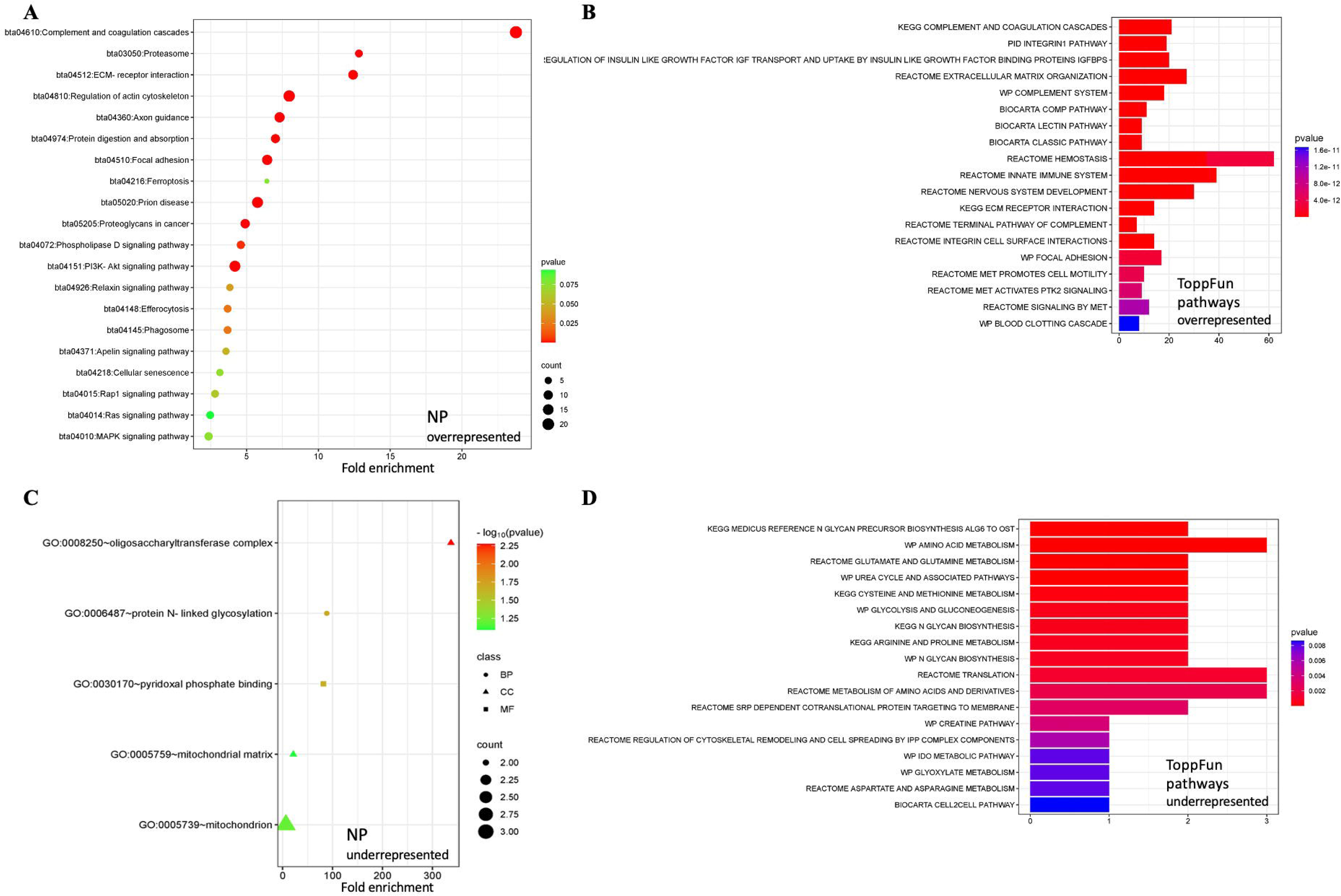
Quantitative functional enrichment analysis of nucleus pulposus sEV proteins: A) KEGG pathway analysis and (B) ToppFun pathway analysis of 141 proteins overrepresented in NP sEVs as compared to their parent cells. C) Functional enrichment analysis and (D) ToppFun pathway analysis of ten proteins reduced in NP sEVs as compared to their parent cells. DAVID: Database for annotation, visualization, and integrated discovery; KEGG: Kyoto encyclopedia of genes and genomes; NP: Nucleus pulposus.

**Figure 7:**
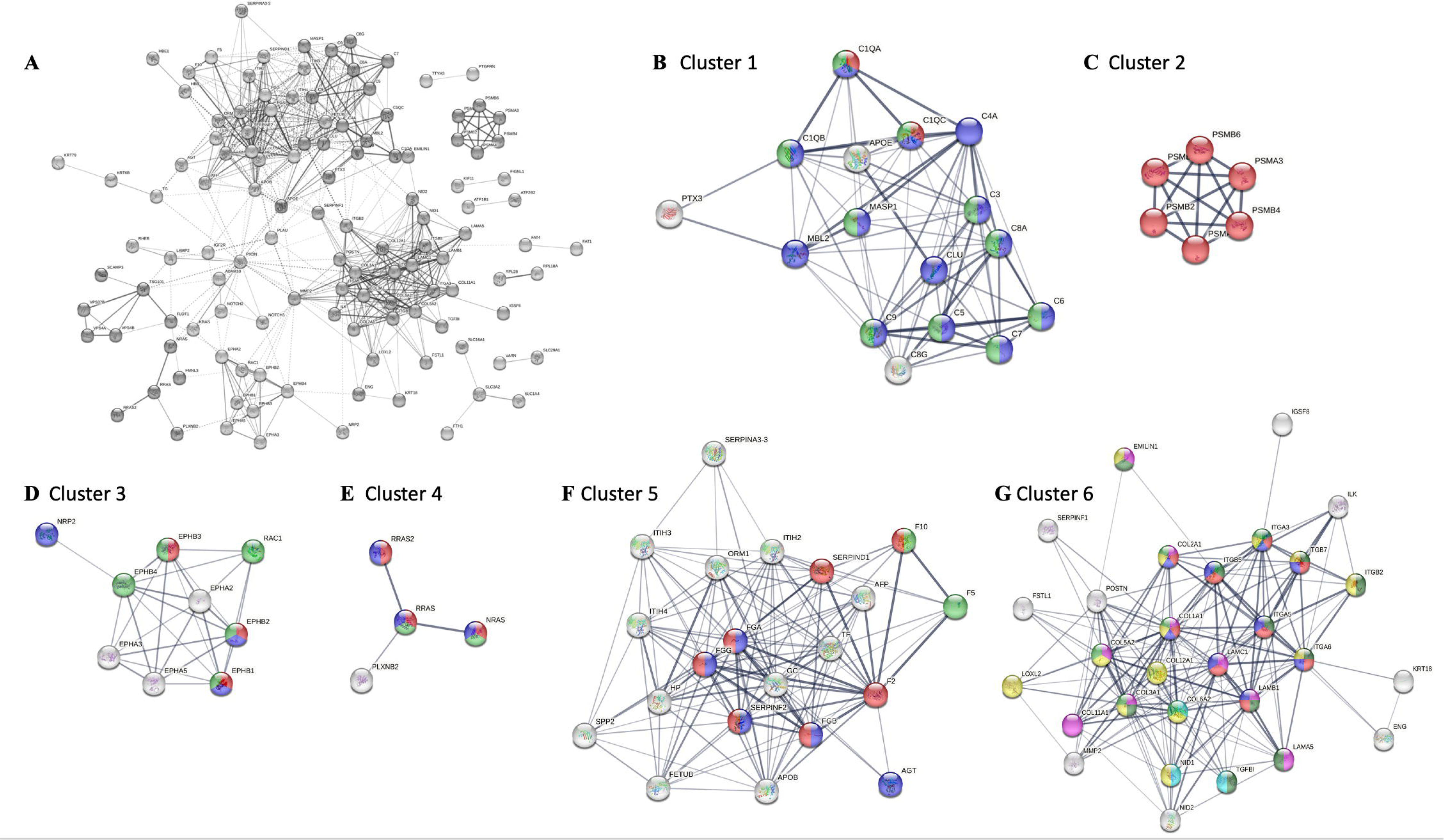
Functional clusters of overrepresented NP sEV proteins: A) STRING network of 141 overrepresented NP sEV proteins; B) Cluster 1: Complement component C1q complex (red), complement and coagulation cascades (blue), complement activation classical pathway (green); C) Cluster 2: Proteasome (red); D) Cluster 3: Axon (blue), axon guidance receptor activity (red), Ephrin signaling (green); E) Cluster 4: Ras signaling (red), axon guidance (green) and MAPK signaling pathway (blue); F) Cluster 5: Hemostasis (green), regulation of MAPK cascade (blue), complement and coagulation cascades (red); G) Cluster 6: ECM (pink), integrin binding (light green), collagen binding (sky blue), ECM receptor interaction (red), PI3K/AKT signaling pathway (blue), ECM organization (yellow), signaling by RTKs (dark green). STRING: Search tool for the retrieval of interacting genes/proteins; NP: Nucleus pulposus.

**Table 4:**
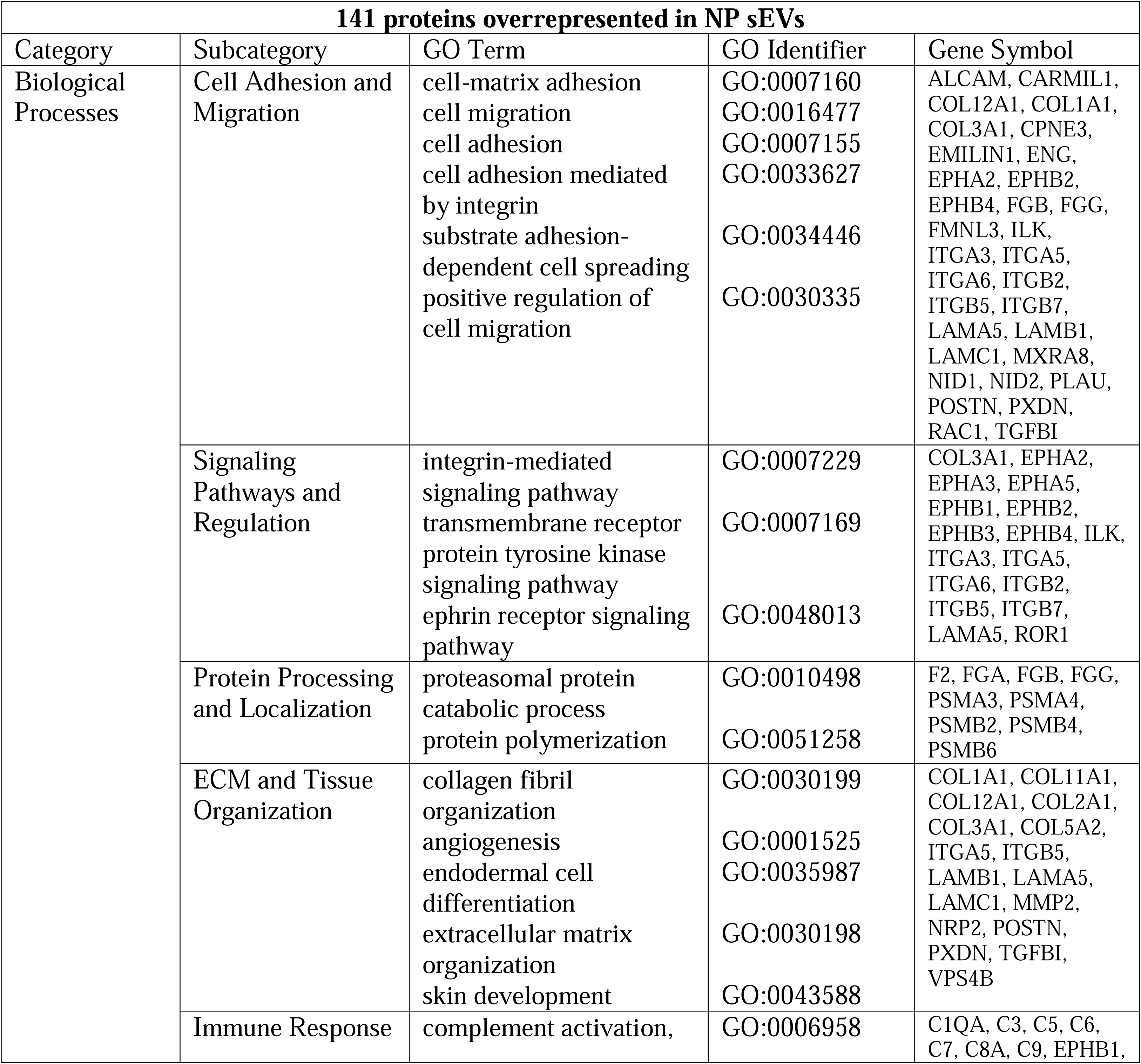
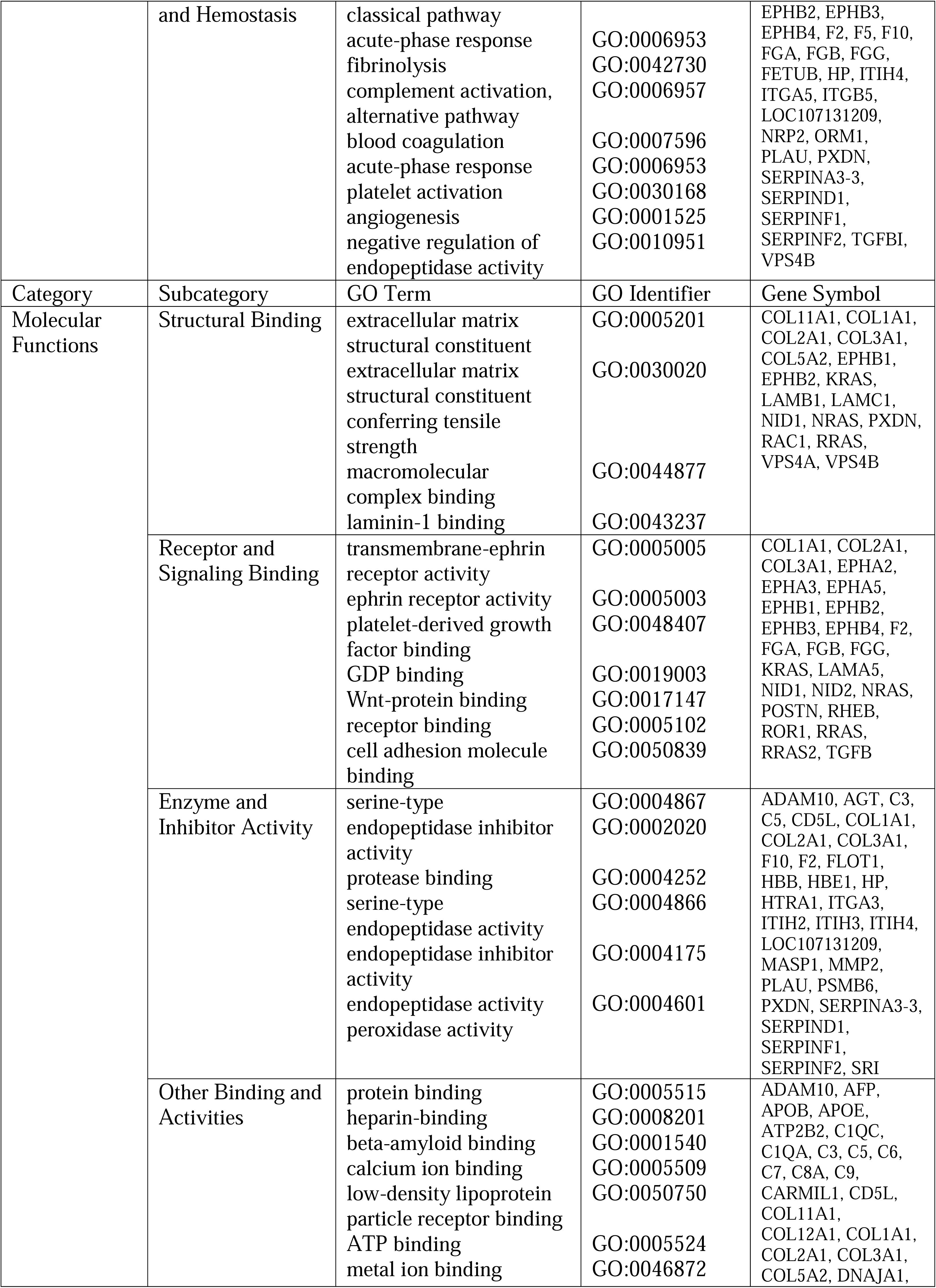
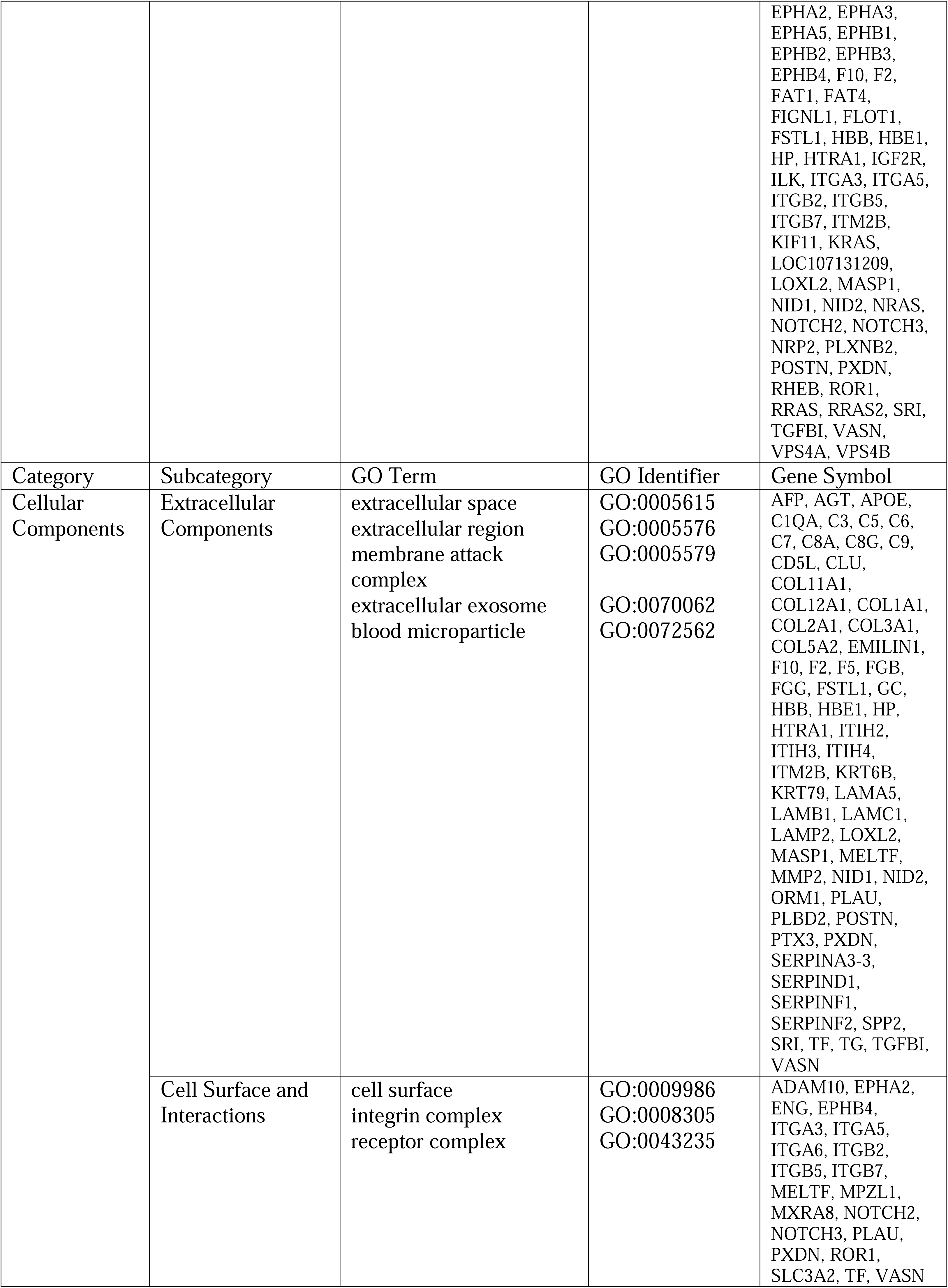
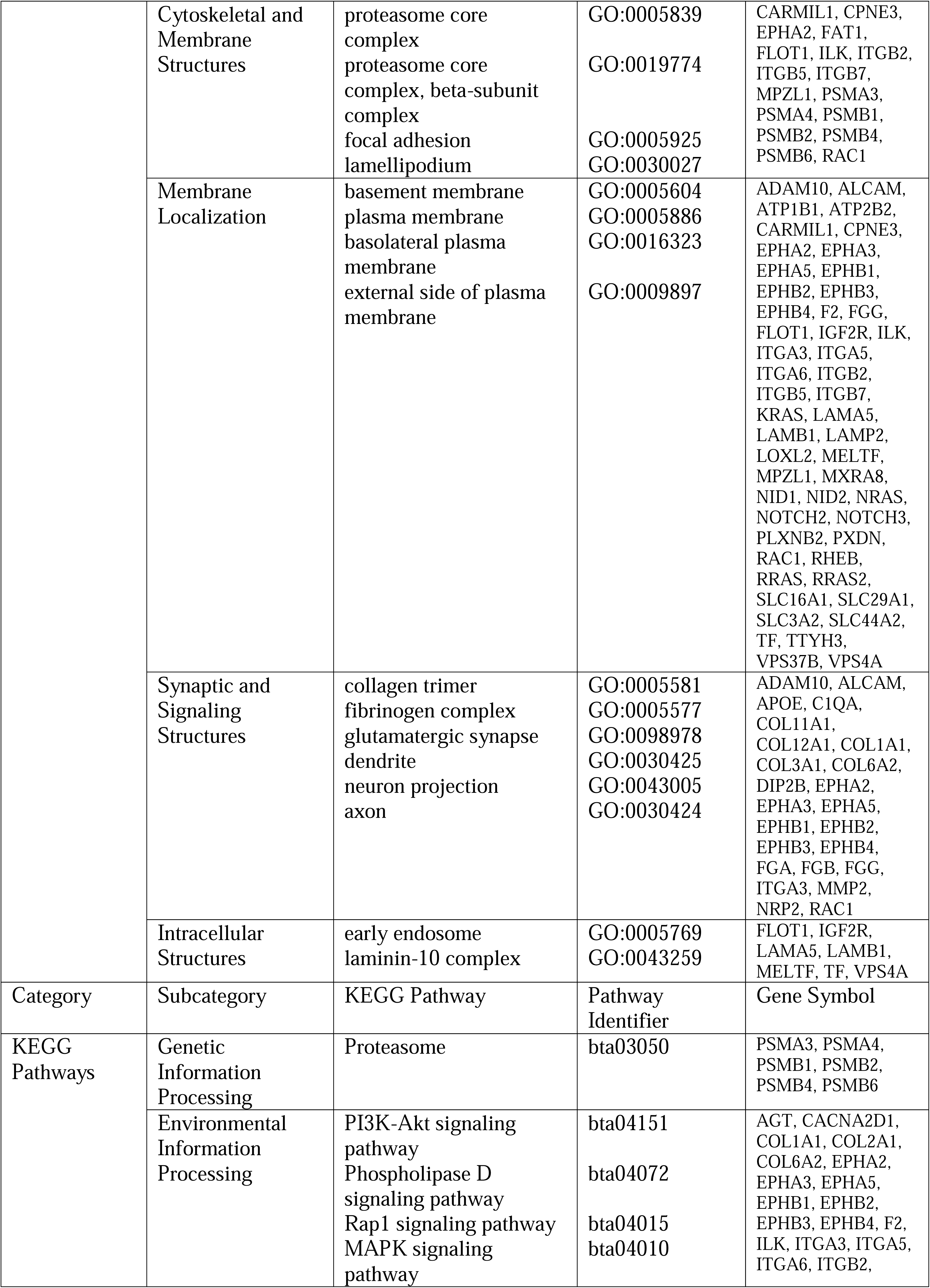
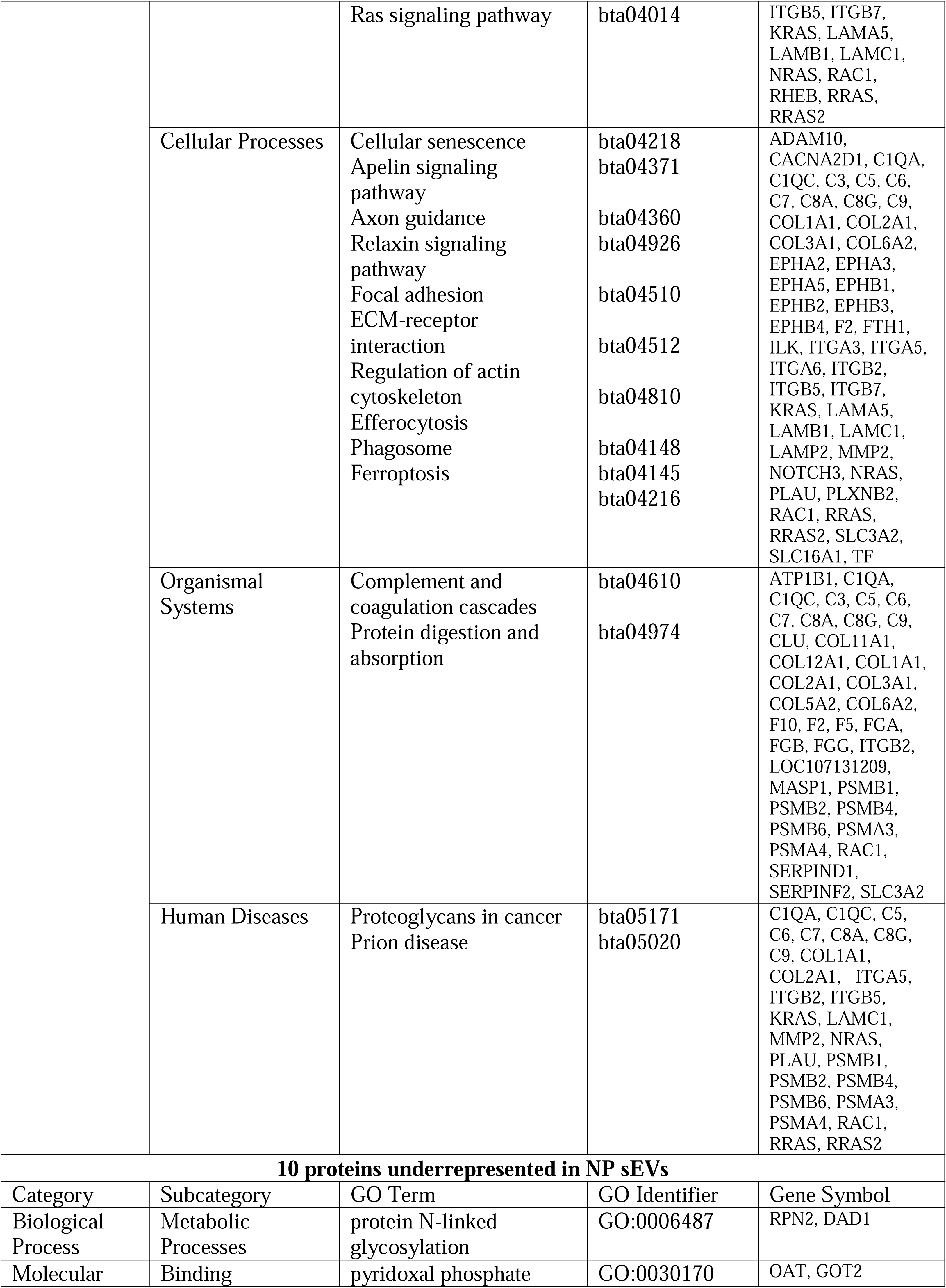
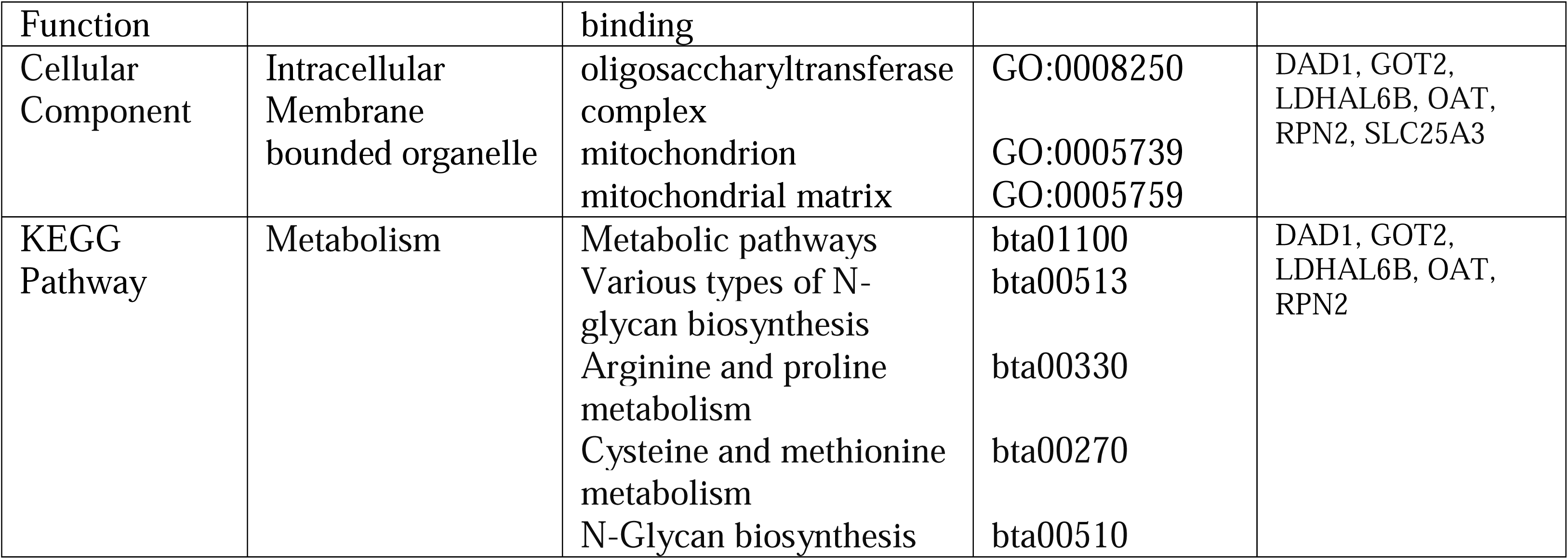
DAVID based functional enrichment after quantitative analysis of NP sEV proteins. bta: Bos taurus; DAVID: Database for annotation, visualization, and integrated discovery; GO: Gene ontology; KEGG: Kyoto encyclopedia of genes and genomes; NP: Nucleus pulposus.

#### 3.4.1. Differentially represented NP sEV proteins

Of the 151 proteins identified as differentially represented, 141 proteins were overrepresented in NP sEVs and ten underrepresented relative to their parent cells (Table 4, Supplementary table 6). Amongst the overrepresented proteins were Syntenin 1 or syndecan binding protein (SDCBP), integrin subunit β1 (ITGB1) and many GTPases, consistent with the sEV core proteome (133). Pathways were associated with important cell signaling cascades as well as cell and niche homeostasis as described in more detail below (Figure 6, Supplementary figure 4). While no pathways were identified for the ten underrepresented proteins using DAVID, these proteins were involved with posttranslational modifications (PTM) such as oligosaccharyltransferase (GO:0008250) and N-linked glycosylation (GO:0006487), and as coenzyme in the biosynthesis of amino acids, neurotransmitters and sphingolipids via pyridoxal phosphate (PLP) binding (GO:0030170). They further associated with mitochondria (GO:0005739) and mitochondria matrix (GO:0005759) (Figure 6C; Table 4, Supplementary table 6). Functional enrichment using ToppFun identified amino acid metabolism (WP, M39570), transaminase activity (GO:0008483) and glutamate and glutamine metabolism (Reactome, M27851) as underrepresented through an association with glutamic-oxaloacetic transaminase 2 (GOT2), tryptophanyl-tRNA sythetase 1 (WARS1) and ornithine aminotransferase (OAT). The mitochondrial lactate dehydrogenase A like 6B (LDHAL6B) was also underrepresented (Figure 6D, Supplementary table 6).

#### 3.4.2 Overrepresented NP sEV Proteins with Immunomodulatory Functions

STRING analysis of the 141 overrepresented NP sEV proteins revealed at least six distinct clusters (Figure 7A). The healthy NP is considered immune privileged (138), yet important elements in the innate immune response were identified in cluster 1 as relevant for complement activation and opsonization, including proteins associated with the complement component C1Q complex (GO:0062167), complement and coagulation cascades (bta04610) and complement activation classical pathway (WP977). The cluster contained most members of the complement cascade (C1 to C9) alongside the opsonins Pentraxin 3 (PTX3) and mannose binding lectin 2 (MBL2); the complement activating component mannose-binding lectin-associated serine protease 1 (MASP1); and Apolipoprotein E (APOE), an inhibitor of the classic complement cascade via C1Q binding (139) (Figure 7B). Proteasomes, intracellular non-lysosomal protein degradation complexes, as seen in cluster 2 have important functions in cell homeostasis, physiology and development (140).This cluster was comprised of subunits of the standard 20S proteasome, namely α3, α4, β1, β2, β4, and β6 (Figure 7C, Table 4).

#### 3.4.3 Overrepresented NP sEV Proteins Participate in Cell Signaling

Small EV proteins can be associated with the sEV membrane as ion channels, transporters, receptors or internal cargo. Of the 141overrepresented NP sEV proteins 38% were membrane associated. Amongst those were seven receptor tyrosine kinases (RTK), all of the EPH receptor type, two Notch receptors (NOTCH 2 and 3), one chloride anion channel (Tweety homolog 3 (TTYH3)), the auxiliary subunit of a calcium-voltage gated channel (CACNA2D1), and the insulin-like growth factor receptor 2 (IGFR2). All proteins have been previously identified as sEV associated and were listed in ExoCarta (Table 4) (124) .

##### 3.4.3.1 EPH Receptor Tyrosine Kinase (RKT) - RAS/MAPK Signaling Axis

Cluster 3 was composed of nine NP sEV proteins. Of those the Ephrin receptors EPHB1, EPHB2, EPHB3, EPHB4 and the RAC family small GTPase 1 (RAC1), a plasma membrane associated small GTPase involved in a range of cellular events (141–143), associated with the term ephrin signaling (bta3928664). EPHB1, EPHB2 and EPHB3 additionally associated with the term axon guidance receptor activity (GO:0008046), and EPHB1, EPHB2 and Neuropilin 2 (NRP2), a transmembrane protein and high-affinity receptor for some semaphorins (144), with the term axon (GO:0030424). Several components of the RAS pathway signaling cascade were also enriched in NP sEVs, forming the small cluster 4 including the RAS proto-oncogenes and GTPases NRAS, RRAS and RRAS2 and PlexinB2 (PLXNB2) a cell surface receptor for semaphorins, and RAC1 binding RAS homolog (RHEB1), and the RTH-like orphan receptor (ROR1), associating with the RAS (bta04014) and the MAPK (bta04010) signaling pathway. NRAS and RRAS also associated with axon guidance (bta04360). Cluster 5 consisted of 20 NP sEV proteins associated with the regulation of MAPK cascade (GO:0043408), namely the fibrinogen α (FGA), β (FGB) and γ (FGG) chains and angiotensinogen (AGT) (Figures 7D to F).

##### 3.4.3.2 ECM-Integrin Signaling Axis

The largest cluster 6 contained 29 overrepresented NP sEV proteins associated with ECM structural constituents (GO:0005201), ECM organization (GO:0030198) and ECM receptor interaction (bta04512), integrin-(GO:00005178) and collagen-(GO:00005518) binding. Amongst the proteins in cluster 6 were collagens (COL) 1, 2, 3, 5, 6, 11 and 12, various integrin subunits (ITG), laminins (LAMA5, LAMB1, LAMC1), and keratin 18 (KRT18) an intermediate filament with a long history in IVD research including being of NP biomarker status (77). Also, matrix metalloproteinase 2 (MMP2), transforming growth factor b induced (TGFbI) and more (Figure 7G). Especially the laminins and integrins but also COL1 and COL2 were associated with PI3K/AKT signaling (bta04151) through RTK signaling (bta9006934) and ECM receptor interaction (Figure 7G). .

##### 3.4.4 Other Overrepresented NP Small EV Proteins

Amongst proteins overrepresented in NP sEVs was SERPINF1 known as pigment epithelium-derived factor (PEDF), a protein with neurotrophic and antiangiogenic properties (135,145) and six proteins not represented in the ExoCarta database at the time, namely adiponectin (ADIPOQ), capping protein and regulator and myosin 1 linker 1 (CARMIL1), Fetuin B (FETUB), MBL, melanotransferrin (MELTF) and thyroglobulin (TG) (Table 4, Supplementary table 2).

### 3.5 NP Small EV Proteins with Links to Major Cellular Events

STRING was also used to identify NP sEV protein clusters (Figure 8A) based on all 484 NP sEV proteins identified here, not limited to those enriched in NP sEVs over their parent cells described above (Supplementary Table 2). The cluster of 22 proteins in Figure 8B, identified proteins involved in sEV biogenesis and the endosomal sorting complex (ESCRT). The largest cluster in Figure 8C contained 87 NP sEV proteins and was dominated by carbon metabolism (bta01200) with associations to metabolic processes including but not limited to core processes in energy production such as glycolysis/gluconeogenesis (bta00010), with GAPDH taking on a central role (Figure 8C). Also, part of this cluster were proteins involved in oxidative stress associated hypoxia inducible factors (HIF) signaling (bta04066), glutathione metabolism (bta00480), and proteins associated with pathways downstream of PI3K/AKT signaling (R-bta-111447, R-bta-9614399). A cluster of eight proteins involved in GPCR signal transduction in Figure 8D showed ties to axon guidance and Schwann cell migration. The cluster of 11 proteins in Figure 8E identified small GTPases such as CDC42 and RAC1 associated with EPHB-mediated forward signaling (R-bta-3928662) and connected to other important signaling pathways such as signaling by RHO GTPases (R-bta-194315) and VEGFA/VEGFR2 (R-bta-4420097). The small cluster of six proteins in Figure 8F related to the RAS proto-oncogenes which are crucial in many signal transduction pathways including the RAS (bta04014), PI3K/AKT (bta04151), MAPK signaling pathway (bta04010,) and related MAP2K and MAPK activation (R-bta-5674135). The cluster of 11 proteins in Figures 8G and H was largely composed of different integrin subunits, fibronectin (FN1) and the integrin-linked kinase (ILK), all involved in a range of biological processes including cell adhesion (GO:0033627), mesodermal cell differentiation (GO:0048333), angiogenesis (GO:0001525), regulation of small GTPase mediated signal transduction (GO:0051056), negative regulation of apoptotic process (GO:0043066) and ties to ECM receptor interaction (bta04512), focal adhesion (bta04510), TGFβ signaling (WP1045), the PI3K/AKT signaling pathway and axon guidance (bta04360). The cluster of nine proteins seen in Figure 8I was associated with EPHB-mediated forward signaling (R-bta-3928662) through subunits of the Arp2/3 protein complex that mediate actin polymerization. This cluster associated further with biological processes such as actin filament polymerization (GO:0030041), actin cytoskeleton organization (GO:0030036,) and positive regulation of lamellipodium (GO0010592), hence affiliated with cell migration. The cluster of 26 NP sEV proteins shown in Figure 8J associated with the PI3K/AKT signaling pathway affecting a range of biological processes including ECM assembly (GO:0085029), cartilage development (GO:0051216), angiogenesis (GO:0001525), blood vessel development (GO:0001568) and cell adhesion (GO:0007155).

**Figure 8:**
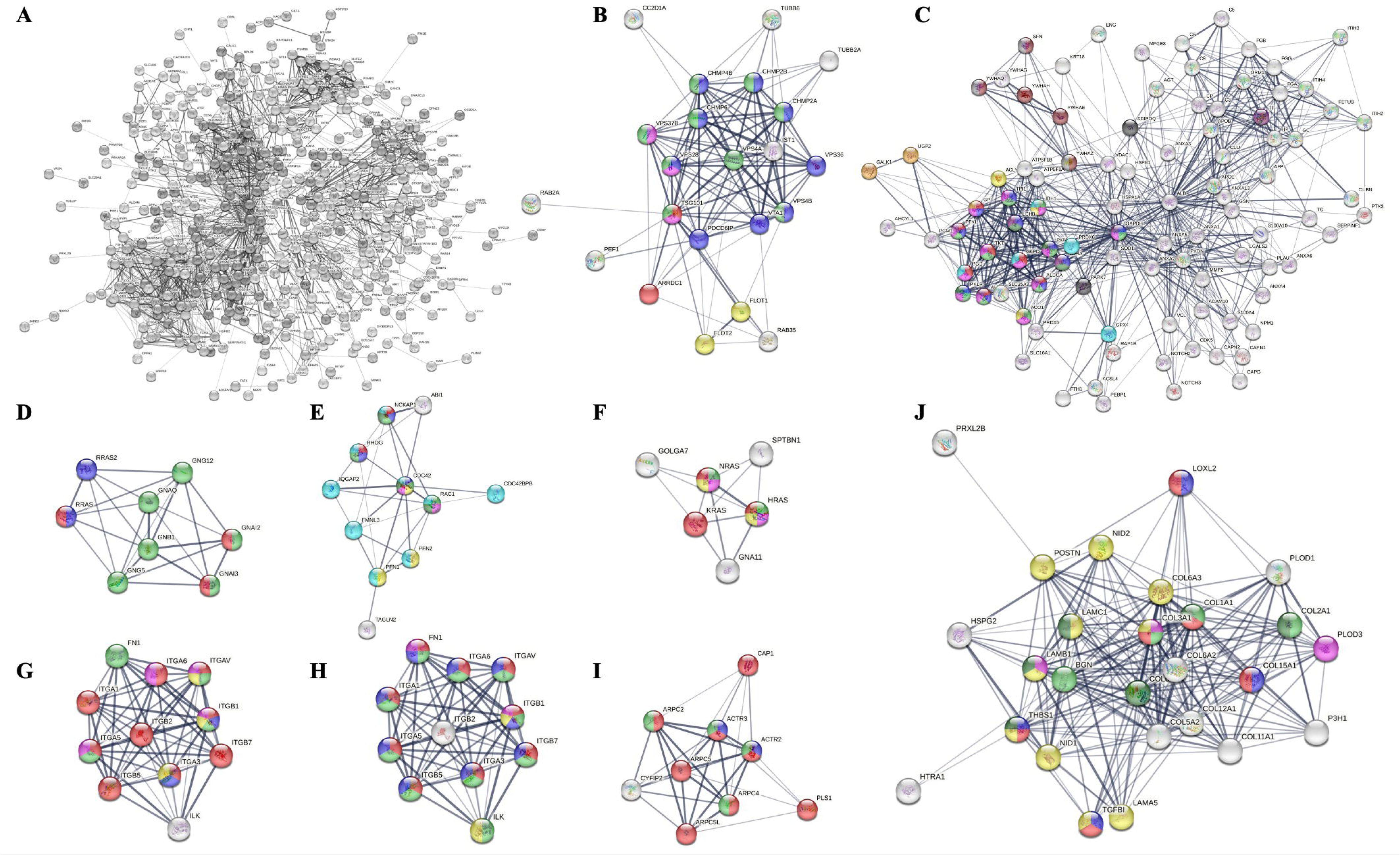
Functional clusters of all identified NP sEV proteins: A) STRING network of 484 NP sEV proteins; B) Cluster 1: Extracellular vesicle biogenesis (red), multivesicular body sorting pathway (blue), ESCRT I complex (pink), ESCRT (green), Flotillin complex (yellow); C) Cluster 2: Detection of oxidative stress (black), pentose phosphate pathway (red), glycolysis/gluconeogenesis (blue), biosynthesis of amino acids (green), carbon metabolism (pink), pyruvate metabolism (dark green), TCA cycle (yellow), glutathione metabolism (sky blue), HIF1 signaling pathway (purple), amino sugar and nucleotide sugar metabolism (ochre yellow), activation of BAD and translocation to mitochondria (brown), regulation of localization of FOXO transcription factors (grey); D) Cluster 3: Regulation of Schwann cell migration (blue), G-protein coupled receptor signaling pathway (green), axon guidance (red); E) Cluster 4: Rac protein signal transduction (red), RAS protein signal transduction (blue), small GTPase mediated signal transduction (green), RAP1 signaling pathway (yellow), EPHB-mediated forward signaling (pink), VEGFA/VEGFR2 pathway (dark green), signaling by Rho GTPases (sky blue); F); Cluster 5: MAP2K and MAPK activation (red), RAS signaling pathway (green), MAPK signaling pathway (pink), PI3K/AKT signaling pathway (yellow); G) Cluster 6 version 1: Cell adhesion mediated by integrin (red), mesodermal cell differentiation (blue), angiogenesis (green), regulation of small GTPase mediated signal transduction (yellow), negative regulation of apoptotic process (pink); H) Cluster 6 version 2: ECM receptor interaction (red), focal adhesion (green), PI3K/AKT signaling pathway (blue), axon guidance (yellow), TGFβ signaling pathway (pink); I) Cluster 7: Positive regulation of lamellipodium assembly (blue), actin cytoskeleton organization (red), EPHB mediated forward signaling (green); J) Cluster 8: ECM assembly (pink), cartilage development (green), angiogenesis (blue), blood vessel development (red), cell adhesion (yellow), PI3K/AKT signaling pathway (dark green). ECM: Extracellular matrix; ESCRT I: Endosomal sorting complex required for transport I; STRING: Search tool for the retrieval of interacting genes/proteins; NP: Nucleus pulposus.

Aside from gaining functional insight through the cluster analysis, the NP sEV proteins were also investigated in the context of specific intracellular signaling pathways and related key words using KEGG. In particular the RAS proto-oncogenes HRAS, KRAS and NRAS alongside other RAS homologs, members of the RAS and RAS-like oncogene families, GTPases and G-proteins with critical roles in many cell signaling events were identified in cell fate determining pathways such as RAS, MAPK, PI3K/AKT and MTOR signaling (Supplementary figure 5A). With an interest in cell fate decisions in tissue de- and regeneration, RAS proto-oncogenes via PI3K/AKT signaling and RHEB via MTOR signaling (bta04150) were associated with longevity regulating pathways (bta04211); while the calcium activated calpains CAPN1 and CAPN2 alongside RHEB and the RAS proto-oncogenes were linked to cellular senescence (bta04218). Other NP sEV proteins had ties to ferroptosis (bta04216) or the P53 signaling pathway (bta04115) (Supplementary figure 5B). Since communication with progenitor or stem cells in situ could be a key to regeneration, we investigated links between NP sEV proteins and pluripotency of stem cells (bta04550) but identified only the three RAS proto-oncogenes. In comparison, many proteins were identified for UCMSC derived sEVs all signaling through diverse pathways such as MAPK (map04010), Jak-STAT (map04630), Tgfβ (map04350) and Wnt (map04310) to activate the expression of pluripotency factors. Lastly, in the context of maintaining the unique avascular and aneural NP niche conditions, axon guidance and angiogenesis were investigated. Proteins were associated with cell adhesion (bta04514), axon guidance (bta04360) and RAP1 signaling (bta04015) with the integrin subunit β 1 involved in all three pathways. Several RAS family members alongside CDC42 associated with axon guidance and RAP1 signaling and many EPH-RTKs were involved in axon guidance by impacting on axon outgrowth and repulsion (Supplementary figure 5C). Lastly, Apelin (bta04371), VEGF (bta04370) and chemokine (bta04062) signaling, and proteoglycans (bta05205) were investigated in the context of vascularization, highlighting again a central role of the RAS proto-oncogenes and players in the RAS signaling pathway (Supplementary figure 5D).

### 3.6 Removing the Influence of a Serum Corona

Comparing sEV proteins identified for all parent cell types, including those enriched in NP sEVs and the ExoCarta database, we routinely identified commonly shared pathway associations in ToppFun, namely: The Reactome pathways neutrophil degranulation, innate immune system, immune system, platelet activation signaling and aggregation, nervous system and WP VEGFA/VEGFR2 signaling, WP focal adhesion as well as KEGG focal adhesion and KEGG regulation of actin cytoskeleton. Reactome pathway associations with all three pathways of the complement system were only found in the dataset from overrepresented NP sEV proteins with clusterin (CLU) being an inhibitor of the complement membrane attack complex (MAC) and therefore cell lysis (Supplementary figure 6, Supplementary table 7). Yet recent work demonstrated that proteins spontaneously form a corona around the surface of EVs or virus particles when maintained in plasma (146). Many cell culture media contain serum in form of fetal bovine serum (FBS) and that, though depleted of sEVs, contains plasma proteins.

Mass spectrometry analysis of the FBS used here for culture identified a list of 523 proteins (Supplementary table 8). Upon removing proteins shared between the serum and sEVs, functional enrichment analysis was repeated for 119 remaining overrepresented NP sEV proteins (Supplementary figure 7). The serum protein depleted dataset still identified complement and coagulation cascades (bta04610) and axon guidance (bta04360) but now also affiliated with longevity regulating pathway (bta04211), efferocytosis (bta04148), cellular senescence (bta04218) and more. The longevity connection was established through the RAS GTPases NRAS, KRAS and RHEB and ADIPOQ, a plasma protein with anti-inflammatory and antioxidant effects (147) involved in AMP-activated protein kinase (AMPK) signaling (bta04152) for which we previously detected transcripts in cultured NP cells (148). Efferocytosis enables tissue homeostasis following distinct phases of finding, binding, internalization and breakdown of an apoptotic cells (149). The proteins involved were chains of serum complement subcomponent C1q (C1QA, C1QC); the solute carrier SLC16A1 involved in transport of monocarboxylates like pyruvate and lactate and driven by the proton motive force (150); integrin subunit (ITGB5); the disintegrin and metalloproteinase domain-containing protein (ADAM10) and the small GTPase RAC1. The association with cellular senescence was established through the small GTPases (NRAS, RRAS, RHEB, RRAS2, KRAS). ToppFun analysis still identified complement and coagulation cascades (KEGG M16894, WP M39649), integrin pathway (PID M18), ECM organization (Reactome M610, MM14572) and hemostasis (M8925) amongst the top pathways (Supplementary table 8). Interestingly, EPH-ephrin mediated repulsion of cells (Reactome, M27311), EPH forward pathway (PID M62), EPH-ephrin signaling (Reactome, M27201) and axon guidance (KEGG M5539) remained significant, essentially through the EPH receptor-RAS signaling axis and ECM components (Supplementary table 8).

Similar associations were observed for all 300 NP sEV proteins after removing the serum proteins from the dataset. DAVID analysis still identified complement and coagulation cascades, axon guidance, glycolysis/gluconeogenesis, PPP, and PI3K/AKT signaling and ToppFun essentially resembled what was seen for overrepresented NP sEV proteins without serum proteins. Now significant additional pathways of interest like signaling by NOTCH4 (Reactome M954), regulation of RUNX3 expression (MM15536), somitogenesis (Reactome, M48031), and stabilization of P53 (Reactome M27670) were essentially based on an affiliation with the proteasome 20 subunits (Supplementary figure 7, Supplementary table 8).

Of the corona proteins reported for the serum of healthy participants (146), 11 were identified in our NP sEV protein data set, namely clusterin (CLU); fibronectin chains α, β and γ (FGA, FGB, FGG); complements C3 and C4; α1-acid protein 1 (ORM1); ceruloplasmin (CP); inter-alpha-trypsin inhibitor heavy chains ITIH2, ITIH4; and apolipoprotein E (APOE). Amongst proteins found in viral coronas (146) APOB, APOE, C3 and C4 overlapped with our data set of NP sEV proteins.

## DISCUSSION

Extracellular vesicle research is an evolving field, even more when it comes to cells of the IVD. Amongst spine related health concerns IVDD is one of the major causes of LBP, a degenerative disease currently lacking a cure (6). Regenerative stem cell therapies have been considered for IVDD; however, the harsh NP microenvironment poses a challenge. The NP is of great functional importance, by volume the largest tissue type in the IVD and due to its avascular and aneural nature the most likely tissue to depend on sEV exchange for communication between isolated NP cells in the ECM. A strategy in tissue regeneration is the activation of endogenous progenitor cells. It has been acknowledged that the MSC secretome, including EVs such as sEVs, plays an important part in immune modulation and regeneration (151,152). Small EVs, as part of many cells’ secretome, showed favorable therapeutic properties in various applications including increased cell proliferation, ECM synthesis, migration of repair-involved cells to an injury site, and promoted the differentiation of MSCs into NP cells while also showing reduced apoptosis, inflammation, ECM degradation, and cell senescence in NP cells (93,104,153). A recent study highlighted the therapeutic potential of implementing a TIE2-positive(+) cell-enhancing protocol, emphasizing the importance of optimizing culture conditions. The EV product isolated from TIE2^+^ NP cells demonstrated promising regenerative outcomes, yet its cargo was not specifically characterized (108). Small EVs are regarded a tool of cell communication and their cargo is comprised of various bioactive molecules such as nucleic acids, lipids and the here investigated proteins. The focus of our analysis was to specifically investigate the NP sEV proteome for content and anticipated function in comparison to other sEV sources such as autologous AF and adipose cells, fetal NP cells and MSC. Our interpretation relays on multiple functional enrichment and pathway tools (DAVID, ToppFun, KEGG, STRING) with KEGG being more focused on disease related databases, ToppFun most up to date in its annotations and STRING considering known and predicted protein/protein interactions.

The bovine coccygeal IVD is an accepted model for a young, healthy human IVD (148). While bovine sEVs might show species and/or tissue specific variations that could affect their makeup and functionality the extensive overlap with existing sEV data is supportive. The majority of sEV proteins identified here were contained in the ExoCarta and Vesiclepedia databases. A core sEV proteome of 1212 proteins from a range of parent cells was established previously (133) and the ExoCarta database contained >5.3× as many identified sEV proteins by now (124). Despite much of the existing data being based on body fluids such as urine and serum, or cancer cell lines, we found significant overlap and only few proteins, previously not associated with sEVs. While we have not discovered any novel NP biomarkers based on their sEV proteome, we believe that the NP sEV proteome cargo makes important contributions to NP cell and tissue homeostasis (Figure 9)

**Figure 9:**
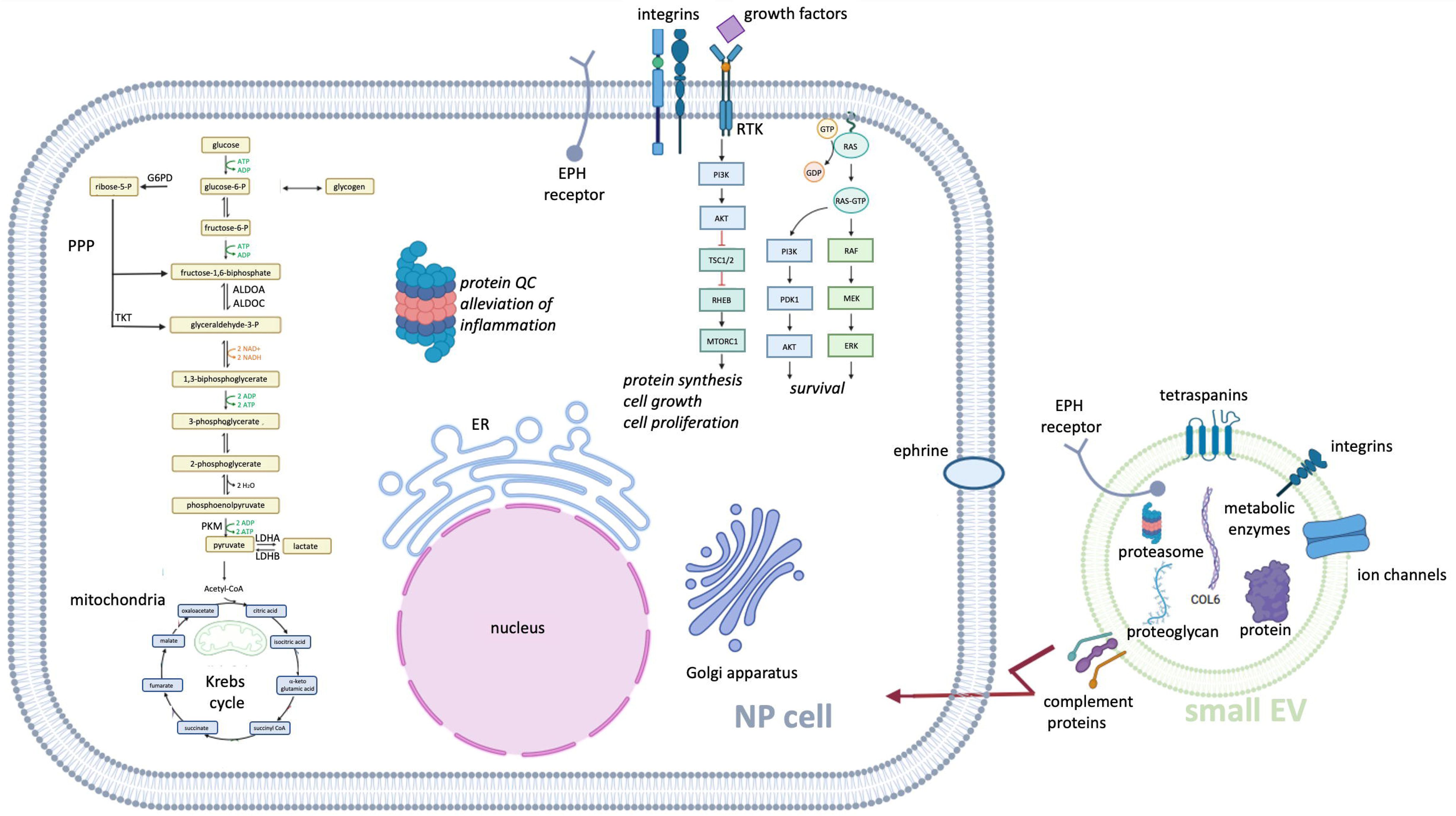
Multifaceted roles of the sEV proteome in NP cell and tissue homeostasis. NP sEV protein cargo and membrane constituents are involved in key metabolic pathways, including glycolysis, gluconeogenesis, Krebs cycle and PPP, crucial for energy production and to maintain the cellular redox balance. Small EVs with the aid of the 20S proteasome, ensure protein quality control and reduced inflammation in a recipient NP cell. Small EVs modulate signaling through EPH receptors, impacting cellular communication and tissue organization. Additionally, sEV interact with the complement system, influencing the classical, alternative, and lectin pathways involved in immune responses and inflammation. The PI3K/AKT/RAS signaling pathway is axis is impacted by the NP sEV proteome, which could promote ECM synthesis, cell growth and proliferation. Together, these processes underscore the essential role of the NP sEV proteome in sustaining NP cell function and NP niche homeostasis. ECM: Extracellular matrix; EPH: Ephrin receptor, ER: endoplasmic reticulum, ERK: extracellular signal-regulated kinase, MEK: mitogen-activated protein kinase, NP: nucleus pulposus, PDK1: phosphoinositide-dependent protein kinase 1, PPP: pentose phosphate pathway, PI3K/AKT: phosphoinositide 3-kinase/protein kinase B, RAF: rapidly accelerated fibrosarcoma, RHEB: RAS homolog enriched in brain, mTORC1: mammalian target of rapamycin complex 1, sEV: small extracellular vesicles, TSC1/2: tuberous sclerosis proteins 1 and 2. This illustration was created on Biorender. (www.biorender.com/).

### 4.1 Small EVs Engage with Cell Metabolism

Disc homeostasis depends on proper metabolism in the NP, which is linked to all cellular activities, and connected to various cellular dysfunctions, including degenerative changes such as IVDD (154). The primary energy source for hypoxic NP cells is glucose, which can generate ATP without oxygen (155). Without glucose, NP cells in culture will lose viability (156). Glucose is transported into the cell using glucose transporters to undergo glycolysis, resulting in the production of pyruvate (157,158). Degenerative changes in the CEPs or the AF could impact nutrient supply, resulting in cell apoptosis and further progression of IVDD (154). Cause-effect relationships between metabolic pathways, NP function and the utilization of other essential nutrients, such as amino acids and fats in the NP remain under investigation (154,159). Proteome profiling indicated a role of NP sEVs in glycolysis, gluconeogenesis, PPP and glutathione metabolism. Glutathione was previously considered as a potential treatment option for IVDD as it protects the IVD from oxidative stress (154,160). Glycolytic enzymes were identified by sEV proteomic studies (161). While these enzymes were not overrepresented among NP sEV proteins, we found several in our profiling data sets from fetal and adult NP parent cells, including LDHA, which has high affinity for pyruvate and preferentially converts pyruvate to lactate as an important step to restore the redox balance (162). GAPDH, widely regarded as a housekeeping gene, fulfills diverse functions such as influencing cell fate by regulating cell death (163). The contribution of functional glycolytic enzymes to a recipient cells’ energy and redox balance was suggested as one of the beneficial therapeutic outcomes of MSC sEV (161). In addition to glycolytic enzymes TKT and G6PD were identified as part of the PPP. The presence of PPP enzymes in metastatic ovarian cancer cells was proposed as a prognostic cancer marker (164), however, their presence in sEVs of primary cells here suggests that this is not a reliable diagnostic tool, rather it might enable a form of metabolic coupling between parent and recipient cells (165). Glycolytic intermediates such as glucose-6-phosphate (G6P) are utilized in the PPP to produce NADPH and ribose-5-phosphate, which are crucial for maintaining cellular redox homeostasis and nucleotide synthesis, respectively. The glycolytic intermediate 3-phosphoglyceric acid, is used in the serine biosynthesis pathway, which is essential for various biosynthesis processes, including the production of proteins, nucleotides, and sphingolipids (154,166).

### 4.2 Housekeeping via Proteosome Delivery

NP cells are resilient *in vitro*. The proteasome pathway was linked to sEV proteins of bovine parent cells and the ExoCarta database, generally associated with NP sEVs and also overrepresented in NP sEVs over parent cells. Proteasomes play crucial roles in cellular proteostasis maintaining functional protein quality and removing protein waste (167,168). Proteasome subunits were detected in EVs present in the growth media of mammalian cells and in body fluids (169–184). Studies suggested a parent cell specific encapsulation of proteasome subunits *in vitro* (170,184). The exact mechanism by which proteasomes are targeted to EVs and their specific function within the EV context remains unclear, however EVs facilitate cargo delivery to recipient cells through several mechanisms, including endocytosis, fusion to the plasma membrane, or phagocytosis (170,185,186). Small EV proteasomes could maintain recipient cell proteostasis upon introduction into target cells. Alternatively, sEVs may also facilitate the export of proteasomes into the ECM to act on extracellular targets. For this, the content of the EVs should be emptied into the extracellular space. Studies have shown that lipid-degrading enzymes such as secretory phospholipase A2 and sphingomyelinase can dismantle EVs, liberating proteasomes into the extracellular environment (187). Also inflammatory mediators could facilitate the release of lipid-degrading enzymes in response to cytokine activation, such as IL1, Interferon γ, and TNFα (188). Potential therapeutic benefits of MSC sEVs with functional 20S proteasomes were demonstrated in a mouse model of myocardial ischemia, reducing the extent of damaged heart tissue and lowering levels of misfolded proteins (189). Proteasomes in NP sEVs might be instrumental for NP homeostasis and to alleviate inflammation.

### 4.3 Immunomodulation in an Immune Privileged Environment

The healthy NP is considered immune privileged due to the physical blood-NP barrier established by the dense lamellar structure of the AF and the PG-rich CEPs, resulting in an absence of blood and lymphatic vessels in the healthy adult NP (138). Nerves and vessels however are present in its periphery. Immune suppressive molecular factors like the cell surface death receptor Fas ligand (FASLG) were described for the NP and NP cells *in vivo* (190,191). We could not detect FAS or FASLG in NP cells or NP sEVs generated *in vitro* by proteome analysis, however *FAS* but not *FASLG* transcripts were detected in cultured NP cells previously, alongside transcripts for the migration inhibitory factor (*MIF*) (148).

Small EVs were previously associated with coagulation, immune modulation and the complement system (192,193). Small EV proteins from all sources investigated here associated with pathways of the (innate) immune system and neutrophil degranulation amongst others. Neutrophils are known as the most abundant type of immune cell in the blood. A connection between IVDD and the infiltration of immune cells including different neutrophil subpopulations was described previously (24,194). Neutrophils generate sEVs (195) and sEVs likely diffuse into NP tissue from the periphery (196). However, neutrophils have a short lifespan *in vitro* and their research generally depends on freshly isolated cells from blood (197). The cells cultured here, though of low passage number, were rinsed profoundly prior to exosome collection and unlikely contained neutrophils or their sEVs. The pathway association could therefore be partially artificial due to the detection of serum corona proteins. However, this does not provide sufficient explanation for a persistent association with those pathways after removing the list of FBS associated proteins.

The complement system represents a group of distinct plasma proteins as part of the innate immune response, essentially complementing antibodies with opsonization by acting through three possible pathways: classic, alternative or lectin (198). Complement proteins and related pathways were predominantly associated with overrepresented NP sEV proteins and also present in the ExoCarta data set. While NP cells are not a classic source for complement proteins, dysregulation of complement proteins was previously linked to inflammatory diseases such as IVDD, CEP lesions, osteo- and rheumatoid arthritis and a potential connection to neovascularization and innervation (198). We found complements C5 and C9 protein in NP sEVs but not the cultured parent cells, however we previously detected transcripts for *C1QBP*, *C1S*, *C1R*, *C2*, *C3* and *C8G* in NP cells (148).While the contribution of NP sEVs to the homeostasis in an immune privileged NP is an intriguing concept, the association with the (innate) immune or the complement system could still be attributable to serum proteins from FBS. This is supported by the spontaneous corona formation on sEV and viral surfaces (146). Other generally observed pathway associations such as focal adhesion and regulation of the cytoskeleton could also be an *in vitro* artefact, as NP cells are known to be very motile in 2D culture (199).

### 4.4 Receptor Shuffling for Niche Homeostasis

Conserved during the evolution of domains and species, yet initially viewed as cellular debris, increasing evidence suggests that (s)EVs play a crucial role in the horizontal transfer of molecules between cells and as mediators of cell-cell communication modulating mammalian cell signaling pathways such as PI3K/AKT, MAPK, AMPK, and cAMP which regulate a variety of cellular functions such as proliferation, survival, metabolism, and cell death (200–203).

Vesicle mediated transport and membrane trafficking were among top pathways identified for bovine sEV proteins and listed in the ExoCarta database. Small EVs are likely essential for promoting intercellular communication in the microenvironment of the IVD, facilitating a flow of bioactive molecules among disc cells with impact on cellular processes to preserve tissue homeostasis. Small EVs could be instrumental in maintaining an avascular and aneural environment in the healthy NP. Recent work demonstrated the theoretical possibility for sEVs to travel through the dense ECM meshwork *in vivo* (196) and the described bidirectional exchange of membrane components by multisized vesicles during NP cell and MSC coculture (204) supports sEVs as communication form. More recently, sEVs isolated from NC cell conditioned medium showed transforming properties and sEVs isolated from MSC had positive effects on NP cells (120,205,206). With a size limited capacity of sEVs, it is reasonable to assume that some proteins are actively deposited or removed while others are simply trapped during vesicle biogenesis. Membrane surface proteins might be put in place by the parent cell prior to vesicle release (207). Secretory vesicles such as sEVs are an important source of cell receptors (208). Small EVs can interact with their target cells through cell surface receptors and signaling cascades, fusion with the target cell or via endocytosis (209). Several of the here identified NP sEV proteins associated with EPH-RTK signaling (210,211). EPH-ephrin domain compositions and complex forward (receptor) and reverse (ligand) signaling events are reviewed in great detail elsewhere (212–214). This signaling pathway is known for its role during embryonic development including the partitioning of the dorsal mesoderm into NC and paraxial mesoderm (215–217), and its’ implication with tissue homeostasis and repair in the adult, maintaining skeletal, hematopoietic and neuronal integrity (213,218). EPH-ephrin interaction typically requires cell-cell contact as receptor and ligand are membrane bound, allowing for uni- or bidirectional signaling (213). Signaling robustness and strength is correlated with the extend of cluster formation (213,219). EPH receptor and ephrin ligand transcripts were detected in NP cells *in vitro* (148,211), however only EPH receptors were identified in the NP sEV proteome. Endocytosis of EPH-ephrin complexes as a mean to regulate signaling was described (220). Though EPH-ephrin signaling in the adult lacks behind its understanding during embryogenesis, it could be relevant for IVD homeostasis that this pathway is reestablished to control stem cell niches; cell homeostasis; tissue boundary formation; cell proliferation, differentiation and migration; metabolism and the cytoskeleton with an antiproliferative effect on the neural stem cell niche, inhibiting neurogenesis beyond just impacting axon guidance as reviewed in (213,218). Furthermore EPH-ephrin, NOTCH and VEGF signaling play important roles in angiogenesis and arteriovenus patterning through the control of cell fate decision (221). Here we found an association between proteins of sEVs from NP parent cells and EPH-receptors. We therefore propose the following potential mechanisms: A) “Down-tuning” of signals in NP parent cells: Studies have suggested a function for EVs in removing undesired compounds, as such contributing to cellular homeostasis by protecting cells (222,223). Packaging EPH receptors into sEV membranes could remove them from NP parent cells and prevent excessive proliferation in the presence of plentiful growth factors in culture medium, which could otherwise trigger non-ligand induced forward signaling (211,212). B) “Amplifying” signals in target cells: On the contrary, shuffling EPH receptors to NP target cells could intensify a downstream forward signaling response in target cells. C) “Avoiding consequences” by inducing reverse signaling in target cells without a forward response in the parent cell and circumventing the requirement for close cell contact by enabling a unique form of long-range communication still based on membrane contact (219).

In addition to EPH receptors, we identified many components of the RAS signaling cascades. It was suggested that RAS downstream signaling is important for sEV biogenesis, release, maintenance and signaling. RAS engagement in RTK signaling can lead to cell fate decisions between survival and death, essentially through the activation of the ERK/MAPK or PI3K/AKT pathways (214,224,225). Other RTKs were identified as well, such as VEGFR2 which can activate downstream PI3K/AKT and mammalian target of rapamycin (MTOR) signaling, impacting angiogenesis via cell proliferation and survival (226). The RTK epidermal growth factor receptor (ERBB1) downstream pathways can initiate the mitogen-activated protein kinase (MAPK) cascade or PI3K/AKT signaling (227). The PI3K/AKT signaling pathway, can influence cell proliferation, apoptosis, autophagy, and differentiation under physiological and pathological conditions by interacting with a number of downstream target proteins, including MTOR and forkhead box O1 (FOXO1) (31,228). Also triggered by RAS signaling events, the MAPK/extracellular signal-regulated kinase (ERK) pathway is an essential intracellular signal transduction pathway that is critical in maintaining IVD homeostasis (229–233).

### 4.5 Promoting ECM Homeostasis

Harnessing “stemness” is an intriguing approach taken by the field of regenerative medicine to treat degenerative diseases or tissue loss. This can include the transplantation of transdifferentiated somatic cells (TDSC), induced pluripotent stem cells (iPSC) or embryonic stem cells (ESC), all though posing a risk of tumorigenesis. Additionally, some cell types are deemed uneconomical on an individualized basis, are not fully understood in their differentiation potential, or face ethical concerns (234–237). Autologous or allogeneic MSC gained popularity since their less-tumorigenic multipotent potential might be directed into the appropriate cell type via endogenous cues from the recipient tissue. However, transplanted stem cells often face delivery and survival challenges, especially in tissues with a naturally harsh micro niche environment such as the IVD. MSC likely signal to resident stem cell populations to kick-start refurbishment via sEVs (206,238–240). Given the adaption of NP cells to their niche, healthy NP parent cells seem a preferential source for sEV production. However, no striking association of NP sEV proteins with stemness were identified here beyond a link to longevity and cellular senescence. This suggests their main contribution is the maintenance of cell and tissue homeostasis in the NP.

During IVDD progression, not only is the expression of pro-inflammatory markers and ECM degrading enzymes noted, but also the exhaustion of progenitor cells. At the same time ECM synthesis reflected by structural collagens, aggrecan and small proteoglycans declines as complex events of cellular senescence and regulated cell death rise (27,34,36,241–247). ECM components like fibronectin connect with cell adhesion receptors and integrins to facilitate cell-cell and cell-matrix interactions, which too have an impact on cell differentiation, proliferation, and function (248–253). Integrin receptors too possess unique bidirectional signaling properties, coordinating extracellular events with intracellular changes (254,255). Integrin can bind with laminin, a component of the ECM activating PI3K/AKT signaling (256–265). Fibronectin-associated with EVs plays a role in maintaining pluripotency and stemness of embryonic stem cells (ESC) (266). Moreover, fibronectin is a significant heparan sulfate ligand, mediating cell-sEV interactions (267). Heparan sulfate proteoglycan 2 (HSPG2), an overrepresented NP sEV protein, plays a crucial role in facilitating sEV uptake by cells. The stiffness of the ECM can enhance integrin signaling and activate the PI3K/AKT signaling pathway (266–268). Fibronectin fragments increase with IVDD progression (269,270). These fragments may contribute to the disease by inducing ECM degradation while suppressing PG synthesis (190,269,271–274).

Collagen VI (COLVI) which was overrepresented in NP sEVs is a multifaceted protein associated with ECM organization, nervous system development, focal adhesion and PI3K/AKT-MTOR signaling, maintaining tissue and cell homeostasis (275). It provides tensile strength and structural integrity to the ECM, and is involved in regulating apoptosis, reducing oxidative stress, and maintaining cell stemness (276). By modulating signaling cascades involved in programmed cell death, COLVI maintains tissue integrity and function (275,277). Its antioxidative properties are responsible for scavenging reactive oxygen species (ROS), thereby protecting cells from oxidative damage (278,279). In stem cell niches, COLVI maintains stemness by providing a supportive microenvironment. This regulatory role ensures the maintenance of stem cell populations and supports tissue regeneration processes (275,277). By modulating signaling pathways involved in cell cycle regulation and ECM proliferation, COLVI promotes tissue homeostasis (277,280). Through interactions with fibronectin, laminin and other ECM molecules, COLVI influences cell-ECM interactions, thereby modulating cellular behavior and tissue function (281–283).

We have investigated sEV proteins from NP and other autologous parent cell sources, fetal NP tissue and MSC derived data. While there is a large overlap in function, we suggest that NP sEVs proteins contribute in particular to niche homeostasis in the NP by maintaining the unique avascular, aneural and immune privileged environment through anti-angiogenic and axon growth inhibitory processes. Lastly, we believe that in the metabolically challenged environment of the NP, sEVs provide resources for an effective redox, energy and pH balance through the delivery of key metabolic enzymes. The repeated association with EPH-RTK signaling, suggests that NP sEVs serve as a tool to regulate cell surface membrane receptor densities and therefore the transduction of extracellular signals to the NP cell. Small EVs from NP parent cells likely also deliver proteasome subunits, which could be of great importance in regulating crucial signaling cascades such as RUNX, NOTCH4, P53, PTEN, Hedgehog and others. We further propose that sEV from NP parent cells contribute to NP cell metabolism as well as NP cell and niche homeostasis beyond MSC.

## Conclusions

This work investigated the composition and possible function of the bovine sEV proteome with a focus on NP parent cells. Small EV research in the IVD recently gained momentum due to advantages over cell based regenerative therapies. Proteome profiling of IVD derived sEVs by state-of-the-art mass spectrometry enables the elucidation of proteome related molecular mechanisms behind cell and tissue homeostasis in this unique organ. However, sEVs can also impact on a transcriptome level that was not assessed here. Based on our findings, there is no obvious indication that the NP sEV proteins identified under these standard culture conditions mobilize progenitor cells other than through the RAS signaling cascade; neither is there any reason to believe they initiate senescence. More interestingly, the NP sEV proteome appears to mediate NP niche and ECM homeostasis including cell fate decisions. Proteins of NP sEVs, whether enriched or not, engaged with major cellular events such as metabolism, pH and redox balance, and signal transduction in particular through the EPH-RAS-PI3K/AKT axis, affecting NP niche homeostasis via axon guidance and angiogenesis.

## Supporting information

Supplementary Figure 1: Original uncropped Western blot membranes. Cropped areas as displayed in Figure 2C (CD63) and Figure 2 E (TSG101) are indicate

Supplementary Figure 2: A) Biological processes (BP); (B) molecular functions (MF) and (C) cellular components (CC) associated with 102 shared sEVs p

Supplementary Figure 3: A) Biological processes (BP); (B) molecular functions (MF) and (C) cellular components (CC) associated with 206 shared sEV pro

Supplementary Figure 4: Quantitative functional enrichment analysis of 141 proteins overrepresented in NP sEVs as compared to their parent cells. A) B

Supplementary Figure 5: Venn diagrams identifying sEV proteins from NP parent cells participating in multiple pathways: A) Cell signaling pathways su

Supplementary Figure 6: Venn diagrams identifying shared ToppFun pathways for the top 200 pathways associated with the different sEV protein sources.

Supplementary Figure 7: Comparison of fetal bovine serum proteins and sEV proteins by various NP parent cell sources. A) Comparison of sEV proteins f

Supplementary Table 1: Protein Concentration of sEV Fractions

Supplementary Table 2: Identified sEV Proteins

Supplementary Table 3: ToppFun functional enrichment analysis for pathways associated with sEV proteins versus ExoCarta and/or Vesiclepedia datasets s

Supplementary Table 4: ToppFun analysis of shared pathways between sEV proteins from autologous bovine NP, AF, and FAT parent cells. AF: Annulus fibro

Supplementary Table 5: ToppFun analysis of shared pathways between sEV proteins from bovine adult and fetal NP, and UCMSC parent cells. bta: Bos tauru

Supplementary Table 6: ToppFun pathway analysis of NP over- or underrepresented sEV proteins.

Supplementary Table 7: Comparison of top 200 results of ToppFun pathways for each sEV source identifies nine pathways shared by all sources

Supplementary Table 8: ToppFun pathway analysis for fetal bovine serum (FBS) proteins and all NP sEV proteins after FBS protein removal.

## List of Abbreviations

AF: annulus fibrosus
BLAST: basic local alignment search tool
BP: biological processes
CC: cellular components
CEP: cartilaginous endplates
CLC: chondrocyte like cells
DAVID: database for annotation, visualization, and integrated discovery
DLS: dynamic light scattering
DMEM: Dulbecco’s modified Eagles medium
DUC: differential ultracentrifugation
ECM: extracellular matrix
EPH: Ephrin receptor
ESCRT: endosomal sorting complex
EV: extracellular vesicle
FAT: adipose tissue
FBS: fetal bovine serum
FC: fold change
FDR: false discovery rate
GO: gene ontology
IVD: intervertebral disc
IVDD: intervertebral disc degeneration
KEGG: Kyoto Encyclopedia of genes and genomes
LBP: low back pain
LC-MS/MS: liquid chromatography-tandem mass spectrometry
MCL: Markov clustering algorithm
MF: molecular function
MSC: mesenchymal stem cell
NC: Notochord
NP: nucleus pulposus
p: p-value
PG: proteoglycan
PID: Pathway interaction database
PPP: pentose phosphate pathway
PTM: posttranslational modification
RTK: receptor tyrosine kinase
SA: senescence associated
SEM: scanning electron microscopy
sEV: small extracellular versicle
STRING: search tool for the retrieval of interacting genes/proteins
UCMSC: umbilical cord mesenchymal stem cells
WB: Western blot

For gene/protein symbol abbreviation please visit gene cards (https://www.genecards.org) or the National Center of Biotechnology Information (NCBI) (https://www.ncbi.nlm.nih.gov)

## Declarations

### Ethics Approval and Consent to Participate

No human materials or subjects were used, no IRB required. No live animals were used. The work has been reported in line with the ARRIVE guidelines 2.0 and is covered under IACUC protocol/approval number 19-04, approved 10/24/2021. Project title: Analysis of Gene Expression. Institution: Clarkson University.

### Consent for Publication

Not applicable.

### Author’s contributions

This project was conceived by PK and carried out in the laboratory of TL. PK, AS, JK and MJY conducted experiments and data analysis. All authors contributed to writing of the manuscript.

### Artificial Intelligence (AI)

The authors declare that they have not use AI-generated work in this manuscript.

## Acknowledgements

We are grateful to Shane Rogers for access to an ultracentrifuge, Daniel Andreescu for access to DLS, Hubert Bilan for expertise in SEM, the Darie lab for preliminary MS, as well as Peter Braun of Woodcrest Dairy (Lisbon, NY), Tritown Meat Packing (Brasher Falls, NY) and Willard & Sons (Heuvelton, NY, USA) for bovine specimens. We thank Sina Lufkin for the help with figures. We also thank the Interdisciplinary Center for Biotechnology Research (ICBR, RRID: SCR_01951) for their invaluable support.

