## Supplementary figures and images for "Small Extracellular Vesicles by Nucleus Pulposus Cells Maintain Niche and Cell Homeostasis via Receptor Shuffling and Metabolic Enzyme Supplements"

### Supplementary Figure 1: Original uncropped Western blot membranes. Cropped areas as displayed in Figure 2C (CD63) and Figure 2 E (TSG101) are indicate

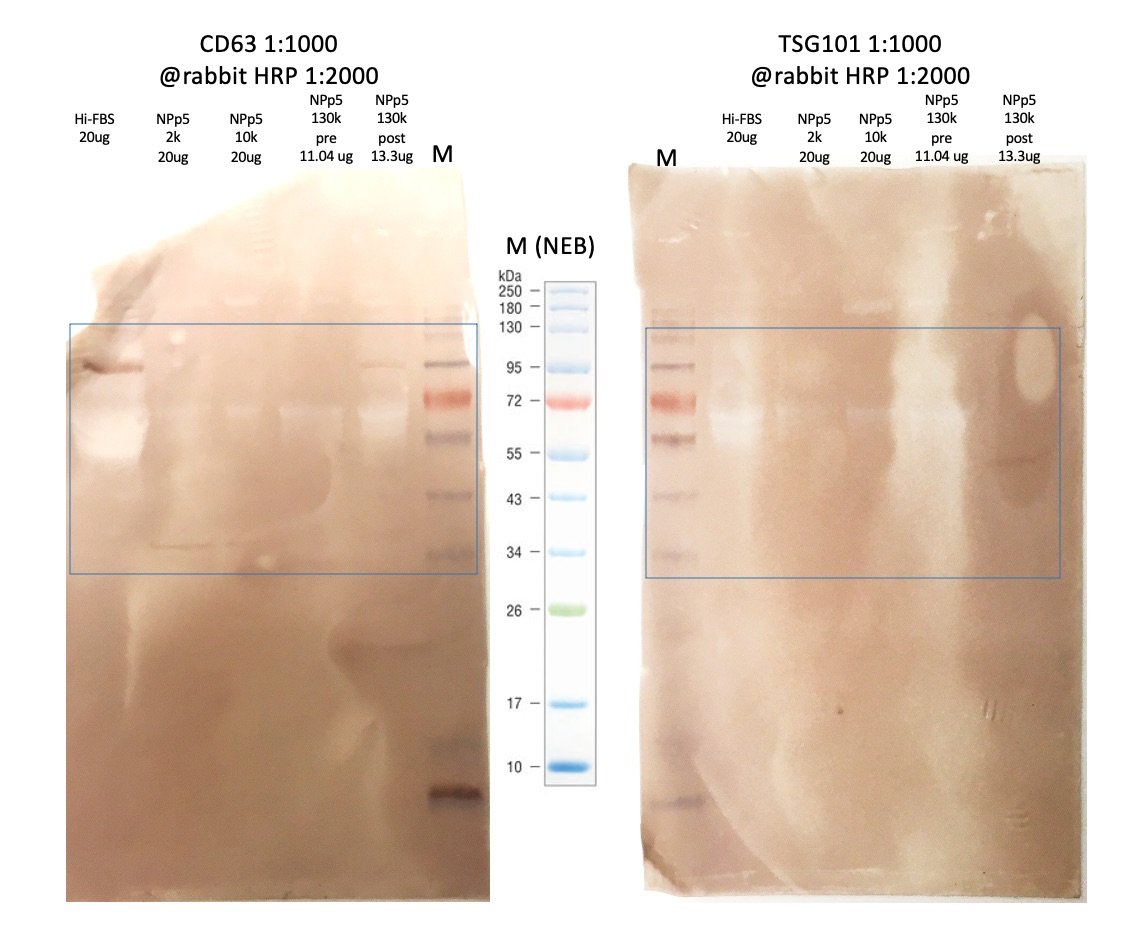

### Supplementary Figure 2: A) Biological processes (BP); (B) molecular functions (MF) and (C) cellular components (CC) associated with 102 shared sEVs p

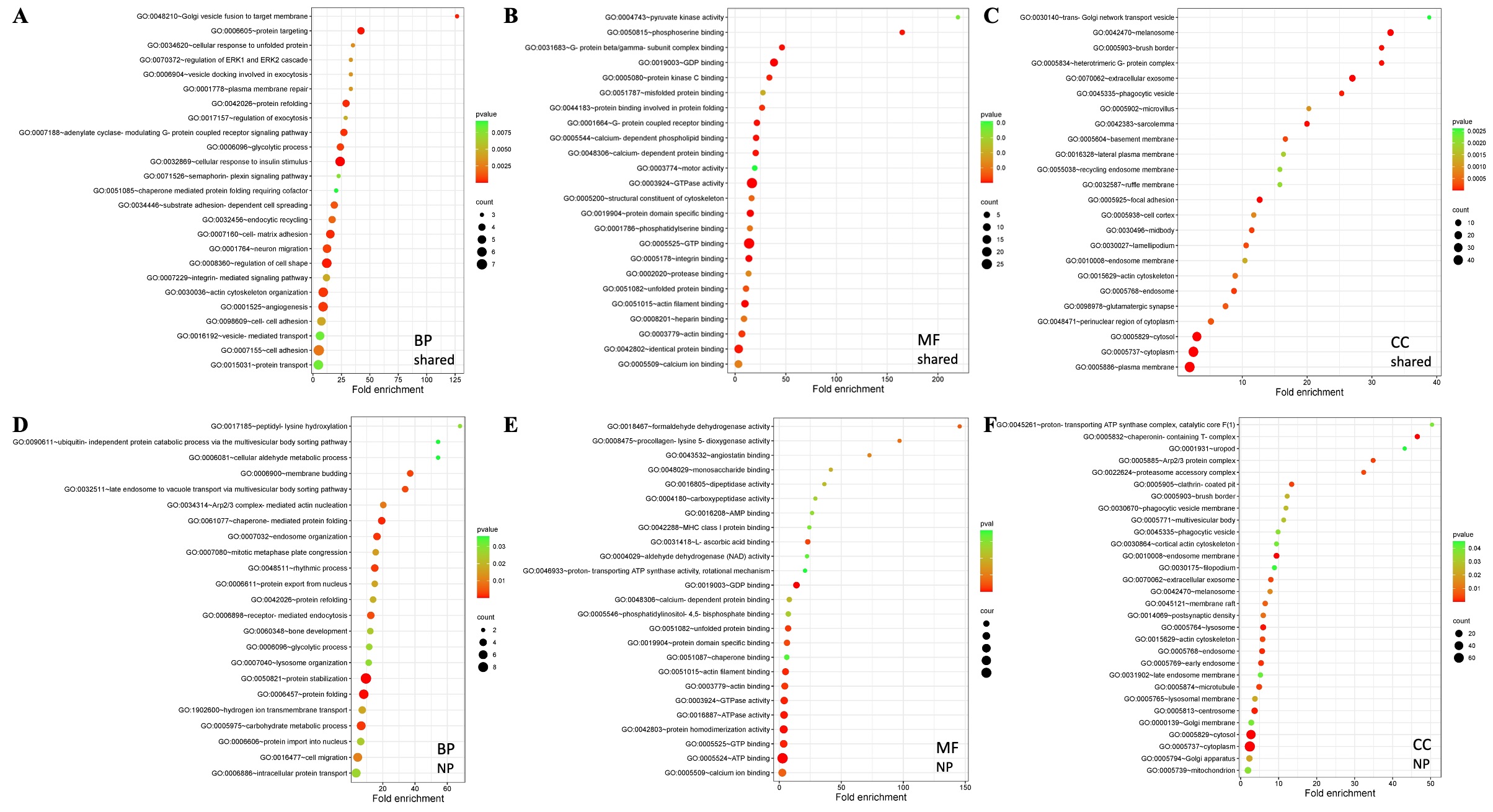

### Supplementary Figure 3: A) Biological processes (BP); (B) molecular functions (MF) and (C) cellular components (CC) associated with 206 shared sEV pro

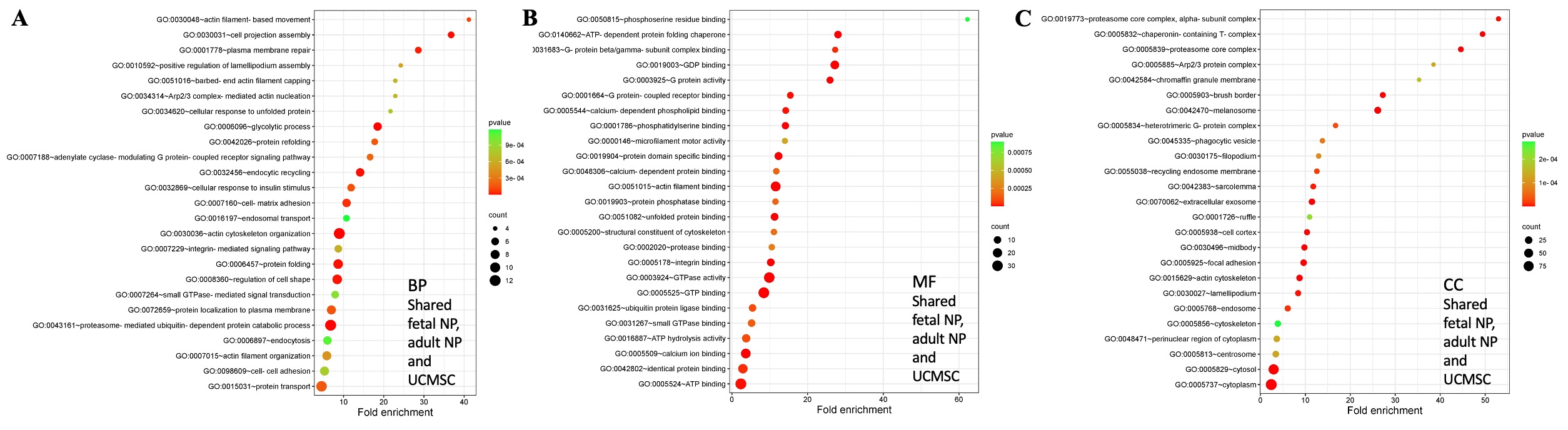

### Supplementary Figure 4: Quantitative functional enrichment analysis of 141 proteins overrepresented in NP sEVs as compared to their parent cells. A) B

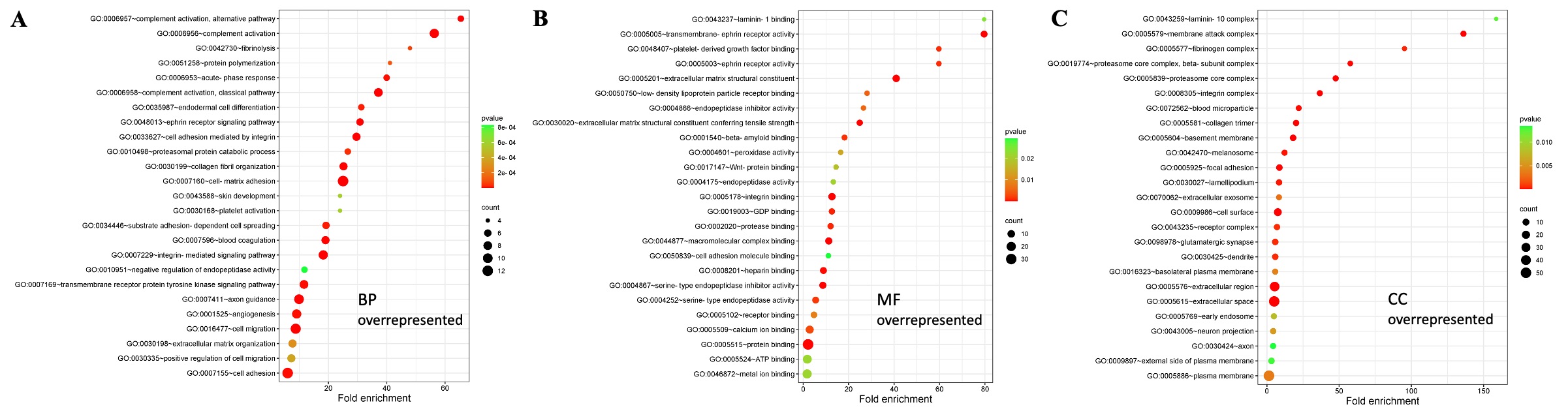

### Supplementary Figure 5: Venn diagrams identifying sEV proteins from NP parent cells participating in multiple pathways: A) Cell signaling pathways su

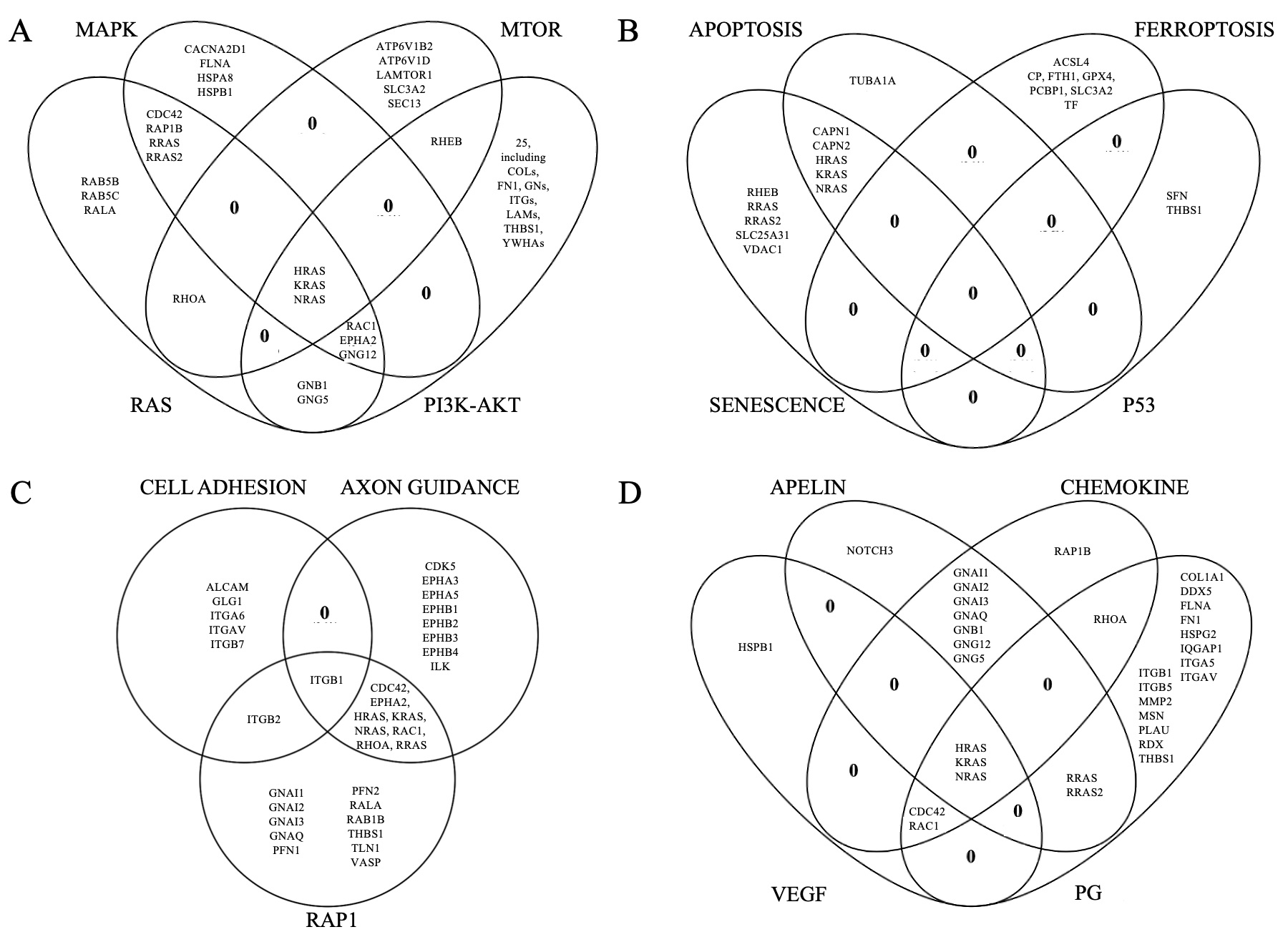

### Supplementary Figure 6: Venn diagrams identifying shared ToppFun pathways for the top 200 pathways associated with the different sEV protein sources.

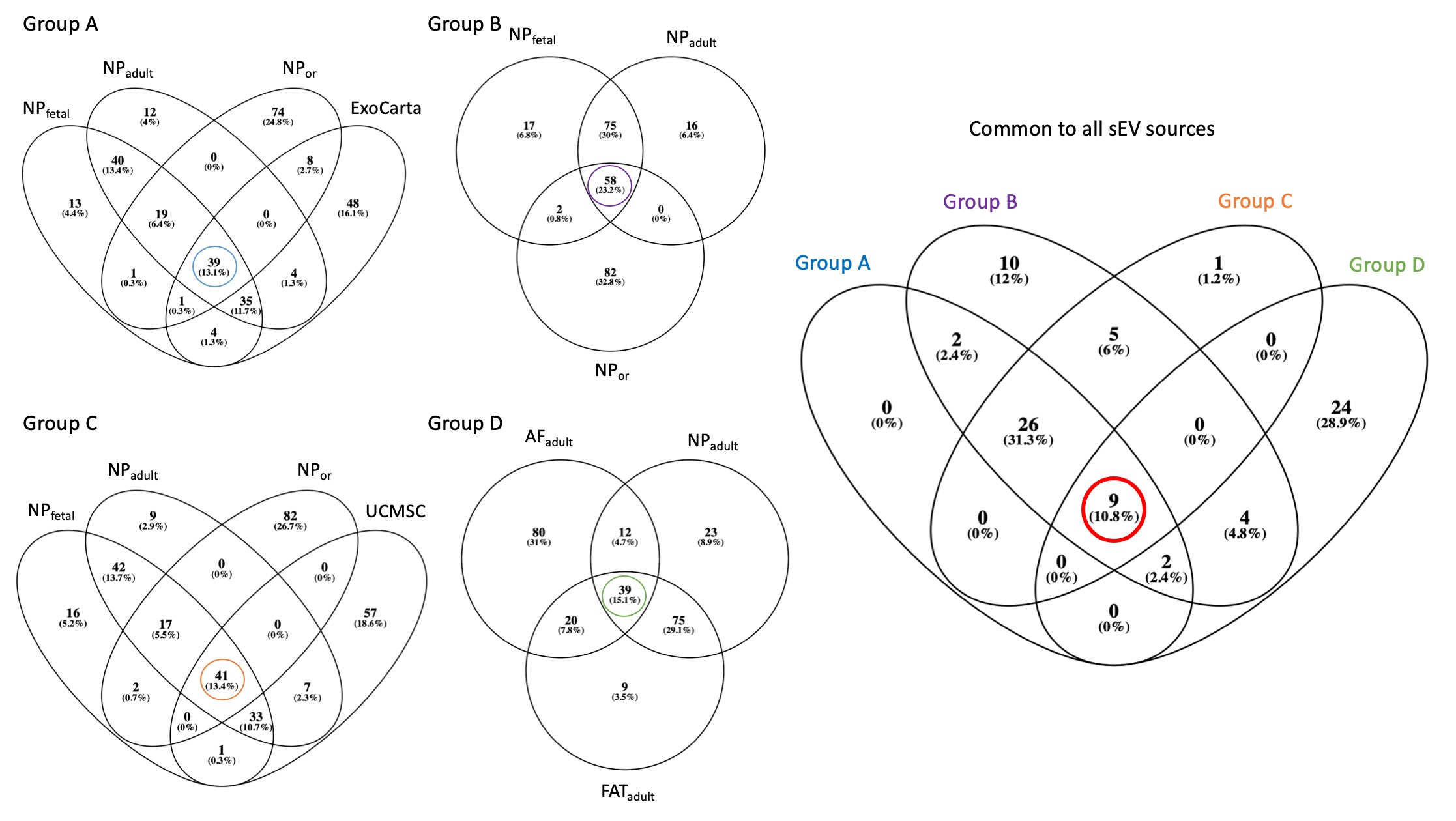

### Supplementary Figure 7: Comparison of fetal bovine serum proteins and sEV proteins by various NP parent cell sources. A) Comparison of sEV proteins f

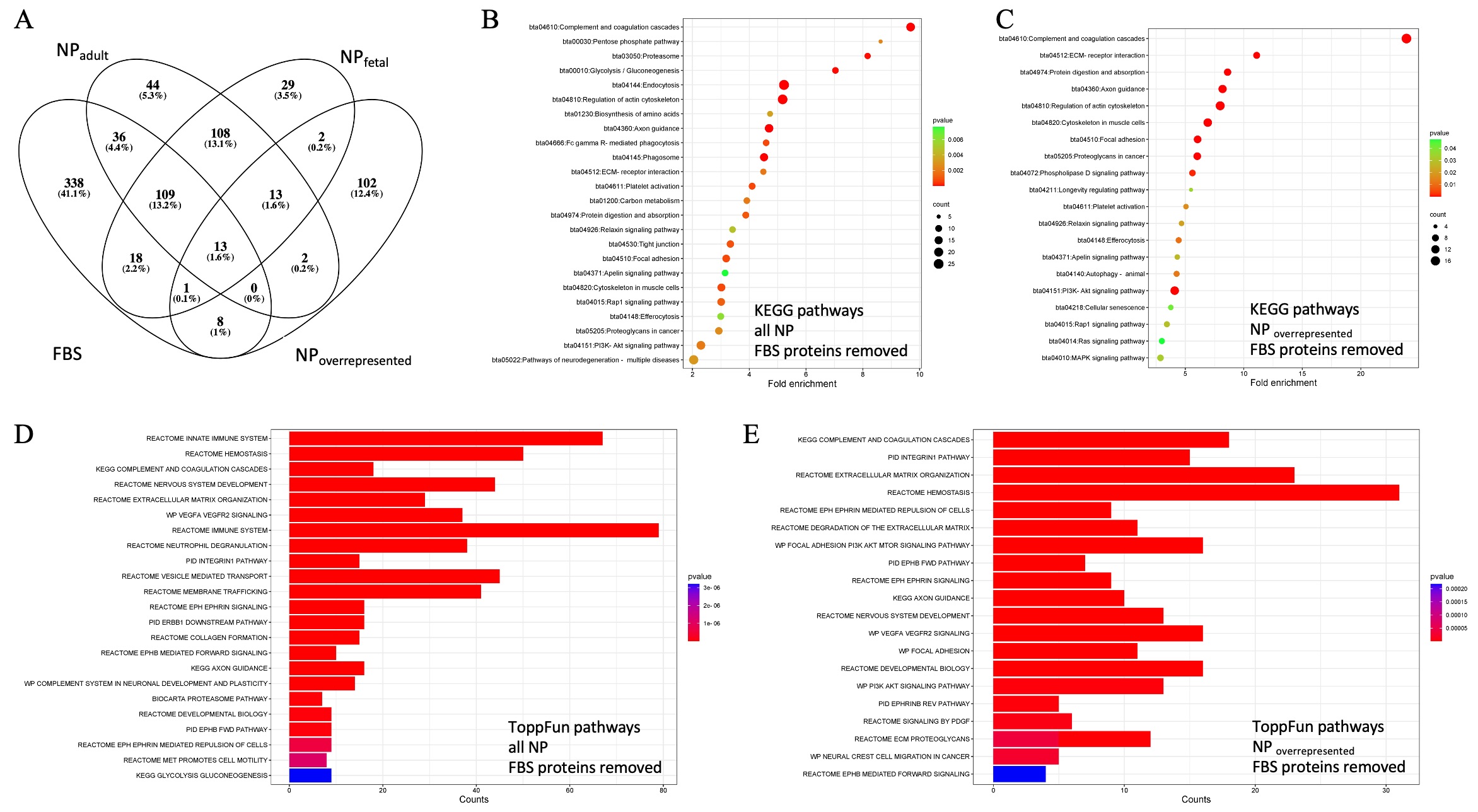
