## Supplementary Table 1: Protein Concentration of sEV Fractions for "Small Extracellular Vesicles by Nucleus Pulposus Cells Maintain Niche and Cell Homeostasis via Receptor Shuffling and Metabolic Enzyme Supplements"

| **Cell Line** | **ng/cell** |
| --- | --- |
| NP (TT39) p5 | 7.72x10^-4^ |
| NP (TT32) p4 | 3.25x10^-3^ |
| NP (TT33) p4 | 2.68x10^-3^ |
| AF (TT39) p6 | 4.7x10^-4^ |
| FAT (TT39) p12 | 2.27x10^-3^ |
| fetal NP p6 | 9.34x10^-3^ |
