## Supplementary Table 2: Identified sEV Proteins for "Small Extracellular Vesicles by Nucleus Pulposus Cells Maintain Niche and Cell Homeostasis via Receptor Shuffling and Metabolic Enzyme Supplements"

| all bovine sEV proteins | all bovine sEV proteins versus ExoCarta and/or Vesiclepedia | | adult profiling | | | fetal profiling | NP sEV enrichment | | pathways |
| --- | --- | --- | --- | --- | --- | --- | --- | --- | --- |
|  | 479 shared | 7 bovine sEV proteins | AF sEV proteins | NP sEV proteins | FAT sEV proteins | NP fetal sEV proteins | over-represented sEV proteins | under-represented sEV proteins | all NP sEV proteins |
| ABI1 | ABI1 | ATP6V0E2 | YWHAH | YWHAH | YWHAH | ABI1 | ADAM10 | ACTN1 | ABI1  |
| ACLY | ACLY | H2AB2 | ACTBL2 | ACLY | ACLY | ACLY | ADIPOQ | CLIC1 | ACLY |
| ACO1 | ACO1 | MBL | ACTN1 | ACO1 | ACTBL2 | ACO1 | AFP | DAD1 | ACO1 |
| ACP1 | ACP1 | NARS1 | ACTR2 | ACP1 | ACTN1 | ACTBL2 | AGT | GOT2 | ACP1 |
| ACSL4 | ACSL4 | PLS2 | AHNAK | ACSL4 | ACTR1A | ACTN1 | ALCAM | LDHAL6B | ACSL4 |
| ACTBL2 | ACTBL2 | RARS1 | ALCAM | ACTBL2 | ACTR3 | ACTR1A | APOB | OAT | ACTBL2 |
| ACTN1 | ACTN1 | STOML1 | ALDOA | ACTN1 | AHNAK | ACTR2 | APOE | RPLP2 | ACTN1 |
| ACTR1A | ACTR1A |  | ANXA1 | ACTR1A | AKR1A1 | ACTR3 | ATP1B1 | RPN2 | ACTR1A |
| ACTR2 | ACTR2 |  | ANXA2 | ACTR2 | ALB | AHCYL1 | ATP2B2 | SLC25A3 | ACTR2 |
| ACTR3 | ACTR3 |  | ANXA4 | ACTR3 | ALCAM | AHNAK | C1QA | WARS1 | ACTR3 |
| ADAM10 | ADAM10 |  | ANXA5 | ADGRV1 | ALDOA | ALB | C1QC |  | ADAM10 |
| ADGRV1 | ADGRV1 |  | ANXA6 | ADH5 | ANGPTL2 | ALCAM | C3 |  | ADGRV1 |
| ADH5 | ADH5 |  | APOE | AHCYL1 | ANXA1 | ALDH1L1 | C4 |  | ADH5 |
| ADIPOQ | ADIPOQ |  | ARF1 | AHNAK | ANXA11 | ALDH9A1 | C5 |  | ADIPOQ |
| AFP | AFP |  | BASP1 | AKR1A1 | ANXA2 | ALDOA | C6 |  | AFP |
| AGT | AGT |  | C9 | ALB | ANXA4 | ANGPTL2 | C7 |  | AGT |
| AHCYL1 | AHCYL1 |  | CAPG | ALCAM | ANXA5 | ANXA1 | C8A |  | AHCYL1 |
| AHNAK | AHNAK |  | CAPZB | ALDH3B1 | ANXA6 | ANXA11 | C8G |  | AHNAK |
| AKR1A1 | AKR1A1 |  | CCT2 | ALDH9A1 | ANXA7 | ANXA13 | C9 |  | AKR1A1 |
| ALB | ALB |  | CHMP1B | ALDOA | APOE | ANXA2 | CACNA2D1 |  | ALB |
| ALCAM | ALCAM |  | CHMP2A | ALDOC | ARF1 | ANXA3 | CARMIL1 |  | ALCAM |
| ALDH1L1 | ALDH1L1 |  | CLIC1 | ANGPTL2 | ARHGDIB | ANXA4 | CD5L |  | ALDH1L1 |
| ALDH3B1 | ALDH3B1 |  | CLIC4 | ANXA1 | ARL8A | ANXA5 | CLU |  | ALDH3B1 |
| ALDH9A1 | ALDH9A1 |  | COL15A1 | ANXA11 | ATP6V1B2 | ANXA6 | COL11A1 |  | ALDH9A1 |
| ALDOA | ALDOA |  | COL6A1 | ANXA2 | ATP6V1D | ANXA7 | COL12A1 |  | ALDOA |
| ALDOC | ALDOC |  | COL6A3 | ANXA3 | BASP1 | APOE | COL1A1 |  | ALDOC |
| ANGPTL2 | ANGPTL2 |  | CROCC | ANXA4 | C5 | APRT | COL2A1 |  | ANGPTL2 |
| ANXA1 | ANXA1 |  | CSE1L | ANXA5 | C9 | ARF1 | COL3A1 |  | ANXA1 |
| ANXA11 | ANXA11 |  | CLTC | ANXA6 | CAPN2 | ARHGDIB | COL5A2 |  | ANXA11 |
| ANXA13 | ANXA13 |  | CYSTM1 | ANXA7 | CCT2 | ARL8A | COL6A2 |  | ANXA13 |
| ANXA2 | ANXA2 |  | DDX19B | AP2M1 | CCT5 | ARPC2 | CPNE3 |  | ANXA2 |
| ANXA3 | ANXA3 |  | DSC2 | APOE | CCT6A | ARPC4 | DIP2B |  | ANXA3 |
| ANXA4 | ANXA4 |  | EEF1A1 | APPL2 | CLIC1 | ARPC5 | DNAJA1 |  | ANXA4 |
| ANXA5 | ANXA5 |  | EEF2 | APRT | CLIC4 | ARRDC1 | DNPEP |  | ANXA5 |
| ANXA6 | ANXA6 |  | EHBP1 | ARF1 | CLTC | ATIC | EMILIN1 |  | ANXA6 |
| ANXA7 | ANXA7 |  | EHD2 | ARHGDIB | COL15A1 | ATP5F1A | ENG |  | ANXA7 |
| AP2M1 | AP2M1 |  | EIF3H | ARL3 | COL6A1 | ATP5F1B | EPB41L2 |  | AP2M1 |
| APOB | APOB |  | FCGBP | ARL8A | COL6A3 | ATP6AP1 | EPHA2 |  | APOB |
| APOE | APOE |  | FLNA | ARPC2 | CROCC | ATP6V0A1 | EPHA3 |  | APOE |
| APPL2 | APPL2 |  | FLNB | ARPC4 | CSE1L | ATP6V1D | EPHA5 |  | APPL2 |
| APRT | APRT |  | FN1 | ARPC5L | CYFIP2 | BASP1 | EPHB1 |  | APRT |
| ARF1 | ARF1 |  | GAPDH | ATIC | CYSTM1 | BGN | EPHB2 |  | ARF1 |
| ARHGDIB | ARHGDIB |  | GDI2 | ATP5F1A | EEF1A1 | C5 | EPHB3 |  | ARHGDIB |
| ARL3 | ARL3 |  | GLIPR2 | ATP5F1B | EEF2 | C9 | EPHB4 |  | ARL3 |
| ARL8A | ARL8A |  | GNAI1 | ATP6AP1 | EHBP1 | CAND1 | F10 |  | ARL8A |
| ARPC2 | ARPC2 |  | GNAI2 | ATP6V0E2 | EHD2 | CAP1 | F2 |  | ARPC2 |
| ARPC4 | ARPC4 |  | GNAI3 | ATP6V1B2 | FAT1 | CAPG | F5 |  | ARPC4 |
| ARPC5 | ARPC5 |  | GNAQ | BASP1 | FLNA | CAPN1 | FAT1 |  | ARPC5 |
| ARPC5L | ARPC5L |  | GNB1 | BGN | FLOT2 | CAPN2 | FAT4 |  | ARPC5L |
| ARRDC1 | ARRDC1 |  | GNG12 | C5 | FN1 | CAPZA2 | FETUB |  | ARRDC1 |
| ATIC | ATIC |  | GSN | C9 | GAPDH | CAPZB | FGA |  | ATIC |
| ATP1B1 | ATP1B1 |  | H2AB2 | CAND1 | GDI2 | CC2D1A | FGB |  | ATP1B1 |
| ATP2B2 | ATP2B2 |  | H2BC18 | CAP1 | GLIPR2 | CCT2 | FGG |  | ATP2B2 |
| ATP5F1A | ATP5F1A |  | H4C4 | CAPG | GNAI1 | CCT3 | FIGNL1 |  | ATP5F1A |
| ATP5F1B | ATP5F1B |  | HRAS | CAPN1 | GNAI2 | CCT5 | FLOT1 |  | ATP5F1B |
| ATP6AP1 | ATP6AP1 |  | HSPA1B | CAPN2 | GNAI3 | CCT6A | FMNL3 |  | ATP6AP1 |
| ATP6V0A1 | ATP6V0A1 |  | HSPA2 | CAPZA2 | GNAQ | CCT8 | FSTL1 |  | ATP6V0A1 |
| ATP6V0E2 | ATP6V1B2 |  | HSPA5 | CAPZB | GNB1 | CDC42 | FTH1 |  | ATP6V0E2 |
| ATP6V1B2 | ATP6V1D |  | HSPA8 | CC2D1A | GNG12 | CEP250 | GC |  | ATP6V1B2 |
| ATP6V1D | BASP1 |  | HSPB1 | CCT2 | GSN | CHMP1B | HBB |  | ATP6V1D |
| BASP1 | BGN |  | HSPG2 | CCT3 | H2AB2 | CHMP2A | HBE1 |  | BASP1 |
| BGN | C1QA |  | IQGAP1 | CCT5 | H2BC18 | CHMP2B | HP |  | BGN |
| C1QA | C1QC |  | IQGAP2 | CCT6A | H4C4 | CHMP4B | HTRA1 |  | C1QA |
| C1QC | C3 |  | ITGA1 | CCT8 | HBB | CLIC1 | IGF2R |  | C1QC |
| C3 | C4 |  | ITGA3 | CDC42 | HSPA1B | CLIC4 | IGSF8 |  | C3 |
| C4 | C5 |  | ITGAV | CDC42BPB | HSPA2 | CLTC | ILK |  | C4 |
| C5 | C6 |  | ITGB1 | CDK5 | HSPA5 | CNDP2 | ITGA3 |  | C5 |
| C6 | C7 |  | ITM2B | CEP250 | HSPA8 | CNTLN | ITGA5 |  | C6 |
| C7 | C8A |  | KIF3B | CHMP2A | HSPB1 | COL15A1 | ITGA6 |  | C7 |
| C8A | C8G |  | LAMC1 | CHMP4B | HSPG2 | COL6A1 | ITGB2 |  | C8A |
| C8G | C9 |  | LAMP1 | CHMP6 | IDH1 | COL6A3 | ITGB5 |  | C8G |
| C9 | CACNA2D1 |  | MARCKS | CHP1 | ITGA3 | CPNE3 | ITGB7 |  | C9 |
| CACNA2D1 | CAND1 |  | MSN | CKAP4 | ITGAV | CPNE5 | ITIH2 |  | CACNA2D1 |
| CAND1 | CAP1 |  | MVP | CLIC1 | ITGB1 | CROCC | ITIH3 |  | CAND1 |
| CAP1 | CAPG |  | MYH9 | CLIC4 | ITM2B | CRYAB | ITIH4 |  | CAP1 |
| CAPG | CAPN1 |  | MYO1C | CLTC | LAMC1 | CSE1L | ITM2B |  | CAPG |
| CAPN1 | CAPN2 |  | MYO1D | CNDP2 | LAMP1 | CUBN | KIF11 |  | CAPN1 |
| CAPN2 | CAPZA2 |  | MYOF | CNTLN | LDHA | CYFIP2 | KRAS |  | CAPN2 |
| CAPZA2 | CAPZB |  | NIBAN2 | COL15A1 | LMAN2 | CYSTM1 | KRT18 |  | CAPZA2 |
| CAPZB | CARMIL1 |  | NT5E | COL6A1 | M6PR | DDAH1 | KRT6B |  | CAPZB |
| CARMIL1 | CC2D1A |  | PDCD6IP | COL6A3 | MARCKS | DDAH2 | KRT79 |  | CARMIL1 |
| CC2D1A | CCT2 |  | PEBP1 | CP | MINK1 | DDB1 | LAMA5 |  | CC2D1A |
| CCT2 | CCT3 |  | PFN1 | CRELD1 | MSN | DDX19B | LAMB1 |  | CCT2 |
| CCT3 | CCT5 |  | PGD | CROCC | MVP | DSTN | LAMC1 |  | CCT3 |
| CCT5 | CCT6A |  | PHGDH | CRYAB | MYH9 | DYNC1H1 | LAMP2 |  | CCT5 |
| CCT6A | CCT8 |  | PKLR | CSE1L | MYO1C | ECE1 | LOXL2 |  | CCT6A |
| CCT8 | CD5L |  | PKM | CSRP1 | MYO1D | EEA1 | MAMDC2 |  | CCT8 |
| CD5L | CDC42 |  | PPIL1 | CUBN | MYOF | EEF1 | MASP1 |  | CD5L |
| CDC42 | CDC42BPB |  | RAB10 | CYFIP2 | NIBAN2 | EEF2 | MBL |  | CDC42 |
| CDC42BPB | CDK5 |  | RAB13 | CYSTM1 | NID1 | EHBP1 | MELTF |  | CDC42BPB |
| CDK5 | CEP250 |  | RAB15 | DCTN2 | NPM1 | EHD2 | MMP2 |  | CDK5 |
| CEP250 | CHMP1B |  | RAB35 | DDAH2 | NT5E | EHD4 | MPZL1 |  | CEP250 |
| CHMP1B | CHMP2A |  | RAB3D | DDB1 | PDCD6IP | EIF3H | MXRA8 |  | CHMP1B |
| CHMP2A | CHMP4B |  | RAB5B | DDX19B | PEBP1 | EPPK1 | NID1 |  | CHMP2A |
| CHMP4B | CHMP6 |  | RAB5C | DDX3X | PFN1 | FASN | NID2 |  | CHMP2B |
| CHMP6 | CHP1 |  | RAB7A | DDX5 | PGD | FAT1 | NOTCH2 |  | CHMP4B |
| CHP1 | CKAP4 |  | RAB8A | DNAJA1 | PHGDH | FCN2 | NOTCH3 |  | CHMP6 |
| CKAP4 | CLIC1 |  | RAB8B | DNAJC13 | PKLR | FIGNL1 | NRAS |  | CHP1 |
| CLIC1 | CLIC4 |  | RAC1 | DNM2 | PKM | FLNA | NRP2 |  | CKAP4 |
| CLIC4 | CLTC |  | RACK1 | DSTN | PLS1 | FLNB | ORM1 |  | CLIC1 |
| CLTC | CLU |  | RALA | DYNC1H1 | PPIL1 | FLOT1 | PLAU |  | CLIC4 |
| CLU | CNDP2 |  | RAN | EEF1A1 | PSMA1 | FLOT2 | PLBD2 |  | CLTC |
| CNDP2 | CNTLN |  | RAP1B | EEF2 | PSMA2 | FN1 | PLXNB2 |  | CLU |
| CNTLN | COL11A1 |  | RAP2B | EHBP1 | PSMA3 | FTH1 | POSTN |  | CNDP2 |
| COL11A1 | COL12A1 |  | RAPGEFL1 | EHD2 | PSMA4 | G6PD | PSMA3 |  | CNTLN |
| COL12A1 | COL15A1 |  | RDX | EIF3H | PSMA5 | GAPDH | PSMA4 |  | COL11A1 |
| COL15A1 | COL1A1 |  | RHOA | EPPK1 | PSMB2 | GDI2 | PSMB1 |  | COL12A1 |
| COL1A1 | COL2A1 |  | RPS11 | EVPL | PSMB5 | GLIPR2 | PSMB2 |  | COL15A1 |
| COL2A1 | COL3A1 |  | RPS18 | FASN | PSMB6 | GNA11 | PSMB4 |  | COL1A1 |
| COL3A1 | COL5A2 |  | RRAS | FAT1 | RAB10 | GNAI1 | PSMB6 |  | COL2A1 |
| COL5A2 | COL6A1 |  | RRAS2 | FCN2 | RAB13 | GNAI2 | PTGFRN |  | COL3A1 |
| COL6A1 | COL6A2 |  | S100A4 | FIGNL1 | RAB15 | GNAI3 | PTX3 |  | COL5A2 |
| COL6A2 | COL6A3 |  | SDCBP | FKBP4 | RAB35 | GNAQ | PXDN |  | COL6A1 |
| COL6A3 | CP |  | SFN | FLNA | RAB5B | GNB1 | RAC1 |  | COL6A2 |
| CP | CPNE3 |  | SH3 | FLNB | RAB5C | GNG12 | RHEB |  | COL6A3 |
| CPNE3 | CPNE5 |  | SRI | FLOT1 | RAB7A | GNG5 | ROR1 |  | CP |
| CPNE5 | CRELD1 |  | STOML1 | FN1 | RAB8A | GOLGA7 | RPL18A |  | CPNE3 |
| CRELD1 | CROCC |  | TAGLN2 | FTH1 | RAB8B | GPI | RPL28 |  | CPNE5 |
| CROCC | CRYAB |  | TLN1 | FUCA1 | RAC1 | GPX4 | RRAS |  | CRELD1 |
| CRYAB | CSE1L |  | TMEM59 | G6PD | RAN | GSN | RRAS2 |  | CROCC |
| CSE1L | CSRP1 |  | TMEM63A | GAA | RAP1B | H2AB2 | SCAMP3 |  | CRYAB |
| CSRP1 | CUBN |  | TMX1 | GALK1 | RAP2B | H2BC18 | SERPINA3 |  | CSE1L |
| CUBN | CYFIP2 |  | TPI1 | GANAB | RAPGEFL1 | H4C4 | SERPIND1 |  | CSRP1 |
| CYFIP2 | CYSTM1 |  | TTN | GAPDH | RHEB | HBB | SERPINF1 |  | CUBN |
| CYSTM1 | DCTN2 |  | TUBA1A | GDI2 | RHOA | HP | SERPINF2 |  | CYFIP2 |
| DCTN2 | DDAH1 |  | TUBB2A | GET3 | RPS18 | HRAS | SLC16A1 |  | CYSTM1 |
| DDAH1 | DDAH2 |  | TUBB8 | GLB1L3 | RRAS | HSPA1B | SLC1A4 |  | DCTN2 |
| DDAH2 | DDB1 |  | VAT1 | GLG1 | RRAS2 | HSPA2 | SLC29A1 |  | DDAH1 |
| DDB1 | DDX19B |  | VCP | GLIPR2 | S100A16 | HSPA5 | SLC3A2 |  | DDAH2 |
| DDX19B | DDX3X |  | WDR1 | GNA11 | S100A4 | HSPA8 | SLC44A2 |  | DDB1 |
| DDX3X | DDX5 |  | YWHAE | GNAI1 | SDCBP | HSPB1 | SPP2 |  | DDX19B |
| DDX5 | DIP2B |  | YWHAG | GNAI2 | SH3 | HSPD1 | SRI |  | DDX3X |
| DIP2B | DNAJA1 |  | YWHAQ | GNAI3 | SNX18 | HSPG2 | TF |  | DDX5 |
| DNAJA1 | DNAJC13 |  | YWHAZ | GNAQ | STOML1 | IDH1 | TG |  | DIP2B |
| DNAJC13 | DNM2 |  |  | GNB1 | TCP1 | IQGAP1 | TGFBI |  | DNAJA1 |
| DNM2 | DNPEP |  |  | GPI | THBS1 | IQGAP2 | TSG101 |  | DNAJC13 |
| DNPEP | DSC2 |  |  | GRHPR | TLN1 | IST1 | TTYH3 |  | DNM2 |
| DSC2 | DSTN |  |  | GSN | TMX1 | ITGA1 | VASN |  | DNPEP |
| DSTN | DYNC1H1 |  |  | GSTP1 | TUBA1A | ITGA3 | VPS37B |  | DSTN |
| DYNC1H1 | ECE1 |  |  | H2AB2 | TUBB2A | ITGAV | VPS4A |  | DYNC1H1 |
| ECE1 | EEA1 |  |  | H2BC18 | UGP2 | ITGB1 | VPS4B |  | ECE1 |
| EEA1 | EEF1 |  |  | H4C4 | VAMP3 | ITM2B |  |  | EEA1 |
| EEF1 | EEF1A1 |  |  | HBB | VAT1 | JADE2 |  |  | EEF1 |
| EEF1A1 | EEF2 |  |  | HNRNPL | VCL | KIF3B |  |  | EEF1A1 |
| EEF2 | EHBP1 |  |  | HRAS | VCP | KPNB1 |  |  | EEF2 |
| EHBP1 | EHD2 |  |  | HSPA12A | YWHAE | LAMA5 |  |  | EHBP1 |
| EHD2 | EHD4 |  |  | HSPA1B | YWHAG | LAMC1 |  |  | EHD2 |
| EHD4 | EIF3H |  |  | HSPA2 | YWHAQ | LAMTOR1 |  |  | EHD4 |
| EIF3H | EMILIN1 |  |  | HSPA5 | YWHAZ | LDHA |  |  | EIF3H |
| EMILIN1 | ENG |  |  | HSPA8 |  | LDHB |  |  | EMILIN1 |
| ENG | EPB41L2 |  |  | HSPB1 |  | LGALS3 |  |  | ENG |
| EPB41L2 | EPHA2 |  |  | HSPD1 |  | LMAN2 |  |  | EPB41L2 |
| EPHA2 | EPHA3 |  |  | HSPG2 |  | M6PR |  |  | EPHA2 |
| EPHA3 | EPHA5 |  |  | IDH1 |  | MAN1A1 |  |  | EPHA3 |
| EPHA5 | EPHB1 |  |  | IQGAP1 |  | MARCKS |  |  | EPHA5 |
| EPHB1 | EPHB2 |  |  | IQGAP2 |  | MFGE8 |  |  | EPHB1 |
| EPHB2 | EPHB3 |  |  | ITGA1 |  | MINK1 |  |  | EPHB2 |
| EPHB3 | EPHB4 |  |  | ITGA3 |  | MSN |  |  | EPHB3 |
| EPHB4 | EPPK1 |  |  | ITGAV |  | MVP |  |  | EPHB4 |
| EPPK1 | EVPL |  |  | ITGB1 |  | MXRA8 |  |  | EPPK1 |
| EVPL | F10 |  |  | ITM2B |  | MYH14 |  |  | EVPL |
| F10 | F2 |  |  | ITM2C |  | MYH9 |  |  | F10 |
| F2 | F5 |  |  | ITSN2 |  | MYO1B |  |  | F2 |
| F5 | FASN |  |  | JADE2 |  | MYO1C |  |  | F5 |
| FASN | FAT1 |  |  | KPNB1 |  | MYO1D |  |  | FASN |
| FAT1 | FAT4 |  |  | LAMA5 |  | MYOF |  |  | FAT1 |
| FAT4 | FCGBP |  |  | LAMC1 |  | NAPA |  |  | FAT4 |
| FCGBP | FCN2 |  |  | LAMP2 |  | NARS1 |  |  | FCN2 |
| FCN2 | FETUB |  |  | LAMTOR1 |  | NCKAP1 |  |  | FETUB |
| FETUB | FGA |  |  | LAP3 |  | NIBAN2 |  |  | FGA |
| FGA | FGB |  |  | LDHA |  | NID1 |  |  | FGB |
| FGB | FGG |  |  | LDHB |  | NPM1 |  |  | FGG |
| FGG | FIGNL1 |  |  | LGALS3 |  | NT5E |  |  | FIGNL1 |
| FIGNL1 | FKBP4 |  |  | LMAN2 |  | NUTF2 |  |  | FKBP4 |
| FKBP4 | FLNA |  |  | M6PR |  | OTUB1 |  |  | FLNA |
| FLNA | FLNB |  |  | MAN1A1 |  | PACSIN2 |  |  | FLNB |
| FLNB | FLOT1 |  |  | MAN2B2 |  | PAFAH1B1 |  |  | FLOT1 |
| FLOT1 | FLOT2 |  |  | MARCKS |  | PARK7 |  |  | FLOT2 |
| FLOT2 | FMNL3 |  |  | MINK1 |  | PDCD10 |  |  | FMNL3 |
| FMNL3 | FN1 |  |  | MOGS |  | PDCD6IP |  |  | FN1 |
| FN1 | FSTL1 |  |  | MSN |  | PEBP1 |  |  | FSTL1 |
| FSTL1 | FTH1 |  |  | MTHFD1 |  | PEDF |  |  | FTH1 |
| FTH1 | FUCA1 |  |  | MVP |  | PEF1 |  |  | FUCA1 |
| FUCA1 | G6PD |  |  | MYH14 |  | PFKL |  |  | G6PD |
| G6PD | GAA |  |  | MYH9 |  | PFN1 |  |  | GAA |
| GAA | GALK1 |  |  | MYO1B |  | PGD |  |  | GALK1 |
| GALK1 | GANAB |  |  | MYO1C |  | PGM1 |  |  | GANAB |
| GANAB | GAPDH |  |  | MYO1D |  | PHGDH |  |  | GAPDH |
| GAPDH | GC |  |  | MYOF |  | PKLR |  |  | GC |
| GC | GDI2 |  |  | N4BP2L2 |  | PKM |  |  | GDI2 |
| GDI2 | GET3 |  |  | NAA50 |  | PLD3 |  |  | GET3 |
| GET3 | GLB1L3 |  |  | NAGK |  | PLS1 |  |  | GLB1L3 |
| GLB1L3 | GLG1 |  |  | NAPA |  | PLS2 |  |  | GLG1 |
| GLG1 | GLIPR2 |  |  | NARS1 |  | PNP |  |  | GLIPR2 |
| GLIPR2 | GNA11 |  |  | NCKAP1 |  | PPFIA2 |  |  | GNA11 |
| GNA11 | GNAI1 |  |  | NIBAN2 |  | PPIB |  |  | GNAI1 |
| GNAI1 | GNAI2 |  |  | NID1 |  | PPIL1 |  |  | GNAI2 |
| GNAI2 | GNAI3 |  |  | NONO |  | PPL |  |  | GNAI3 |
| GNAI3 | GNAQ |  |  | NPEPPS |  | PRDX6 |  |  | GNAQ |
| GNAQ | GNB1 |  |  | NPM1 |  | PSAT1 |  |  | GNB1 |
| GNB1 | GNG12 |  |  | NT5E |  | PSMA1 |  |  | GNG12 |
| GNG12 | GNG5 |  |  | NUTF2 |  | PSMA2 |  |  | GNG5 |
| GNG5 | GOLGA7 |  |  | OLA1 |  | PSMA3 |  |  | GOLGA7 |
| GOLGA7 | GPI |  |  | OTUB1 |  | PSMA4 |  |  | GPI |
| GPI | GPX4 |  |  | OXSR1 |  | PSMA5 |  |  | GPX4 |
| GPX4 | GRHPR |  |  | P3H1 |  | PSMB2 |  |  | GRHPR |
| GRHPR | GSN |  |  | PACSIN2 |  | PSMB5 |  |  | GSN |
| GSN | GSTP1 |  |  | PAFAH1B2 |  | PSMB6 |  |  | GSTP1 |
| GSTP1 | H2BC18 |  |  | PCBP1 |  | PSMC6 |  |  | H2AB2 |
| H2AB2 | H4C4 |  |  | PDCD6 |  | PSMD12 |  |  | H2BC18 |
| H2BC18 | HBB |  |  | PDCD6IP |  | PSMD8 |  |  | H4C4 |
| H4C4 | HBE1 |  |  | PEBP1 |  | RAB10 |  |  | HBB |
| HBB | HNRNPL |  |  | PEDF |  | RAB13 |  |  | HBE1 |
| HBE1 | HP |  |  | PFKL |  | RAB14 |  |  | HNRNPL |
| HNRNPL | HRAS |  |  | PFN1 |  | RAB15 |  |  | HP |
| HP | HSPA12A |  |  | PFN2 |  | RAB2A |  |  | HRAS |
| HRAS | HSPA1B |  |  | PGD |  | RAB33B |  |  | HSPA12A |
| HSPA12A | HSPA2 |  |  | PGM1 |  | RAB35 |  |  | HSPA1B |
| HSPA1B | HSPA5 |  |  | PHGDH |  | RAB3D |  |  | HSPA2 |
| HSPA2 | HSPA8 |  |  | PKLR |  | RAB5B |  |  | HSPA5 |
| HSPA5 | HSPB1 |  |  | PKM |  | RAB5C |  |  | HSPA8 |
| HSPA8 | HSPD1 |  |  | PLOD1 |  | RAB7A |  |  | HSPB1 |
| HSPB1 | HSPG2 |  |  | PLOD3 |  | RAB8A |  |  | HSPD1 |
| HSPD1 | HTRA1 |  |  | PLS1 |  | RAB8B |  |  | HSPG2 |
| HSPG2 | IDH1 |  |  | PLS2 |  | RAC1 |  |  | HTRA1 |
| HTRA1 | IGF2R |  |  | PNP |  | RACK1 |  |  | IDH1 |
| IDH1 | IGSF8 |  |  | PPIB |  | RALA |  |  | IGF2R |
| IGF2R | ILK |  |  | PPIL1 |  | RAN |  |  | IGSF8 |
| IGSF8 | IQGAP1 |  |  | PPP1R7 |  | RAP1B |  |  | ILK |
| ILK | IQGAP2 |  |  | PRDX5 |  | RAP2B |  |  | IQGAP1 |
| IQGAP1 | IST1 |  |  | PRDX6 |  | RAPGEFL1 |  |  | IQGAP2 |
| IQGAP2 | ITGA1 |  |  | PRKAR2A |  | RDX |  |  | IST1 |
| IST1 | ITGA3 |  |  | PRXL2B |  | RENBP |  |  | ITGA1 |
| ITGA1 | ITGA5 |  |  | PSAT1 |  | RHEB |  |  | ITGA3 |
| ITGA3 | ITGA6 |  |  | PSMA1 |  | RHOA |  |  | ITGA5 |
| ITGA5 | ITGAV |  |  | PSMA2 |  | RHOG |  |  | ITGA6 |
| ITGA6 | ITGB1 |  |  | PSMA3 |  | RPS11 |  |  | ITGAV |
| ITGAV | ITGB2 |  |  | PSMA4 |  | RPS18 |  |  | ITGB1 |
| ITGB1 | ITGB5 |  |  | PSMA5 |  | RRAS |  |  | ITGB2 |
| ITGB2 | ITGB7 |  |  | PSMB2 |  | RRAS2 |  |  | ITGB5 |
| ITGB5 | ITIH2 |  |  | PSMB3 |  | S100A10 |  |  | ITGB7 |
| ITGB7 | ITIH3 |  |  | PSMB5 |  | S100A14 |  |  | ITIH2 |
| ITIH2 | ITIH4 |  |  | PSMB6 |  | S100A16 |  |  | ITIH3 |
| ITIH3 | ITM2B |  |  | PSMC6 |  | S100A4 |  |  | ITIH4 |
| ITIH4 | ITM2C |  |  | PSMD12 |  | SDCBP |  |  | ITM2B |
| ITM2B | ITSN2 |  |  | PSMD8 |  | SFN |  |  | ITM2C |
| ITM2C | JADE2 |  |  | PWWP3B |  | SH3BGRL3 |  |  | ITSN2 |
| ITSN2 | KIF11 |  |  | RAB10 |  | SLC25A3 |  |  | JADE2 |
| JADE2 | KIF3B |  |  | RAB13 |  | SLC3A2 |  |  | KIF11 |
| KIF11 | KPNB1 |  |  | RAB14 |  | SMURF1 |  |  | KIF3B |
| KIF3B | KRAS |  |  | RAB15 |  | SNX12 |  |  | KPNB1 |
| KPNB1 | KRT18 |  |  | RAB21 |  | SNX3 |  |  | KRAS |
| KRAS | KRT6B |  |  | RAB22A |  | SOD1 |  |  | KRT18 |
| KRT18 | KRT79 |  |  | RAB2A |  | SPTBN1 |  |  | KRT6B |
| KRT6B | LAMA5 |  |  | RAB33B |  | SRI |  |  | KRT79 |
| KRT79 | LAMB1 |  |  | RAB35 |  | ST13 |  |  | LAMA5 |
| LAMA5 | LAMC1 |  |  | RAB3D |  | STOML1 |  |  | LAMB1 |
| LAMB1 | LAMP1 |  |  | RAB5B |  | STXBP1 |  |  | LAMC1 |
| LAMC1 | LAMP2 |  |  | RAB5C |  | STXBP2 |  |  | LAMP2 |
| LAMP1 | LAMTOR1 |  |  | RAB7A |  | TAGLN2 |  |  | LAMTOR1 |
| LAMP2 | LAP3 |  |  | RAB8A |  | TAX1BP3 |  |  | LAP3 |
| LAMTOR1 | LDHA |  |  | RAB8B |  | TBC1D21 |  |  | LDHA |
| LAP3 | LDHB |  |  | RAC1 |  | TCP1 |  |  | LDHB |
| LDHA | LGALS3 |  |  | RACK1 |  | THBS1 |  |  | LGALS3 |
| LDHB | LMAN2 |  |  | RALA |  | TLN1 |  |  | LMAN2 |
| LGALS3 | LOXL2 |  |  | RAN |  | TMEM59 |  |  | LOXL2 |
| LMAN2 | M6PR |  |  | RAP1B |  | TMX1 |  |  | M6PR |
| LOXL2 | MAMDC2 |  |  | RAP2B |  | TOLLIP |  |  | MAMDC2 |
| M6PR | MAN1A1 |  |  | RAPGEFL1 |  | TPI1 |  |  | MAN1A1 |
| MAMDC2 | MAN2B2 |  |  | RARS1 |  | TSG101 |  |  | MAN2B2 |
| MAN1A1 | MARCKS |  |  | RDX |  | TTN |  |  | MARCKS |
| MAN2B2 | MASP1 |  |  | RENBP |  | TUBA1A |  |  | MASP1 |
| MARCKS | MELTF |  |  | RHEB |  | TUBB2A |  |  | MBL |
| MASP1 | MFGE8 |  |  | RHOA |  | TUBB6 |  |  | MELTF |
| MBL | MINK1 |  |  | RPS11 |  | TUBB8 |  |  | MFGE8 |
| MELTF | MMP2 |  |  | RPS18 |  | UGP2 |  |  | MINK1 |
| MFGE8 | MOGS |  |  | RRAS |  | VAMP3 |  |  | MMP2 |
| MINK1 | MPZL1 |  |  | RRAS2 |  | VAMP7 |  |  | MOGS |
| MMP2 | MSN |  |  | S100A10 |  | VAMP8 |  |  | MPZL1 |
| MOGS | MTHFD1 |  |  | S100A16 |  | VAT1 |  |  | MSN |
| MPZL1 | MVP |  |  | S100A4 |  | VCL |  |  | MTHFD1 |
| MSN | MXRA8 |  |  | SDCBP |  | VCP |  |  | MVP |
| MTHFD1 | MYH14 |  |  | SEC13 |  | VDAC1 |  |  | MXRA8 |
| MVP | MYH9 |  |  | SFN |  | VPS28 |  |  | MYH14 |
| MXRA8 | MYO1B |  |  | SH3 |  | VPS35 |  |  | MYH9 |
| MYH14 | MYO1C |  |  | SLC22A11 |  | VPS4B |  |  | MYO1B |
| MYH9 | MYO1D |  |  | SLC3A2 |  | VTA1 |  |  | MYO1C |
| MYO1B | MYOF |  |  | SNX18 |  | WDR1 |  |  | MYO1D |
| MYO1C | N4BP2L2 |  |  | SPAG9 |  | YWHAE |  |  | MYOF |
| MYO1D | NAA50 |  |  | SRI |  | YWHAG |  |  | N4BP2L2 |
| MYOF | NAGK |  |  | ST13 |  | YWHAH |  |  | NAA50 |
| N4BP2L2 | NAPA |  |  | STK24 |  | YWHAQ |  |  | NAGK |
| NAA50 | NCKAP1 |  |  | STOML1 |  | YWHAZ |  |  | NAPA |
| NAGK | NIBAN2 |  |  | TAGLN2 |  |  |  |  | NARS1 |
| NAPA | NID1 |  |  | TAX1BP3 |  |  |  |  | NCKAP1 |
| NARS1 | NID2 |  |  | TCP1 |  |  |  |  | NIBAN2 |
| NCKAP1 | NONO |  |  | THBS1 |  |  |  |  | NID1 |
| NIBAN2 | NOTCH2 |  |  | TKT |  |  |  |  | NID2 |
| NID1 | NOTCH3 |  |  | TLN1 |  |  |  |  | NONO |
| NID2 | NPEPPS |  |  | TMEM59 |  |  |  |  | NOTCH2 |
| NONO | NPM1 |  |  | TMX1 |  |  |  |  | NOTCH3 |
| NOTCH2 | NRAS |  |  | TPI1 |  |  |  |  | NPEPPS |
| NOTCH3 | NRP2 |  |  | TPP1 |  |  |  |  | NPM1 |
| NPEPPS | NT5E |  |  | TSG101 |  |  |  |  | NRAS |
| NPM1 | NUTF2 |  |  | TTN |  |  |  |  | NRP2 |
| NRAS | OLA1 |  |  | TUBA1A |  |  |  |  | NT5E |
| NRP2 | ORM1 |  |  | TUBB2A |  |  |  |  | NUTF2 |
| NT5E | OTUB1 |  |  | TUBB6 |  |  |  |  | OLA1 |
| NUTF2 | OXSR1 |  |  | TUBB8 |  |  |  |  | ORM1 |
| OLA1 | P3H1 |  |  | UBA1 |  |  |  |  | OTUB1 |
| ORM1 | PACSIN2 |  |  | UGP2 |  |  |  |  | OXSR1 |
| OTUB1 | PAFAH1B1 |  |  | VAMP3 |  |  |  |  | P3H1 |
| OXSR1 | PAFAH1B2 |  |  | VASP |  |  |  |  | PACSIN2 |
| P3H1 | PARK7 |  |  | VAT1 |  |  |  |  | PAFAH1B1 |
| PACSIN2 | PCBP1 |  |  | VCL |  |  |  |  | PAFAH1B2 |
| PAFAH1B1 | PDCD10 |  |  | VCP |  |  |  |  | PARK7 |
| PAFAH1B2 | PDCD6 |  |  | VPS35 |  |  |  |  | PCBP1 |
| PARK7 | PDCD6IP |  |  | VPS36 |  |  |  |  | PDCD10 |
| PCBP1 | PEBP1 |  |  | VPS4B |  |  |  |  | PDCD6 |
| PDCD10 | PEF1 |  |  | VTA1 |  |  |  |  | PDCD6IP |
| PDCD6 | PFKL |  |  | WDR1 |  |  |  |  | PEBP1 |
| PDCD6IP | PFN1 |  |  | YWHAE |  |  |  |  | PEF1 |
| PEBP1 | PFN2 |  |  | YWHAG |  |  |  |  | PFKL |
| PEF1 | PGD |  |  | YWHAQ |  |  |  |  | PFN1 |
| PFKL | PGM1 |  |  | YWHAZ |  |  |  |  | PFN2 |
| PFN1 | PHGDH |  |  |  |  |  |  |  | PGD |
| PFN2 | PKLR |  |  |  |  |  |  |  | PGM1 |
| PGD | PKM |  |  |  |  |  |  |  | PHGDH |
| PGM1 | PLAU |  |  |  |  |  |  |  | PKLR |
| PHGDH | PLBD2 |  |  |  |  |  |  |  | PKM |
| PKLR | PLD3 |  |  |  |  |  |  |  | PLAU |
| PKM | PLOD1 |  |  |  |  |  |  |  | PLBD2 |
| PLAU | PLOD3 |  |  |  |  |  |  |  | PLD3 |
| PLBD2 | PLS1 |  |  |  |  |  |  |  | PLOD1 |
| PLD3 | PLXNB2 |  |  |  |  |  |  |  | PLOD3 |
| PLOD1 | PNP |  |  |  |  |  |  |  | PLS1 |
| PLOD3 | POSTN |  |  |  |  |  |  |  | PLS2 |
| PLS1 | PPFIA2 |  |  |  |  |  |  |  | PLXNB2 |
| PLS2 | PPIB |  |  |  |  |  |  |  | PNP |
| PLXNB2 | PPIL1 |  |  |  |  |  |  |  | POSTN |
| PNP | PPL |  |  |  |  |  |  |  | PPFIA2 |
| POSTN | PPP1R7 |  |  |  |  |  |  |  | PPIB |
| PPFIA2 | PRDX5 |  |  |  |  |  |  |  | PPIL1 |
| PPIB | PRDX6 |  |  |  |  |  |  |  | PPL |
| PPIL1 | PRKAR2A |  |  |  |  |  |  |  | PPP1R7 |
| PPL | PRXL2B |  |  |  |  |  |  |  | PRDX5 |
| PPP1R7 | PSAT1 |  |  |  |  |  |  |  | PRDX6 |
| PRDX5 | PSMA1 |  |  |  |  |  |  |  | PRKAR2A |
| PRDX6 | PSMA2 |  |  |  |  |  |  |  | PRXL2B |
| PRKAR2A | PSMA3 |  |  |  |  |  |  |  | PSAT1 |
| PRXL2B | PSMA4 |  |  |  |  |  |  |  | PSMA1 |
| PSAT1 | PSMA5 |  |  |  |  |  |  |  | PSMA2 |
| PSMA1 | PSMB1 |  |  |  |  |  |  |  | PSMA3 |
| PSMA2 | PSMB2 |  |  |  |  |  |  |  | PSMA4 |
| PSMA3 | PSMB3 |  |  |  |  |  |  |  | PSMA5 |
| PSMA4 | PSMB4 |  |  |  |  |  |  |  | PSMB1 |
| PSMA5 | PSMB5 |  |  |  |  |  |  |  | PSMB2 |
| PSMB1 | PSMB6 |  |  |  |  |  |  |  | PSMB3 |
| PSMB2 | PSMC6 |  |  |  |  |  |  |  | PSMB4 |
| PSMB3 | PSMD12 |  |  |  |  |  |  |  | PSMB5 |
| PSMB4 | PSMD8 |  |  |  |  |  |  |  | PSMB6 |
| PSMB5 | PTGFRN |  |  |  |  |  |  |  | PSMC6 |
| PSMB6 | PTX3 |  |  |  |  |  |  |  | PSMD12 |
| PSMC6 | PWWP3B |  |  |  |  |  |  |  | PSMD8 |
| PSMD12 | PXDN |  |  |  |  |  |  |  | PTGFRN |
| PSMD8 | RAB10 |  |  |  |  |  |  |  | PTX3 |
| PTGFRN | RAB13 |  |  |  |  |  |  |  | PWWP3B |
| PTX3 | RAB14 |  |  |  |  |  |  |  | PXDN |
| PWWP3B | RAB15 |  |  |  |  |  |  |  | RAB10 |
| PXDN | RAB21 |  |  |  |  |  |  |  | RAB13 |
| RAB10 | RAB22A |  |  |  |  |  |  |  | RAB14 |
| RAB13 | RAB2A |  |  |  |  |  |  |  | RAB15 |
| RAB14 | RAB33B |  |  |  |  |  |  |  | RAB21 |
| RAB15 | RAB35 |  |  |  |  |  |  |  | RAB22A |
| RAB21 | RAB3D |  |  |  |  |  |  |  | RAB2A |
| RAB22A | RAB5B |  |  |  |  |  |  |  | RAB33B |
| RAB2A | RAB5C |  |  |  |  |  |  |  | RAB35 |
| RAB33B | RAB7A |  |  |  |  |  |  |  | RAB3D |
| RAB35 | RAB8A |  |  |  |  |  |  |  | RAB5B |
| RAB3D | RAB8B |  |  |  |  |  |  |  | RAB5C |
| RAB5B | RAC1 |  |  |  |  |  |  |  | RAB7A |
| RAB5C | RACK1 |  |  |  |  |  |  |  | RAB8A |
| RAB7A | RALA |  |  |  |  |  |  |  | RAB8B |
| RAB8A | RAN |  |  |  |  |  |  |  | RAC1 |
| RAB8B | RAP1B |  |  |  |  |  |  |  | RACK1 |
| RAC1 | RAP2B |  |  |  |  |  |  |  | RALA |
| RACK1 | RAPGEFL1 |  |  |  |  |  |  |  | RAN |
| RALA | RDX |  |  |  |  |  |  |  | RAP1B |
| RAN | RENBP |  |  |  |  |  |  |  | RAP2B |
| RAP1B | RHEB |  |  |  |  |  |  |  | RAPGEFL1 |
| RAP2B | RHOA |  |  |  |  |  |  |  | RARS1 |
| RAPGEFL1 | RHOG |  |  |  |  |  |  |  | RDX |
| RARS1 | ROR1 |  |  |  |  |  |  |  | RENBP |
| RDX | RPL18A |  |  |  |  |  |  |  | RHEB |
| RENBP | RPL28 |  |  |  |  |  |  |  | RHOA |
| RHEB | RPS11 |  |  |  |  |  |  |  | RHOG |
| RHOA | RPS18 |  |  |  |  |  |  |  | ROR1 |
| RHOG | RRAS |  |  |  |  |  |  |  | RPL18A |
| ROR1 | RRAS2 |  |  |  |  |  |  |  | RPL28 |
| RPL18A | S100A10 |  |  |  |  |  |  |  | RPS11 |
| RPL28 | S100A14 |  |  |  |  |  |  |  | RPS18 |
| RPS11 | S100A16 |  |  |  |  |  |  |  | RRAS |
| RPS18 | S100A4 |  |  |  |  |  |  |  | RRAS2 |
| RRAS | SCAMP3 |  |  |  |  |  |  |  | S100A10 |
| RRAS2 | SDCBP |  |  |  |  |  |  |  | S100A14 |
| S100A10 | SEC13 |  |  |  |  |  |  |  | S100A16 |
| S100A14 | SERPINA3 |  |  |  |  |  |  |  | S100A4 |
| S100A16 | SERPIND1 |  |  |  |  |  |  |  | SCAMP3 |
| S100A4 | SERPINF1 |  |  |  |  |  |  |  | SDCBP |
| SCAMP3 | SERPINF2 |  |  |  |  |  |  |  | SEC13 |
| SDCBP | SFN |  |  |  |  |  |  |  | SERPINA3 |
| SEC13 | SH3BGRL3 |  |  |  |  |  |  |  | SERPIND1 |
| SERPINA3 | SLC16A1 |  |  |  |  |  |  |  | SERPINF1 |
| SERPIND1 | SLC1A4 |  |  |  |  |  |  |  | SERPINF2 |
| SERPINF1 | SLC22A11 |  |  |  |  |  |  |  | SFN |
| SERPINF2 | SLC25A3 |  |  |  |  |  |  |  | SH3 |
| SFN | SLC29A1 |  |  |  |  |  |  |  | SH3BGRL3 |
| SH3BGRL3 | SLC3A2 |  |  |  |  |  |  |  | SLC16A1 |
| SLC16A1 | SLC44A2 |  |  |  |  |  |  |  | SLC1A4 |
| SLC1A4 | SMURF1 |  |  |  |  |  |  |  | SLC22A11 |
| SLC22A11 | SNX12 |  |  |  |  |  |  |  | SLC25A3 |
| SLC25A3 | SNX18 |  |  |  |  |  |  |  | SLC29A1 |
| SLC29A1 | SNX3 |  |  |  |  |  |  |  | SLC3A2 |
| SLC3A2 | SOD1 |  |  |  |  |  |  |  | SLC44A2 |
| SLC44A2 | SPAG9 |  |  |  |  |  |  |  | SMURF1 |
| SMURF1 | SPP2 |  |  |  |  |  |  |  | SNX12 |
| SNX12 | SPTBN1 |  |  |  |  |  |  |  | SNX18 |
| SNX18 | SRI |  |  |  |  |  |  |  | SNX3 |
| SNX3 | ST13 |  |  |  |  |  |  |  | SOD1 |
| SOD1 | STK24 |  |  |  |  |  |  |  | SPAG9 |
| SPAG9 | STXBP1 |  |  |  |  |  |  |  | SPP2 |
| SPP2 | STXBP2 |  |  |  |  |  |  |  | SPTBN1 |
| SPTBN1 | TAGLN2 |  |  |  |  |  |  |  | SRI |
| SRI | TAX1BP3 |  |  |  |  |  |  |  | ST13 |
| ST13 | TBC1D21 |  |  |  |  |  |  |  | STK24 |
| STK24 | TCP1 |  |  |  |  |  |  |  | STOML1 |
| STOML1 | TF |  |  |  |  |  |  |  | STXBP1 |
| STXBP1 | TG |  |  |  |  |  |  |  | STXBP2 |
| STXBP2 | TGFBI |  |  |  |  |  |  |  | TAGLN2 |
| TAGLN2 | THBS1 |  |  |  |  |  |  |  | TAX1BP3 |
| TAX1BP3 | TKT |  |  |  |  |  |  |  | TBC1D21 |
| TBC1D21 | TLN1 |  |  |  |  |  |  |  | TCP1 |
| TCP1 | TMEM59 |  |  |  |  |  |  |  | TF |
| TF | TMEM63A |  |  |  |  |  |  |  | TG |
| TG | TMX1 |  |  |  |  |  |  |  | TGFBI |
| TGFBI | TOLLIP |  |  |  |  |  |  |  | THBS1 |
| THBS1 | TPI1 |  |  |  |  |  |  |  | TKT |
| TKT | TPP1 |  |  |  |  |  |  |  | TLN1 |
| TLN1 | TSG101 |  |  |  |  |  |  |  | TMEM59 |
| TMEM59 | TTN |  |  |  |  |  |  |  | TMX1 |
| TMEM63A | TTYH3 |  |  |  |  |  |  |  | TOLLIP |
| TMX1 | TUBA1A |  |  |  |  |  |  |  | TPI1 |
| TOLLIP | TUBB2A |  |  |  |  |  |  |  | TPP1 |
| TPI1 | TUBB6 |  |  |  |  |  |  |  | TSG101 |
| TPP1 | TUBB8 |  |  |  |  |  |  |  | TTN |
| TSG101 | UBA1 |  |  |  |  |  |  |  | TTYH3 |
| TTN | UGP2 |  |  |  |  |  |  |  | TUBA1A |
| TTYH3 | VAMP3 |  |  |  |  |  |  |  | TUBB2A |
| TUBA1A | VAMP7 |  |  |  |  |  |  |  | TUBB6 |
| TUBB2A | VAMP8 |  |  |  |  |  |  |  | TUBB8 |
| TUBB6 | VASN |  |  |  |  |  |  |  | UBA1 |
| TUBB8 | VASP |  |  |  |  |  |  |  | UGP2 |
| UBA1 | VAT1 |  |  |  |  |  |  |  | VAMP3 |
| UGP2 | VCL |  |  |  |  |  |  |  | VAMP7 |
| VAMP3 | VCP |  |  |  |  |  |  |  | VAMP8 |
| VAMP7 | VDAC1 |  |  |  |  |  |  |  | VASN |
| VAMP8 | VPS28 |  |  |  |  |  |  |  | VASP |
| VASN | VPS35 |  |  |  |  |  |  |  | VAT1 |
| VASP | VPS36 |  |  |  |  |  |  |  | VCL |
| VAT1 | VPS37B |  |  |  |  |  |  |  | VCP |
| VCL | VPS4A |  |  |  |  |  |  |  | VDAC1 |
| VCP | VPS4B |  |  |  |  |  |  |  | VPS28 |
| VDAC1 | VTA1 |  |  |  |  |  |  |  | VPS35 |
| VPS28 | WDR1 |  |  |  |  |  |  |  | VPS36 |
| VPS35 | YWHAE |  |  |  |  |  |  |  | VPS37B |
| VPS36 | YWHAG |  |  |  |  |  |  |  | VPS4A |
| VPS37B | YWHAH |  |  |  |  |  |  |  | VPS4B |
| VPS4A | YWHAQ |  |  |  |  |  |  |  | VTA1 |
| VPS4B | YWHAZ |  |  |  |  |  |  |  | WDR1 |
| VTA1 |  |  |  |  |  |  |  |  | YWHAE |
| WDR1 |  |  |  |  |  |  |  |  | YWHAG |
| YWHAE |  |  |  |  |  |  |  |  | YWHAH |
| YWHAG |  |  |  |  |  |  |  |  | YWHAQ |
| YWHAH |  |  |  |  |  |  |  |  | YWHAZ |
| YWHAQ |  |  |  |  |  |  |  |  |  |
| YWHAZ |  |  |  |  |  |  |  |  |  |
