## Supplementary Table 3: ToppFun functional enrichment analysis for pathways associated with sEV proteins versus ExoCarta and/or Vesiclepedia datasets s for "Small Extracellular Vesicles by Nucleus Pulposus Cells Maintain Niche and Cell Homeostasis via Receptor Shuffling and Metabolic Enzyme Supplements"

**Supplementary Table 3:** ToppFun functional enrichment analysis for pathways associated with sEV proteins versus ExoCarta and/or Vesiclepedia datasets showing a selection of the top 200 pathways.

| **#** | **ID** | **Name** | **p-value** | **Bonferroni** | **# Genes from input** |
| --- | --- | --- | --- | --- | --- |
| 1 | MM14661 | REACTOME INNATE IMMUNE SYSTEM | 6.09E-43 | 1.89E-39 | [126](https://toppgene.cchmc.org/showQueryTerms.jsp?userdata_id=f84adcdb-6f56-4f32-8800-82bb00b8d437&feature=pt&row=0) |
| 3 | M27620 | REACTOME NEUTROPHIL DEGRANULATION | 4.26E-40 | 1.32E-36 | [85](https://toppgene.cchmc.org/showQueryTerms.jsp?userdata_id=f84adcdb-6f56-4f32-8800-82bb00b8d437&feature=pt&row=2) |
| 5 | MM14662 | REACTOME IMMUNE SYSTEM | 9.62E-36 | 2.99E-32 | [154](https://toppgene.cchmc.org/showQueryTerms.jsp?userdata_id=f84adcdb-6f56-4f32-8800-82bb00b8d437&feature=pt&row=4) |
| 6 | M8395 | REACTOME HEMOSTASIS | 4.67E-27 | 1.45E-23 | [83](https://toppgene.cchmc.org/showQueryTerms.jsp?userdata_id=f84adcdb-6f56-4f32-8800-82bb00b8d437&feature=pt&row=5) |
| 7 | M29853 | REACTOME NERVOUS SYSTEM DEVELOPMENT | 6.31E-26 | 1.96E-22 | [75](https://toppgene.cchmc.org/showQueryTerms.jsp?userdata_id=f84adcdb-6f56-4f32-8800-82bb00b8d437&feature=pt&row=6) |
| 8 | M27507 | REACTOME VESICLE MEDIATED TRANSPORT | 1.76E-22 | 5.47E-19 | [79](https://toppgene.cchmc.org/showQueryTerms.jsp?userdata_id=f84adcdb-6f56-4f32-8800-82bb00b8d437&feature=pt&row=7) |
| 9 | M1077 | REACTOME PLATELET ACTIVATION SIGNALING AND AGGREGATION | 1.81E-22 | 5.61E-19 | [47](https://toppgene.cchmc.org/showQueryTerms.jsp?userdata_id=f84adcdb-6f56-4f32-8800-82bb00b8d437&feature=pt&row=8) |
| 10 | M18 | PID INTEGRIN1 PATHWAY | 3.31E-22 | 1.03E-18 | [26](https://toppgene.cchmc.org/showQueryTerms.jsp?userdata_id=f84adcdb-6f56-4f32-8800-82bb00b8d437&feature=pt&row=9) |
| 11 | M11480 | REACTOME MEMBRANE TRAFFICKING | 1.07E-21 | 3.33E-18 | [72](https://toppgene.cchmc.org/showQueryTerms.jsp?userdata_id=f84adcdb-6f56-4f32-8800-82bb00b8d437&feature=pt&row=10) |
| 12 | M39729 | WP VEGFA VEGFR2 SIGNALING | 1.09E-21 | 3.37E-18 | [59](https://toppgene.cchmc.org/showQueryTerms.jsp?userdata_id=f84adcdb-6f56-4f32-8800-82bb00b8d437&feature=pt&row=11) |
| 16 | M610 | REACTOME EXTRACELLULAR MATRIX ORGANIZATION | 1.61E-17 | 5.00E-14 | [44](https://toppgene.cchmc.org/showQueryTerms.jsp?userdata_id=f84adcdb-6f56-4f32-8800-82bb00b8d437&feature=pt&row=15) |
| 17 | MM15676 | REACTOME NERVOUS SYSTEM DEVELOPMENT | 2.33E-17 | 7.24E-14 | [41](https://toppgene.cchmc.org/showQueryTerms.jsp?userdata_id=f84adcdb-6f56-4f32-8800-82bb00b8d437&feature=pt&row=16) |
| 22 | M7253 | KEGG FOCAL ADHESION | 8.21E-16 | 2.55E-12 | [34](https://toppgene.cchmc.org/showQueryTerms.jsp?userdata_id=f84adcdb-6f56-4f32-8800-82bb00b8d437&feature=pt&row=21) |
| 25 | M27201 | REACTOME EPH EPHRIN SIGNALING | 8.57E-15 | 2.66E-11 | [23](https://toppgene.cchmc.org/showQueryTerms.jsp?userdata_id=f84adcdb-6f56-4f32-8800-82bb00b8d437&feature=pt&row=24) |
| 28 | M16894 | KEGG COMPLEMENT AND COAGULATION CASCADES | 2.24E-14 | 6.95E-11 | [20](https://toppgene.cchmc.org/showQueryTerms.jsp?userdata_id=f84adcdb-6f56-4f32-8800-82bb00b8d437&feature=pt&row=27) |
| 29 | M194 | BIOCARTA PROTEASOME PATHWAY | 4.07E-14 | 1.27E-10 | [12](https://toppgene.cchmc.org/showQueryTerms.jsp?userdata_id=f84adcdb-6f56-4f32-8800-82bb00b8d437&feature=pt&row=28) |
| 32 | M39581 | WP COMPLEMENT SYSTEM | 1.45E-13 | 4.49E-10 | [22](https://toppgene.cchmc.org/showQueryTerms.jsp?userdata_id=f84adcdb-6f56-4f32-8800-82bb00b8d437&feature=pt&row=31) |
| 34 | M164 | PID ERBB1 DOWNSTREAM PATHWAY | 1.83E-13 | 5.67E-10 | [23](https://toppgene.cchmc.org/showQueryTerms.jsp?userdata_id=f84adcdb-6f56-4f32-8800-82bb00b8d437&feature=pt&row=33) |
| 35 | MM14537 | REACTOME DEVELOPMENTAL BIOLOGY | 2.39E-13 | 7.43E-10 | [48](https://toppgene.cchmc.org/showQueryTerms.jsp?userdata_id=f84adcdb-6f56-4f32-8800-82bb00b8d437&feature=pt&row=34) |
| 39 | M26954 | REACTOME TRANSLOCATION OF SLC2A4 GLUT4 TO THE PLASMA MEMBRANE | 6.61E-13 | 2.05E-09 | [19](https://toppgene.cchmc.org/showQueryTerms.jsp?userdata_id=f84adcdb-6f56-4f32-8800-82bb00b8d437&feature=pt&row=38) |
| 51 | M27436 | REACTOME PROGRAMMED CELL DEATH | 5.98E-12 | 1.86E-08 | [30](https://toppgene.cchmc.org/showQueryTerms.jsp?userdata_id=f84adcdb-6f56-4f32-8800-82bb00b8d437&feature=pt&row=50) |
| 73 | M27080 | REACTOME RHO GTPASE EFFECTORS | 6.24E-11 | 1.94E-07 | [36](https://toppgene.cchmc.org/showQueryTerms.jsp?userdata_id=f84adcdb-6f56-4f32-8800-82bb00b8d437&feature=pt&row=72) |
| 108 | M11521 | KEGG GLYCOLYSIS GLUCONEOGENESIS | 1.98E-08 | 2.16E-06 | [15](https://toppgene.cchmc.org/showQueryTerms.jsp?userdata_id=f84adcdb-6f56-4f32-8800-82bb00b8d437&feature=pt&row=107) |
| 133 | M27219 | REACTOME ECM PROTEOGLYCANS | 1.98E-08 | 5.29E-06 | [16](https://toppgene.cchmc.org/showQueryTerms.jsp?userdata_id=f84adcdb-6f56-4f32-8800-82bb00b8d437&feature=pt&row=132) |
| 143 | M62 | PID EPHB FWD PATHWAY | 2.45E-09 | 7.61E-06 | [12](https://toppgene.cchmc.org/showQueryTerms.jsp?userdata_id=f84adcdb-6f56-4f32-8800-82bb00b8d437&feature=pt&row=142) |
| 157 | M5539 | KEGG AXON GUIDANCE | 2.45E-09 | 1.45E-05 | [20](https://toppgene.cchmc.org/showQueryTerms.jsp?userdata_id=f84adcdb-6f56-4f32-8800-82bb00b8d437&feature=pt&row=156) |
| 191 | M124 | PID CXCR4 PATHWAY | 1.64E-08 | 5.08E-05 | [17](https://toppgene.cchmc.org/showQueryTerms.jsp?userdata_id=f84adcdb-6f56-4f32-8800-82bb00b8d437&feature=pt&row=190) |
