## Supplementary Table 4: ToppFun analysis of shared pathways between sEV proteins from autologous bovine NP, AF, and FAT parent cells. AF: Annulus fibro for "Small Extracellular Vesicles by Nucleus Pulposus Cells Maintain Niche and Cell Homeostasis via Receptor Shuffling and Metabolic Enzyme Supplements"

**Supplementary Table 4:** ToppFun analysis of shared pathways between sEV proteins from autologous bovine NP, AF, and FAT parent cells. AF: Annulus fibrosus; FAT: Subcutaneous adipose tissue; GO: gene ontology; bta: Bos taurus; cAMP: Cyclic adenosine monophosphate; ECM: Extracellular matrix; KEGG: Kyoto Encyclopedia of Genes and Genomes; FAT: Subcutaneous adipose tissue; MAPK: Mitogen-activated protein kinase; NP: Nucleus pulposus. *denotes 20 pathways detected for all investigated bovine parent cells in this study; ^#^denotes 9 pathways detected for all investigated sEV proteins in this study.

| **39 common pathways for sEV proteins shared by AF, NP FAT parent cells** | **75 common pathways for sEV proteins shared only by NP and FAT parent cells** | **12 common pathways for sEV proteins shared only by AF and NP parent cells** | **23 pathways for sEV proteins only from NP parent cells** |
| --- | --- | --- | --- |
| REACTOME TRANSLOCATION OF SLC2A4 GLUT4 TO THE PLASMA MEMBRANE | REACTOME CELLULAR RESPONSES TO STIMULI | REACTOME SARS COV 1 INFECTION | KEGG GLYCOLYSIS GLUCONEOGENESIS |
| *^#^REACTOME NEUTROPHIL DEGRANULATION | REACTOME AUF1 HNRNP D0 BINDS AND DESTABILIZES MRNA | WP PATHOGENIC ESCHERICHIA COLI INFECTION | WP CLEAR CELL RENAL CELL CARCINOMA PATHWAYS |
| REACTOME HEMOSTASIS | KEGG MEDICUS VARIANT MUTATION INACTIVATED VCP TO 26S PROTEASOME MEDIATED PROTEIN DEGRADATION | KEGG PATHOGENIC ESCHERICHIA COLI INFECTION | WP METABOLIC REPROGRAMMING IN COLON CANCER |
| *^#^WP VEGFA VEGFR2 SIGNALING | BIOCARTA PROTEASOME PATHWAY | KEGG ENDOCYTOSIS | KEGG MEDICUS REFERENCE GLYCOLYSIS |
| *^#^REACTOME PLATELET ACTIVATION SIGNALING AND AGGREGATION | REACTOME PCP CE PATHWAY | REACTOME HSP90 CHAPERONE CYCLE FOR STEROID HORMONE RECEPTORS SHR IN THE PRESENCE OF LIGAND | REACTOME MITOTIC METAPHASE AND ANAPHASE |
| *VKEGG REGULATION OF ACTIN CYTOSKELETON | REACTOME HIV INFECTION | WP 17P13 3 YWHAE COPY NUMBER VARIATION | REACTOME SIGNALING BY ROBO RECEPTORS |
| *REACTOME MEMBRANE TRAFFICKING | KEGG MEDICUS VARIANT SCRAPIE CONFORMATION PRPSC TO 26S PROTEASOME MEDIATED PROTEIN DEGRADATION | WP AEROBIC GLYCOLYSIS | KEGG MEDICUS PATHOGEN SALMONELLA SOPE TO RAC SIGNALING PATHWAY |
| *REACTOME VESICLE MEDIATED TRANSPORT | KEGG MEDICUS VARIANT MUTATION CAUSED ABERRANT ABETA TO 26S PROTEASOME MEDIATED PROTEIN DEGRADATION | WP PARKIN UBIQUITIN PROTEASOMAL SYSTEM PATHWAY | REACTOME ASSEMBLY OF THE PRE REPLICATIVE COMPLEX |
| *REACTOME INFECTIOUS DISEASE | KEGG MEDICUS VARIANT MUTATION CAUSED ABERRANT HTT TO 26S PROTEASOME MEDIATED PROTEIN DEGRADATION | REACTOME INTEGRATION OF ENERGY METABOLISM | REACTOME FOLDING OF ACTIN BY CCT TRIC |
| REACTOME L1CAM INTERACTIONS | KEGG MEDICUS VARIANT MUTATION CAUSED ABERRANT SNCA TO 26S PROTEASOME MEDIATED PROTEIN DEGRADATION | REACTOME MHC CLASS II ANTIGEN PRESENTATION | REACTOME ASSOCIATION OF TRIC CCT WITH TARGET PROTEINS DURING BIOSYNTHESIS |
| *^#^REACTOME IMMUNE SYSTEM | REACTOME THE ROLE OF GTSE1 IN G2 M PROGRESSION AFTER G2 CHECKPOINT | REACTOME EPH EPHRIN SIGNALING | WP GLYCOLYSIS AND GLUCONEOGENESIS |
| *^#^REACTOME INNATE IMMUNE SYSTEM | REACTOME MITOTIC G2 G2 M PHASES | BIOCARTA MCALPAIN PATHWAY | KEGG MEDICUS REFERENCE ARNO ARF ACTB G SIGNALING PATHWAY |
| PID A6B1 A6B4 INTEGRIN PATHWAY | KEGG MEDICUS REFERENCE 26S PROTEASOME MEDIATED PROTEIN DEGRADATION |  | SIG REGULATION OF THE ACTIN CYTOSKELETON BY RHO GTPASES |
| REACTOME RHO GTPASES ACTIVATE PKNS | KEGG MEDICUS VARIANT MUTATION INACTIVATED UBQLN2 TO 26S PROTEASOME MEDIATED PROTEIN DEGRADATION |  | KEGG PENTOSE PHOSPHATE PATHWAY |
| *^#^KEGG FOCAL ADHESION | REACTOME MAPK6 MAPK4 SIGNALING |  | WP METABOLIC EPILEPTIC DISORDERS |
| *REACTOME SIGNALING BY RHO GTPASES MIRO GTPASES AND RHOBTB3 | KEGG MEDICUS VARIANT MUTATION CAUSED ABERRANT SOD1 TO 26S PROTEASOME MEDIATED PROTEIN DEGRADATION |  | PID LIS1 PATHWAY |
| *^#^REACTOME NERVOUS SYSTEM DEVELOPMENT | REACTOME REGULATION OF MRNA STABILITY BY PROTEINS THAT BIND AU RICH ELEMENTS |  | KEGG MEDICUS PATHOGEN SHIGELLA IPGD TO ARNO ARF ACTB G SIGNALING PATHWAY |
| *REACTOME RHO GTPASE EFFECTORS | REACTOME HOST INTERACTIONS OF HIV FACTORS |  | KEGG MEDICUS PATHOGEN ESCHERICHIA MAP TO CDC42 SIGNALING PATHWAY |
| PID CXCR4 PATHWAY | REACTOME NEGATIVE REGULATION OF NOTCH4 SIGNALING |  | REACTOME DNA REPLICATION PRE INITIATION |
| *WP EBOLA VIRUS INFECTION IN HOST | REACTOME BETA CATENIN INDEPENDENT WNT SIGNALING |  | REACTOME HEDGEHOG ON STATE |
| *PID PDGFRB PATHWAY | KEGG PROTEASOME |  | REACTOME COPI INDEPENDENT GOLGI TO ER RETROGRADE TRAFFIC |
| *PID ERBB1 DOWNSTREAM PATHWAY | REACTOME NUCLEAR EVENTS MEDIATED BY NFE2L2 |  | KEGG MEDICUS PATHOGEN SALMONELLA SOPB TO ARNO ARF ACTB G SIGNALING PATHWAY |
| REACTOME RAB GERANYLGERANYLATION | REACTOME CROSS PRESENTATION OF SOLUBLE EXOGENOUS ANTIGENS ENDOSOMES |  | REACTOME PTEN REGULATION |
| REACTOME ACTIVATION OF BAD AND TRANSLOCATION TO MITOCHONDRIA | REACTOME APOPTOSIS |  |  |
| REACTOME SARS COV 2 TARGETS HOST INTRACELLULAR SIGNALLING AND REGULATORY PATHWAYS | REACTOME G2 M CHECKPOINTS |  |  |
| REACTOME CHK1 CHK2 CDS1 MEDIATED INACTIVATION OF CYCLIN B CDK1 COMPLEX | REACTOME M PHASE |  |  |
| WP REGULATION OF ACTIN CYTOSKELETON | REACTOME DEFECTIVE CFTR CAUSES CYSTIC FIBROSIS |  |  |
| PID ER NONGENOMIC PATHWAY | REACTOME REGULATION OF RUNX2 EXPRESSION AND ACTIVITY |  |  |
| *^#^WP FOCAL ADHESION | REACTOME UBIQUITIN MEDIATED DEGRADATION OF PHOSPHORYLATED CDC25A |  |  |
| REACTOME SARS COV 1 TARGETS HOST INTRACELLULAR SIGNALLING AND REGULATORY PATHWAYS | WP PROTEASOME DEGRADATION |  |  |
| PID S1P S1P3 PATHWAY | REACTOME HEDGEHOG LIGAND BIOGENESIS |  |  |
| KEGG MEDICUS REFERENCE ITGA B TALIN VINCULIN SIGNALING PATHWAY | REACTOME ANTIGEN PROCESSING CROSS PRESENTATION |  |  |
| *REACTOME SARS COV 1 HOST INTERACTIONS | REACTOME REGULATION OF RUNX3 EXPRESSION AND ACTIVITY |  |  |
| REACTOME RESPONSE TO ELEVATED PLATELET CYTOSOLIC CA2 | REACTOME DEGRADATION OF AXIN |  |  |
| PID INTEGRIN1 PATHWAY | REACTOME STABILIZATION OF P53 |  |  |
| *REACTOME VIRAL INFECTION PATHWAYS | REACTOME REGULATION OF RAS BY GAPS |  |  |
| REACTOME LAMININ INTERACTIONS | REACTOME CELLULAR RESPONSE TO CHEMICAL STRESS |  |  |
| WP INTEGRIN MEDIATED CELL ADHESION | REACTOME KEAP1 NFE2L2 PATHWAY |  |  |
| *REACTOME PROGRAMMED CELL DEATH | REACTOME SOMITOGENESIS |  |  |
|  | REACTOME DEGRADATION OF DVL |  |  |
|  | REACTOME SCF BETA TRCP MEDIATED DEGRADATION OF EMI1 |  |  |
|  | REACTOME DECTIN 1 MEDIATED NONCANONICAL NF KB SIGNALING |  |  |
|  | REACTOME FORMATION OF TUBULIN FOLDING INTERMEDIATES BY CCT TRIC |  |  |
|  | REACTOME GLI3 IS PROCESSED TO GLI3R BY THE PROTEASOME |  |  |
|  | REACTOME METABOLISM OF POLYAMINES |  |  |
|  | REACTOME HEDGEHOG OFF STATE |  |  |
|  | REACTOME SCF SKP2 MEDIATED DEGRADATION OF P27 P21 |  |  |
|  | REACTOME DEGRADATION OF GLI1 BY THE PROTEASOME |  |  |
|  | REACTOME ASYMMETRIC LOCALIZATION OF PCP PROTEINS |  |  |
|  | REACTOME POST TRANSLATIONAL PROTEIN MODIFICATION |  |  |
|  | REACTOME TRANSCRIPTIONAL REGULATION BY RUNX2 |  |  |
|  | REACTOME ACTIVATION OF NF KAPPAB IN B CELLS |  |  |
|  | REACTOME G1 S DNA DAMAGE CHECKPOINTS |  |  |
|  | REACTOME ABC TRANSPORTER DISORDERS |  |  |
|  | REACTOME DOWNSTREAM SIGNALING EVENTS OF B CELL RECEPTOR BCR |  |  |
|  | REACTOME RUNX1 REGULATES TRANSCRIPTION OF GENES INVOLVED IN DIFFERENTIATION OF HSCS |  |  |
|  | REACTOME METABOLISM OF PROTEINS |  |  |
|  | WP ALZHEIMER 39 S DISEASE |  |  |
|  | WP ALZHEIMER 39 S DISEASE AND MIRNA EFFECTS |  |  |
|  | REACTOME SIGNALING BY NOTCH4 |  |  |
|  | REACTOME COOPERATION OF PREFOLDIN AND TRIC CCT IN ACTIN AND TUBULIN FOLDING |  |  |
|  | REACTOME ORC1 REMOVAL FROM CHROMATIN |  |  |
|  | REACTOME REGULATION OF PTEN STABILITY AND ACTIVITY |  |  |
|  | REACTOME CELLULAR RESPONSE TO HYPOXIA |  |  |
|  | REACTOME FORMATION OF PARAXIAL MESODERM |  |  |
|  | REACTOME SIGNALING BY HEDGEHOG |  |  |
|  | REACTOME CYCLIN A CDK2 ASSOCIATED EVENTS AT S PHASE ENTRY |  |  |
|  | REACTOME CELL CYCLE |  |  |
|  | REACTOME CDK MEDIATED PHOSPHORYLATION AND REMOVAL OF CDC6 |  |  |
|  | REACTOME APC C CDH1 MEDIATED DEGRADATION OF CDC20 AND OTHER APC C CDH1 TARGETED PROTEINS IN LATE MITOSIS EARLY G1 |  |  |
|  | REACTOME TRANSCRIPTIONAL REGULATION BY RUNX3 |  |  |
|  | REACTOME COOPERATION OF PDCL PHLP1 AND TRIC CCT IN G PROTEIN BETA FOLDING |  |  |
|  | REACTOME ACTIVATION OF APC C AND APC C CDC20 MEDIATED DEGRADATION OF MITOTIC PROTEINS |  |  |
|  | REACTOME CHAPERONIN MEDIATED PROTEIN FOLDING |  |  |
|  | REACTOME DEGRADATION OF BETA CATENIN BY THE DESTRUCTION COMPLEX |  |  |
