## Supplementary Table 5: ToppFun analysis of shared pathways between sEV proteins from bovine adult and fetal NP, and UCMSC parent cells. bta: Bos tauru for "Small Extracellular Vesicles by Nucleus Pulposus Cells Maintain Niche and Cell Homeostasis via Receptor Shuffling and Metabolic Enzyme Supplements"

**Supplementary Table 5:** ToppFun analysis of shared pathways between sEV proteins from bovine adult and fetal NP, and UCMSC parent cells. bta: Bos taurus; ECM: Extracellular matrix; KEGG: Kyoto Encyclopedia of Genes and Genomes; NP: Nucleus pulposus. *denotes 20 pathways detected for all investigated bovine parent cells in this study; ^#^denotes 9 pathways detected for all investigated sEV proteins in this study.

| **74 common pathways for sEV proteins shared by adult and fetal NP and UCMSC parent cells** | **59 common pathways for sEV proteins shared only by adult and fetal NP parent cells** | **7 common pathways for sEV proteins shared only by adult NP and UCMSC parent cells** | **1 common pathway for sEV proteins shared only by fetal NP and UCMSC parent cells** |
| --- | --- | --- | --- |
| *^#^REACTOME NEUTROPHIL DEGRANULATION | REACTOME TRANSLOCATION OF SLC2A4 GLUT4 TO THE PLASMA MEMBRANE | REACTOME SIGNALING BY ROBO RECEPTORS | REACTOME DISEASES OF SIGNAL TRANSDUCTION BY GROWTH FACTOR RECEPTORS AND SECOND MESSENGERS |
| *^#^REACTOME INNATE IMMUNE SYSTEM | REACTOME HEMOSTASIS | REACTOME CDK MEDIATED PHOSPHORYLATION AND REMOVAL OF CDC6 |  |
| *^#^REACTOME IMMUNE SYSTEM | BIOCARTA PROTEASOME PATHWAY | REACTOME APC C CDH1 MEDIATED DEGRADATION OF CDC20 AND OTHER APC C CDH1 TARGETED PROTEINS IN LATE MITOSIS EARLY G1 |  |
| *REACTOME MEMBRANE TRAFFICKING | KEGG GLYCOLYSIS GLUCONEOGENESIS | REACTOME ACTIVATION OF APC C AND APC C CDC20 MEDIATED DEGRADATION OF MITOTIC PROTEINS |  |
| *REACTOME VESICLE MEDIATED TRANSPORT | REACTOME RAB GERANYLGERANYLATION | REACTOME DEGRADATION OF BETA CATENIN BY THE DESTRUCTION COMPLEX |  |
| *REACTOME INFECTIOUS DISEASE | KEGG MEDICUS VARIANT SCRAPIE CONFORMATION PRPSC TO 26S PROTEASOME MEDIATED PROTEIN DEGRADATION | REACTOME DNA REPLICATION PRE INITIATION |  |
| *^#^REACTOME NERVOUS SYSTEM DEVELOPMENT | KEGG MEDICUS VARIANT MUTATION CAUSED ABERRANT ABETA TO 26S PROTEASOME MEDIATED PROTEIN DEGRADATION | REACTOME PTEN REGULATION |  |
| *^#^WP VEGFA VEGFR2 SIGNALING | KEGG MEDICUS VARIANT MUTATION CAUSED ABERRANT HTT TO 26S PROTEASOME MEDIATED PROTEIN DEGRADATION |  |  |
| REACTOME CELLULAR RESPONSES TO STIMULI | KEGG MEDICUS VARIANT MUTATION CAUSED ABERRANT SNCA TO 26S PROTEASOME MEDIATED PROTEIN DEGRADATION |  |  |
| *REACTOME PROGRAMMED CELL DEATH | REACTOME HSP90 CHAPERONE CYCLE FOR STEROID HORMONE RECEPTORS SHR IN THE PRESENCE OF LIGAND |  |  |
| *^#^KEGG REGULATION OF ACTIN CYTOSKELETON | WP CLEAR CELL RENAL CELL CARCINOMA PATHWAYS |  |  |
| *REACTOME SIGNALING BY RHO GTPASES MIRO GTPASES AND RHOBTB3 | KEGG MEDICUS VARIANT MUTATION CAUSED ABERRANT SOD1 TO 26S PROTEASOME MEDIATED PROTEIN DEGRADATION |  |  |
| REACTOME AUF1 HNRNP D0 BINDS AND DESTABILIZES MRNA | REACTOME L1CAM INTERACTIONS |  |  |
| *REACTOME VIRAL INFECTION PATHWAYS | KEGG MEDICUS REFERENCE GLYCOLYSIS |  |  |
| *PID ERBB1 DOWNSTREAM PATHWAY | REACTOME SOMITOGENESIS |  |  |
| KEGG MEDICUS VARIANT MUTATION INACTIVATED VCP TO 26S PROTEASOME MEDIATED PROTEIN DEGRADATION | WP PATHOGENIC ESCHERICHIA COLI INFECTION |  |  |
| *REACTOME RHO GTPASE EFFECTORS | KEGG PATHOGENIC ESCHERICHIA COLI INFECTION |  |  |
| REACTOME PCP CE PATHWAY | REACTOME FORMATION OF TUBULIN FOLDING INTERMEDIATES BY CCT TRIC |  |  |
| *^#^REACTOME PLATELET ACTIVATION SIGNALING AND AGGREGATION | REACTOME METABOLISM OF POLYAMINES |  |  |
| *REACTOME SARS COV 1 INFECTION | PID A6B1 A6B4 INTEGRIN PATHWAY |  |  |
| REACTOME HIV INFECTION | REACTOME RESPONSE TO ELEVATED PLATELET CYTOSOLIC CA2 |  |  |
| REACTOME THE ROLE OF GTSE1 IN G2 M PROGRESSION AFTER G2 CHECKPOINT | KEGG MEDICUS PATHOGEN SALMONELLA SOPE TO RAC SIGNALING PATHWAY |  |  |
| REACTOME MITOTIC G2 G2 M PHASES | WP 17P13 3 YWHAE COPY NUMBER VARIATION |  |  |
| KEGG MEDICUS REFERENCE 26S PROTEASOME MEDIATED PROTEIN DEGRADATION | REACTOME G1 S DNA DAMAGE CHECKPOINTS |  |  |
| KEGG MEDICUS VARIANT MUTATION INACTIVATED UBQLN2 TO 26S PROTEASOME MEDIATED PROTEIN DEGRADATION | REACTOME ABC TRANSPORTER DISORDERS |  |  |
| REACTOME MAPK6 MAPK4 SIGNALING | REACTOME RUNX1 REGULATES TRANSCRIPTION OF GENES INVOLVED IN DIFFERENTIATION OF HSCS |  |  |
| REACTOME REGULATION OF MRNA STABILITY BY PROTEINS THAT BIND AU RICH ELEMENTS | WP ALZHEIMER 39 S DISEASE |  |  |
| *PID PDGFRB PATHWAY | REACTOME RHO GTPASES ACTIVATE PKNS |  |  |
| REACTOME HOST INTERACTIONS OF HIV FACTORS | REACTOME FOLDING OF ACTIN BY CCT TRIC |  |  |
| REACTOME NEGATIVE REGULATION OF NOTCH4 SIGNALING | REACTOME ASSOCIATION OF TRIC CCT WITH TARGET PROTEINS DURING BIOSYNTHESIS |  |  |
| REACTOME BETA CATENIN INDEPENDENT WNT SIGNALING | WP ALZHEIMER 39 S DISEASE AND MIRNA EFFECTS |  |  |
| KEGG PROTEASOME | REACTOME SARS COV 1 TARGETS HOST INTRACELLULAR SIGNALLING AND REGULATORY PATHWAYS |  |  |
| REACTOME NUCLEAR EVENTS MEDIATED BY NFE2L2 | REACTOME SIGNALING BY NOTCH4 |  |  |
| REACTOME CROSS PRESENTATION OF SOLUBLE EXOGENOUS ANTIGENS ENDOSOMES | REACTOME COOPERATION OF PREFOLDIN AND TRIC CCT IN ACTIN AND TUBULIN FOLDING |  |  |
| REACTOME APOPTOSIS | REACTOME CELLULAR RESPONSE TO HYPOXIA |  |  |
| REACTOME G2 M CHECKPOINTS | REACTOME FORMATION OF PARAXIAL MESODERM |  |  |
| *^#^KEGG FOCAL ADHESION | REACTOME SIGNALING BY HEDGEHOG |  |  |
| REACTOME M PHASE | PID CXCR4 PATHWAY |  |  |
| REACTOME DEFECTIVE CFTR CAUSES CYSTIC FIBROSIS | KEGG MEDICUS REFERENCE ARNO ARF ACTB G SIGNALING PATHWAY |  |  |
| REACTOME REGULATION OF RUNX2 EXPRESSION AND ACTIVITY | WP INTEGRIN MEDIATED CELL ADHESION |  |  |
| REACTOME UBIQUITIN MEDIATED DEGRADATION OF PHOSPHORYLATED CDC25A | REACTOME TRANSCRIPTIONAL REGULATION BY RUNX3 |  |  |
| WP PROTEASOME DEGRADATION | REACTOME ACTIVATION OF BAD AND TRANSLOCATION TO MITOCHONDRIA |  |  |
| REACTOME HEDGEHOG LIGAND BIOGENESIS | REACTOME SARS COV 2 TARGETS HOST INTRACELLULAR SIGNALLING AND REGULATORY PATHWAYS |  |  |
| REACTOME ANTIGEN PROCESSING CROSS PRESENTATION | WP AEROBIC GLYCOLYSIS |  |  |
| REACTOME REGULATION OF RUNX3 EXPRESSION AND ACTIVITY | REACTOME CHK1 CHK2 CDS1 MEDIATED INACTIVATION OF CYCLIN B CDK1 COMPLEX |  |  |
| WP METABOLIC REPROGRAMMING IN COLON CANCER | REACTOME INTEGRATION OF ENERGY METABOLISM |  |  |
| REACTOME DEGRADATION OF AXIN | REACTOME COOPERATION OF PDCL PHLP1 AND TRIC CCT IN G PROTEIN BETA FOLDING |  |  |
| REACTOME STABILIZATION OF P53 | REACTOME EPH EPHRIN SIGNALING |  |  |
| REACTOME REGULATION OF RAS BY GAPS | REACTOME CHAPERONIN MEDIATED PROTEIN FOLDING |  |  |
| REACTOME CELLULAR RESPONSE TO CHEMICAL STRESS | PID S1P S1P3 PATHWAY |  |  |
| REACTOME KEAP1 NFE2L2 PATHWAY | KEGG MEDICUS PATHOGEN SHIGELLA IPGD TO ARNO ARF ACTB G SIGNALING PATHWAY |  |  |
| REACTOME DEGRADATION OF DVL | PID ER NONGENOMIC PATHWAY |  |  |
| REACTOME SCF BETA TRCP MEDIATED DEGRADATION OF EMI1 | KEGG MEDICUS PATHOGEN ESCHERICHIA MAP TO CDC42 SIGNALING PATHWAY |  |  |
| REACTOME MITOTIC METAPHASE AND ANAPHASE | PID INTEGRIN1 PATHWAY |  |  |
| REACTOME DECTIN 1 MEDIATED NONCANONICAL NF KB SIGNALING | REACTOME HEDGEHOG ON STATE |  |  |
| KEGG ENDOCYTOSIS | WP REGULATION OF ACTIN CYTOSKELETON |  |  |
| REACTOME GLI3 IS PROCESSED TO GLI3R BY THE PROTEASOME | KEGG MEDICUS REFERENCE ITGA B TALIN VINCULIN SIGNALING PATHWAY |  |  |
| REACTOME HEDGEHOG OFF STATE | KEGG MEDICUS PATHOGEN SALMONELLA SOPB TO ARNO ARF ACTB G SIGNALING PATHWAY |  |  |
| REACTOME SCF SKP2 MEDIATED DEGRADATION OF P27 P21 | REACTOME LAMININ INTERACTIONS |  |  |
| REACTOME DEGRADATION OF GLI1 BY THE PROTEASOME |  |  |  |
| REACTOME ASYMMETRIC LOCALIZATION OF PCP PROTEINS |  |  |  |
| REACTOME POST TRANSLATIONAL PROTEIN MODIFICATION |  |  |  |
| REACTOME TRANSCRIPTIONAL REGULATION BY RUNX2 |  |  |  |
| REACTOME ACTIVATION OF NF KAPPAB IN B CELLS |  |  |  |
| REACTOME DOWNSTREAM SIGNALING EVENTS OF B CELL RECEPTOR BCR |  |  |  |
| REACTOME METABOLISM OF PROTEINS |  |  |  |
| *^#^WP FOCAL ADHESION |  |  |  |
| REACTOME ASSEMBLY OF THE PRE REPLICATIVE COMPLEX |  |  |  |
| REACTOME SARS COV 1 HOST INTERACTIONS |  |  |  |
| REACTOME ORC1 REMOVAL FROM CHROMATIN |  |  |  |
| REACTOME REGULATION OF PTEN STABILITY AND ACTIVITY |  |  |  |
| REACTOME CYCLIN A CDK2 ASSOCIATED EVENTS AT S PHASE ENTRY |  |  |  |
| REACTOME CELL CYCLE |  |  |  |
| *WP EBOLA VIRUS INFECTION IN HOST |  |  |  |
