## Supplementary Table 6: ToppFun pathway analysis of NP over- or underrepresented sEV proteins. for "Small Extracellular Vesicles by Nucleus Pulposus Cells Maintain Niche and Cell Homeostasis via Receptor Shuffling and Metabolic Enzyme Supplements"

| **Type** | **#** | **ID** | **Name** | **p-value** | **Bonferroni** | **# Genes from input** |
| --- | --- | --- | --- | --- | --- | --- |
| over-represented | 1 | M16894 | KEGG COMPLEMENT AND COAGULATION CASCADES | 1.52E-26 | 2.48E-23 | [21](https://toppgene.cchmc.org/showQueryTerms.jsp?userdata_id=114a21ba-041d-4c33-9b52-1d71268ed0d5&feature=pt&row=0) |
|  | 2 | M18 | PID INTEGRIN1 PATHWAY | 1.45E-23 | 2.36E-20 | [19](https://toppgene.cchmc.org/showQueryTerms.jsp?userdata_id=114a21ba-041d-4c33-9b52-1d71268ed0d5&feature=pt&row=1) |
|  | 3 | M39649 | WP COMPLEMENT AND COAGULATION CASCADES | 1.18E-19 | 1.92E-16 | [16](https://toppgene.cchmc.org/showQueryTerms.jsp?userdata_id=114a21ba-041d-4c33-9b52-1d71268ed0d5&feature=pt&row=2) |
|  | 4 | M27285 | REACTOME REGULATION OF INSULIN LIKE GROWTH FACTOR IGF TRANSPORT AND UPTAKE BY INSULIN LIKE GROWTH FACTOR BINDING PROTEINS IGFBPS | 2.67E-19 | 4.34E-16 | [20](https://toppgene.cchmc.org/showQueryTerms.jsp?userdata_id=114a21ba-041d-4c33-9b52-1d71268ed0d5&feature=pt&row=3) |
|  | 5 | M610 | REACTOME EXTRACELLULAR MATRIX ORGANIZATION | 5.49E-19 | 8.93E-16 | [27](https://toppgene.cchmc.org/showQueryTerms.jsp?userdata_id=114a21ba-041d-4c33-9b52-1d71268ed0d5&feature=pt&row=4) |
|  | 6 | M39581 | WP COMPLEMENT SYSTEM | 7.21E-19 | 1.17E-15 | [18](https://toppgene.cchmc.org/showQueryTerms.jsp?userdata_id=114a21ba-041d-4c33-9b52-1d71268ed0d5&feature=pt&row=5) |
|  | 7 | M917 | BIOCARTA COMP PATHWAY | 6.56E-18 | 1.07E-14 | [11](https://toppgene.cchmc.org/showQueryTerms.jsp?userdata_id=114a21ba-041d-4c33-9b52-1d71268ed0d5&feature=pt&row=6) |
|  | 8 | M39502 | WP COMPLEMENT ACTIVATION | 2.71E-17 | 4.41E-14 | [11](https://toppgene.cchmc.org/showQueryTerms.jsp?userdata_id=114a21ba-041d-4c33-9b52-1d71268ed0d5&feature=pt&row=7) |
|  | 9 | MM15944 | WP COMPLEMENT ACTIVATION CLASSICAL PATHWAY | 8.69E-17 | 1.41E-13 | [10](https://toppgene.cchmc.org/showQueryTerms.jsp?userdata_id=114a21ba-041d-4c33-9b52-1d71268ed0d5&feature=pt&row=8) |
|  | 10 | M8395 | REACTOME HEMOSTASIS | 9.62E-17 | 1.57E-13 | [35](https://toppgene.cchmc.org/showQueryTerms.jsp?userdata_id=114a21ba-041d-4c33-9b52-1d71268ed0d5&feature=pt&row=9) |
|  | 11 | MM14572 | REACTOME EXTRACELLULAR MATRIX ORGANIZATION | 3.03E-16 | 4.93E-13 | [23](https://toppgene.cchmc.org/showQueryTerms.jsp?userdata_id=114a21ba-041d-4c33-9b52-1d71268ed0d5&feature=pt&row=10) |
|  | 12 | M4732 | BIOCARTA LECTIN PATHWAY | 3.66E-16 | 5.95E-13 | [9](https://toppgene.cchmc.org/showQueryTerms.jsp?userdata_id=114a21ba-041d-4c33-9b52-1d71268ed0d5&feature=pt&row=11) |
|  | 13 | M7146 | BIOCARTA CLASSIC PATHWAY | 2.52E-15 | 4.10E-12 | [9](https://toppgene.cchmc.org/showQueryTerms.jsp?userdata_id=114a21ba-041d-4c33-9b52-1d71268ed0d5&feature=pt&row=12) |
|  | 14 | MM14661 | REACTOME INNATE IMMUNE SYSTEM | 1.96E-14 | 3.18E-11 | [39](https://toppgene.cchmc.org/showQueryTerms.jsp?userdata_id=114a21ba-041d-4c33-9b52-1d71268ed0d5&feature=pt&row=13) |
|  | 15 | M29853 | REACTOME NERVOUS SYSTEM DEVELOPMENT | 2.17E-14 | 3.52E-11 | [30](https://toppgene.cchmc.org/showQueryTerms.jsp?userdata_id=114a21ba-041d-4c33-9b52-1d71268ed0d5&feature=pt&row=14) |
| under-represented | 1 | M47613 | KEGG MEDICUS REFERENCE N GLYCAN PRECURSOR BIOSYNTHESIS ALG6 TO OST | 2.57E-05 | 3.88E-03 | [2](https://toppgene.cchmc.org/showQueryTerms.jsp?userdata_id=6ebe1e92-1766-4dfb-b172-50859905198d&feature=pt&row=0) |
|  | 2 | M39570 | WP AMINO ACID METABOLISM | 3.19E-05 | 4.81E-03 | [3](https://toppgene.cchmc.org/showQueryTerms.jsp?userdata_id=6ebe1e92-1766-4dfb-b172-50859905198d&feature=pt&row=1) |
|  | 3 | MM15942 | WP AMINO ACID METABOLISM | 3.98E-05 | 6.01E-03 | [3](https://toppgene.cchmc.org/showQueryTerms.jsp?userdata_id=6ebe1e92-1766-4dfb-b172-50859905198d&feature=pt&row=2) |
|  | 4 | M27851 | REACTOME GLUTAMATE AND GLUTAMINE METABOLISM | 4.25E-05 | 6.42E-03 | [2](https://toppgene.cchmc.org/showQueryTerms.jsp?userdata_id=6ebe1e92-1766-4dfb-b172-50859905198d&feature=pt&row=3) |
|  | 5 | MM15570 | REACTOME GLUTAMATE AND GLUTAMINE METABOLISM | 4.25E-05 | 6.42E-03 | [2](https://toppgene.cchmc.org/showQueryTerms.jsp?userdata_id=6ebe1e92-1766-4dfb-b172-50859905198d&feature=pt&row=4) |
|  | 6 | M39850 | WP UREA CYCLE AND ASSOCIATED PATHWAYS | 9.78E-05 | 1.48E-02 | [2](https://toppgene.cchmc.org/showQueryTerms.jsp?userdata_id=6ebe1e92-1766-4dfb-b172-50859905198d&feature=pt&row=5) |
|  | 7 | M10911 | KEGG CYSTEINE AND METHIONINE METABOLISM | 2.60E-04 | 3.93E-02 | [2](https://toppgene.cchmc.org/showQueryTerms.jsp?userdata_id=6ebe1e92-1766-4dfb-b172-50859905198d&feature=pt&row=6) |
|  | 8 | M41728 | REACTOME MATURATION OF SARS COV 2 SPIKE PROTEIN | 3.08E-04 | 4.66E-02 | [2](https://toppgene.cchmc.org/showQueryTerms.jsp?userdata_id=6ebe1e92-1766-4dfb-b172-50859905198d&feature=pt&row=7) |
|  | 9 | M39474 | WP GLYCOLYSIS AND GLUCONEOGENESIS | 4.57E-04 | 6.90E-02 | [2](https://toppgene.cchmc.org/showQueryTerms.jsp?userdata_id=6ebe1e92-1766-4dfb-b172-50859905198d&feature=pt&row=8) |
|  | 10 | M11079 | KEGG N GLYCAN BIOSYNTHESIS | 4.78E-04 | 7.21E-02 | [2](https://toppgene.cchmc.org/showQueryTerms.jsp?userdata_id=6ebe1e92-1766-4dfb-b172-50859905198d&feature=pt&row=9) |
|  | 11 | MM15928 | WP GLYCOLYSIS AND GLUCONEOGENESIS | 5.87E-04 | 8.87E-02 | [2](https://toppgene.cchmc.org/showQueryTerms.jsp?userdata_id=6ebe1e92-1766-4dfb-b172-50859905198d&feature=pt&row=10) |
