## Supplementary Table 7: Comparison of top 200 results of ToppFun pathways for each sEV source identifies nine pathways shared by all sources for "Small Extracellular Vesicles by Nucleus Pulposus Cells Maintain Niche and Cell Homeostasis via Receptor Shuffling and Metabolic Enzyme Supplements"

| **Group** | **Shared ToppFun Pathways** |
| --- | --- |
| A: sEV source: NP fetal, NP adult, NP overrepresented and ExoCarta | REACTOME NEUTROPHIL DEGRANULATION |
|  | REACTOME INNATE IMMUNE SYSTEM |
|  | REACTOME IMMUNE SYSTEM |
|  | REACTOME PLATELET ACTIVATION SIGNALING AND AGGREGATION |
|  | REACTOME HEMOSTASIS |
|  | REACTOME NERVOUS SYSTEM DEVELOPMENT |
|  | KEGG REGULATION OF ACTIN CYTOSKELETON |
|  | WP VEGFA VEGFR2 SIGNALING |
|  | REACTOME AUF1 HNRNP D0 BINDS AND DESTABILIZES MRNA |
|  | REACTOME BETA CATENIN INDEPENDENT WNT SIGNALING |
|  | REACTOME PCP CE PATHWAY |
|  | KEGG MEDICUS VARIANT MUTATION INACTIVATED VCP TO 26S PROTEASOME MEDIATED PROTEIN DEGRADATION |
|  | KEGG MEDICUS VARIANT MUTATION CAUSED ABERRANT SOD1 TO 26S PROTEASOME MEDIATED PROTEIN DEGRADATION |
|  | REACTOME HIV INFECTION |
|  | REACTOME MAPK6 MAPK4 SIGNALING |
|  | KEGG MEDICUS REFERENCE 26S PROTEASOME MEDIATED PROTEIN DEGRADATION |
|  | KEGG MEDICUS VARIANT MUTATION INACTIVATED UBQLN2 TO 26S PROTEASOME MEDIATED PROTEIN DEGRADATION |
|  | REACTOME ANTIGEN PROCESSING CROSS PRESENTATION |
|  | REACTOME REGULATION OF RUNX3 EXPRESSION AND ACTIVITY |
|  | KEGG FOCAL ADHESION |
|  | REACTOME NEGATIVE REGULATION OF NOTCH4 SIGNALING |
|  | REACTOME RESPONSE TO ELEVATED PLATELET CYTOSOLIC CA2 |
|  | REACTOME ASYMMETRIC LOCALIZATION OF PCP PROTEINS |
|  | REACTOME DEFECTIVE CFTR CAUSES CYSTIC FIBROSIS |
|  | REACTOME REGULATION OF RUNX2 EXPRESSION AND ACTIVITY |
|  | REACTOME UBIQUITIN MEDIATED DEGRADATION OF PHOSPHORYLATED CDC25A |
|  | REACTOME REGULATION OF RAS BY GAPS |
|  | REACTOME NUCLEAR EVENTS MEDIATED BY NFE2L2 |
|  | REACTOME DEGRADATION OF AXIN |
|  | REACTOME DEGRADATION OF DVL |
|  | REACTOME SCF BETA TRCP MEDIATED DEGRADATION OF EMI1 |
|  | REACTOME DECTIN 1 MEDIATED NONCANONICAL NF KB SIGNALING |
|  | REACTOME GLI3 IS PROCESSED TO GLI3R BY THE PROTEASOME |
|  | WP FOCAL ADHESION |
|  | REACTOME DEGRADATION OF GLI1 BY THE PROTEASOME |
|  | WP PROTEASOME DEGRADATION |
|  | REACTOME TRANSCRIPTIONAL REGULATION BY RUNX2 |
|  | REACTOME ACTIVATION OF NF KAPPAB IN B CELLS |
|  | REACTOME DOWNSTREAM SIGNALING EVENTS OF B CELL RECEPTOR BCR |
|  | REACTOME NEUTROPHIL DEGRANULATION |
| B: sEV source: NP fetal, NP adult, NP overrepresented | REACTOME NEUTROPHIL DEGRANULATION |
|  | REACTOME INNATE IMMUNE SYSTEM |
|  | REACTOME IMMUNE SYSTEM |
|  | REACTOME PLATELET ACTIVATION SIGNALING AND AGGREGATION |
|  | REACTOME HEMOSTASIS |
|  | REACTOME NERVOUS SYSTEM DEVELOPMENT |
|  | KEGG REGULATION OF ACTIN CYTOSKELETON |
|  | WP VEGFA VEGFR2 SIGNALING |
|  | REACTOME AUF1 HNRNP D0 BINDS AND DESTABILIZES MRNA |
|  | REACTOME BETA CATENIN INDEPENDENT WNT SIGNALING |
|  | REACTOME PCP CE PATHWAY |
|  | KEGG MEDICUS VARIANT MUTATION INACTIVATED VCP TO 26S PROTEASOME MEDIATED PROTEIN DEGRADATION |
|  | KEGG MEDICUS VARIANT MUTATION CAUSED ABERRANT SOD1 TO 26S PROTEASOME MEDIATED PROTEIN DEGRADATION |
|  | REACTOME HIV INFECTION |
|  | KEGG MEDICUS VARIANT SCRAPIE CONFORMATION PRPSC TO 26S PROTEASOME MEDIATED PROTEIN DEGRADATION |
|  | KEGG MEDICUS VARIANT MUTATION CAUSED ABERRANT ABETA TO 26S PROTEASOME MEDIATED PROTEIN DEGRADATION |
|  | KEGG MEDICUS VARIANT MUTATION CAUSED ABERRANT HTT TO 26S PROTEASOME MEDIATED PROTEIN DEGRADATION |
|  | KEGG MEDICUS VARIANT MUTATION CAUSED ABERRANT SNCA TO 26S PROTEASOME MEDIATED PROTEIN DEGRADATION |
|  | REACTOME MAPK6 MAPK4 SIGNALING |
|  | KEGG MEDICUS REFERENCE 26S PROTEASOME MEDIATED PROTEIN DEGRADATION |
|  | KEGG MEDICUS VARIANT MUTATION INACTIVATED UBQLN2 TO 26S PROTEASOME MEDIATED PROTEIN DEGRADATION |
|  | REACTOME ANTIGEN PROCESSING CROSS PRESENTATION |
|  | REACTOME REGULATION OF RUNX3 EXPRESSION AND ACTIVITY |
|  | KEGG FOCAL ADHESION |
|  | REACTOME NEGATIVE REGULATION OF NOTCH4 SIGNALING |
|  | REACTOME RESPONSE TO ELEVATED PLATELET CYTOSOLIC CA2 |
|  | KEGG PROTEASOME |
|  | BIOCARTA PROTEASOME PATHWAY |
|  | REACTOME ASYMMETRIC LOCALIZATION OF PCP PROTEINS |
|  | REACTOME CROSS PRESENTATION OF SOLUBLE EXOGENOUS ANTIGENS ENDOSOMES |
|  | REACTOME DEFECTIVE CFTR CAUSES CYSTIC FIBROSIS |
|  | REACTOME HEDGEHOG LIGAND BIOGENESIS |
|  | REACTOME REGULATION OF RUNX2 EXPRESSION AND ACTIVITY |
|  | REACTOME UBIQUITIN MEDIATED DEGRADATION OF PHOSPHORYLATED CDC25A |
|  | REACTOME REGULATION OF RAS BY GAPS |
|  | REACTOME NUCLEAR EVENTS MEDIATED BY NFE2L2 |
|  | REACTOME DEGRADATION OF AXIN |
|  | REACTOME STABILIZATION OF P53 |
|  | REACTOME SOMITOGENESIS |
|  | REACTOME DEGRADATION OF DVL |
|  | REACTOME SCF BETA TRCP MEDIATED DEGRADATION OF EMI1 |
|  | REACTOME DECTIN 1 MEDIATED NONCANONICAL NF KB SIGNALING |
|  | REACTOME GLI3 IS PROCESSED TO GLI3R BY THE PROTEASOME |
|  | REACTOME METABOLISM OF POLYAMINES |
|  | WP FOCAL ADHESION |
|  | REACTOME SCF SKP2 MEDIATED DEGRADATION OF P27 P21 |
|  | REACTOME DEGRADATION OF GLI1 BY THE PROTEASOME |
|  | REACTOME EPH EPHRIN SIGNALING |
|  | WP PROTEASOME DEGRADATION |
|  | REACTOME TRANSCRIPTIONAL REGULATION BY RUNX2 |
|  | REACTOME ACTIVATION OF NF KAPPAB IN B CELLS |
|  | REACTOME DOWNSTREAM SIGNALING EVENTS OF B CELL RECEPTOR BCR |
|  | REACTOME G1 S DNA DAMAGE CHECKPOINTS |
|  | PID INTEGRIN1 PATHWAY |
|  | REACTOME SIGNALING BY NOTCH4 |
|  | KEGG MEDICUS REFERENCE ITGA B TALIN VINCULIN SIGNALING PATHWAY |
|  | REACTOME LAMININ INTERACTIONS |
|  | WP INTEGRIN MEDIATED CELL ADHESION |
| C: sEV source: NP fetal, NP adult, NP overrepresented, UCMSC | REACTOME NEUTROPHIL DEGRANULATION |
|  | REACTOME INNATE IMMUNE SYSTEM |
|  | REACTOME IMMUNE SYSTEM |
|  | REACTOME PLATELET ACTIVATION SIGNALING AND AGGREGATION |
|  | REACTOME NERVOUS SYSTEM DEVELOPMENT |
|  | KEGG REGULATION OF ACTIN CYTOSKELETON |
|  | WP VEGFA VEGFR2 SIGNALING |
|  | REACTOME AUF1 HNRNP D0 BINDS AND DESTABILIZES MRNA |
|  | REACTOME BETA CATENIN INDEPENDENT WNT SIGNALING |
|  | REACTOME PCP CE PATHWAY |
|  | KEGG MEDICUS VARIANT MUTATION INACTIVATED VCP TO 26S PROTEASOME MEDIATED PROTEIN DEGRADATION |
|  | REACTOME HIV INFECTION |
|  | REACTOME MAPK6 MAPK4 SIGNALING |
|  | KEGG MEDICUS REFERENCE 26S PROTEASOME MEDIATED PROTEIN DEGRADATION |
|  | KEGG MEDICUS VARIANT MUTATION INACTIVATED UBQLN2 TO 26S PROTEASOME MEDIATED PROTEIN DEGRADATION |
|  | REACTOME ANTIGEN PROCESSING CROSS PRESENTATION |
|  | REACTOME REGULATION OF RUNX3 EXPRESSION AND ACTIVITY |
|  | KEGG FOCAL ADHESION |
|  | REACTOME NEGATIVE REGULATION OF NOTCH4 SIGNALING |
|  | KEGG PROTEASOME |
|  | REACTOME ASYMMETRIC LOCALIZATION OF PCP PROTEINS |
|  | REACTOME CROSS PRESENTATION OF SOLUBLE EXOGENOUS ANTIGENS ENDOSOMES |
|  | REACTOME DEFECTIVE CFTR CAUSES CYSTIC FIBROSIS |
|  | REACTOME HEDGEHOG LIGAND BIOGENESIS |
|  | REACTOME REGULATION OF RUNX2 EXPRESSION AND ACTIVITY |
|  | REACTOME UBIQUITIN MEDIATED DEGRADATION OF PHOSPHORYLATED CDC25A |
|  | REACTOME REGULATION OF RAS BY GAPS |
|  | REACTOME NUCLEAR EVENTS MEDIATED BY NFE2L2 |
|  | REACTOME DEGRADATION OF AXIN |
|  | REACTOME STABILIZATION OF P53 |
|  | REACTOME DEGRADATION OF DVL |
|  | REACTOME SCF BETA TRCP MEDIATED DEGRADATION OF EMI1 |
|  | REACTOME DECTIN 1 MEDIATED NONCANONICAL NF KB SIGNALING |
|  | REACTOME GLI3 IS PROCESSED TO GLI3R BY THE PROTEASOME |
|  | WP FOCAL ADHESION |
|  | REACTOME SCF SKP2 MEDIATED DEGRADATION OF P27 P21 |
|  | REACTOME DEGRADATION OF GLI1 BY THE PROTEASOME |
|  | WP PROTEASOME DEGRADATION |
|  | REACTOME TRANSCRIPTIONAL REGULATION BY RUNX2 |
|  | REACTOME ACTIVATION OF NF KAPPAB IN B CELLS |
|  | REACTOME DOWNSTREAM SIGNALING EVENTS OF B CELL RECEPTOR |
|  | REACTOME NEUTROPHIL DEGRANULATION |
| D: sEV source: AF adult, NP adult, FAT adult | REACTOME TRANSLOCATION OF SLC2A4 GLUT4 TO THE PLASMA MEMBRANE |
|  | REACTOME NEUTROPHIL DEGRANULATION |
|  | REACTOME HEMOSTASIS |
|  | WP VEGFA VEGFR2 SIGNALING |
|  | REACTOME PLATELET ACTIVATION SIGNALING AND AGGREGATION |
|  | KEGG REGULATION OF ACTIN CYTOSKELETON |
|  | REACTOME MEMBRANE TRAFFICKING |
|  | REACTOME VESICLE MEDIATED TRANSPORT |
|  | REACTOME INFECTIOUS DISEASE |
|  | REACTOME L1CAM INTERACTIONS |
|  | REACTOME IMMUNE SYSTEM |
|  | REACTOME INNATE IMMUNE SYSTEM |
|  | PID A6B1 A6B4 INTEGRIN PATHWAY |
|  | REACTOME RHO GTPASES ACTIVATE PKNS |
|  | KEGG FOCAL ADHESION |
|  | REACTOME SIGNALING BY RHO GTPASES MIRO GTPASES AND RHOBTB3 |
|  | REACTOME NERVOUS SYSTEM DEVELOPMENT |
|  | REACTOME RHO GTPASE EFFECTORS |
|  | PID CXCR4 PATHWAY |
|  | WP EBOLA VIRUS INFECTION IN HOST |
|  | PID PDGFRB PATHWAY |
|  | PID ERBB1 DOWNSTREAM PATHWAY |
|  | REACTOME RAB GERANYLGERANYLATION |
|  | REACTOME ACTIVATION OF BAD AND TRANSLOCATION TO MITOCHONDRIA |
|  | REACTOME SARS COV 2 TARGETS HOST INTRACELLULAR SIGNALLING AND REGULATORY PATHWAYS |
|  | REACTOME CHK1 CHK2 CDS1 MEDIATED INACTIVATION OF CYCLIN B CDK1 COMPLEX |
|  | WP REGULATION OF ACTIN CYTOSKELETON |
|  | PID ER NONGENOMIC PATHWAY |
|  | WP FOCAL ADHESION |
|  | REACTOME SARS COV 1 TARGETS HOST INTRACELLULAR SIGNALLING AND REGULATORY PATHWAYS |
|  | PID S1P S1P3 PATHWAY |
|  | KEGG MEDICUS REFERENCE ITGA B TALIN VINCULIN SIGNALING PATHWAY |
|  | REACTOME SARS COV 1 HOST INTERACTIONS |
|  | REACTOME RESPONSE TO ELEVATED PLATELET CYTOSOLIC CA2 |
|  | PID INTEGRIN1 PATHWAY |
|  | REACTOME VIRAL INFECTION PATHWAYS |
|  | REACTOME LAMININ INTERACTIONS |
|  | WP INTEGRIN MEDIATED CELL ADHESION |
|  | REACTOME PROGRAMMED CELL DEATH |
|  | REACTOME TRANSLOCATION OF SLC2A4 GLUT4 TO THE PLASMA MEMBRANE |
| Shared by all sEV sources | REACTOME NEUTROPHIL DEGRANULATION |
|  | REACTOME INNATE IMMUNE SYSTEM |
|  | REACTOME IMMUNE SYSTEM |
|  | REACTOME PLATELET ACTIVATION SIGNALING AND AGGREGATION |
|  | REACTOME NERVOUS SYSTEM DEVELOPMENT |
|  | KEGG REGULATION OF ACTIN CYTOSKELETON |
|  | WP VEGFA VEGFR2 SIGNALING |
|  | KEGG FOCAL ADHESION |
|  | WP FOCAL ADHESION |
| Exclusively associated with Group B | KEGG MEDICUS VARIANT SCRAPIE CONFORMATION PRPSC TO 26S PROTEASOME MEDIATED PROTEIN DEGRADATION |
|  | KEGG MEDICUS VARIANT MUTATION CAUSED ABERRANT ABETA TO 26S PROTEASOME MEDIATED PROTEIN DEGRADATION |
|  | KEGG MEDICUS VARIANT MUTATION CAUSED ABERRANT HTT TO 26S PROTEASOME MEDIATED PROTEIN DEGRADATION |
|  | KEGG MEDICUS VARIANT MUTATION CAUSED ABERRANT SNCA TO 26S PROTEASOME MEDIATED PROTEIN DEGRADATION |
|  | BIOCARTA PROTEASOME PATHWAY |
|  | REACTOME SOMITOGENESIS |
|  | REACTOME METABOLISM OF POLYAMINES |
|  | REACTOME EPH EPHRIN SIGNALING |
|  | REACTOME G1 S DNA DAMAGE CHECKPOINTS |
|  | REACTOME SIGNALING BY NOTCH4 |
