## Supplementary Table 8: ToppFun pathway analysis for fetal bovine serum (FBS) proteins and all NP sEV proteins after FBS protein removal. for "Small Extracellular Vesicles by Nucleus Pulposus Cells Maintain Niche and Cell Homeostasis via Receptor Shuffling and Metabolic Enzyme Supplements"

| FBS serum proteins | | | | | | | | All NP sEV proteins with FBS serum proteins removed | | | | | | | |
| --- | --- | --- | --- | --- | --- | --- | --- | --- | --- | --- | --- | --- | --- | --- | --- |
| FBS Protein ID | ToppFun - Top 200 pathway affiliations | | | | | | | All NP protein ID adjusted | ToppFun - Top 200 pathway affiliations of all NP proteins corrected for serum proteins from FBS | | | | | | |
|  | ID | Name | pValue | FDR B&H | FDR B&Y | Bonferroni | Genes from Input |  | ID | Name | pValue | FDR B&H | FDR B&Y | Bonferroni | Genes from Input |
| A1BG | M27620 | REACTOME NEUTROPHIL DEGRANULATION | 3.43E-35 | 1.23E-31 | 1.08E-30 | 1.23E-31 | [80](https://toppgene.cchmc.org/showQueryTerms.jsp?userdata_id=677e6385-c8f1-4db4-acb1-4b26e7d30bc3&feature=pt&row=0) | ACP1 | MM14661 | REACTOME INNATE IMMUNE SYSTEM | 3.64E-19 | 9.01E-16 | 7.56E-15 | 9.01E-16 | [67](https://toppgene.cchmc.org/showQueryTerms.jsp?userdata_id=e0bdc6b9-955d-48cd-95d3-4ccb1cbe49c7&feature=pt&row=0) |
| A2ML1 | MM15330 | REACTOME NEUTROPHIL DEGRANULATION | 1.37E-31 | 2.47E-28 | 2.17E-27 | 4.95E-28 | [79](https://toppgene.cchmc.org/showQueryTerms.jsp?userdata_id=677e6385-c8f1-4db4-acb1-4b26e7d30bc3&feature=pt&row=1) | ADH5 | M1036 | REACTOME INNATE IMMUNE SYSTEM | 5.82E-18 | 7.19E-15 | 6.04E-14 | 1.44E-14 | [69](https://toppgene.cchmc.org/showQueryTerms.jsp?userdata_id=e0bdc6b9-955d-48cd-95d3-4ccb1cbe49c7&feature=pt&row=1) |
| ABCC11 | M1036 | REACTOME INNATE IMMUNE SYSTEM | 6.38E-27 | 7.66E-24 | 6.72E-23 | 2.30E-23 | [110](https://toppgene.cchmc.org/showQueryTerms.jsp?userdata_id=677e6385-c8f1-4db4-acb1-4b26e7d30bc3&feature=pt&row=2) | AKR1A1 | M8395 | REACTOME HEMOSTASIS | 3.90E-16 | 3.22E-13 | 2.70E-12 | 9.65E-13 | [50](https://toppgene.cchmc.org/showQueryTerms.jsp?userdata_id=e0bdc6b9-955d-48cd-95d3-4ccb1cbe49c7&feature=pt&row=2) |
| ABHD14B | MM14661 | REACTOME INNATE IMMUNE SYSTEM | 1.64E-26 | 1.47E-23 | 1.29E-22 | 5.89E-23 | [103](https://toppgene.cchmc.org/showQueryTerms.jsp?userdata_id=677e6385-c8f1-4db4-acb1-4b26e7d30bc3&feature=pt&row=3) | ALDOC | M16894 | KEGG COMPLEMENT AND COAGULATION CASCADES | 8.60E-16 | 5.32E-13 | 4.46E-12 | 2.13E-12 | [18](https://toppgene.cchmc.org/showQueryTerms.jsp?userdata_id=e0bdc6b9-955d-48cd-95d3-4ccb1cbe49c7&feature=pt&row=3) |
| ACAA2 | MM14662 | REACTOME IMMUNE SYSTEM | 2.54E-20 | 1.83E-17 | 1.60E-16 | 9.15E-17 | [127](https://toppgene.cchmc.org/showQueryTerms.jsp?userdata_id=677e6385-c8f1-4db4-acb1-4b26e7d30bc3&feature=pt&row=4) | APPL2 | M29853 | REACTOME NERVOUS SYSTEM DEVELOPMENT | 1.08E-14 | 5.34E-12 | 4.48E-11 | 2.67E-11 | [44](https://toppgene.cchmc.org/showQueryTerms.jsp?userdata_id=e0bdc6b9-955d-48cd-95d3-4ccb1cbe49c7&feature=pt&row=4) |
| ACLY | M1077 | REACTOME PLATELET ACTIVATION SIGNALING AND AGGREGATION | 9.54E-11 | 5.72E-08 | 5.02E-07 | 3.43E-07 | [32](https://toppgene.cchmc.org/showQueryTerms.jsp?userdata_id=677e6385-c8f1-4db4-acb1-4b26e7d30bc3&feature=pt&row=5) | ARPC5L | MM14572 | REACTOME EXTRACELLULAR MATRIX ORGANIZATION | 2.52E-14 | 1.04E-11 | 8.72E-11 | 6.23E-11 | [29](https://toppgene.cchmc.org/showQueryTerms.jsp?userdata_id=e0bdc6b9-955d-48cd-95d3-4ccb1cbe49c7&feature=pt&row=5) |
| ACOT11 | M11480 | REACTOME MEMBRANE TRAFFICKING | 1.62E-10 | 8.34E-08 | 7.31E-07 | 5.83E-07 | [53](https://toppgene.cchmc.org/showQueryTerms.jsp?userdata_id=677e6385-c8f1-4db4-acb1-4b26e7d30bc3&feature=pt&row=6) | ATP6V0E2 | M610 | REACTOME EXTRACELLULAR MATRIX ORGANIZATION | 3.90E-14 | 1.26E-11 | 1.05E-10 | 9.65E-11 | [31](https://toppgene.cchmc.org/showQueryTerms.jsp?userdata_id=e0bdc6b9-955d-48cd-95d3-4ccb1cbe49c7&feature=pt&row=6) |
| ACOT8 | M724 | REACTOME RESPONSE TO ELEVATED PLATELET CYTOSOLIC CA2 | 2.51E-10 | 1.13E-07 | 9.92E-07 | 9.05E-07 | [22](https://toppgene.cchmc.org/showQueryTerms.jsp?userdata_id=677e6385-c8f1-4db4-acb1-4b26e7d30bc3&feature=pt&row=7) | ATP6V1B2 | M39729 | WP VEGFA VEGFR2 SIGNALING | 4.06E-14 | 1.26E-11 | 1.05E-10 | 1.00E-10 | [37](https://toppgene.cchmc.org/showQueryTerms.jsp?userdata_id=e0bdc6b9-955d-48cd-95d3-4ccb1cbe49c7&feature=pt&row=7) |
| ACP2 | M27507 | REACTOME VESICLE MEDIATED TRANSPORT | 4.24E-10 | 1.70E-07 | 1.49E-06 | 1.53E-06 | [57](https://toppgene.cchmc.org/showQueryTerms.jsp?userdata_id=677e6385-c8f1-4db4-acb1-4b26e7d30bc3&feature=pt&row=8) | CDC42BPB | MM14662 | REACTOME IMMUNE SYSTEM | 1.29E-13 | 3.54E-11 | 2.97E-10 | 3.19E-10 | [79](https://toppgene.cchmc.org/showQueryTerms.jsp?userdata_id=e0bdc6b9-955d-48cd-95d3-4ccb1cbe49c7&feature=pt&row=8) |
| ACSL4 | M18306 | KEGG REGULATION OF ACTIN CYTOSKELETON | 1.38E-09 | 4.97E-07 | 4.36E-06 | 4.97E-06 | [27](https://toppgene.cchmc.org/showQueryTerms.jsp?userdata_id=677e6385-c8f1-4db4-acb1-4b26e7d30bc3&feature=pt&row=9) | CDK5 | M27620 | REACTOME NEUTROPHIL DEGRANULATION | 2.03E-13 | 5.01E-11 | 4.20E-10 | 5.01E-10 | [38](https://toppgene.cchmc.org/showQueryTerms.jsp?userdata_id=e0bdc6b9-955d-48cd-95d3-4ccb1cbe49c7&feature=pt&row=9) |
| ACTBL2 | M11266 | KEGG LYSOSOME | 1.93E-09 | 6.31E-07 | 5.53E-06 | 6.94E-06 | [20](https://toppgene.cchmc.org/showQueryTerms.jsp?userdata_id=677e6385-c8f1-4db4-acb1-4b26e7d30bc3&feature=pt&row=10) | CHMP6 | M39649 | WP COMPLEMENT AND COAGULATION CASCADES | 2.64E-13 | 5.69E-11 | 4.78E-10 | 6.52E-10 | [15](https://toppgene.cchmc.org/showQueryTerms.jsp?userdata_id=e0bdc6b9-955d-48cd-95d3-4ccb1cbe49c7&feature=pt&row=10) |
| ACTN1 | M8395 | REACTOME HEMOSTASIS | 6.57E-09 | 1.97E-06 | 1.73E-05 | 2.37E-05 | [52](https://toppgene.cchmc.org/showQueryTerms.jsp?userdata_id=677e6385-c8f1-4db4-acb1-4b26e7d30bc3&feature=pt&row=11) | CHP1 | M1077 | REACTOME PLATELET ACTIVATION SIGNALING AND AGGREGATION | 2.84E-13 | 5.69E-11 | 4.78E-10 | 7.03E-10 | [28](https://toppgene.cchmc.org/showQueryTerms.jsp?userdata_id=e0bdc6b9-955d-48cd-95d3-4ccb1cbe49c7&feature=pt&row=11) |
| ADCY1 | MM15232 | REACTOME VESICLE MEDIATED TRANSPORT | 1.32E-08 | 3.67E-06 | 3.21E-05 | 4.77E-05 | [50](https://toppgene.cchmc.org/showQueryTerms.jsp?userdata_id=677e6385-c8f1-4db4-acb1-4b26e7d30bc3&feature=pt&row=12) | CRELD1 | MM14472 | REACTOME HEMOSTASIS | 2.99E-13 | 5.69E-11 | 4.78E-10 | 7.40E-10 | [41](https://toppgene.cchmc.org/showQueryTerms.jsp?userdata_id=e0bdc6b9-955d-48cd-95d3-4ccb1cbe49c7&feature=pt&row=12) |
| ADGRV1 | MM15454 | REACTOME PLATELET ACTIVATION SIGNALING AND AGGREGATION | 2.06E-08 | 5.31E-06 | 4.66E-05 | 7.43E-05 | [28](https://toppgene.cchmc.org/showQueryTerms.jsp?userdata_id=677e6385-c8f1-4db4-acb1-4b26e7d30bc3&feature=pt&row=13) | DCTN2 | MM15454 | REACTOME PLATELET ACTIVATION SIGNALING AND AGGREGATION | 1.28E-12 | 2.26E-10 | 1.90E-09 | 3.17E-09 | [27](https://toppgene.cchmc.org/showQueryTerms.jsp?userdata_id=e0bdc6b9-955d-48cd-95d3-4ccb1cbe49c7&feature=pt&row=13) |
| ADIRF | MM14786 | REACTOME MEMBRANE TRAFFICKING | 2.48E-08 | 5.96E-06 | 5.23E-05 | 8.94E-05 | [46](https://toppgene.cchmc.org/showQueryTerms.jsp?userdata_id=677e6385-c8f1-4db4-acb1-4b26e7d30bc3&feature=pt&row=14) | GALK1 | M18 | PID INTEGRIN1 PATHWAY | 2.07E-12 | 3.41E-10 | 2.86E-09 | 5.11E-09 | [15](https://toppgene.cchmc.org/showQueryTerms.jsp?userdata_id=e0bdc6b9-955d-48cd-95d3-4ccb1cbe49c7&feature=pt&row=14) |
| AHA1 | M232 | PID ECADHERIN STABILIZATION PATHWAY | 4.94E-08 | 1.05E-05 | 9.21E-05 | 1.78E-04 | [11](https://toppgene.cchmc.org/showQueryTerms.jsp?userdata_id=677e6385-c8f1-4db4-acb1-4b26e7d30bc3&feature=pt&row=15) | GANAB | M27507 | REACTOME VESICLE MEDIATED TRANSPORT | 6.00E-12 | 9.28E-10 | 7.79E-09 | 1.49E-08 | [45](https://toppgene.cchmc.org/showQueryTerms.jsp?userdata_id=e0bdc6b9-955d-48cd-95d3-4ccb1cbe49c7&feature=pt&row=15) |
| AHCTF1 | MM14472 | REACTOME HEMOSTASIS | 4.96E-08 | 1.05E-05 | 9.21E-05 | 1.79E-04 | [44](https://toppgene.cchmc.org/showQueryTerms.jsp?userdata_id=677e6385-c8f1-4db4-acb1-4b26e7d30bc3&feature=pt&row=16) | GET3 | MM15330 | REACTOME NEUTROPHIL DEGRANULATION | 1.34E-11 | 1.85E-09 | 1.55E-08 | 3.32E-08 | [37](https://toppgene.cchmc.org/showQueryTerms.jsp?userdata_id=e0bdc6b9-955d-48cd-95d3-4ccb1cbe49c7&feature=pt&row=16) |
| AHCY | M41838 | REACTOME SIGNALING BY RHO GTPASES MIRO GTPASES AND RHOBTB3 | 1.22E-07 | 2.43E-05 | 2.13E-04 | 4.38E-04 | [51](https://toppgene.cchmc.org/showQueryTerms.jsp?userdata_id=677e6385-c8f1-4db4-acb1-4b26e7d30bc3&feature=pt&row=17) | GLB1L3 | M11480 | REACTOME MEMBRANE TRAFFICKING | 1.35E-11 | 1.85E-09 | 1.55E-08 | 3.33E-08 | [41](https://toppgene.cchmc.org/showQueryTerms.jsp?userdata_id=e0bdc6b9-955d-48cd-95d3-4ccb1cbe49c7&feature=pt&row=17) |
| AHNAK | M7253 | KEGG FOCAL ADHESION | 1.32E-07 | 2.49E-05 | 2.19E-04 | 4.74E-04 | [23](https://toppgene.cchmc.org/showQueryTerms.jsp?userdata_id=677e6385-c8f1-4db4-acb1-4b26e7d30bc3&feature=pt&row=18) | GLG1 | M27201 | REACTOME EPH EPHRIN SIGNALING | 2.92E-11 | 3.80E-09 | 3.19E-08 | 7.21E-08 | [16](https://toppgene.cchmc.org/showQueryTerms.jsp?userdata_id=e0bdc6b9-955d-48cd-95d3-4ccb1cbe49c7&feature=pt&row=18) |
| AKR1B1 | M27623 | REACTOME SIGNALING BY MODERATE KINASE ACTIVITY BRAF MUTANTS | 1.41E-07 | 2.54E-05 | 2.23E-04 | 5.08E-04 | [11](https://toppgene.cchmc.org/showQueryTerms.jsp?userdata_id=677e6385-c8f1-4db4-acb1-4b26e7d30bc3&feature=pt&row=19) | GRHPR | M917 | BIOCARTA COMP PATHWAY | 5.65E-11 | 6.76E-09 | 5.68E-08 | 1.40E-07 | [9](https://toppgene.cchmc.org/showQueryTerms.jsp?userdata_id=e0bdc6b9-955d-48cd-95d3-4ccb1cbe49c7&feature=pt&row=19) |
| ALAD | M39729 | WP VEGFA VEGFR2 SIGNALING | 2.07E-07 | 3.40E-05 | 2.98E-04 | 7.46E-04 | [36](https://toppgene.cchmc.org/showQueryTerms.jsp?userdata_id=677e6385-c8f1-4db4-acb1-4b26e7d30bc3&feature=pt&row=20) | HSPA12A | M47725 | KEGG MEDICUS PATHOGEN ESCHERICHIA ESPT TO RAC SIGNALING PATHWAY | 5.74E-11 | 6.76E-09 | 5.68E-08 | 1.42E-07 | [8](https://toppgene.cchmc.org/showQueryTerms.jsp?userdata_id=e0bdc6b9-955d-48cd-95d3-4ccb1cbe49c7&feature=pt&row=20) |
| ALB | MM15455 | REACTOME RESPONSE TO ELEVATED PLATELET CYTOSOLIC CA2 | 2.08E-07 | 3.40E-05 | 2.98E-04 | 7.47E-04 | [18](https://toppgene.cchmc.org/showQueryTerms.jsp?userdata_id=677e6385-c8f1-4db4-acb1-4b26e7d30bc3&feature=pt&row=21) | LAP3 | M39502 | WP COMPLEMENT ACTIVATION | 1.62E-10 | 1.82E-08 | 1.53E-07 | 4.00E-07 | [9](https://toppgene.cchmc.org/showQueryTerms.jsp?userdata_id=e0bdc6b9-955d-48cd-95d3-4ccb1cbe49c7&feature=pt&row=21) |
| ALCAM | MM15690 | REACTOME SIGNALING BY RHO GTPASES MIRO GTPASES AND RHOBTB3 | 2.17E-07 | 3.40E-05 | 2.98E-04 | 7.83E-04 | [47](https://toppgene.cchmc.org/showQueryTerms.jsp?userdata_id=677e6385-c8f1-4db4-acb1-4b26e7d30bc3&feature=pt&row=22) | MAN2B2 | M164 | PID ERBB1 DOWNSTREAM PATHWAY | 2.30E-10 | 2.38E-08 | 1.99E-07 | 5.69E-07 | [16](https://toppgene.cchmc.org/showQueryTerms.jsp?userdata_id=e0bdc6b9-955d-48cd-95d3-4ccb1cbe49c7&feature=pt&row=22) |
| ALDH3B1 | MM15548 | REACTOME CELLULAR RESPONSES TO STIMULI | 2.83E-07 | 4.02E-05 | 3.53E-04 | 1.02E-03 | [38](https://toppgene.cchmc.org/showQueryTerms.jsp?userdata_id=677e6385-c8f1-4db4-acb1-4b26e7d30bc3&feature=pt&row=23) | N4BP2L2 | M631 | REACTOME COLLAGEN FORMATION | 2.31E-10 | 2.38E-08 | 1.99E-07 | 5.70E-07 | [15](https://toppgene.cchmc.org/showQueryTerms.jsp?userdata_id=e0bdc6b9-955d-48cd-95d3-4ccb1cbe49c7&feature=pt&row=23) |
| ALDOA | M142 | PID AJDISS 2PATHWAY | 2.88E-07 | 4.02E-05 | 3.53E-04 | 1.04E-03 | [11](https://toppgene.cchmc.org/showQueryTerms.jsp?userdata_id=677e6385-c8f1-4db4-acb1-4b26e7d30bc3&feature=pt&row=24) | NONO | MM14573 | REACTOME COLLAGEN FORMATION | 3.33E-10 | 3.30E-08 | 2.77E-07 | 8.25E-07 | [14](https://toppgene.cchmc.org/showQueryTerms.jsp?userdata_id=e0bdc6b9-955d-48cd-95d3-4ccb1cbe49c7&feature=pt&row=24) |
| ALK | M47723 | KEGG MEDICUS REFERENCE ITGA B TALIN VINCULIN SIGNALING PATHWAY | 2.90E-07 | 4.02E-05 | 3.53E-04 | 1.05E-03 | [9](https://toppgene.cchmc.org/showQueryTerms.jsp?userdata_id=677e6385-c8f1-4db4-acb1-4b26e7d30bc3&feature=pt&row=25) | NPEPPS | MM14911 | REACTOME EPH EPHRIN SIGNALING | 3.51E-10 | 3.34E-08 | 2.80E-07 | 8.68E-07 | [13](https://toppgene.cchmc.org/showQueryTerms.jsp?userdata_id=e0bdc6b9-955d-48cd-95d3-4ccb1cbe49c7&feature=pt&row=25) |
| ALPL | MM15272 | REACTOME MAP2K AND MAPK ACTIVATION | 3.24E-07 | 4.32E-05 | 3.79E-04 | 1.17E-03 | [10](https://toppgene.cchmc.org/showQueryTerms.jsp?userdata_id=677e6385-c8f1-4db4-acb1-4b26e7d30bc3&feature=pt&row=26) | OLA1 | M39581 | WP COMPLEMENT SYSTEM | 4.36E-10 | 3.90E-08 | 3.27E-07 | 1.08E-06 | [15](https://toppgene.cchmc.org/showQueryTerms.jsp?userdata_id=e0bdc6b9-955d-48cd-95d3-4ccb1cbe49c7&feature=pt&row=26) |
| AMBP | M872 | REACTOME L1CAM INTERACTIONS | 3.66E-07 | 4.70E-05 | 4.12E-04 | 1.32E-03 | [17](https://toppgene.cchmc.org/showQueryTerms.jsp?userdata_id=677e6385-c8f1-4db4-acb1-4b26e7d30bc3&feature=pt&row=27) | OXSR1 | MM15944 | WP COMPLEMENT ACTIVATION CLASSICAL PATHWAY | 4.42E-10 | 3.90E-08 | 3.27E-07 | 1.09E-06 | [8](https://toppgene.cchmc.org/showQueryTerms.jsp?userdata_id=e0bdc6b9-955d-48cd-95d3-4ccb1cbe49c7&feature=pt&row=27) |
| AMN | M27436 | REACTOME PROGRAMMED CELL DEATH | 3.82E-07 | 4.74E-05 | 4.15E-04 | 1.37E-03 | [23](https://toppgene.cchmc.org/showQueryTerms.jsp?userdata_id=677e6385-c8f1-4db4-acb1-4b26e7d30bc3&feature=pt&row=28) | P3H1 | MM15676 | REACTOME NERVOUS SYSTEM DEVELOPMENT | 4.57E-10 | 3.90E-08 | 3.27E-07 | 1.13E-06 | [24](https://toppgene.cchmc.org/showQueryTerms.jsp?userdata_id=e0bdc6b9-955d-48cd-95d3-4ccb1cbe49c7&feature=pt&row=28) |
| ANKFY1 | M39520 | WP REGULATION OF ACTIN CYTOSKELETON | 3.96E-07 | 4.75E-05 | 4.17E-04 | 1.43E-03 | [19](https://toppgene.cchmc.org/showQueryTerms.jsp?userdata_id=677e6385-c8f1-4db4-acb1-4b26e7d30bc3&feature=pt&row=29) | PDCD6 | M27548 | REACTOME INFECTIOUS DISEASE | 5.43E-10 | 4.48E-08 | 3.76E-07 | 1.34E-06 | [51](https://toppgene.cchmc.org/showQueryTerms.jsp?userdata_id=e0bdc6b9-955d-48cd-95d3-4ccb1cbe49c7&feature=pt&row=29) |
| ANO6 | M27557 | REACTOME MAP2K AND MAPK ACTIVATION | 4.19E-07 | 4.87E-05 | 4.27E-04 | 1.51E-03 | [10](https://toppgene.cchmc.org/showQueryTerms.jsp?userdata_id=677e6385-c8f1-4db4-acb1-4b26e7d30bc3&feature=pt&row=30) | PFN2 | MM15022 | REACTOME EPHB MEDIATED FORWARD SIGNALING | 1.70E-09 | 1.28E-07 | 1.07E-06 | 4.19E-06 | [10](https://toppgene.cchmc.org/showQueryTerms.jsp?userdata_id=e0bdc6b9-955d-48cd-95d3-4ccb1cbe49c7&feature=pt&row=30) |
| ANPEP | M1519 | KEGG ENDOCYTOSIS | 4.35E-07 | 4.90E-05 | 4.29E-04 | 1.57E-03 | [21](https://toppgene.cchmc.org/showQueryTerms.jsp?userdata_id=677e6385-c8f1-4db4-acb1-4b26e7d30bc3&feature=pt&row=31) | PLOD1 | M47729 | KEGG MEDICUS PATHOGEN ESCHERICHIA MAP TO CDC42 SIGNALING PATHWAY | 1.70E-09 | 1.28E-07 | 1.07E-06 | 4.21E-06 | [7](https://toppgene.cchmc.org/showQueryTerms.jsp?userdata_id=e0bdc6b9-955d-48cd-95d3-4ccb1cbe49c7&feature=pt&row=31) |
| ANXA11 | MM15834 | WP REGULATION OF ACTIN CYTOSKELETON | 4.88E-07 | 5.16E-05 | 4.52E-04 | 1.76E-03 | [19](https://toppgene.cchmc.org/showQueryTerms.jsp?userdata_id=677e6385-c8f1-4db4-acb1-4b26e7d30bc3&feature=pt&row=32) | PLOD3 | M4732 | BIOCARTA LECTIN PATHWAY | 1.70E-09 | 1.28E-07 | 1.07E-06 | 4.21E-06 | [7](https://toppgene.cchmc.org/showQueryTerms.jsp?userdata_id=e0bdc6b9-955d-48cd-95d3-4ccb1cbe49c7&feature=pt&row=32) |
| ANXA2 | M41836 | REACTOME CELLULAR RESPONSE TO CHEMICAL STRESS | 4.91E-07 | 5.16E-05 | 4.52E-04 | 1.77E-03 | [23](https://toppgene.cchmc.org/showQueryTerms.jsp?userdata_id=677e6385-c8f1-4db4-acb1-4b26e7d30bc3&feature=pt&row=33) | PRXL2B | M47736 | KEGG MEDICUS PATHOGEN SALMONELLA SOPE TO RAC SIGNALING PATHWAY | 2.17E-09 | 1.58E-07 | 1.33E-06 | 5.37E-06 | [8](https://toppgene.cchmc.org/showQueryTerms.jsp?userdata_id=e0bdc6b9-955d-48cd-95d3-4ccb1cbe49c7&feature=pt&row=33) |
| ANXA3 | M27548 | REACTOME INFECTIOUS DISEASE | 5.02E-07 | 5.16E-05 | 4.52E-04 | 1.81E-03 | [63](https://toppgene.cchmc.org/showQueryTerms.jsp?userdata_id=677e6385-c8f1-4db4-acb1-4b26e7d30bc3&feature=pt&row=34) | PSMB3 | M27285 | REACTOME REGULATION OF INSULIN LIKE GROWTH FACTOR IGF TRANSPORT AND UPTAKE BY INSULIN LIKE GROWTH FACTOR BINDING PROTEINS IGFBPS | 2.85E-09 | 2.02E-07 | 1.69E-06 | 7.05E-06 | [16](https://toppgene.cchmc.org/showQueryTerms.jsp?userdata_id=e0bdc6b9-955d-48cd-95d3-4ccb1cbe49c7&feature=pt&row=34) |
| ANXA4 | MM14949 | REACTOME HSP90 CHAPERONE CYCLE FOR STEROID HORMONE RECEPTORS SHR IN THE PRESENCE OF LIGAND | 6.84E-07 | 6.84E-05 | 6.00E-04 | 2.46E-03 | [11](https://toppgene.cchmc.org/showQueryTerms.jsp?userdata_id=677e6385-c8f1-4db4-acb1-4b26e7d30bc3&feature=pt&row=35) | RAB22A | MM14637 | REACTOME COLLAGEN BIOSYNTHESIS AND MODIFYING ENZYMES | 4.48E-09 | 3.08E-07 | 2.58E-06 | 1.11E-05 | [12](https://toppgene.cchmc.org/showQueryTerms.jsp?userdata_id=e0bdc6b9-955d-48cd-95d3-4ccb1cbe49c7&feature=pt&row=35) |
| ANXA5 | M22064 | BIOCARTA EEA1 PATHWAY | 7.05E-07 | 6.86E-05 | 6.01E-04 | 2.54E-03 | [5](https://toppgene.cchmc.org/showQueryTerms.jsp?userdata_id=677e6385-c8f1-4db4-acb1-4b26e7d30bc3&feature=pt&row=36) | RARS1 | M5539 | KEGG AXON GUIDANCE | 5.12E-09 | 3.42E-07 | 2.87E-06 | 1.27E-05 | [16](https://toppgene.cchmc.org/showQueryTerms.jsp?userdata_id=e0bdc6b9-955d-48cd-95d3-4ccb1cbe49c7&feature=pt&row=36) |
| ANXA6 | MM14563 | REACTOME METABOLISM | 8.17E-07 | 7.74E-05 | 6.79E-04 | 2.94E-03 | [95](https://toppgene.cchmc.org/showQueryTerms.jsp?userdata_id=677e6385-c8f1-4db4-acb1-4b26e7d30bc3&feature=pt&row=37) | SH3 | M7146 | BIOCARTA CLASSIC PATHWAY | 6.18E-09 | 4.02E-07 | 3.37E-06 | 1.53E-05 | [7](https://toppgene.cchmc.org/showQueryTerms.jsp?userdata_id=e0bdc6b9-955d-48cd-95d3-4ccb1cbe49c7&feature=pt&row=37) |
| ANXA7 | M29853 | REACTOME NERVOUS SYSTEM DEVELOPMENT | 9.63E-07 | 8.90E-05 | 7.80E-04 | 3.47E-03 | [42](https://toppgene.cchmc.org/showQueryTerms.jsp?userdata_id=677e6385-c8f1-4db4-acb1-4b26e7d30bc3&feature=pt&row=38) | SLC22A11 | M26999 | REACTOME COLLAGEN BIOSYNTHESIS AND MODIFYING ENZYMES | 6.44E-09 | 4.06E-07 | 3.41E-06 | 1.59E-05 | [12](https://toppgene.cchmc.org/showQueryTerms.jsp?userdata_id=e0bdc6b9-955d-48cd-95d3-4ccb1cbe49c7&feature=pt&row=38) |
| AOX1 | M27753 | REACTOME CLATHRIN MEDIATED ENDOCYTOSIS | 1.08E-06 | 9.73E-05 | 8.53E-04 | 3.89E-03 | [18](https://toppgene.cchmc.org/showQueryTerms.jsp?userdata_id=677e6385-c8f1-4db4-acb1-4b26e7d30bc3&feature=pt&row=39) | SPAG9 | M27308 | REACTOME EPHB MEDIATED FORWARD SIGNALING | 6.57E-09 | 4.06E-07 | 3.41E-06 | 1.62E-05 | [10](https://toppgene.cchmc.org/showQueryTerms.jsp?userdata_id=e0bdc6b9-955d-48cd-95d3-4ccb1cbe49c7&feature=pt&row=39) |
| AP1M2 | M18 | PID INTEGRIN1 PATHWAY | 1.20E-06 | 1.05E-04 | 9.22E-04 | 4.31E-03 | [12](https://toppgene.cchmc.org/showQueryTerms.jsp?userdata_id=677e6385-c8f1-4db4-acb1-4b26e7d30bc3&feature=pt&row=40) | STK24 | M724 | REACTOME RESPONSE TO ELEVATED PLATELET CYTOSOLIC CA2 | 7.18E-09 | 4.33E-07 | 3.63E-06 | 1.77E-05 | [16](https://toppgene.cchmc.org/showQueryTerms.jsp?userdata_id=e0bdc6b9-955d-48cd-95d3-4ccb1cbe49c7&feature=pt&row=40) |
| AP2M1 | MM1434 | BIOCARTA MCALPAIN PATHWAY | 1.31E-06 | 1.12E-04 | 9.85E-04 | 4.72E-03 | [7](https://toppgene.cchmc.org/showQueryTerms.jsp?userdata_id=677e6385-c8f1-4db4-acb1-4b26e7d30bc3&feature=pt&row=41) | TKT | MM14786 | REACTOME MEMBRANE TRAFFICKING | 1.66E-08 | 9.62E-07 | 8.07E-06 | 4.10E-05 | [34](https://toppgene.cchmc.org/showQueryTerms.jsp?userdata_id=e0bdc6b9-955d-48cd-95d3-4ccb1cbe49c7&feature=pt&row=41) |
| APAF1 | M48078 | WP METABOLIC EPILEPTIC DISORDERS | 1.48E-06 | 1.24E-04 | 1.09E-03 | 5.34E-03 | [14](https://toppgene.cchmc.org/showQueryTerms.jsp?userdata_id=677e6385-c8f1-4db4-acb1-4b26e7d30bc3&feature=pt&row=42) | TPP1 | M198 | PID SYNDECAN 1 PATHWAY | 1.70E-08 | 9.62E-07 | 8.07E-06 | 4.19E-05 | [10](https://toppgene.cchmc.org/showQueryTerms.jsp?userdata_id=e0bdc6b9-955d-48cd-95d3-4ccb1cbe49c7&feature=pt&row=42) |
| APBA2 | M27078 | REACTOME RHO GTPASE CYCLE | 1.63E-06 | 1.33E-04 | 1.17E-03 | 5.87E-03 | [35](https://toppgene.cchmc.org/showQueryTerms.jsp?userdata_id=677e6385-c8f1-4db4-acb1-4b26e7d30bc3&feature=pt&row=43) | VPS36 | M47715 | KEGG MEDICUS REFERENCE ARNO ARF ACTB G SIGNALING PATHWAY | 1.71E-08 | 9.62E-07 | 8.07E-06 | 4.23E-05 | [8](https://toppgene.cchmc.org/showQueryTerms.jsp?userdata_id=e0bdc6b9-955d-48cd-95d3-4ccb1cbe49c7&feature=pt&row=43) |
| APOE | MM14921 | REACTOME NUCLEAR ENVELOPE NE REASSEMBLY | 1.67E-06 | 1.34E-04 | 1.17E-03 | 6.01E-03 | [12](https://toppgene.cchmc.org/showQueryTerms.jsp?userdata_id=677e6385-c8f1-4db4-acb1-4b26e7d30bc3&feature=pt&row=44) | ACO1 | M42535 | WP COMPLEMENT SYSTEM IN NEURONAL DEVELOPMENT AND PLASTICITY | 2.10E-08 | 1.13E-06 | 9.47E-06 | 5.19E-05 | [14](https://toppgene.cchmc.org/showQueryTerms.jsp?userdata_id=e0bdc6b9-955d-48cd-95d3-4ccb1cbe49c7&feature=pt&row=44) |
| ARG1 | M27251 | REACTOME HSP90 CHAPERONE CYCLE FOR STEROID HORMONE RECEPTORS SHR IN THE PRESENCE OF LIGAND | 1.80E-06 | 1.41E-04 | 1.24E-03 | 6.50E-03 | [11](https://toppgene.cchmc.org/showQueryTerms.jsp?userdata_id=677e6385-c8f1-4db4-acb1-4b26e7d30bc3&feature=pt&row=45) | ACTR1A | MM14571 | REACTOME DEGRADATION OF THE EXTRACELLULAR MATRIX | 2.10E-08 | 1.13E-06 | 9.47E-06 | 5.19E-05 | [14](https://toppgene.cchmc.org/showQueryTerms.jsp?userdata_id=e0bdc6b9-955d-48cd-95d3-4ccb1cbe49c7&feature=pt&row=45) |
| ARHGEF12 | M39581 | WP COMPLEMENT SYSTEM | 1.93E-06 | 1.48E-04 | 1.30E-03 | 6.96E-03 | [14](https://toppgene.cchmc.org/showQueryTerms.jsp?userdata_id=677e6385-c8f1-4db4-acb1-4b26e7d30bc3&feature=pt&row=46) | ACTR2 | MM15232 | REACTOME VESICLE MEDIATED TRANSPORT | 2.27E-08 | 1.19E-06 | 1.00E-05 | 5.61E-05 | [36](https://toppgene.cchmc.org/showQueryTerms.jsp?userdata_id=e0bdc6b9-955d-48cd-95d3-4ccb1cbe49c7&feature=pt&row=46) |
| ARL3 | M39402 | WP FOCAL ADHESION | 2.08E-06 | 1.56E-04 | 1.37E-03 | 7.49E-03 | [21](https://toppgene.cchmc.org/showQueryTerms.jsp?userdata_id=677e6385-c8f1-4db4-acb1-4b26e7d30bc3&feature=pt&row=47) | ACTR3 | M27103 | REACTOME ASSEMBLY OF COLLAGEN FIBRILS AND OTHER MULTIMERIC STRUCTURES | 2.59E-08 | 1.33E-06 | 1.11E-05 | 6.40E-05 | [11](https://toppgene.cchmc.org/showQueryTerms.jsp?userdata_id=e0bdc6b9-955d-48cd-95d3-4ccb1cbe49c7&feature=pt&row=47) |
| ARL8A | M27287 | REACTOME TRANSPORT OF SMALL MOLECULES | 2.13E-06 | 1.56E-04 | 1.37E-03 | 7.67E-03 | [48](https://toppgene.cchmc.org/showQueryTerms.jsp?userdata_id=677e6385-c8f1-4db4-acb1-4b26e7d30bc3&feature=pt&row=48) | AHCYL1 | MM14796 | REACTOME ASSEMBLY OF COLLAGEN FIBRILS AND OTHER MULTIMERIC STRUCTURES | 2.63E-08 | 1.33E-06 | 1.11E-05 | 6.50E-05 | [10](https://toppgene.cchmc.org/showQueryTerms.jsp?userdata_id=e0bdc6b9-955d-48cd-95d3-4ccb1cbe49c7&feature=pt&row=48) |
| ARMC3 | MM1390 | BIOCARTA ECM PATHWAY | 2.86E-06 | 2.04E-04 | 1.79E-03 | 1.03E-02 | [7](https://toppgene.cchmc.org/showQueryTerms.jsp?userdata_id=677e6385-c8f1-4db4-acb1-4b26e7d30bc3&feature=pt&row=49) | ALDH9A1 | M39737 | WP INHIBITION OF EXOSOME BIOGENESIS AND SECRETION BY MANUMYCIN A IN CRPC CELLS | 2.90E-08 | 1.44E-06 | 1.20E-05 | 7.18E-05 | [7](https://toppgene.cchmc.org/showQueryTerms.jsp?userdata_id=e0bdc6b9-955d-48cd-95d3-4ccb1cbe49c7&feature=pt&row=49) |
| ARSF | M7098 | KEGG ECM RECEPTOR INTERACTION | 2.89E-06 | 2.04E-04 | 1.79E-03 | 1.04E-02 | [13](https://toppgene.cchmc.org/showQueryTerms.jsp?userdata_id=677e6385-c8f1-4db4-acb1-4b26e7d30bc3&feature=pt&row=50) | ANGPTL2 | MM15455 | REACTOME RESPONSE TO ELEVATED PLATELET CYTOSOLIC CA2 | 4.19E-08 | 2.03E-06 | 1.71E-05 | 1.04E-04 | [15](https://toppgene.cchmc.org/showQueryTerms.jsp?userdata_id=e0bdc6b9-955d-48cd-95d3-4ccb1cbe49c7&feature=pt&row=50) |
| ASAH1 | M16441 | REACTOME INTEGRIN CELL SURFACE INTERACTIONS | 3.32E-06 | 2.30E-04 | 2.01E-03 | 1.19E-02 | [13](https://toppgene.cchmc.org/showQueryTerms.jsp?userdata_id=677e6385-c8f1-4db4-acb1-4b26e7d30bc3&feature=pt&row=51) | ANXA1 | M194 | BIOCARTA PROTEASOME PATHWAY | 4.52E-08 | 2.15E-06 | 1.80E-05 | 1.12E-04 | [7](https://toppgene.cchmc.org/showQueryTerms.jsp?userdata_id=e0bdc6b9-955d-48cd-95d3-4ccb1cbe49c7&feature=pt&row=51) |
| ASAT | MM14867 | REACTOME INTEGRIN CELL SURFACE INTERACTIONS | 3.64E-06 | 2.47E-04 | 2.17E-03 | 1.31E-02 | [12](https://toppgene.cchmc.org/showQueryTerms.jsp?userdata_id=677e6385-c8f1-4db4-acb1-4b26e7d30bc3&feature=pt&row=52) | APRT | M47716 | KEGG MEDICUS PATHOGEN SHIGELLA IPGD TO ARNO ARF ACTB G SIGNALING PATHWAY | 6.34E-08 | 2.93E-06 | 2.46E-05 | 1.57E-04 | [8](https://toppgene.cchmc.org/showQueryTerms.jsp?userdata_id=e0bdc6b9-955d-48cd-95d3-4ccb1cbe49c7&feature=pt&row=52) |
| ASL | M39693 | WP EBOLA VIRUS INFECTION IN HOST | 4.32E-06 | 2.83E-04 | 2.48E-03 | 1.56E-02 | [16](https://toppgene.cchmc.org/showQueryTerms.jsp?userdata_id=677e6385-c8f1-4db4-acb1-4b26e7d30bc3&feature=pt&row=53) | ARF1 | M62 | PID EPHB FWD PATHWAY | 6.51E-08 | 2.93E-06 | 2.46E-05 | 1.61E-04 | [9](https://toppgene.cchmc.org/showQueryTerms.jsp?userdata_id=e0bdc6b9-955d-48cd-95d3-4ccb1cbe49c7&feature=pt&row=53) |
| ATAD2 | M186 | PID PDGFRB PATHWAY | 4.32E-06 | 2.83E-04 | 2.48E-03 | 1.56E-02 | [16](https://toppgene.cchmc.org/showQueryTerms.jsp?userdata_id=677e6385-c8f1-4db4-acb1-4b26e7d30bc3&feature=pt&row=54) | ARHGDIB | M47747 | KEGG MEDICUS VARIANT MUTATION INACTIVATED VCP TO 26S PROTEASOME MEDIATED PROTEIN DEGRADATION | 6.51E-08 | 2.93E-06 | 2.46E-05 | 1.61E-04 | [9](https://toppgene.cchmc.org/showQueryTerms.jsp?userdata_id=e0bdc6b9-955d-48cd-95d3-4ccb1cbe49c7&feature=pt&row=54) |
| ATIC | M1009 | REACTOME ENDOSOMAL SORTING COMPLEX REQUIRED FOR TRANSPORT ESCRT | 4.77E-06 | 3.01E-04 | 2.64E-03 | 1.72E-02 | [8](https://toppgene.cchmc.org/showQueryTerms.jsp?userdata_id=677e6385-c8f1-4db4-acb1-4b26e7d30bc3&feature=pt&row=55) | ARPC2 | M47746 | KEGG MEDICUS VARIANT MUTATION CAUSED ABERRANT SOD1 TO 26S PROTEASOME MEDIATED PROTEIN DEGRADATION | 8.20E-08 | 3.62E-06 | 3.04E-05 | 2.03E-04 | [9](https://toppgene.cchmc.org/showQueryTerms.jsp?userdata_id=e0bdc6b9-955d-48cd-95d3-4ccb1cbe49c7&feature=pt&row=55) |
| ATP5F1A | MM15637 | REACTOME ENDOSOMAL SORTING COMPLEX REQUIRED FOR TRANSPORT ESCRT | 4.77E-06 | 3.01E-04 | 2.64E-03 | 1.72E-02 | [8](https://toppgene.cchmc.org/showQueryTerms.jsp?userdata_id=677e6385-c8f1-4db4-acb1-4b26e7d30bc3&feature=pt&row=56) | ARPC4 | M47737 | KEGG MEDICUS PATHOGEN SALMONELLA SOPB TO ARNO ARF ACTB G SIGNALING PATHWAY | 8.50E-08 | 3.69E-06 | 3.10E-05 | 2.10E-04 | [8](https://toppgene.cchmc.org/showQueryTerms.jsp?userdata_id=e0bdc6b9-955d-48cd-95d3-4ccb1cbe49c7&feature=pt&row=56) |
| ATP5F1B | MM14855 | REACTOME MHC CLASS II ANTIGEN PRESENTATION | 5.12E-06 | 3.18E-04 | 2.78E-03 | 1.84E-02 | [15](https://toppgene.cchmc.org/showQueryTerms.jsp?userdata_id=677e6385-c8f1-4db4-acb1-4b26e7d30bc3&feature=pt&row=57) | ATP6AP1 | M47722 | KEGG MEDICUS PATHOGEN SHIGELLA IPAC TO ACTIN SIGNALING PATHWAY | 8.67E-08 | 3.70E-06 | 3.10E-05 | 2.14E-04 | [6](https://toppgene.cchmc.org/showQueryTerms.jsp?userdata_id=e0bdc6b9-955d-48cd-95d3-4ccb1cbe49c7&feature=pt&row=57) |
| ATP6V0A1 | MM15830 | WP INTEGRIN MEDIATED CELL ADHESION | 5.20E-06 | 3.18E-04 | 2.78E-03 | 1.87E-02 | [14](https://toppgene.cchmc.org/showQueryTerms.jsp?userdata_id=677e6385-c8f1-4db4-acb1-4b26e7d30bc3&feature=pt&row=58) | BASP1 | MM14537 | REACTOME DEVELOPMENTAL BIOLOGY | 9.87E-08 | 4.14E-06 | 3.47E-05 | 2.44E-04 | [28](https://toppgene.cchmc.org/showQueryTerms.jsp?userdata_id=e0bdc6b9-955d-48cd-95d3-4ccb1cbe49c7&feature=pt&row=58) |
| ATP6V0A4 | M27215 | REACTOME NUCLEAR ENVELOPE NE REASSEMBLY | 5.62E-06 | 3.31E-04 | 2.90E-03 | 2.02E-02 | [12](https://toppgene.cchmc.org/showQueryTerms.jsp?userdata_id=677e6385-c8f1-4db4-acb1-4b26e7d30bc3&feature=pt&row=59) | BGN | M48087 | WP 17P13 3 YWHAE COPY NUMBER VARIATION | 1.01E-07 | 4.15E-06 | 3.48E-05 | 2.49E-04 | [7](https://toppgene.cchmc.org/showQueryTerms.jsp?userdata_id=e0bdc6b9-955d-48cd-95d3-4ccb1cbe49c7&feature=pt&row=59) |
| ATP6V0C | M47719 | KEGG MEDICUS REFERENCE ITGA B RHOGEF RHOA SIGNALING PATHWAY | 5.69E-06 | 3.31E-04 | 2.90E-03 | 2.05E-02 | [7](https://toppgene.cchmc.org/showQueryTerms.jsp?userdata_id=677e6385-c8f1-4db4-acb1-4b26e7d30bc3&feature=pt&row=60) | CAP1 | M16441 | REACTOME INTEGRIN CELL SURFACE INTERACTIONS | 1.03E-07 | 4.15E-06 | 3.48E-05 | 2.55E-04 | [12](https://toppgene.cchmc.org/showQueryTerms.jsp?userdata_id=e0bdc6b9-955d-48cd-95d3-4ccb1cbe49c7&feature=pt&row=60) |
| ATP6V0D2 | M47536 | KEGG MEDICUS REFERENCE ITGA B RHOGAP RHOA SIGNALING PATHWAY | 5.69E-06 | 3.31E-04 | 2.90E-03 | 2.05E-02 | [7](https://toppgene.cchmc.org/showQueryTerms.jsp?userdata_id=677e6385-c8f1-4db4-acb1-4b26e7d30bc3&feature=pt&row=61) | CAPG | M39613 | WP PATHOGENIC ESCHERICHIA COLI INFECTION | 1.04E-07 | 4.15E-06 | 3.48E-05 | 2.57E-04 | [10](https://toppgene.cchmc.org/showQueryTerms.jsp?userdata_id=e0bdc6b9-955d-48cd-95d3-4ccb1cbe49c7&feature=pt&row=61) |
| ATP6V1A | M27771 | REACTOME RAB GERANYLGERANYLATION | 6.92E-06 | 3.90E-04 | 3.42E-03 | 2.49E-02 | [11](https://toppgene.cchmc.org/showQueryTerms.jsp?userdata_id=677e6385-c8f1-4db4-acb1-4b26e7d30bc3&feature=pt&row=62) | CAPN1 | M587 | REACTOME DEGRADATION OF THE EXTRACELLULAR MATRIX | 1.14E-07 | 4.46E-06 | 3.74E-05 | 2.81E-04 | [15](https://toppgene.cchmc.org/showQueryTerms.jsp?userdata_id=e0bdc6b9-955d-48cd-95d3-4ccb1cbe49c7&feature=pt&row=62) |
| ATP6V1H | MM15511 | REACTOME RAB GERANYLGERANYLATION | 6.92E-06 | 3.90E-04 | 3.42E-03 | 2.49E-02 | [11](https://toppgene.cchmc.org/showQueryTerms.jsp?userdata_id=677e6385-c8f1-4db4-acb1-4b26e7d30bc3&feature=pt&row=63) | CAPN2 | M2333 | KEGG PATHOGENIC ESCHERICHIA COLI INFECTION | 1.25E-07 | 4.81E-06 | 4.04E-05 | 3.08E-04 | [10](https://toppgene.cchmc.org/showQueryTerms.jsp?userdata_id=e0bdc6b9-955d-48cd-95d3-4ccb1cbe49c7&feature=pt&row=63) |
| ATRN | MM15595 | REACTOME RHO GTPASE CYCLE | 7.07E-06 | 3.92E-04 | 3.43E-03 | 2.55E-02 | [33](https://toppgene.cchmc.org/showQueryTerms.jsp?userdata_id=677e6385-c8f1-4db4-acb1-4b26e7d30bc3&feature=pt&row=64) | CC2D1A | M39480 | WP BLOOD CLOTTING CASCADE | 1.45E-07 | 5.53E-06 | 4.64E-05 | 3.59E-04 | [7](https://toppgene.cchmc.org/showQueryTerms.jsp?userdata_id=e0bdc6b9-955d-48cd-95d3-4ccb1cbe49c7&feature=pt&row=64) |
| B4GALT1 | M164 | PID ERBB1 DOWNSTREAM PATHWAY | 7.35E-06 | 4.01E-04 | 3.51E-03 | 2.65E-02 | [14](https://toppgene.cchmc.org/showQueryTerms.jsp?userdata_id=677e6385-c8f1-4db4-acb1-4b26e7d30bc3&feature=pt&row=65) | CCT2 | M47728 | KEGG MEDICUS PATHOGEN ESCHERICHIA EAE TIR TCCP TO ACTIN SIGNALING PATHWAY | 1.49E-07 | 5.59E-06 | 4.69E-05 | 3.69E-04 | [6](https://toppgene.cchmc.org/showQueryTerms.jsp?userdata_id=e0bdc6b9-955d-48cd-95d3-4ccb1cbe49c7&feature=pt&row=65) |
| BAIAP2 | M27299 | REACTOME FOLDING OF ACTIN BY CCT TRIC | 7.80E-06 | 4.13E-04 | 3.62E-03 | 2.81E-02 | [5](https://toppgene.cchmc.org/showQueryTerms.jsp?userdata_id=677e6385-c8f1-4db4-acb1-4b26e7d30bc3&feature=pt&row=66) | CCT6A | M26954 | REACTOME TRANSLOCATION OF SLC2A4 GLUT4 TO THE PLASMA MEMBRANE | 1.54E-07 | 5.59E-06 | 4.69E-05 | 3.82E-04 | [11](https://toppgene.cchmc.org/showQueryTerms.jsp?userdata_id=e0bdc6b9-955d-48cd-95d3-4ccb1cbe49c7&feature=pt&row=66) |
| BAIAP2L1 | MM15004 | REACTOME ASSOCIATION OF TRIC CCT WITH TARGET PROTEINS DURING BIOSYNTHESIS | 7.80E-06 | 4.13E-04 | 3.62E-03 | 2.81E-02 | [5](https://toppgene.cchmc.org/showQueryTerms.jsp?userdata_id=677e6385-c8f1-4db4-acb1-4b26e7d30bc3&feature=pt&row=67) | CDC42 | M27006 | REACTOME TERMINAL PATHWAY OF COMPLEMENT | 1.56E-07 | 5.59E-06 | 4.69E-05 | 3.86E-04 | [5](https://toppgene.cchmc.org/showQueryTerms.jsp?userdata_id=e0bdc6b9-955d-48cd-95d3-4ccb1cbe49c7&feature=pt&row=67) |
| BBOX1 | MM1426 | BIOCARTA INTEGRIN PATHWAY | 7.92E-06 | 4.14E-04 | 3.63E-03 | 2.85E-02 | [8](https://toppgene.cchmc.org/showQueryTerms.jsp?userdata_id=677e6385-c8f1-4db4-acb1-4b26e7d30bc3&feature=pt&row=68) | CLIC1 | MM14656 | REACTOME TERMINAL PATHWAY OF COMPLEMENT | 1.56E-07 | 5.59E-06 | 4.69E-05 | 3.86E-04 | [5](https://toppgene.cchmc.org/showQueryTerms.jsp?userdata_id=e0bdc6b9-955d-48cd-95d3-4ccb1cbe49c7&feature=pt&row=68) |
| BEND7 | M3342 | BIOCARTA INTEGRIN PATHWAY | 1.01E-05 | 5.11E-04 | 4.48E-03 | 3.63E-02 | [8](https://toppgene.cchmc.org/showQueryTerms.jsp?userdata_id=677e6385-c8f1-4db4-acb1-4b26e7d30bc3&feature=pt&row=69) | CLIC4 | M518 | REACTOME ANTIGEN PROCESSING CROSS PRESENTATION | 1.65E-07 | 5.84E-06 | 4.90E-05 | 4.08E-04 | [13](https://toppgene.cchmc.org/showQueryTerms.jsp?userdata_id=e0bdc6b9-955d-48cd-95d3-4ccb1cbe49c7&feature=pt&row=69) |
| BLVRA | M27942 | REACTOME LATE ENDOSOMAL MICROAUTOPHAGY | 1.01E-05 | 5.11E-04 | 4.48E-03 | 3.63E-02 | [8](https://toppgene.cchmc.org/showQueryTerms.jsp?userdata_id=677e6385-c8f1-4db4-acb1-4b26e7d30bc3&feature=pt&row=70) | CNDP2 | MM15285 | REACTOME MAPK6 MAPK4 SIGNALING | 1.79E-07 | 6.22E-06 | 5.22E-05 | 4.42E-04 | [11](https://toppgene.cchmc.org/showQueryTerms.jsp?userdata_id=e0bdc6b9-955d-48cd-95d3-4ccb1cbe49c7&feature=pt&row=70) |
| BLVRB | MM14968 | REACTOME L1CAM INTERACTIONS | 1.09E-05 | 5.38E-04 | 4.72E-03 | 3.91E-02 | [11](https://toppgene.cchmc.org/showQueryTerms.jsp?userdata_id=677e6385-c8f1-4db4-acb1-4b26e7d30bc3&feature=pt&row=71) | COL15A1 | MM15913 | WP FOCAL ADHESION | 1.81E-07 | 6.23E-06 | 5.22E-05 | 4.48E-04 | [17](https://toppgene.cchmc.org/showQueryTerms.jsp?userdata_id=e0bdc6b9-955d-48cd-95d3-4ccb1cbe49c7&feature=pt&row=71) |
| BROX | M99 | PID TXA2PATHWAY | 1.11E-05 | 5.38E-04 | 4.72E-03 | 3.98E-02 | [10](https://toppgene.cchmc.org/showQueryTerms.jsp?userdata_id=677e6385-c8f1-4db4-acb1-4b26e7d30bc3&feature=pt&row=72) | CRYAB | MM15265 | REACTOME RHO GTPASES ACTIVATE WASPS AND WAVES | 1.92E-07 | 6.49E-06 | 5.45E-05 | 4.74E-04 | [8](https://toppgene.cchmc.org/showQueryTerms.jsp?userdata_id=e0bdc6b9-955d-48cd-95d3-4ccb1cbe49c7&feature=pt&row=72) |
| C16orf89 | MM15638 | REACTOME IRON UPTAKE AND TRANSPORT | 1.11E-05 | 5.38E-04 | 4.72E-03 | 3.98E-02 | [10](https://toppgene.cchmc.org/showQueryTerms.jsp?userdata_id=677e6385-c8f1-4db4-acb1-4b26e7d30bc3&feature=pt&row=73) | CSE1L | MM15936 | WP BLOOD CLOTTING CASCADE | 2.05E-07 | 6.80E-06 | 5.70E-05 | 5.08E-04 | [7](https://toppgene.cchmc.org/showQueryTerms.jsp?userdata_id=e0bdc6b9-955d-48cd-95d3-4ccb1cbe49c7&feature=pt&row=73) |
| C17orf80 | M127 | PID ERBB1 RECEPTOR PROXIMAL PATHWAY | 1.27E-05 | 6.09E-04 | 5.34E-03 | 4.57E-02 | [8](https://toppgene.cchmc.org/showQueryTerms.jsp?userdata_id=677e6385-c8f1-4db4-acb1-4b26e7d30bc3&feature=pt&row=74) | CYSTM1 | M160 | PID AVB3 INTEGRIN PATHWAY | 2.06E-07 | 6.80E-06 | 5.70E-05 | 5.10E-04 | [11](https://toppgene.cchmc.org/showQueryTerms.jsp?userdata_id=e0bdc6b9-955d-48cd-95d3-4ccb1cbe49c7&feature=pt&row=74) |
| C19orf18 | M45021 | REACTOME KEAP1 NFE2L2 PATHWAY | 1.29E-05 | 6.11E-04 | 5.36E-03 | 4.65E-02 | [15](https://toppgene.cchmc.org/showQueryTerms.jsp?userdata_id=677e6385-c8f1-4db4-acb1-4b26e7d30bc3&feature=pt&row=75) | DDAH2 | M18306 | KEGG REGULATION OF ACTIN CYTOSKELETON | 2.37E-07 | 7.70E-06 | 6.46E-05 | 5.85E-04 | [18](https://toppgene.cchmc.org/showQueryTerms.jsp?userdata_id=e0bdc6b9-955d-48cd-95d3-4ccb1cbe49c7&feature=pt&row=75) |
| C5 | M45026 | REACTOME NUCLEAR EVENTS MEDIATED BY NFE2L2 | 1.31E-05 | 6.13E-04 | 5.37E-03 | 4.72E-02 | [13](https://toppgene.cchmc.org/showQueryTerms.jsp?userdata_id=677e6385-c8f1-4db4-acb1-4b26e7d30bc3&feature=pt&row=76) | DSTN | M186 | PID PDGFRB PATHWAY | 2.58E-07 | 8.29E-06 | 6.96E-05 | 6.38E-04 | [14](https://toppgene.cchmc.org/showQueryTerms.jsp?userdata_id=e0bdc6b9-955d-48cd-95d3-4ccb1cbe49c7&feature=pt&row=76) |
| C9 | M9648 | REACTOME FORMATION OF TUBULIN FOLDING INTERMEDIATES BY CCT TRIC | 1.40E-05 | 6.40E-04 | 5.61E-03 | 5.05E-02 | [7](https://toppgene.cchmc.org/showQueryTerms.jsp?userdata_id=677e6385-c8f1-4db4-acb1-4b26e7d30bc3&feature=pt&row=77) | EHBP1 | M27572 | REACTOME MAPK6 MAPK4 SIGNALING | 2.83E-07 | 8.98E-06 | 7.54E-05 | 7.01E-04 | [12](https://toppgene.cchmc.org/showQueryTerms.jsp?userdata_id=e0bdc6b9-955d-48cd-95d3-4ccb1cbe49c7&feature=pt&row=77) |
| CAB39L | M239 | PID A6B1 A6B4 INTEGRIN PATHWAY | 1.41E-05 | 6.40E-04 | 5.61E-03 | 5.07E-02 | [9](https://toppgene.cchmc.org/showQueryTerms.jsp?userdata_id=677e6385-c8f1-4db4-acb1-4b26e7d30bc3&feature=pt&row=78) | G6PD | M47880 | KEGG MEDICUS REFERENCE REGULATION OF COMPLEMENT CASCADE MAC INHIBITION | 3.45E-07 | 1.08E-05 | 9.06E-05 | 8.53E-04 | [5](https://toppgene.cchmc.org/showQueryTerms.jsp?userdata_id=e0bdc6b9-955d-48cd-95d3-4ccb1cbe49c7&feature=pt&row=78) |
| CAMK4 | M705 | REACTOME MHC CLASS II ANTIGEN PRESENTATION | 1.42E-05 | 6.40E-04 | 5.61E-03 | 5.12E-02 | [15](https://toppgene.cchmc.org/showQueryTerms.jsp?userdata_id=677e6385-c8f1-4db4-acb1-4b26e7d30bc3&feature=pt&row=79) | GAPDH | MM14653 | REACTOME COMPLEMENT CASCADE | 3.59E-07 | 1.11E-05 | 9.31E-05 | 8.87E-04 | [12](https://toppgene.cchmc.org/showQueryTerms.jsp?userdata_id=e0bdc6b9-955d-48cd-95d3-4ccb1cbe49c7&feature=pt&row=79) |
| CAMP | MM15499 | REACTOME CLATHRIN MEDIATED ENDOCYTOSIS | 1.50E-05 | 6.62E-04 | 5.81E-03 | 5.39E-02 | [16](https://toppgene.cchmc.org/showQueryTerms.jsp?userdata_id=677e6385-c8f1-4db4-acb1-4b26e7d30bc3&feature=pt&row=80) | GLIPR2 | M27549 | REACTOME RHO GTPASES ACTIVATE WASPS AND WAVES | 3.97E-07 | 1.15E-05 | 9.68E-05 | 9.81E-04 | [8](https://toppgene.cchmc.org/showQueryTerms.jsp?userdata_id=e0bdc6b9-955d-48cd-95d3-4ccb1cbe49c7&feature=pt&row=80) |
| CAND1 | M27827 | REACTOME CELLULAR RESPONSES TO STIMULI | 1.51E-05 | 6.62E-04 | 5.81E-03 | 5.43E-02 | [50](https://toppgene.cchmc.org/showQueryTerms.jsp?userdata_id=677e6385-c8f1-4db4-acb1-4b26e7d30bc3&feature=pt&row=81) | GNA11 | M47757 | KEGG MEDICUS VARIANT SCRAPIE CONFORMATION PRPSC TO 26S PROTEASOME MEDIATED PROTEIN DEGRADATION | 3.97E-07 | 1.15E-05 | 9.68E-05 | 9.81E-04 | [8](https://toppgene.cchmc.org/showQueryTerms.jsp?userdata_id=e0bdc6b9-955d-48cd-95d3-4ccb1cbe49c7&feature=pt&row=81) |
| CAPN5 | M962 | REACTOME IRON UPTAKE AND TRANSPORT | 1.53E-05 | 6.63E-04 | 5.81E-03 | 5.51E-02 | [10](https://toppgene.cchmc.org/showQueryTerms.jsp?userdata_id=677e6385-c8f1-4db4-acb1-4b26e7d30bc3&feature=pt&row=82) | GNAI1 | M47712 | KEGG MEDICUS VARIANT MUTATION CAUSED ABERRANT ABETA TO 26S PROTEASOME MEDIATED PROTEIN DEGRADATION | 3.97E-07 | 1.15E-05 | 9.68E-05 | 9.81E-04 | [8](https://toppgene.cchmc.org/showQueryTerms.jsp?userdata_id=e0bdc6b9-955d-48cd-95d3-4ccb1cbe49c7&feature=pt&row=82) |
| CAPN7 | MM1577 | BIOCARTA EEA1 PATHWAY | 1.57E-05 | 6.71E-04 | 5.89E-03 | 5.64E-02 | [4](https://toppgene.cchmc.org/showQueryTerms.jsp?userdata_id=677e6385-c8f1-4db4-acb1-4b26e7d30bc3&feature=pt&row=83) | GNAI2 | M47713 | KEGG MEDICUS VARIANT MUTATION CAUSED ABERRANT HTT TO 26S PROTEASOME MEDIATED PROTEIN DEGRADATION | 3.97E-07 | 1.15E-05 | 9.68E-05 | 9.81E-04 | [8](https://toppgene.cchmc.org/showQueryTerms.jsp?userdata_id=e0bdc6b9-955d-48cd-95d3-4ccb1cbe49c7&feature=pt&row=83) |
| CAPZA2 | M39006 | REACTOME SARS COV 1 INFECTION | 1.64E-05 | 6.85E-04 | 6.01E-03 | 5.89E-02 | [16](https://toppgene.cchmc.org/showQueryTerms.jsp?userdata_id=677e6385-c8f1-4db4-acb1-4b26e7d30bc3&feature=pt&row=84) | GNAI3 | M47701 | KEGG MEDICUS VARIANT MUTATION CAUSED ABERRANT SNCA TO 26S PROTEASOME MEDIATED PROTEIN DEGRADATION | 3.97E-07 | 1.15E-05 | 9.68E-05 | 9.81E-04 | [8](https://toppgene.cchmc.org/showQueryTerms.jsp?userdata_id=e0bdc6b9-955d-48cd-95d3-4ccb1cbe49c7&feature=pt&row=84) |
| CAPZB | MM15676 | REACTOME NERVOUS SYSTEM DEVELOPMENT | 1.64E-05 | 6.85E-04 | 6.01E-03 | 5.89E-02 | [23](https://toppgene.cchmc.org/showQueryTerms.jsp?userdata_id=677e6385-c8f1-4db4-acb1-4b26e7d30bc3&feature=pt&row=85) | H2AB2 | M26953 | REACTOME COLLAGEN DEGRADATION | 4.61E-07 | 1.32E-05 | 1.11E-04 | 1.14E-03 | [10](https://toppgene.cchmc.org/showQueryTerms.jsp?userdata_id=e0bdc6b9-955d-48cd-95d3-4ccb1cbe49c7&feature=pt&row=85) |
| CCDC105 | MM15041 | REACTOME PCP CE PATHWAY | 1.83E-05 | 7.45E-04 | 6.53E-03 | 6.59E-02 | [12](https://toppgene.cchmc.org/showQueryTerms.jsp?userdata_id=677e6385-c8f1-4db4-acb1-4b26e7d30bc3&feature=pt&row=86) | H2BC18 | MM15538 | REACTOME COLLAGEN CHAIN TRIMERIZATION | 4.97E-07 | 1.41E-05 | 1.19E-04 | 1.23E-03 | [8](https://toppgene.cchmc.org/showQueryTerms.jsp?userdata_id=e0bdc6b9-955d-48cd-95d3-4ccb1cbe49c7&feature=pt&row=86) |
| CCDC180 | M760 | REACTOME INTEGRIN SIGNALING | 1.84E-05 | 7.45E-04 | 6.53E-03 | 6.63E-02 | [7](https://toppgene.cchmc.org/showQueryTerms.jsp?userdata_id=677e6385-c8f1-4db4-acb1-4b26e7d30bc3&feature=pt&row=87) | H4C4 | M509 | REACTOME DEVELOPMENTAL BIOLOGY | 5.25E-07 | 1.47E-05 | 1.24E-04 | 1.30E-03 | [51](https://toppgene.cchmc.org/showQueryTerms.jsp?userdata_id=e0bdc6b9-955d-48cd-95d3-4ccb1cbe49c7&feature=pt&row=87) |
| CCT3 | MM15674 | REACTOME SEALING OF THE NUCLEAR ENVELOPE NE BY ESCRT III | 1.84E-05 | 7.45E-04 | 6.53E-03 | 6.63E-02 | [7](https://toppgene.cchmc.org/showQueryTerms.jsp?userdata_id=677e6385-c8f1-4db4-acb1-4b26e7d30bc3&feature=pt&row=88) | HSPB1 | MM1369 | BIOCARTA COMP PATHWAY | 5.85E-07 | 1.60E-05 | 1.34E-04 | 1.45E-03 | [6](https://toppgene.cchmc.org/showQueryTerms.jsp?userdata_id=e0bdc6b9-955d-48cd-95d3-4ccb1cbe49c7&feature=pt&row=88) |
| CCT5 | M539 | REACTOME TRANS GOLGI NETWORK VESICLE BUDDING | 1.91E-05 | 7.54E-04 | 6.61E-03 | 6.86E-02 | [11](https://toppgene.cchmc.org/showQueryTerms.jsp?userdata_id=677e6385-c8f1-4db4-acb1-4b26e7d30bc3&feature=pt&row=89) | HSPD1 | M47727 | KEGG MEDICUS PATHOGEN ESCHERICHIA EAE TIR TO ACTIN SIGNALING PATHWAY | 5.85E-07 | 1.60E-05 | 1.34E-04 | 1.45E-03 | [6](https://toppgene.cchmc.org/showQueryTerms.jsp?userdata_id=e0bdc6b9-955d-48cd-95d3-4ccb1cbe49c7&feature=pt&row=89) |
| CCT8 | M39403 | WP PRIMARY FOCAL SEGMENTAL GLOMERULOSCLEROSIS FSGS | 1.91E-05 | 7.54E-04 | 6.61E-03 | 6.86E-02 | [11](https://toppgene.cchmc.org/showQueryTerms.jsp?userdata_id=677e6385-c8f1-4db4-acb1-4b26e7d30bc3&feature=pt&row=90) | JADE2 | M27311 | REACTOME EPH EPHRIN MEDIATED REPULSION OF CELLS | 5.99E-07 | 1.60E-05 | 1.34E-04 | 1.48E-03 | [9](https://toppgene.cchmc.org/showQueryTerms.jsp?userdata_id=e0bdc6b9-955d-48cd-95d3-4ccb1cbe49c7&feature=pt&row=90) |
| CCTB | M6355 | BIOCARTA ECM PATHWAY | 2.15E-05 | 8.17E-04 | 7.16E-03 | 7.76E-02 | [6](https://toppgene.cchmc.org/showQueryTerms.jsp?userdata_id=677e6385-c8f1-4db4-acb1-4b26e7d30bc3&feature=pt&row=91) | LDHA | MM14566 | REACTOME COLLAGEN DEGRADATION | 5.99E-07 | 1.60E-05 | 1.34E-04 | 1.48E-03 | [9](https://toppgene.cchmc.org/showQueryTerms.jsp?userdata_id=e0bdc6b9-955d-48cd-95d3-4ccb1cbe49c7&feature=pt&row=91) |
| CCTZ | M39388 | WP GLUTATHIONE METABOLISM | 2.15E-05 | 8.17E-04 | 7.16E-03 | 7.76E-02 | [6](https://toppgene.cchmc.org/showQueryTerms.jsp?userdata_id=677e6385-c8f1-4db4-acb1-4b26e7d30bc3&feature=pt&row=92) | LMAN2 | M594 | REACTOME SIGNALING BY NOTCH4 | 6.00E-07 | 1.60E-05 | 1.34E-04 | 1.48E-03 | [11](https://toppgene.cchmc.org/showQueryTerms.jsp?userdata_id=e0bdc6b9-955d-48cd-95d3-4ccb1cbe49c7&feature=pt&row=92) |
| CDC42BPA | M8719 | BIOCARTA MCALPAIN PATHWAY | 2.15E-05 | 8.17E-04 | 7.16E-03 | 7.76E-02 | [6](https://toppgene.cchmc.org/showQueryTerms.jsp?userdata_id=677e6385-c8f1-4db4-acb1-4b26e7d30bc3&feature=pt&row=93) | M6PR | M47700 | KEGG MEDICUS REFERENCE 26S PROTEASOME MEDIATED PROTEIN DEGRADATION | 7.66E-07 | 2.00E-05 | 1.68E-04 | 1.89E-03 | [8](https://toppgene.cchmc.org/showQueryTerms.jsp?userdata_id=e0bdc6b9-955d-48cd-95d3-4ccb1cbe49c7&feature=pt&row=93) |
| CDHR2 | M15303 | REACTOME APOPTOSIS | 2.16E-05 | 8.17E-04 | 7.16E-03 | 7.76E-02 | [18](https://toppgene.cchmc.org/showQueryTerms.jsp?userdata_id=677e6385-c8f1-4db4-acb1-4b26e7d30bc3&feature=pt&row=94) | MAN1A1 | M7098 | KEGG ECM RECEPTOR INTERACTION | 7.69E-07 | 2.00E-05 | 1.68E-04 | 1.90E-03 | [11](https://toppgene.cchmc.org/showQueryTerms.jsp?userdata_id=e0bdc6b9-955d-48cd-95d3-4ccb1cbe49c7&feature=pt&row=94) |
| CDK1 | MM15906 | WP PRIMARY FOCAL SEGMENTAL GLOMERULOSCLEROSIS FSGS | 2.18E-05 | 8.17E-04 | 7.17E-03 | 7.85E-02 | [11](https://toppgene.cchmc.org/showQueryTerms.jsp?userdata_id=677e6385-c8f1-4db4-acb1-4b26e7d30bc3&feature=pt&row=95) | MARCKS | MM14982 | REACTOME REGULATION OF INSULIN LIKE GROWTH FACTOR IGF TRANSPORT AND UPTAKE BY INSULIN LIKE GROWTH FACTOR BINDING PROTEINS IGFBPS | 7.82E-07 | 2.02E-05 | 1.69E-04 | 1.93E-03 | [13](https://toppgene.cchmc.org/showQueryTerms.jsp?userdata_id=e0bdc6b9-955d-48cd-95d3-4ccb1cbe49c7&feature=pt&row=95) |
| CDK5RAP2 | MM14958 | REACTOME INTEGRIN SIGNALING | 2.39E-05 | 8.69E-04 | 7.62E-03 | 8.60E-02 | [7](https://toppgene.cchmc.org/showQueryTerms.jsp?userdata_id=677e6385-c8f1-4db4-acb1-4b26e7d30bc3&feature=pt&row=96) | MVP | MM15145 | REACTOME MITOTIC G2 G2 M PHASES | 8.23E-07 | 2.08E-05 | 1.75E-04 | 2.04E-03 | [16](https://toppgene.cchmc.org/showQueryTerms.jsp?userdata_id=e0bdc6b9-955d-48cd-95d3-4ccb1cbe49c7&feature=pt&row=96) |
| CEACAM5 | M26982 | REACTOME BUDDING AND MATURATION OF HIV VIRION | 2.39E-05 | 8.69E-04 | 7.62E-03 | 8.60E-02 | [7](https://toppgene.cchmc.org/showQueryTerms.jsp?userdata_id=677e6385-c8f1-4db4-acb1-4b26e7d30bc3&feature=pt&row=97) | NAPA | M27539 | REACTOME REGULATION OF RAS BY GAPS | 8.25E-07 | 2.08E-05 | 1.75E-04 | 2.04E-03 | [10](https://toppgene.cchmc.org/showQueryTerms.jsp?userdata_id=e0bdc6b9-955d-48cd-95d3-4ccb1cbe49c7&feature=pt&row=97) |
| CEP250 | M39572 | WP NANOPARTICLE MEDIATED ACTIVATION OF RECEPTOR SIGNALING | 2.39E-05 | 8.69E-04 | 7.62E-03 | 8.60E-02 | [7](https://toppgene.cchmc.org/showQueryTerms.jsp?userdata_id=677e6385-c8f1-4db4-acb1-4b26e7d30bc3&feature=pt&row=98) | NARS1 | M26982 | REACTOME BUDDING AND MATURATION OF HIV VIRION | 9.11E-07 | 2.28E-05 | 1.91E-04 | 2.25E-03 | [7](https://toppgene.cchmc.org/showQueryTerms.jsp?userdata_id=e0bdc6b9-955d-48cd-95d3-4ccb1cbe49c7&feature=pt&row=98) |
| CETP | M27656 | REACTOME COOPERATION OF PDCL PHLP1 AND TRIC CCT IN G PROTEIN BETA FOLDING | 2.42E-05 | 8.71E-04 | 7.64E-03 | 8.71E-02 | [8](https://toppgene.cchmc.org/showQueryTerms.jsp?userdata_id=677e6385-c8f1-4db4-acb1-4b26e7d30bc3&feature=pt&row=99) | NIBAN2 | M47748 | KEGG MEDICUS VARIANT MUTATION INACTIVATED UBQLN2 TO 26S PROTEASOME MEDIATED PROTEIN DEGRADATION | 9.41E-07 | 2.33E-05 | 1.95E-04 | 2.33E-03 | [8](https://toppgene.cchmc.org/showQueryTerms.jsp?userdata_id=e0bdc6b9-955d-48cd-95d3-4ccb1cbe49c7&feature=pt&row=99) |
| CFAP20 | M39577 | WP INTEGRIN MEDIATED CELL ADHESION | 2.54E-05 | 9.07E-04 | 7.95E-03 | 9.16E-02 | [13](https://toppgene.cchmc.org/showQueryTerms.jsp?userdata_id=677e6385-c8f1-4db4-acb1-4b26e7d30bc3&feature=pt&row=100) | NT5E | M27931 | REACTOME NEGATIVE REGULATION OF NOTCH4 SIGNALING | 9.93E-07 | 2.43E-05 | 2.04E-04 | 2.46E-03 | [9](https://toppgene.cchmc.org/showQueryTerms.jsp?userdata_id=e0bdc6b9-955d-48cd-95d3-4ccb1cbe49c7&feature=pt&row=100) |
| CFH | MM15003 | REACTOME CHAPERONIN MEDIATED PROTEIN FOLDING | 2.96E-05 | 1.03E-03 | 9.00E-03 | 1.07E-01 | [8](https://toppgene.cchmc.org/showQueryTerms.jsp?userdata_id=677e6385-c8f1-4db4-acb1-4b26e7d30bc3&feature=pt&row=101) | NUTF2 | M27436 | REACTOME PROGRAMMED CELL DEATH | 1.01E-06 | 2.44E-05 | 2.05E-04 | 2.49E-03 | [17](https://toppgene.cchmc.org/showQueryTerms.jsp?userdata_id=e0bdc6b9-955d-48cd-95d3-4ccb1cbe49c7&feature=pt&row=101) |
| CFI | MM15587 | REACTOME SIGNALING BY RECEPTOR TYROSINE KINASES | 2.96E-05 | 1.03E-03 | 9.00E-03 | 1.07E-01 | [30](https://toppgene.cchmc.org/showQueryTerms.jsp?userdata_id=677e6385-c8f1-4db4-acb1-4b26e7d30bc3&feature=pt&row=102) | OTUB1 | M27778 | REACTOME MET PROMOTES CELL MOTILITY | 1.15E-06 | 2.75E-05 | 2.31E-04 | 2.84E-03 | [8](https://toppgene.cchmc.org/showQueryTerms.jsp?userdata_id=e0bdc6b9-955d-48cd-95d3-4ccb1cbe49c7&feature=pt&row=102) |
| CHMP1A | M27440 | REACTOME SIGNALING BY HEDGEHOG | 2.98E-05 | 1.03E-03 | 9.00E-03 | 1.07E-01 | [16](https://toppgene.cchmc.org/showQueryTerms.jsp?userdata_id=677e6385-c8f1-4db4-acb1-4b26e7d30bc3&feature=pt&row=103) | PACSIN2 | MM15137 | REACTOME AUF1 HNRNP D0 BINDS AND DESTABILIZES MRNA | 1.17E-06 | 2.75E-05 | 2.31E-04 | 2.89E-03 | [9](https://toppgene.cchmc.org/showQueryTerms.jsp?userdata_id=e0bdc6b9-955d-48cd-95d3-4ccb1cbe49c7&feature=pt&row=103) |
| CHMP2A | MM1476 | BIOCARTA RHO PATHWAY | 2.99E-05 | 1.03E-03 | 9.00E-03 | 1.08E-01 | [6](https://toppgene.cchmc.org/showQueryTerms.jsp?userdata_id=677e6385-c8f1-4db4-acb1-4b26e7d30bc3&feature=pt&row=104) | PDCD6IP | M998 | REACTOME AUF1 HNRNP D0 BINDS AND DESTABILIZES MRNA | 1.17E-06 | 2.75E-05 | 2.31E-04 | 2.89E-03 | [9](https://toppgene.cchmc.org/showQueryTerms.jsp?userdata_id=e0bdc6b9-955d-48cd-95d3-4ccb1cbe49c7&feature=pt&row=104) |
| CHMP2B | M48033 | REACTOME VIRAL INFECTION PATHWAYS | 3.02E-05 | 1.03E-03 | 9.00E-03 | 1.09E-01 | [49](https://toppgene.cchmc.org/showQueryTerms.jsp?userdata_id=677e6385-c8f1-4db4-acb1-4b26e7d30bc3&feature=pt&row=105) | PEDF | M169 | PID INTEGRIN2 PATHWAY | 1.18E-06 | 2.76E-05 | 2.31E-04 | 2.92E-03 | [7](https://toppgene.cchmc.org/showQueryTerms.jsp?userdata_id=e0bdc6b9-955d-48cd-95d3-4ccb1cbe49c7&feature=pt&row=105) |
| CHMP3 | MM15913 | WP FOCAL ADHESION | 3.88E-05 | 1.29E-03 | 1.13E-02 | 1.40E-01 | [18](https://toppgene.cchmc.org/showQueryTerms.jsp?userdata_id=677e6385-c8f1-4db4-acb1-4b26e7d30bc3&feature=pt&row=106) | PFKL | MM1430 | BIOCARTA LECTIN PATHWAY | 1.23E-06 | 2.83E-05 | 2.38E-04 | 3.03E-03 | [5](https://toppgene.cchmc.org/showQueryTerms.jsp?userdata_id=e0bdc6b9-955d-48cd-95d3-4ccb1cbe49c7&feature=pt&row=106) |
| CHMP4B | M27216 | REACTOME LAMININ INTERACTIONS | 3.88E-05 | 1.29E-03 | 1.13E-02 | 1.40E-01 | [7](https://toppgene.cchmc.org/showQueryTerms.jsp?userdata_id=677e6385-c8f1-4db4-acb1-4b26e7d30bc3&feature=pt&row=107) | PGM1 | MM14867 | REACTOME INTEGRIN CELL SURFACE INTERACTIONS | 1.62E-06 | 3.70E-05 | 3.10E-04 | 4.00E-03 | [10](https://toppgene.cchmc.org/showQueryTerms.jsp?userdata_id=e0bdc6b9-955d-48cd-95d3-4ccb1cbe49c7&feature=pt&row=107) |
| CKAP4 | M1001 | BIOCARTA RHO PATHWAY | 4.08E-05 | 1.35E-03 | 1.18E-02 | 1.47E-01 | [6](https://toppgene.cchmc.org/showQueryTerms.jsp?userdata_id=677e6385-c8f1-4db4-acb1-4b26e7d30bc3&feature=pt&row=108) | PHGDH | MM15637 | REACTOME ENDOSOMAL SORTING COMPLEX REQUIRED FOR TRANSPORT ESCRT | 1.92E-06 | 4.33E-05 | 3.63E-04 | 4.76E-03 | [7](https://toppgene.cchmc.org/showQueryTerms.jsp?userdata_id=e0bdc6b9-955d-48cd-95d3-4ccb1cbe49c7&feature=pt&row=108) |
| CLIC6 | M27322 | REACTOME PCP CE PATHWAY | 4.11E-05 | 1.35E-03 | 1.18E-02 | 1.48E-01 | [12](https://toppgene.cchmc.org/showQueryTerms.jsp?userdata_id=677e6385-c8f1-4db4-acb1-4b26e7d30bc3&feature=pt&row=109) | PLS2 | M1009 | REACTOME ENDOSOMAL SORTING COMPLEX REQUIRED FOR TRANSPORT ESCRT | 1.92E-06 | 4.33E-05 | 3.63E-04 | 4.76E-03 | [7](https://toppgene.cchmc.org/showQueryTerms.jsp?userdata_id=e0bdc6b9-955d-48cd-95d3-4ccb1cbe49c7&feature=pt&row=109) |
| CLTC | MM15182 | REACTOME SIGNALING BY HEDGEHOG | 4.22E-05 | 1.37E-03 | 1.20E-02 | 1.52E-01 | [15](https://toppgene.cchmc.org/showQueryTerms.jsp?userdata_id=677e6385-c8f1-4db4-acb1-4b26e7d30bc3&feature=pt&row=110) | PPIB | M27812 | REACTOME COLLAGEN CHAIN TRIMERIZATION | 2.03E-06 | 4.51E-05 | 3.79E-04 | 5.01E-03 | [8](https://toppgene.cchmc.org/showQueryTerms.jsp?userdata_id=e0bdc6b9-955d-48cd-95d3-4ccb1cbe49c7&feature=pt&row=110) |
| CNKSR2 | M38994 | REACTOME SIGNALING BY BRAF AND RAF1 FUSIONS | 4.28E-05 | 1.38E-03 | 1.21E-02 | 1.54E-01 | [10](https://toppgene.cchmc.org/showQueryTerms.jsp?userdata_id=677e6385-c8f1-4db4-acb1-4b26e7d30bc3&feature=pt&row=111) | PPIL1 | MM1368 | BIOCARTA CLASSIC PATHWAY | 2.07E-06 | 4.52E-05 | 3.79E-04 | 5.11E-03 | [5](https://toppgene.cchmc.org/showQueryTerms.jsp?userdata_id=e0bdc6b9-955d-48cd-95d3-4ccb1cbe49c7&feature=pt&row=111) |
| CNTLN | M27870 | REACTOME SIGNALING BY RECEPTOR TYROSINE KINASES | 4.41E-05 | 1.41E-03 | 1.23E-02 | 1.59E-01 | [35](https://toppgene.cchmc.org/showQueryTerms.jsp?userdata_id=677e6385-c8f1-4db4-acb1-4b26e7d30bc3&feature=pt&row=112) | PRDX6 | M39408 | WP RALA DOWNSTREAM REGULATED GENES | 2.07E-06 | 4.52E-05 | 3.79E-04 | 5.11E-03 | [5](https://toppgene.cchmc.org/showQueryTerms.jsp?userdata_id=e0bdc6b9-955d-48cd-95d3-4ccb1cbe49c7&feature=pt&row=112) |
| COASY | MM14988 | REACTOME BETA CATENIN INDEPENDENT WNT SIGNALING | 4.54E-05 | 1.43E-03 | 1.26E-02 | 1.63E-01 | [14](https://toppgene.cchmc.org/showQueryTerms.jsp?userdata_id=677e6385-c8f1-4db4-acb1-4b26e7d30bc3&feature=pt&row=113) | PSMA1 | M7253 | KEGG FOCAL ADHESION | 2.17E-06 | 4.70E-05 | 3.94E-04 | 5.36E-03 | [16](https://toppgene.cchmc.org/showQueryTerms.jsp?userdata_id=e0bdc6b9-955d-48cd-95d3-4ccb1cbe49c7&feature=pt&row=113) |
| COBLL1 | M991 | REACTOME TRANSFERRIN ENDOCYTOSIS AND RECYCLING | 4.87E-05 | 1.51E-03 | 1.33E-02 | 1.76E-01 | [7](https://toppgene.cchmc.org/showQueryTerms.jsp?userdata_id=677e6385-c8f1-4db4-acb1-4b26e7d30bc3&feature=pt&row=114) | PSMA5 | M864 | REACTOME MITOTIC G2 G2 M PHASES | 2.31E-06 | 4.97E-05 | 4.17E-04 | 5.72E-03 | [16](https://toppgene.cchmc.org/showQueryTerms.jsp?userdata_id=e0bdc6b9-955d-48cd-95d3-4ccb1cbe49c7&feature=pt&row=114) |
| COL6A1 | MM15639 | REACTOME TRANSFERRIN ENDOCYTOSIS AND RECYCLING | 4.87E-05 | 1.51E-03 | 1.33E-02 | 1.76E-01 | [7](https://toppgene.cchmc.org/showQueryTerms.jsp?userdata_id=677e6385-c8f1-4db4-acb1-4b26e7d30bc3&feature=pt&row=115) | PSMB5 | MM14526 | REACTOME ANTIGEN PROCESSING CROSS PRESENTATION | 2.36E-06 | 5.02E-05 | 4.21E-04 | 5.83E-03 | [10](https://toppgene.cchmc.org/showQueryTerms.jsp?userdata_id=e0bdc6b9-955d-48cd-95d3-4ccb1cbe49c7&feature=pt&row=115) |
| COL6A3 | M39740 | WP METABOLIC REPROGRAMMING IN COLON CANCER | 5.21E-05 | 1.60E-03 | 1.41E-02 | 1.88E-01 | [8](https://toppgene.cchmc.org/showQueryTerms.jsp?userdata_id=677e6385-c8f1-4db4-acb1-4b26e7d30bc3&feature=pt&row=116) | PSMC6 | M47721 | KEGG MEDICUS REFERENCE EGF EGFR ACTIN SIGNALING PATHWAY | 2.40E-06 | 5.07E-05 | 4.26E-04 | 5.93E-03 | [6](https://toppgene.cchmc.org/showQueryTerms.jsp?userdata_id=e0bdc6b9-955d-48cd-95d3-4ccb1cbe49c7&feature=pt&row=116) |
| COTL1 | M27080 | REACTOME RHO GTPASE EFFECTORS | 5.55E-05 | 1.69E-03 | 1.49E-02 | 2.00E-01 | [25](https://toppgene.cchmc.org/showQueryTerms.jsp?userdata_id=677e6385-c8f1-4db4-acb1-4b26e7d30bc3&feature=pt&row=117) | RAB13 | MM15517 | REACTOME MET PROMOTES CELL MOTILITY | 2.42E-06 | 5.07E-05 | 4.26E-04 | 5.99E-03 | [7](https://toppgene.cchmc.org/showQueryTerms.jsp?userdata_id=e0bdc6b9-955d-48cd-95d3-4ccb1cbe49c7&feature=pt&row=117) |
| CP | M29848 | REACTOME SEALING OF THE NUCLEAR ENVELOPE NE BY ESCRT III | 6.07E-05 | 1.82E-03 | 1.60E-02 | 2.18E-01 | [7](https://toppgene.cchmc.org/showQueryTerms.jsp?userdata_id=677e6385-c8f1-4db4-acb1-4b26e7d30bc3&feature=pt&row=118) | RAB14 | M48033 | REACTOME VIRAL INFECTION PATHWAYS | 2.50E-06 | 5.20E-05 | 4.36E-04 | 6.19E-03 | [37](https://toppgene.cchmc.org/showQueryTerms.jsp?userdata_id=e0bdc6b9-955d-48cd-95d3-4ccb1cbe49c7&feature=pt&row=118) |
| CPNE3 | M165 | PID SYNDECAN 4 PATHWAY | 6.07E-05 | 1.82E-03 | 1.60E-02 | 2.18E-01 | [7](https://toppgene.cchmc.org/showQueryTerms.jsp?userdata_id=677e6385-c8f1-4db4-acb1-4b26e7d30bc3&feature=pt&row=119) | RAB35 | M19752 | REACTOME COMPLEMENT CASCADE | 2.85E-06 | 5.84E-05 | 4.90E-04 | 7.05E-03 | [12](https://toppgene.cchmc.org/showQueryTerms.jsp?userdata_id=e0bdc6b9-955d-48cd-95d3-4ccb1cbe49c7&feature=pt&row=119) |
| CPVL | M42521 | REACTOME SIGNALING BY ALK IN CANCER | 6.27E-05 | 1.87E-03 | 1.64E-02 | 2.26E-01 | [9](https://toppgene.cchmc.org/showQueryTerms.jsp?userdata_id=677e6385-c8f1-4db4-acb1-4b26e7d30bc3&feature=pt&row=120) | RAB5B | M27563 | REACTOME DEFECTIVE CFTR CAUSES CYSTIC FIBROSIS | 2.87E-06 | 5.84E-05 | 4.90E-04 | 7.10E-03 | [9](https://toppgene.cchmc.org/showQueryTerms.jsp?userdata_id=e0bdc6b9-955d-48cd-95d3-4ccb1cbe49c7&feature=pt&row=120) |
| CREB5 | M27626 | REACTOME ONCOGENIC MAPK SIGNALING | 6.60E-05 | 1.95E-03 | 1.71E-02 | 2.38E-01 | [11](https://toppgene.cchmc.org/showQueryTerms.jsp?userdata_id=677e6385-c8f1-4db4-acb1-4b26e7d30bc3&feature=pt&row=121) | RAB5C | M10680 | KEGG PROTEASOME | 2.88E-06 | 5.84E-05 | 4.90E-04 | 7.13E-03 | [8](https://toppgene.cchmc.org/showQueryTerms.jsp?userdata_id=e0bdc6b9-955d-48cd-95d3-4ccb1cbe49c7&feature=pt&row=121) |
| CREBBP | M12469 | REACTOME HIV INFECTION | 6.67E-05 | 1.95E-03 | 1.71E-02 | 2.40E-01 | [20](https://toppgene.cchmc.org/showQueryTerms.jsp?userdata_id=677e6385-c8f1-4db4-acb1-4b26e7d30bc3&feature=pt&row=122) | RAB8A | M17879 | REACTOME COMMON PATHWAY OF FIBRIN CLOT FORMATION | 3.24E-06 | 6.38E-05 | 5.35E-04 | 8.02E-03 | [6](https://toppgene.cchmc.org/showQueryTerms.jsp?userdata_id=e0bdc6b9-955d-48cd-95d3-4ccb1cbe49c7&feature=pt&row=122) |
| CRNN | M237 | PID VEGFR1 2 PATHWAY | 7.24E-05 | 2.09E-03 | 1.83E-02 | 2.61E-01 | [10](https://toppgene.cchmc.org/showQueryTerms.jsp?userdata_id=677e6385-c8f1-4db4-acb1-4b26e7d30bc3&feature=pt&row=123) | RALA | MM14558 | REACTOME COMMON PATHWAY OF FIBRIN CLOT FORMATION | 3.24E-06 | 6.38E-05 | 5.35E-04 | 8.02E-03 | [6](https://toppgene.cchmc.org/showQueryTerms.jsp?userdata_id=e0bdc6b9-955d-48cd-95d3-4ccb1cbe49c7&feature=pt&row=123) |
| CROCC | M1877 | REACTOME GOLGI ASSOCIATED VESICLE BIOGENESIS | 7.26E-05 | 2.09E-03 | 1.83E-02 | 2.62E-01 | [9](https://toppgene.cchmc.org/showQueryTerms.jsp?userdata_id=677e6385-c8f1-4db4-acb1-4b26e7d30bc3&feature=pt&row=124) | RAP1B | M11521 | KEGG GLYCOLYSIS GLUCONEOGENESIS | 3.30E-06 | 6.38E-05 | 5.35E-04 | 8.16E-03 | [9](https://toppgene.cchmc.org/showQueryTerms.jsp?userdata_id=e0bdc6b9-955d-48cd-95d3-4ccb1cbe49c7&feature=pt&row=124) |
| CSK | M240 | PID SYNDECAN 2 PATHWAY | 7.49E-05 | 2.12E-03 | 1.86E-02 | 2.70E-01 | [7](https://toppgene.cchmc.org/showQueryTerms.jsp?userdata_id=677e6385-c8f1-4db4-acb1-4b26e7d30bc3&feature=pt&row=125) | RAP2B | MM15181 | REACTOME HEDGEHOG LIGAND BIOGENESIS | 3.30E-06 | 6.38E-05 | 5.35E-04 | 8.16E-03 | [9](https://toppgene.cchmc.org/showQueryTerms.jsp?userdata_id=e0bdc6b9-955d-48cd-95d3-4ccb1cbe49c7&feature=pt&row=125) |
| CSRP1 | M27297 | REACTOME COOPERATION OF PREFOLDIN AND TRIC CCT IN ACTIN AND TUBULIN FOLDING | 7.49E-05 | 2.12E-03 | 1.86E-02 | 2.70E-01 | [7](https://toppgene.cchmc.org/showQueryTerms.jsp?userdata_id=677e6385-c8f1-4db4-acb1-4b26e7d30bc3&feature=pt&row=126) | RAPGEFL1 | MM1395 | BIOCARTA EXTRINSIC PATHWAY | 3.30E-06 | 6.38E-05 | 5.35E-04 | 8.17E-03 | [5](https://toppgene.cchmc.org/showQueryTerms.jsp?userdata_id=e0bdc6b9-955d-48cd-95d3-4ccb1cbe49c7&feature=pt&row=126) |
| CSTB | M27471 | REACTOME HEDGEHOG OFF STATE | 7.55E-05 | 2.13E-03 | 1.86E-02 | 2.72E-01 | [13](https://toppgene.cchmc.org/showQueryTerms.jsp?userdata_id=677e6385-c8f1-4db4-acb1-4b26e7d30bc3&feature=pt&row=127) | RDX | M4470 | BIOCARTA EXTRINSIC PATHWAY | 3.30E-06 | 6.38E-05 | 5.35E-04 | 8.17E-03 | [5](https://toppgene.cchmc.org/showQueryTerms.jsp?userdata_id=e0bdc6b9-955d-48cd-95d3-4ccb1cbe49c7&feature=pt&row=127) |
| CTNS | M1262 | REACTOME GRB2 SOS PROVIDES LINKAGE TO MAPK SIGNALING FOR INTEGRINS | 8.12E-05 | 2.23E-03 | 1.96E-02 | 2.92E-01 | [5](https://toppgene.cchmc.org/showQueryTerms.jsp?userdata_id=677e6385-c8f1-4db4-acb1-4b26e7d30bc3&feature=pt&row=128) | RENBP | M27743 | REACTOME THE ROLE OF GTSE1 IN G2 M PROGRESSION AFTER G2 CHECKPOINT | 3.38E-06 | 6.42E-05 | 5.39E-04 | 8.35E-03 | [10](https://toppgene.cchmc.org/showQueryTerms.jsp?userdata_id=e0bdc6b9-955d-48cd-95d3-4ccb1cbe49c7&feature=pt&row=128) |
| CTSA | M27060 | REACTOME HEME DEGRADATION | 8.12E-05 | 2.23E-03 | 1.96E-02 | 2.92E-01 | [5](https://toppgene.cchmc.org/showQueryTerms.jsp?userdata_id=677e6385-c8f1-4db4-acb1-4b26e7d30bc3&feature=pt&row=129) | RPS11 | M27643 | REACTOME SIGNALING BY MET | 3.38E-06 | 6.42E-05 | 5.39E-04 | 8.35E-03 | [10](https://toppgene.cchmc.org/showQueryTerms.jsp?userdata_id=e0bdc6b9-955d-48cd-95d3-4ccb1cbe49c7&feature=pt&row=129) |
| CTSC | MM14959 | REACTOME GRB2 SOS PROVIDES LINKAGE TO MAPK SIGNALING FOR INTEGRINS | 8.12E-05 | 2.23E-03 | 1.96E-02 | 2.92E-01 | [5](https://toppgene.cchmc.org/showQueryTerms.jsp?userdata_id=677e6385-c8f1-4db4-acb1-4b26e7d30bc3&feature=pt&row=130) | RPS18 | M12469 | REACTOME HIV INFECTION | 3.50E-06 | 6.60E-05 | 5.54E-04 | 8.65E-03 | [17](https://toppgene.cchmc.org/showQueryTerms.jsp?userdata_id=e0bdc6b9-955d-48cd-95d3-4ccb1cbe49c7&feature=pt&row=130) |
| CUBN | M39812 | WP PROXIMAL TUBULE TRANSPORT | 8.38E-05 | 2.29E-03 | 2.00E-02 | 3.02E-01 | [9](https://toppgene.cchmc.org/showQueryTerms.jsp?userdata_id=677e6385-c8f1-4db4-acb1-4b26e7d30bc3&feature=pt&row=131) | S100A10 | MM14528 | REACTOME CROSS PRESENTATION OF SOLUBLE EXOGENOUS ANTIGENS ENDOSOMES | 4.03E-06 | 7.54E-05 | 6.33E-04 | 9.96E-03 | [8](https://toppgene.cchmc.org/showQueryTerms.jsp?userdata_id=e0bdc6b9-955d-48cd-95d3-4ccb1cbe49c7&feature=pt&row=131) |
| CUL3 | MM14898 | REACTOME MITOTIC METAPHASE AND ANAPHASE | 8.47E-05 | 2.29E-03 | 2.01E-02 | 3.05E-01 | [20](https://toppgene.cchmc.org/showQueryTerms.jsp?userdata_id=677e6385-c8f1-4db4-acb1-4b26e7d30bc3&feature=pt&row=132) | S100A16 | M595 | REACTOME DOWNSTREAM SIGNALING EVENTS OF B CELL RECEPTOR BCR | 4.25E-06 | 7.90E-05 | 6.63E-04 | 1.05E-02 | [10](https://toppgene.cchmc.org/showQueryTerms.jsp?userdata_id=e0bdc6b9-955d-48cd-95d3-4ccb1cbe49c7&feature=pt&row=132) |
| CUL4B | M39474 | WP GLYCOLYSIS AND GLUCONEOGENESIS | 8.74E-05 | 2.35E-03 | 2.06E-02 | 3.15E-01 | [8](https://toppgene.cchmc.org/showQueryTerms.jsp?userdata_id=677e6385-c8f1-4db4-acb1-4b26e7d30bc3&feature=pt&row=133) | S100A4 | M13036 | KEGG PRION DISEASES | 4.60E-06 | 8.48E-05 | 7.12E-04 | 1.14E-02 | [7](https://toppgene.cchmc.org/showQueryTerms.jsp?userdata_id=e0bdc6b9-955d-48cd-95d3-4ccb1cbe49c7&feature=pt&row=133) |
| CUTL2 | MM15688 | REACTOME CELLULAR RESPONSE TO CHEMICAL STRESS | 8.82E-05 | 2.35E-03 | 2.06E-02 | 3.17E-01 | [16](https://toppgene.cchmc.org/showQueryTerms.jsp?userdata_id=677e6385-c8f1-4db4-acb1-4b26e7d30bc3&feature=pt&row=134) | SDCBP | M27439 | REACTOME HEDGEHOG LIGAND BIOGENESIS | 4.94E-06 | 9.05E-05 | 7.59E-04 | 1.22E-02 | [9](https://toppgene.cchmc.org/showQueryTerms.jsp?userdata_id=e0bdc6b9-955d-48cd-95d3-4ccb1cbe49c7&feature=pt&row=134) |
| CYBRD1 | M27185 | REACTOME MITOTIC METAPHASE AND ANAPHASE | 8.98E-05 | 2.38E-03 | 2.08E-02 | 3.23E-01 | [20](https://toppgene.cchmc.org/showQueryTerms.jsp?userdata_id=677e6385-c8f1-4db4-acb1-4b26e7d30bc3&feature=pt&row=135) | ST13 | M47872 | KEGG MEDICUS REFERENCE COMMON PATHWAY OF COMPLEMENT CASCADE MAC FORMATION | 5.01E-06 | 9.10E-05 | 7.64E-04 | 1.24E-02 | [4](https://toppgene.cchmc.org/showQueryTerms.jsp?userdata_id=e0bdc6b9-955d-48cd-95d3-4ccb1cbe49c7&feature=pt&row=135) |
| CYFIP2 | M5283 | REACTOME HOST INTERACTIONS OF HIV FACTORS | 9.09E-05 | 2.39E-03 | 2.10E-02 | 3.27E-01 | [14](https://toppgene.cchmc.org/showQueryTerms.jsp?userdata_id=677e6385-c8f1-4db4-acb1-4b26e7d30bc3&feature=pt&row=136) | STOML1 | MM15830 | WP INTEGRIN MEDIATED CELL ADHESION | 5.38E-06 | 9.65E-05 | 8.10E-04 | 1.33E-02 | [11](https://toppgene.cchmc.org/showQueryTerms.jsp?userdata_id=e0bdc6b9-955d-48cd-95d3-4ccb1cbe49c7&feature=pt&row=136) |
| DBNL | M39375 | WP CLEAR CELL RENAL CELL CARCINOMA PATHWAYS | 9.21E-05 | 2.39E-03 | 2.10E-02 | 3.32E-01 | [11](https://toppgene.cchmc.org/showQueryTerms.jsp?userdata_id=677e6385-c8f1-4db4-acb1-4b26e7d30bc3&feature=pt&row=137) | TAGLN2 | M39577 | WP INTEGRIN MEDIATED CELL ADHESION | 5.38E-06 | 9.65E-05 | 8.10E-04 | 1.33E-02 | [11](https://toppgene.cchmc.org/showQueryTerms.jsp?userdata_id=e0bdc6b9-955d-48cd-95d3-4ccb1cbe49c7&feature=pt&row=137) |
| DCCIP13 | MM14787 | REACTOME TRANS GOLGI NETWORK VESICLE BUDDING | 9.28E-05 | 2.39E-03 | 2.10E-02 | 3.34E-01 | [10](https://toppgene.cchmc.org/showQueryTerms.jsp?userdata_id=677e6385-c8f1-4db4-acb1-4b26e7d30bc3&feature=pt&row=138) | TMEM59 | MM17068 | REACTOME REGULATION OF RUNX2 EXPRESSION AND ACTIVITY | 5.54E-06 | 9.71E-05 | 8.15E-04 | 1.37E-02 | [8](https://toppgene.cchmc.org/showQueryTerms.jsp?userdata_id=e0bdc6b9-955d-48cd-95d3-4ccb1cbe49c7&feature=pt&row=138) |
| DCD | M15926 | BIOCARTA TFF PATHWAY | 9.30E-05 | 2.39E-03 | 2.10E-02 | 3.35E-01 | [6](https://toppgene.cchmc.org/showQueryTerms.jsp?userdata_id=677e6385-c8f1-4db4-acb1-4b26e7d30bc3&feature=pt&row=139) | TMX1 | M510 | REACTOME CROSS PRESENTATION OF SOLUBLE EXOGENOUS ANTIGENS ENDOSOMES | 5.54E-06 | 9.71E-05 | 8.15E-04 | 1.37E-02 | [8](https://toppgene.cchmc.org/showQueryTerms.jsp?userdata_id=e0bdc6b9-955d-48cd-95d3-4ccb1cbe49c7&feature=pt&row=139) |
| DDAH1 | MM15278 | REACTOME MAPK FAMILY SIGNALING CASCADES | 9.48E-05 | 2.42E-03 | 2.12E-02 | 3.42E-01 | [24](https://toppgene.cchmc.org/showQueryTerms.jsp?userdata_id=677e6385-c8f1-4db4-acb1-4b26e7d30bc3&feature=pt&row=140) | TPI1 | MM15385 | REACTOME UBIQUITIN MEDIATED DEGRADATION OF PHOSPHORYLATED CDC25A | 5.54E-06 | 9.71E-05 | 8.15E-04 | 1.37E-02 | [8](https://toppgene.cchmc.org/showQueryTerms.jsp?userdata_id=e0bdc6b9-955d-48cd-95d3-4ccb1cbe49c7&feature=pt&row=140) |
| DDB1 | M27050 | REACTOME ORGANELLE BIOGENESIS AND MAINTENANCE | 9.82E-05 | 2.49E-03 | 2.18E-02 | 3.54E-01 | [23](https://toppgene.cchmc.org/showQueryTerms.jsp?userdata_id=677e6385-c8f1-4db4-acb1-4b26e7d30bc3&feature=pt&row=141) | TUBA1A | MM15261 | REACTOME REGULATION OF RAS BY GAPS | 5.62E-06 | 9.79E-05 | 8.21E-04 | 1.39E-02 | [9](https://toppgene.cchmc.org/showQueryTerms.jsp?userdata_id=e0bdc6b9-955d-48cd-95d3-4ccb1cbe49c7&feature=pt&row=141) |
| DDC | M27472 | REACTOME CILIUM ASSEMBLY | 9.96E-05 | 2.51E-03 | 2.20E-02 | 3.59E-01 | [18](https://toppgene.cchmc.org/showQueryTerms.jsp?userdata_id=677e6385-c8f1-4db4-acb1-4b26e7d30bc3&feature=pt&row=142) | TUBB8 | MM14988 | REACTOME BETA CATENIN INDEPENDENT WNT SIGNALING | 5.78E-06 | 9.99E-05 | 8.39E-04 | 1.43E-02 | [12](https://toppgene.cchmc.org/showQueryTerms.jsp?userdata_id=e0bdc6b9-955d-48cd-95d3-4ccb1cbe49c7&feature=pt&row=142) |
| DDX19B | MM14755 | REACTOME RHO GTPASE EFFECTORS | 1.02E-04 | 2.54E-03 | 2.23E-02 | 3.66E-01 | [21](https://toppgene.cchmc.org/showQueryTerms.jsp?userdata_id=677e6385-c8f1-4db4-acb1-4b26e7d30bc3&feature=pt&row=143) | VAMP3 | MM15383 | REACTOME G2 M CHECKPOINTS | 6.04E-06 | 1.04E-04 | 8.70E-04 | 1.49E-02 | [13](https://toppgene.cchmc.org/showQueryTerms.jsp?userdata_id=e0bdc6b9-955d-48cd-95d3-4ccb1cbe49c7&feature=pt&row=143) |
| DDX3X | MM15815 | WP GLUTATHIONE AND ONE CARBON METABOLISM | 1.11E-04 | 2.75E-03 | 2.41E-02 | 4.01E-01 | [7](https://toppgene.cchmc.org/showQueryTerms.jsp?userdata_id=677e6385-c8f1-4db4-acb1-4b26e7d30bc3&feature=pt&row=144) | VCP | MM15041 | REACTOME PCP CE PATHWAY | 6.60E-06 | 1.13E-04 | 9.45E-04 | 1.63E-02 | [10](https://toppgene.cchmc.org/showQueryTerms.jsp?userdata_id=e0bdc6b9-955d-48cd-95d3-4ccb1cbe49c7&feature=pt&row=144) |
| DDX5 | M2881 | REACTOME LYSOSOME VESICLE BIOGENESIS | 1.11E-04 | 2.75E-03 | 2.41E-02 | 4.01E-01 | [7](https://toppgene.cchmc.org/showQueryTerms.jsp?userdata_id=677e6385-c8f1-4db4-acb1-4b26e7d30bc3&feature=pt&row=145) | VPS35 | M2780 | REACTOME SIGNALING BY ROBO RECEPTORS | 7.04E-06 | 1.19E-04 | 1.00E-03 | 1.74E-02 | [16](https://toppgene.cchmc.org/showQueryTerms.jsp?userdata_id=e0bdc6b9-955d-48cd-95d3-4ccb1cbe49c7&feature=pt&row=145) |
| DED81 | M27894 | REACTOME INTERLEUKIN 12 SIGNALING | 1.20E-04 | 2.93E-03 | 2.57E-02 | 4.34E-01 | [8](https://toppgene.cchmc.org/showQueryTerms.jsp?userdata_id=677e6385-c8f1-4db4-acb1-4b26e7d30bc3&feature=pt&row=146) | VTA1 | MM15140 | REACTOME REGULATION OF MRNA STABILITY BY PROTEINS THAT BIND AU RICH ELEMENTS | 7.35E-06 | 1.24E-04 | 1.04E-03 | 1.82E-02 | [10](https://toppgene.cchmc.org/showQueryTerms.jsp?userdata_id=e0bdc6b9-955d-48cd-95d3-4ccb1cbe49c7&feature=pt&row=146) |
| DHPR | M19806 | REACTOME POST TRANSLATIONAL PROTEIN MODIFICATION | 1.21E-04 | 2.93E-03 | 2.57E-02 | 4.34E-01 | [73](https://toppgene.cchmc.org/showQueryTerms.jsp?userdata_id=677e6385-c8f1-4db4-acb1-4b26e7d30bc3&feature=pt&row=147) | WDR1 | MM15536 | REACTOME REGULATION OF RUNX3 EXPRESSION AND ACTIVITY | 7.50E-06 | 1.25E-04 | 1.05E-03 | 1.85E-02 | [8](https://toppgene.cchmc.org/showQueryTerms.jsp?userdata_id=e0bdc6b9-955d-48cd-95d3-4ccb1cbe49c7&feature=pt&row=147) |
| DNAH8 | M26923 | REACTOME ROS AND RNS PRODUCTION IN PHAGOCYTES | 1.35E-04 | 3.25E-03 | 2.85E-02 | 4.84E-01 | [7](https://toppgene.cchmc.org/showQueryTerms.jsp?userdata_id=677e6385-c8f1-4db4-acb1-4b26e7d30bc3&feature=pt&row=148) | YWHAE | M944 | REACTOME REGULATION OF MRNA STABILITY BY PROTEINS THAT BIND AU RICH ELEMENTS | 8.16E-06 | 1.35E-04 | 1.14E-03 | 2.02E-02 | [10](https://toppgene.cchmc.org/showQueryTerms.jsp?userdata_id=e0bdc6b9-955d-48cd-95d3-4ccb1cbe49c7&feature=pt&row=148) |
| DNAJA2 | M47718 | KEGG MEDICUS REFERENCE ITGA B FAK CDC42 SIGNALING PATHWAY | 1.51E-04 | 3.62E-03 | 3.17E-02 | 5.42E-01 | [6](https://toppgene.cchmc.org/showQueryTerms.jsp?userdata_id=677e6385-c8f1-4db4-acb1-4b26e7d30bc3&feature=pt&row=149) | YWHAG | MM16640 | REACTOME NUCLEAR EVENTS MEDIATED BY NFE2L2 | 8.68E-06 | 1.42E-04 | 1.19E-03 | 2.15E-02 | [8](https://toppgene.cchmc.org/showQueryTerms.jsp?userdata_id=e0bdc6b9-955d-48cd-95d3-4ccb1cbe49c7&feature=pt&row=149) |
| DNAJC13 | MM14985 | REACTOME TRANSPORT OF SMALL MOLECULES | 1.55E-04 | 3.70E-03 | 3.24E-02 | 5.59E-01 | [40](https://toppgene.cchmc.org/showQueryTerms.jsp?userdata_id=677e6385-c8f1-4db4-acb1-4b26e7d30bc3&feature=pt&row=150) | YWHAQ | MM15152 | REACTOME DEGRADATION OF AXIN | 8.68E-06 | 1.42E-04 | 1.19E-03 | 2.15E-02 | [8](https://toppgene.cchmc.org/showQueryTerms.jsp?userdata_id=e0bdc6b9-955d-48cd-95d3-4ccb1cbe49c7&feature=pt&row=150) |
| DNAJC7 | M45028 | REACTOME EARLY SARS COV 2 INFECTION EVENTS | 1.61E-04 | 3.82E-03 | 3.35E-02 | 5.81E-01 | [7](https://toppgene.cchmc.org/showQueryTerms.jsp?userdata_id=677e6385-c8f1-4db4-acb1-4b26e7d30bc3&feature=pt&row=151) | YWHAZ | M39719 | WP FOCAL ADHESION PI3K AKT MTOR SIGNALING PATHWAY | 9.23E-06 | 1.50E-04 | 1.26E-03 | 2.28E-02 | [19](https://toppgene.cchmc.org/showQueryTerms.jsp?userdata_id=e0bdc6b9-955d-48cd-95d3-4ccb1cbe49c7&feature=pt&row=151) |
| DNASE1L3 | M11521 | KEGG GLYCOLYSIS GLUCONEOGENESIS | 1.64E-04 | 3.85E-03 | 3.38E-02 | 5.90E-01 | [9](https://toppgene.cchmc.org/showQueryTerms.jsp?userdata_id=677e6385-c8f1-4db4-acb1-4b26e7d30bc3&feature=pt&row=152) | FLOT1 | MM15384 | REACTOME STABILIZATION OF P53 | 1.00E-05 | 1.62E-04 | 1.36E-03 | 2.48E-02 | [8](https://toppgene.cchmc.org/showQueryTerms.jsp?userdata_id=e0bdc6b9-955d-48cd-95d3-4ccb1cbe49c7&feature=pt&row=152) |
| DNM1L | MM15206 | REACTOME HEDGEHOG OFF STATE | 1.66E-04 | 3.87E-03 | 3.39E-02 | 5.96E-01 | [12](https://toppgene.cchmc.org/showQueryTerms.jsp?userdata_id=677e6385-c8f1-4db4-acb1-4b26e7d30bc3&feature=pt&row=153) | FTH1 | M46428 | REACTOME SARS COV 1 TARGETS HOST INTRACELLULAR SIGNALLING AND REGULATORY PATHWAYS | 1.07E-05 | 1.71E-04 | 1.44E-03 | 2.64E-02 | [5](https://toppgene.cchmc.org/showQueryTerms.jsp?userdata_id=e0bdc6b9-955d-48cd-95d3-4ccb1cbe49c7&feature=pt&row=153) |
| DNM2 | M27565 | REACTOME MAPK FAMILY SIGNALING CASCADES | 1.76E-04 | 4.10E-03 | 3.59E-02 | 6.35E-01 | [24](https://toppgene.cchmc.org/showQueryTerms.jsp?userdata_id=677e6385-c8f1-4db4-acb1-4b26e7d30bc3&feature=pt&row=154) | ITM2B | M48031 | REACTOME SOMITOGENESIS | 1.15E-05 | 1.79E-04 | 1.50E-03 | 2.85E-02 | [8](https://toppgene.cchmc.org/showQueryTerms.jsp?userdata_id=e0bdc6b9-955d-48cd-95d3-4ccb1cbe49c7&feature=pt&row=154) |
| DNPEP | M41822 | REACTOME SENSORY PROCESSING OF SOUND | 1.85E-04 | 4.28E-03 | 3.75E-02 | 6.68E-01 | [10](https://toppgene.cchmc.org/showQueryTerms.jsp?userdata_id=677e6385-c8f1-4db4-acb1-4b26e7d30bc3&feature=pt&row=155) | NID1 | MM15153 | REACTOME DEGRADATION OF DVL | 1.15E-05 | 1.79E-04 | 1.50E-03 | 2.85E-02 | [8](https://toppgene.cchmc.org/showQueryTerms.jsp?userdata_id=e0bdc6b9-955d-48cd-95d3-4ccb1cbe49c7&feature=pt&row=155) |
| DOPEY2 | M1088 | REACTOME METABOLISM OF PORPHYRINS | 1.88E-04 | 4.28E-03 | 3.75E-02 | 6.78E-01 | [6](https://toppgene.cchmc.org/showQueryTerms.jsp?userdata_id=677e6385-c8f1-4db4-acb1-4b26e7d30bc3&feature=pt&row=156) | PSMA3 | M27398 | REACTOME DEGRADATION OF AXIN | 1.15E-05 | 1.79E-04 | 1.50E-03 | 2.85E-02 | [8](https://toppgene.cchmc.org/showQueryTerms.jsp?userdata_id=e0bdc6b9-955d-48cd-95d3-4ccb1cbe49c7&feature=pt&row=156) |
| DPP4 | MM15908 | WP GLUTATHIONE METABOLISM | 1.88E-04 | 4.28E-03 | 3.75E-02 | 6.78E-01 | [6](https://toppgene.cchmc.org/showQueryTerms.jsp?userdata_id=677e6385-c8f1-4db4-acb1-4b26e7d30bc3&feature=pt&row=157) | PSMB6 | M27809 | REACTOME REGULATION OF RUNX3 EXPRESSION AND ACTIVITY | 1.15E-05 | 1.79E-04 | 1.50E-03 | 2.85E-02 | [8](https://toppgene.cchmc.org/showQueryTerms.jsp?userdata_id=e0bdc6b9-955d-48cd-95d3-4ccb1cbe49c7&feature=pt&row=157) |
| DSG3 | M1840 | KEGG GLUTATHIONE METABOLISM | 1.89E-04 | 4.28E-03 | 3.75E-02 | 6.80E-01 | [8](https://toppgene.cchmc.org/showQueryTerms.jsp?userdata_id=677e6385-c8f1-4db4-acb1-4b26e7d30bc3&feature=pt&row=158) | RAC1 | M17095 | REACTOME SCF BETA TRCP MEDIATED DEGRADATION OF EMI1 | 1.15E-05 | 1.79E-04 | 1.50E-03 | 2.85E-02 | [8](https://toppgene.cchmc.org/showQueryTerms.jsp?userdata_id=e0bdc6b9-955d-48cd-95d3-4ccb1cbe49c7&feature=pt&row=158) |
| DSP | MM15456 | REACTOME PLATELET AGGREGATION PLUG FORMATION | 1.92E-04 | 4.30E-03 | 3.77E-02 | 6.92E-01 | [7](https://toppgene.cchmc.org/showQueryTerms.jsp?userdata_id=677e6385-c8f1-4db4-acb1-4b26e7d30bc3&feature=pt&row=159) | RHEB | M41 | PID ER NONGENOMIC PATHWAY | 1.17E-05 | 1.81E-04 | 1.52E-03 | 2.90E-02 | [7](https://toppgene.cchmc.org/showQueryTerms.jsp?userdata_id=e0bdc6b9-955d-48cd-95d3-4ccb1cbe49c7&feature=pt&row=159) |
| DYNC1H1 | MM14519 | REACTOME ROS AND RNS PRODUCTION IN PHAGOCYTES | 1.92E-04 | 4.30E-03 | 3.77E-02 | 6.92E-01 | [7](https://toppgene.cchmc.org/showQueryTerms.jsp?userdata_id=677e6385-c8f1-4db4-acb1-4b26e7d30bc3&feature=pt&row=160) | RRAS | MM15491 | REACTOME THE ROLE OF GTSE1 IN G2 M PROGRESSION AFTER G2 CHECKPOINT | 1.31E-05 | 2.01E-04 | 1.68E-03 | 3.23E-02 | [9](https://toppgene.cchmc.org/showQueryTerms.jsp?userdata_id=e0bdc6b9-955d-48cd-95d3-4ccb1cbe49c7&feature=pt&row=160) |
| DYNC2H1 | M27285 | REACTOME REGULATION OF INSULIN LIKE GROWTH FACTOR IGF TRANSPORT AND UPTAKE BY INSULIN LIKE GROWTH FACTOR BINDING PROTEINS IGFBPS | 1.95E-04 | 4.34E-03 | 3.81E-02 | 7.04E-01 | [13](https://toppgene.cchmc.org/showQueryTerms.jsp?userdata_id=677e6385-c8f1-4db4-acb1-4b26e7d30bc3&feature=pt&row=161) | RRAS2 | MM15203 | REACTOME DECTIN 1 MEDIATED NONCANONICAL NF KB SIGNALING | 1.32E-05 | 2.02E-04 | 1.69E-03 | 3.27E-02 | [8](https://toppgene.cchmc.org/showQueryTerms.jsp?userdata_id=e0bdc6b9-955d-48cd-95d3-4ccb1cbe49c7&feature=pt&row=161) |
| DYNLRB1 | M17034 | REACTOME INTEGRATION OF ENERGY METABOLISM | 1.98E-04 | 4.37E-03 | 3.83E-02 | 7.12E-01 | [12](https://toppgene.cchmc.org/showQueryTerms.jsp?userdata_id=677e6385-c8f1-4db4-acb1-4b26e7d30bc3&feature=pt&row=162) | SLC3A2 | M27322 | REACTOME PCP CE PATHWAY | 1.35E-05 | 2.05E-04 | 1.72E-03 | 3.33E-02 | [10](https://toppgene.cchmc.org/showQueryTerms.jsp?userdata_id=e0bdc6b9-955d-48cd-95d3-4ccb1cbe49c7&feature=pt&row=162) |
| DYSF | MM15178 | REACTOME PROGRAMMED CELL DEATH | 2.00E-04 | 4.39E-03 | 3.85E-02 | 7.20E-01 | [14](https://toppgene.cchmc.org/showQueryTerms.jsp?userdata_id=677e6385-c8f1-4db4-acb1-4b26e7d30bc3&feature=pt&row=163) | SRI | MM14755 | REACTOME RHO GTPASE EFFECTORS | 1.44E-05 | 2.17E-04 | 1.82E-03 | 3.56E-02 | [17](https://toppgene.cchmc.org/showQueryTerms.jsp?userdata_id=e0bdc6b9-955d-48cd-95d3-4ccb1cbe49c7&feature=pt&row=163) |
| EEA1 | M39337 | WP VITAMIN B12 METABOLISM | 2.18E-04 | 4.69E-03 | 4.12E-02 | 7.84E-01 | [8](https://toppgene.cchmc.org/showQueryTerms.jsp?userdata_id=677e6385-c8f1-4db4-acb1-4b26e7d30bc3&feature=pt&row=164) | TSG101 | M1519 | KEGG ENDOCYTOSIS | 1.47E-05 | 2.20E-04 | 1.85E-03 | 3.63E-02 | [14](https://toppgene.cchmc.org/showQueryTerms.jsp?userdata_id=e0bdc6b9-955d-48cd-95d3-4ccb1cbe49c7&feature=pt&row=164) |
| EEF1A1 | MM15928 | WP GLYCOLYSIS AND GLUCONEOGENESIS | 2.18E-04 | 4.69E-03 | 4.12E-02 | 7.84E-01 | [8](https://toppgene.cchmc.org/showQueryTerms.jsp?userdata_id=677e6385-c8f1-4db4-acb1-4b26e7d30bc3&feature=pt&row=165) | DNAJA1 | MM15205 | REACTOME GLI3 IS PROCESSED TO GLI3R BY THE PROTEASOME | 1.51E-05 | 2.21E-04 | 1.86E-03 | 3.74E-02 | [8](https://toppgene.cchmc.org/showQueryTerms.jsp?userdata_id=e0bdc6b9-955d-48cd-95d3-4ccb1cbe49c7&feature=pt&row=165) |
| EEF2 | M27480 | REACTOME CARGO TRAFFICKING TO THE PERICILIARY MEMBRANE | 2.18E-04 | 4.69E-03 | 4.12E-02 | 7.84E-01 | [8](https://toppgene.cchmc.org/showQueryTerms.jsp?userdata_id=677e6385-c8f1-4db4-acb1-4b26e7d30bc3&feature=pt&row=166) | LAMP2 | M27399 | REACTOME DEGRADATION OF DVL | 1.51E-05 | 2.21E-04 | 1.86E-03 | 3.74E-02 | [8](https://toppgene.cchmc.org/showQueryTerms.jsp?userdata_id=e0bdc6b9-955d-48cd-95d3-4ccb1cbe49c7&feature=pt&row=166) |
| EGF | M10735 | REACTOME PLATELET AGGREGATION PLUG FORMATION | 2.28E-04 | 4.85E-03 | 4.26E-02 | 8.20E-01 | [7](https://toppgene.cchmc.org/showQueryTerms.jsp?userdata_id=677e6385-c8f1-4db4-acb1-4b26e7d30bc3&feature=pt&row=167) | ABI1 | M27670 | REACTOME STABILIZATION OF P53 | 1.51E-05 | 2.21E-04 | 1.86E-03 | 3.74E-02 | [8](https://toppgene.cchmc.org/showQueryTerms.jsp?userdata_id=e0bdc6b9-955d-48cd-95d3-4ccb1cbe49c7&feature=pt&row=167) |
| EGFR | MM15607 | REACTOME RHOU GTPASE CYCLE | 2.28E-04 | 4.85E-03 | 4.26E-02 | 8.20E-01 | [7](https://toppgene.cchmc.org/showQueryTerms.jsp?userdata_id=677e6385-c8f1-4db4-acb1-4b26e7d30bc3&feature=pt&row=168) | ALDH1L1 | MM14955 | REACTOME METABOLISM OF POLYAMINES | 1.51E-05 | 2.21E-04 | 1.86E-03 | 3.74E-02 | [8](https://toppgene.cchmc.org/showQueryTerms.jsp?userdata_id=e0bdc6b9-955d-48cd-95d3-4ccb1cbe49c7&feature=pt&row=168) |
| EHD2 | M47654 | KEGG MEDICUS REFERENCE ITGA B RHOG RAC SIGNALING PATHWAY | 2.33E-04 | 4.94E-03 | 4.33E-02 | 8.40E-01 | [6](https://toppgene.cchmc.org/showQueryTerms.jsp?userdata_id=677e6385-c8f1-4db4-acb1-4b26e7d30bc3&feature=pt&row=169) | ANXA13 | M174 | PID UPA UPAR PATHWAY | 1.64E-05 | 2.38E-04 | 2.00E-03 | 4.05E-02 | [7](https://toppgene.cchmc.org/showQueryTerms.jsp?userdata_id=e0bdc6b9-955d-48cd-95d3-4ccb1cbe49c7&feature=pt&row=169) |
| EIF3H | M2341 | REACTOME APOPTOTIC EXECUTION PHASE | 2.50E-04 | 5.27E-03 | 4.62E-02 | 9.00E-01 | [8](https://toppgene.cchmc.org/showQueryTerms.jsp?userdata_id=677e6385-c8f1-4db4-acb1-4b26e7d30bc3&feature=pt&row=170) | ARPC5 | MM14715 | REACTOME SCF SKP2 MEDIATED DEGRADATION OF P27 P21 | 1.72E-05 | 2.49E-04 | 2.09E-03 | 4.26E-02 | [8](https://toppgene.cchmc.org/showQueryTerms.jsp?userdata_id=e0bdc6b9-955d-48cd-95d3-4ccb1cbe49c7&feature=pt&row=170) |
| EML5 | M29804 | REACTOME HCMV INFECTION | 2.59E-04 | 5.43E-03 | 4.76E-02 | 9.34E-01 | [15](https://toppgene.cchmc.org/showQueryTerms.jsp?userdata_id=677e6385-c8f1-4db4-acb1-4b26e7d30bc3&feature=pt&row=171) | ARRDC1 | M22072 | BIOCARTA ALTERNATIVE PATHWAY | 1.75E-05 | 2.51E-04 | 2.11E-03 | 4.32E-02 | [4](https://toppgene.cchmc.org/showQueryTerms.jsp?userdata_id=e0bdc6b9-955d-48cd-95d3-4ccb1cbe49c7&feature=pt&row=171) |
| ENPEP | MM15925 | WP ALPHA 6 BETA 4 INTEGRIN SIGNALING PATHWAY | 2.66E-04 | 5.55E-03 | 4.86E-02 | 9.59E-01 | [9](https://toppgene.cchmc.org/showQueryTerms.jsp?userdata_id=677e6385-c8f1-4db4-acb1-4b26e7d30bc3&feature=pt&row=172) | ATP6V1D | M42565 | WP ALZHEIMER 39 S DISEASE | 1.76E-05 | 2.52E-04 | 2.11E-03 | 4.35E-02 | [17](https://toppgene.cchmc.org/showQueryTerms.jsp?userdata_id=e0bdc6b9-955d-48cd-95d3-4ccb1cbe49c7&feature=pt&row=172) |
| ENPP6 | M41816 | REACTOME RHOU GTPASE CYCLE | 2.69E-04 | 5.56E-03 | 4.87E-02 | 9.67E-01 | [7](https://toppgene.cchmc.org/showQueryTerms.jsp?userdata_id=677e6385-c8f1-4db4-acb1-4b26e7d30bc3&feature=pt&row=173) | CHMP1B | M27218 | REACTOME NON INTEGRIN MEMBRANE ECM INTERACTIONS | 1.96E-05 | 2.76E-04 | 2.32E-03 | 4.85E-02 | [8](https://toppgene.cchmc.org/showQueryTerms.jsp?userdata_id=e0bdc6b9-955d-48cd-95d3-4ccb1cbe49c7&feature=pt&row=173) |
| EPCAM | M39777 | WP DISORDERS OF FOLATE METABOLISM AND TRANSPORT | 2.82E-04 | 5.69E-03 | 4.99E-02 | 1.00E+00 | [5](https://toppgene.cchmc.org/showQueryTerms.jsp?userdata_id=677e6385-c8f1-4db4-acb1-4b26e7d30bc3&feature=pt&row=174) | CPNE5 | M747 | REACTOME METABOLISM OF POLYAMINES | 1.96E-05 | 2.76E-04 | 2.32E-03 | 4.85E-02 | [8](https://toppgene.cchmc.org/showQueryTerms.jsp?userdata_id=e0bdc6b9-955d-48cd-95d3-4ccb1cbe49c7&feature=pt&row=174) |
| EPHB1 | M47862 | KEGG MEDICUS REFERENCE ASSEMBLY AND TRAFFICKING OF TELOMERASE | 2.82E-04 | 5.69E-03 | 4.99E-02 | 1.00E+00 | [5](https://toppgene.cchmc.org/showQueryTerms.jsp?userdata_id=677e6385-c8f1-4db4-acb1-4b26e7d30bc3&feature=pt&row=175) | ECE1 | M45026 | REACTOME NUCLEAR EVENTS MEDIATED BY NFE2L2 | 1.97E-05 | 2.76E-04 | 2.32E-03 | 4.86E-02 | [10](https://toppgene.cchmc.org/showQueryTerms.jsp?userdata_id=e0bdc6b9-955d-48cd-95d3-4ccb1cbe49c7&feature=pt&row=175) |
| EPHX2 | M194 | BIOCARTA PROTEASOME PATHWAY | 2.82E-04 | 5.69E-03 | 4.99E-02 | 1.00E+00 | [5](https://toppgene.cchmc.org/showQueryTerms.jsp?userdata_id=677e6385-c8f1-4db4-acb1-4b26e7d30bc3&feature=pt&row=176) | EEF1 | MM15698 | REACTOME KEAP1 NFE2L2 PATHWAY | 2.03E-05 | 2.82E-04 | 2.36E-03 | 5.01E-02 | [9](https://toppgene.cchmc.org/showQueryTerms.jsp?userdata_id=e0bdc6b9-955d-48cd-95d3-4ccb1cbe49c7&feature=pt&row=176) |
| EPPK1 | MM15225 | REACTOME HEDGEHOG ON STATE | 2.83E-04 | 5.69E-03 | 4.99E-02 | 1.00E+00 | [10](https://toppgene.cchmc.org/showQueryTerms.jsp?userdata_id=677e6385-c8f1-4db4-acb1-4b26e7d30bc3&feature=pt&row=177) | EHD4 | MM15024 | REACTOME EPHRIN SIGNALING | 2.03E-05 | 2.82E-04 | 2.36E-03 | 5.01E-02 | [5](https://toppgene.cchmc.org/showQueryTerms.jsp?userdata_id=e0bdc6b9-955d-48cd-95d3-4ccb1cbe49c7&feature=pt&row=177) |
| EPS8 | MM1522 | BIOCARTA TFF PATHWAY | 2.86E-04 | 5.69E-03 | 4.99E-02 | 1.00E+00 | [6](https://toppgene.cchmc.org/showQueryTerms.jsp?userdata_id=677e6385-c8f1-4db4-acb1-4b26e7d30bc3&feature=pt&row=178) | GNG5 | M27470 | REACTOME DEGRADATION OF GLI1 BY THE PROTEASOME | 2.22E-05 | 3.01E-04 | 2.52E-03 | 5.50E-02 | [8](https://toppgene.cchmc.org/showQueryTerms.jsp?userdata_id=e0bdc6b9-955d-48cd-95d3-4ccb1cbe49c7&feature=pt&row=178) |
| ERMN | M113 | PID NFAT 3PATHWAY | 2.86E-04 | 5.69E-03 | 4.99E-02 | 1.00E+00 | [8](https://toppgene.cchmc.org/showQueryTerms.jsp?userdata_id=677e6385-c8f1-4db4-acb1-4b26e7d30bc3&feature=pt&row=179) | GOLGA7 | MM15150 | REACTOME ASYMMETRIC LOCALIZATION OF PCP PROTEINS | 2.22E-05 | 3.01E-04 | 2.52E-03 | 5.50E-02 | [8](https://toppgene.cchmc.org/showQueryTerms.jsp?userdata_id=e0bdc6b9-955d-48cd-95d3-4ccb1cbe49c7&feature=pt&row=179) |
| EVPL | MM15307 | REACTOME POST TRANSLATIONAL PROTEIN MODIFICATION | 2.87E-04 | 5.69E-03 | 4.99E-02 | 1.00E+00 | [68](https://toppgene.cchmc.org/showQueryTerms.jsp?userdata_id=677e6385-c8f1-4db4-acb1-4b26e7d30bc3&feature=pt&row=180) | IST1 | M1081 | REACTOME SCF SKP2 MEDIATED DEGRADATION OF P27 P21 | 2.22E-05 | 3.01E-04 | 2.52E-03 | 5.50E-02 | [8](https://toppgene.cchmc.org/showQueryTerms.jsp?userdata_id=e0bdc6b9-955d-48cd-95d3-4ccb1cbe49c7&feature=pt&row=180) |
| F11 | M27288 | REACTOME BETA CATENIN INDEPENDENT WNT SIGNALING | 2.88E-04 | 5.69E-03 | 4.99E-02 | 1.00E+00 | [14](https://toppgene.cchmc.org/showQueryTerms.jsp?userdata_id=677e6385-c8f1-4db4-acb1-4b26e7d30bc3&feature=pt&row=181) | MFGE8 | M39379 | WP ALZHEIMER 39 S DISEASE AND MIRNA EFFECTS | 2.25E-05 | 3.01E-04 | 2.52E-03 | 5.56E-02 | [17](https://toppgene.cchmc.org/showQueryTerms.jsp?userdata_id=e0bdc6b9-955d-48cd-95d3-4ccb1cbe49c7&feature=pt&row=181) |
| FABP1 | MM14732 | REACTOME GAP JUNCTION DEGRADATION | 3.03E-04 | 5.89E-03 | 5.16E-02 | 1.00E+00 | [4](https://toppgene.cchmc.org/showQueryTerms.jsp?userdata_id=677e6385-c8f1-4db4-acb1-4b26e7d30bc3&feature=pt&row=182) | PAFAH1B1 | M27473 | REACTOME ABC TRANSPORTER DISORDERS | 2.25E-05 | 3.01E-04 | 2.52E-03 | 5.57E-02 | [9](https://toppgene.cchmc.org/showQueryTerms.jsp?userdata_id=e0bdc6b9-955d-48cd-95d3-4ccb1cbe49c7&feature=pt&row=182) |
| FABP3 | M47435 | KEGG MEDICUS REFERENCE EGF EGFR RAS RALGDS SIGNALING PATHWAY | 3.03E-04 | 5.89E-03 | 5.16E-02 | 1.00E+00 | [4](https://toppgene.cchmc.org/showQueryTerms.jsp?userdata_id=677e6385-c8f1-4db4-acb1-4b26e7d30bc3&feature=pt&row=183) | PARK7 | MM14513 | REACTOME DOWNSTREAM SIGNALING EVENTS OF B CELL RECEPTOR BCR | 2.25E-05 | 3.01E-04 | 2.52E-03 | 5.57E-02 | [9](https://toppgene.cchmc.org/showQueryTerms.jsp?userdata_id=e0bdc6b9-955d-48cd-95d3-4ccb1cbe49c7&feature=pt&row=183) |
| FAM209A | M47431 | KEGG MEDICUS REFERENCE EGF EGFR RAS RASSF1 SIGNALING PATHWAY | 3.03E-04 | 5.89E-03 | 5.16E-02 | 1.00E+00 | [4](https://toppgene.cchmc.org/showQueryTerms.jsp?userdata_id=677e6385-c8f1-4db4-acb1-4b26e7d30bc3&feature=pt&row=184) | PEF1 | M27772 | REACTOME MET ACTIVATES PTK2 SIGNALING | 2.26E-05 | 3.01E-04 | 2.52E-03 | 5.59E-02 | [6](https://toppgene.cchmc.org/showQueryTerms.jsp?userdata_id=e0bdc6b9-955d-48cd-95d3-4ccb1cbe49c7&feature=pt&row=184) |
| FAM20C | M776 | REACTOME PROTEIN FOLDING | 3.31E-04 | 6.38E-03 | 5.59E-02 | 1.00E+00 | [11](https://toppgene.cchmc.org/showQueryTerms.jsp?userdata_id=677e6385-c8f1-4db4-acb1-4b26e7d30bc3&feature=pt&row=185) | PLD3 | MM14924 | REACTOME NON INTEGRIN MEMBRANE ECM INTERACTIONS | 2.26E-05 | 3.01E-04 | 2.52E-03 | 5.59E-02 | [6](https://toppgene.cchmc.org/showQueryTerms.jsp?userdata_id=e0bdc6b9-955d-48cd-95d3-4ccb1cbe49c7&feature=pt&row=185) |
| FASN | MM15942 | WP AMINO ACID METABOLISM | 3.31E-04 | 6.38E-03 | 5.59E-02 | 1.00E+00 | [11](https://toppgene.cchmc.org/showQueryTerms.jsp?userdata_id=677e6385-c8f1-4db4-acb1-4b26e7d30bc3&feature=pt&row=186) | RHOG | M27080 | REACTOME RHO GTPASE EFFECTORS | 2.37E-05 | 3.14E-04 | 2.63E-03 | 5.86E-02 | [19](https://toppgene.cchmc.org/showQueryTerms.jsp?userdata_id=e0bdc6b9-955d-48cd-95d3-4ccb1cbe49c7&feature=pt&row=186) |
| FAT1 | M47717 | KEGG MEDICUS REFERENCE ITGA B FAK RAC SIGNALING PATHWAY | 3.48E-04 | 6.63E-03 | 5.81E-02 | 1.00E+00 | [6](https://toppgene.cchmc.org/showQueryTerms.jsp?userdata_id=677e6385-c8f1-4db4-acb1-4b26e7d30bc3&feature=pt&row=187) | S100A14 | MM15499 | REACTOME CLATHRIN MEDIATED ENDOCYTOSIS | 2.51E-05 | 3.30E-04 | 2.77E-03 | 6.20E-02 | [12](https://toppgene.cchmc.org/showQueryTerms.jsp?userdata_id=e0bdc6b9-955d-48cd-95d3-4ccb1cbe49c7&feature=pt&row=187) |
| FBL | M506 | REACTOME INSULIN RECEPTOR RECYCLING | 3.48E-04 | 6.63E-03 | 5.81E-02 | 1.00E+00 | [6](https://toppgene.cchmc.org/showQueryTerms.jsp?userdata_id=677e6385-c8f1-4db4-acb1-4b26e7d30bc3&feature=pt&row=188) | SH3BGRL3 | MM15548 | REACTOME CELLULAR RESPONSES TO STIMULI | 2.54E-05 | 3.32E-04 | 2.79E-03 | 6.28E-02 | [24](https://toppgene.cchmc.org/showQueryTerms.jsp?userdata_id=e0bdc6b9-955d-48cd-95d3-4ccb1cbe49c7&feature=pt&row=188) |
| FBP1 | M39719 | WP FOCAL ADHESION PI3K AKT MTOR SIGNALING PATHWAY | 3.61E-04 | 6.81E-03 | 5.97E-02 | 1.00E+00 | [22](https://toppgene.cchmc.org/showQueryTerms.jsp?userdata_id=677e6385-c8f1-4db4-acb1-4b26e7d30bc3&feature=pt&row=189) | SLC25A3 | MM15917 | WP FOCAL ADHESION PI3K AKT MTOR SIGNALING PATHWAY | 2.69E-05 | 3.49E-04 | 2.93E-03 | 6.66E-02 | [19](https://toppgene.cchmc.org/showQueryTerms.jsp?userdata_id=e0bdc6b9-955d-48cd-95d3-4ccb1cbe49c7&feature=pt&row=189) |
| FBP2 | MM14721 | REACTOME HEME DEGRADATION | 3.66E-04 | 6.81E-03 | 5.97E-02 | 1.00E+00 | [5](https://toppgene.cchmc.org/showQueryTerms.jsp?userdata_id=677e6385-c8f1-4db4-acb1-4b26e7d30bc3&feature=pt&row=190) | SNX12 | M27310 | REACTOME EPHRIN SIGNALING | 2.71E-05 | 3.49E-04 | 2.93E-03 | 6.69E-02 | [5](https://toppgene.cchmc.org/showQueryTerms.jsp?userdata_id=e0bdc6b9-955d-48cd-95d3-4ccb1cbe49c7&feature=pt&row=190) |
| FCGBP | M91 | PID TCPTP PATHWAY | 3.67E-04 | 6.81E-03 | 5.97E-02 | 1.00E+00 | [7](https://toppgene.cchmc.org/showQueryTerms.jsp?userdata_id=677e6385-c8f1-4db4-acb1-4b26e7d30bc3&feature=pt&row=191) | SOD1 | M27221 | REACTOME SCAVENGING BY CLASS A RECEPTORS | 2.71E-05 | 3.49E-04 | 2.93E-03 | 6.69E-02 | [5](https://toppgene.cchmc.org/showQueryTerms.jsp?userdata_id=e0bdc6b9-955d-48cd-95d3-4ccb1cbe49c7&feature=pt&row=191) |
| FCN2 | MM15018 | REACTOME METABOLISM OF PROTEINS | 3.70E-04 | 6.81E-03 | 5.97E-02 | 1.00E+00 | [81](https://toppgene.cchmc.org/showQueryTerms.jsp?userdata_id=677e6385-c8f1-4db4-acb1-4b26e7d30bc3&feature=pt&row=192) | STXBP1 | M19381 | REACTOME G2 M CHECKPOINTS | 2.78E-05 | 3.56E-04 | 2.98E-03 | 6.86E-02 | [13](https://toppgene.cchmc.org/showQueryTerms.jsp?userdata_id=e0bdc6b9-955d-48cd-95d3-4ccb1cbe49c7&feature=pt&row=192) |
| FGA | MM15137 | REACTOME AUF1 HNRNP D0 BINDS AND DESTABILIZES MRNA | 3.72E-04 | 6.81E-03 | 5.97E-02 | 1.00E+00 | [8](https://toppgene.cchmc.org/showQueryTerms.jsp?userdata_id=677e6385-c8f1-4db4-acb1-4b26e7d30bc3&feature=pt&row=193) | TBC1D21 | M39639 | WP PROTEASOME DEGRADATION | 2.84E-05 | 3.58E-04 | 3.01E-03 | 7.02E-02 | [8](https://toppgene.cchmc.org/showQueryTerms.jsp?userdata_id=e0bdc6b9-955d-48cd-95d3-4ccb1cbe49c7&feature=pt&row=193) |
| FIGNL1 | M41823 | REACTOME SENSORY PROCESSING OF SOUND BY OUTER HAIR CELLS OF THE COCHLEA | 3.72E-04 | 6.81E-03 | 5.97E-02 | 1.00E+00 | [8](https://toppgene.cchmc.org/showQueryTerms.jsp?userdata_id=677e6385-c8f1-4db4-acb1-4b26e7d30bc3&feature=pt&row=194) | VAMP8 | M27465 | REACTOME DECTIN 1 MEDIATED NONCANONICAL NF KB SIGNALING | 2.84E-05 | 3.58E-04 | 3.01E-03 | 7.02E-02 | [8](https://toppgene.cchmc.org/showQueryTerms.jsp?userdata_id=e0bdc6b9-955d-48cd-95d3-4ccb1cbe49c7&feature=pt&row=194) |
| FKBP4 | MM15098 | REACTOME GOLGI ASSOCIATED VESICLE BIOGENESIS | 3.72E-04 | 6.81E-03 | 5.97E-02 | 1.00E+00 | [8](https://toppgene.cchmc.org/showQueryTerms.jsp?userdata_id=677e6385-c8f1-4db4-acb1-4b26e7d30bc3&feature=pt&row=195) | VPS28 | MM15521 | REACTOME TRANSCRIPTIONAL REGULATION BY RUNX2 | 2.84E-05 | 3.58E-04 | 3.01E-03 | 7.02E-02 | [8](https://toppgene.cchmc.org/showQueryTerms.jsp?userdata_id=e0bdc6b9-955d-48cd-95d3-4ccb1cbe49c7&feature=pt&row=195) |
| FKBP5 | M16894 | KEGG COMPLEMENT AND COAGULATION CASCADES | 3.74E-04 | 6.81E-03 | 5.97E-02 | 1.00E+00 | [9](https://toppgene.cchmc.org/showQueryTerms.jsp?userdata_id=677e6385-c8f1-4db4-acb1-4b26e7d30bc3&feature=pt&row=196) | HP | M47878 | KEGG MEDICUS REFERENCE LECTIN PATHWAY OF COAGULATION CASCADE FIBRINOGEN TO FIBRIN | 2.87E-05 | 3.60E-04 | 3.02E-03 | 7.09E-02 | [4](https://toppgene.cchmc.org/showQueryTerms.jsp?userdata_id=e0bdc6b9-955d-48cd-95d3-4ccb1cbe49c7&feature=pt&row=196) |
| FLNA | M39617 | WP FOLATE METABOLISM | 3.74E-04 | 6.81E-03 | 5.97E-02 | 1.00E+00 | [9](https://toppgene.cchmc.org/showQueryTerms.jsp?userdata_id=677e6385-c8f1-4db4-acb1-4b26e7d30bc3&feature=pt&row=197) | MXRA8 | M239 | PID A6B1 A6B4 INTEGRIN PATHWAY | 3.04E-05 | 3.80E-04 | 3.19E-03 | 7.52E-02 | [7](https://toppgene.cchmc.org/showQueryTerms.jsp?userdata_id=e0bdc6b9-955d-48cd-95d3-4ccb1cbe49c7&feature=pt&row=197) |
| FLNB | M124 | PID CXCR4 PATHWAY | 3.95E-04 | 7.15E-03 | 6.27E-02 | 1.00E+00 | [11](https://toppgene.cchmc.org/showQueryTerms.jsp?userdata_id=677e6385-c8f1-4db4-acb1-4b26e7d30bc3&feature=pt&row=198) | ADAM10 | M29836 | REACTOME LEISHMANIA INFECTION | 3.08E-05 | 3.82E-04 | 3.20E-03 | 7.63E-02 | [15](https://toppgene.cchmc.org/showQueryTerms.jsp?userdata_id=e0bdc6b9-955d-48cd-95d3-4ccb1cbe49c7&feature=pt&row=198) |
| FLOT2 | M27935 | REACTOME AUTOPHAGY | 4.07E-04 | 7.33E-03 | 6.42E-02 | 1.00E+00 | [14](https://toppgene.cchmc.org/showQueryTerms.jsp?userdata_id=677e6385-c8f1-4db4-acb1-4b26e7d30bc3&feature=pt&row=199) | ADIPOQ | M27753 | REACTOME CLATHRIN MEDIATED ENDOCYTOSIS | 3.09E-05 | 3.82E-04 | 3.20E-03 | 7.64E-02 | [12](https://toppgene.cchmc.org/showQueryTerms.jsp?userdata_id=e0bdc6b9-955d-48cd-95d3-4ccb1cbe49c7&feature=pt&row=199) |
| FN1 |  |  |  |  |  |  |  | AFP |  |  |  |  |  |  |  |
| FRK |  |  |  |  |  |  |  | AGT |  |  |  |  |  |  |  |
| FTCD |  |  |  |  |  |  |  | APOB |  |  |  |  |  |  |  |
| FUCA1 |  |  |  |  |  |  |  | ATP1B1 |  |  |  |  |  |  |  |
| FXC1 |  |  |  |  |  |  |  | ATP2B2 |  |  |  |  |  |  |  |
| GAA |  |  |  |  |  |  |  | C1QA |  |  |  |  |  |  |  |
| GABRB2 |  |  |  |  |  |  |  | C1QC |  |  |  |  |  |  |  |
| GAK |  |  |  |  |  |  |  | C3 |  |  |  |  |  |  |  |
| GALM |  |  |  |  |  |  |  | C4 |  |  |  |  |  |  |  |
| GAPR1 |  |  |  |  |  |  |  | C6 |  |  |  |  |  |  |  |
| GBE1 |  |  |  |  |  |  |  | C7 |  |  |  |  |  |  |  |
| GDI2 |  |  |  |  |  |  |  | C8A |  |  |  |  |  |  |  |
| GEMIN4 |  |  |  |  |  |  |  | C8G |  |  |  |  |  |  |  |
| GFRAL |  |  |  |  |  |  |  | CACNA2D1 |  |  |  |  |  |  |  |
| GIPC2 |  |  |  |  |  |  |  | CARMIL1 |  |  |  |  |  |  |  |
| GK2 |  |  |  |  |  |  |  | CD5L |  |  |  |  |  |  |  |
| GLB1 |  |  |  |  |  |  |  | CLU |  |  |  |  |  |  |  |
| GLO1 |  |  |  |  |  |  |  | COL11A1 |  |  |  |  |  |  |  |
| GM2A |  |  |  |  |  |  |  | COL12A1 |  |  |  |  |  |  |  |
| GNAQ |  |  |  |  |  |  |  | COL1A1 |  |  |  |  |  |  |  |
| GNB1 |  |  |  |  |  |  |  | COL2A1 |  |  |  |  |  |  |  |
| GNG12 |  |  |  |  |  |  |  | COL3A1 |  |  |  |  |  |  |  |
| GPI |  |  |  |  |  |  |  | COL5A2 |  |  |  |  |  |  |  |
| GPR155 |  |  |  |  |  |  |  | COL6A2 |  |  |  |  |  |  |  |
| GPRASP3 |  |  |  |  |  |  |  | DIP2B |  |  |  |  |  |  |  |
| GPT |  |  |  |  |  |  |  | EMILIN1 |  |  |  |  |  |  |  |
| GPX4 |  |  |  |  |  |  |  | ENG |  |  |  |  |  |  |  |
| GRB2 |  |  |  |  |  |  |  | EPB41L2 |  |  |  |  |  |  |  |
| GRID1 |  |  |  |  |  |  |  | EPHA2 |  |  |  |  |  |  |  |
| GSN |  |  |  |  |  |  |  | EPHA3 |  |  |  |  |  |  |  |
| GSR |  |  |  |  |  |  |  | EPHA5 |  |  |  |  |  |  |  |
| GSS |  |  |  |  |  |  |  | EPHB2 |  |  |  |  |  |  |  |
| GSTA1 |  |  |  |  |  |  |  | EPHB3 |  |  |  |  |  |  |  |
| GSTP1 |  |  |  |  |  |  |  | EPHB4 |  |  |  |  |  |  |  |
| GUSB |  |  |  |  |  |  |  | F10 |  |  |  |  |  |  |  |
| HBB |  |  |  |  |  |  |  | F2 |  |  |  |  |  |  |  |
| HBS1L |  |  |  |  |  |  |  | F5 |  |  |  |  |  |  |  |
| HEBP1 |  |  |  |  |  |  |  | FAT4 |  |  |  |  |  |  |  |
| HEBP2 |  |  |  |  |  |  |  | FETUB |  |  |  |  |  |  |  |
| HGS |  |  |  |  |  |  |  | FGB |  |  |  |  |  |  |  |
| HIST1H1C |  |  |  |  |  |  |  | FGG |  |  |  |  |  |  |  |
| HIST1H4A |  |  |  |  |  |  |  | FMNL3 |  |  |  |  |  |  |  |
| HIST2H2AB |  |  |  |  |  |  |  | FSTL1 |  |  |  |  |  |  |  |
| HNRNPL |  |  |  |  |  |  |  | GC |  |  |  |  |  |  |  |
| HPD |  |  |  |  |  |  |  | HBE1 |  |  |  |  |  |  |  |
| HRAS |  |  |  |  |  |  |  | HTRA1 |  |  |  |  |  |  |  |
| HRG |  |  |  |  |  |  |  | IGSF8 |  |  |  |  |  |  |  |
| HSPA1B |  |  |  |  |  |  |  | ILK |  |  |  |  |  |  |  |
| HSPA2 |  |  |  |  |  |  |  | ITGA5 |  |  |  |  |  |  |  |
| HSPA5 |  |  |  |  |  |  |  | ITGA6 |  |  |  |  |  |  |  |
| HSPA8 |  |  |  |  |  |  |  | ITGB2 |  |  |  |  |  |  |  |
| HSPG2 |  |  |  |  |  |  |  | ITGB5 |  |  |  |  |  |  |  |
| HUWE1 |  |  |  |  |  |  |  | ITGB7 |  |  |  |  |  |  |  |
| HYAL2 |  |  |  |  |  |  |  | ITIH2 |  |  |  |  |  |  |  |
| ICAM1 |  |  |  |  |  |  |  | ITIH3 |  |  |  |  |  |  |  |
| IDH1 |  |  |  |  |  |  |  | ITIH4 |  |  |  |  |  |  |  |
| IGF2R |  |  |  |  |  |  |  | KIF11 |  |  |  |  |  |  |  |
| INADL |  |  |  |  |  |  |  | KRAS |  |  |  |  |  |  |  |
| IQGAP1 |  |  |  |  |  |  |  | KRT18 |  |  |  |  |  |  |  |
| IQGAP2 |  |  |  |  |  |  |  | KRT6B |  |  |  |  |  |  |  |
| IRF6 |  |  |  |  |  |  |  | KRT79 |  |  |  |  |  |  |  |
| ITGA1 |  |  |  |  |  |  |  | LAMB1 |  |  |  |  |  |  |  |
| ITGA3 |  |  |  |  |  |  |  | LOXL2 |  |  |  |  |  |  |  |
| ITGAV |  |  |  |  |  |  |  | MAMDC2 |  |  |  |  |  |  |  |
| ITGB1 |  |  |  |  |  |  |  | MASP1 |  |  |  |  |  |  |  |
| ITGB8 |  |  |  |  |  |  |  | MBL |  |  |  |  |  |  |  |
| ITLN1 |  |  |  |  |  |  |  | MMP2 |  |  |  |  |  |  |  |
| ITM2C |  |  |  |  |  |  |  | MPZL1 |  |  |  |  |  |  |  |
| ITSN2 |  |  |  |  |  |  |  | NID2 |  |  |  |  |  |  |  |
| JUP |  |  |  |  |  |  |  | NOTCH2 |  |  |  |  |  |  |  |
| KALRN |  |  |  |  |  |  |  | NOTCH3 |  |  |  |  |  |  |  |
| KCNQ2 |  |  |  |  |  |  |  | NRAS |  |  |  |  |  |  |  |
| KHK |  |  |  |  |  |  |  | NRP2 |  |  |  |  |  |  |  |
| KIF12 |  |  |  |  |  |  |  | ORM1 |  |  |  |  |  |  |  |
| KIF3A |  |  |  |  |  |  |  | PLAU |  |  |  |  |  |  |  |
| KIF3B |  |  |  |  |  |  |  | PLBD2 |  |  |  |  |  |  |  |
| KIFC3 |  |  |  |  |  |  |  | PLXNB2 |  |  |  |  |  |  |  |
| KNG1 |  |  |  |  |  |  |  | POSTN |  |  |  |  |  |  |  |
| KPNB1 |  |  |  |  |  |  |  | PSMB4 |  |  |  |  |  |  |  |
| LAMA5 |  |  |  |  |  |  |  | PTGFRN |  |  |  |  |  |  |  |
| LAMC1 |  |  |  |  |  |  |  | PTX3 |  |  |  |  |  |  |  |
| LAMP1 |  |  |  |  |  |  |  | PXDN |  |  |  |  |  |  |  |
| LAMTOR1 |  |  |  |  |  |  |  | ROR1 |  |  |  |  |  |  |  |
| LCP1 |  |  |  |  |  |  |  | RPL18A |  |  |  |  |  |  |  |
| LDHB |  |  |  |  |  |  |  | RPL28 |  |  |  |  |  |  |  |
| LEG1 |  |  |  |  |  |  |  | SCAMP3 |  |  |  |  |  |  |  |
| LGALS3 |  |  |  |  |  |  |  | SERPINA3 |  |  |  |  |  |  |  |
| LH3 |  |  |  |  |  |  |  | SERPIND1 |  |  |  |  |  |  |  |
| LRAT |  |  |  |  |  |  |  | SERPINF2 |  |  |  |  |  |  |  |
| LRP2 |  |  |  |  |  |  |  | SLC16A1 |  |  |  |  |  |  |  |
| LRRC15 |  |  |  |  |  |  |  | SLC1A4 |  |  |  |  |  |  |  |
| LRRK2 |  |  |  |  |  |  |  | SLC29A1 |  |  |  |  |  |  |  |
| LTA4H |  |  |  |  |  |  |  | SLC44A2 |  |  |  |  |  |  |  |
| LYPLA2 |  |  |  |  |  |  |  | SPP2 |  |  |  |  |  |  |  |
| MAN2A1 |  |  |  |  |  |  |  | TG |  |  |  |  |  |  |  |
| MAPK8IP4 |  |  |  |  |  |  |  | TGFBI |  |  |  |  |  |  |  |
| MBD5 |  |  |  |  |  |  |  | TTYH3 |  |  |  |  |  |  |  |
| MDH2 |  |  |  |  |  |  |  | VASN |  |  |  |  |  |  |  |
| MEGF8 |  |  |  |  |  |  |  | VPS37B |  |  |  |  |  |  |  |
| MELTF |  |  |  |  |  |  |  |  |  |  |  |  |  |  |  |
| MINK1 |  |  |  |  |  |  |  |  |  |  |  |  |  |  |  |
| MITF |  |  |  |  |  |  |  |  |  |  |  |  |  |  |  |
| MMRN2 |  |  |  |  |  |  |  |  |  |  |  |  |  |  |  |
| MNDA |  |  |  |  |  |  |  |  |  |  |  |  |  |  |  |
| MOGS |  |  |  |  |  |  |  |  |  |  |  |  |  |  |  |
| MPO |  |  |  |  |  |  |  |  |  |  |  |  |  |  |  |
| MRCKB |  |  |  |  |  |  |  |  |  |  |  |  |  |  |  |
| MSN |  |  |  |  |  |  |  |  |  |  |  |  |  |  |  |
| MTAP |  |  |  |  |  |  |  |  |  |  |  |  |  |  |  |
| MTHFD1 |  |  |  |  |  |  |  |  |  |  |  |  |  |  |  |
| MUC1 |  |  |  |  |  |  |  |  |  |  |  |  |  |  |  |
| MVB12A |  |  |  |  |  |  |  |  |  |  |  |  |  |  |  |
| MXRA5 |  |  |  |  |  |  |  |  |  |  |  |  |  |  |  |
| MYH13 |  |  |  |  |  |  |  |  |  |  |  |  |  |  |  |
| MYH14 |  |  |  |  |  |  |  |  |  |  |  |  |  |  |  |
| MYH3 |  |  |  |  |  |  |  |  |  |  |  |  |  |  |  |
| MYH9 |  |  |  |  |  |  |  |  |  |  |  |  |  |  |  |
| MYO15A |  |  |  |  |  |  |  |  |  |  |  |  |  |  |  |
| MYO1B |  |  |  |  |  |  |  |  |  |  |  |  |  |  |  |
| MYO1C |  |  |  |  |  |  |  |  |  |  |  |  |  |  |  |
| MYO1D |  |  |  |  |  |  |  |  |  |  |  |  |  |  |  |
| MYO5B |  |  |  |  |  |  |  |  |  |  |  |  |  |  |  |
| MYO6 |  |  |  |  |  |  |  |  |  |  |  |  |  |  |  |
| MYOF |  |  |  |  |  |  |  |  |  |  |  |  |  |  |  |
| NAA16 |  |  |  |  |  |  |  |  |  |  |  |  |  |  |  |
| NAA50 |  |  |  |  |  |  |  |  |  |  |  |  |  |  |  |
| NAGK |  |  |  |  |  |  |  |  |  |  |  |  |  |  |  |
| NAPEPLD |  |  |  |  |  |  |  |  |  |  |  |  |  |  |  |
| NAPSA |  |  |  |  |  |  |  |  |  |  |  |  |  |  |  |
| NBN |  |  |  |  |  |  |  |  |  |  |  |  |  |  |  |
| NCKAP1 |  |  |  |  |  |  |  |  |  |  |  |  |  |  |  |
| NDRG1 |  |  |  |  |  |  |  |  |  |  |  |  |  |  |  |
| NDRG3 |  |  |  |  |  |  |  |  |  |  |  |  |  |  |  |
| NEB |  |  |  |  |  |  |  |  |  |  |  |  |  |  |  |
| NEBL |  |  |  |  |  |  |  |  |  |  |  |  |  |  |  |
| NHERF1 |  |  |  |  |  |  |  |  |  |  |  |  |  |  |  |
| NIPBL |  |  |  |  |  |  |  |  |  |  |  |  |  |  |  |
| NPM1 |  |  |  |  |  |  |  |  |  |  |  |  |  |  |  |
| NQO1 |  |  |  |  |  |  |  |  |  |  |  |  |  |  |  |
| OLFM4 |  |  |  |  |  |  |  |  |  |  |  |  |  |  |  |
| OSBPL1 |  |  |  |  |  |  |  |  |  |  |  |  |  |  |  |
| OSR1 |  |  |  |  |  |  |  |  |  |  |  |  |  |  |  |
| P2RX4 |  |  |  |  |  |  |  |  |  |  |  |  |  |  |  |
| P4HB |  |  |  |  |  |  |  |  |  |  |  |  |  |  |  |
| PAFAH1B2 |  |  |  |  |  |  |  |  |  |  |  |  |  |  |  |
| PALS1 |  |  |  |  |  |  |  |  |  |  |  |  |  |  |  |
| PCBP1 |  |  |  |  |  |  |  |  |  |  |  |  |  |  |  |
| PDCD10 |  |  |  |  |  |  |  |  |  |  |  |  |  |  |  |
| PDE8A |  |  |  |  |  |  |  |  |  |  |  |  |  |  |  |
| PEBP1 |  |  |  |  |  |  |  |  |  |  |  |  |  |  |  |
| PEFLIN |  |  |  |  |  |  |  |  |  |  |  |  |  |  |  |
| PFN1 |  |  |  |  |  |  |  |  |  |  |  |  |  |  |  |
| PGD |  |  |  |  |  |  |  |  |  |  |  |  |  |  |  |
| PHB |  |  |  |  |  |  |  |  |  |  |  |  |  |  |  |
| PIK3C2A |  |  |  |  |  |  |  |  |  |  |  |  |  |  |  |
| PIK3C2B |  |  |  |  |  |  |  |  |  |  |  |  |  |  |  |
| PKD1 |  |  |  |  |  |  |  |  |  |  |  |  |  |  |  |
| PKD1L3 |  |  |  |  |  |  |  |  |  |  |  |  |  |  |  |
| PKD2 |  |  |  |  |  |  |  |  |  |  |  |  |  |  |  |
| PKHD1 |  |  |  |  |  |  |  |  |  |  |  |  |  |  |  |
| PKLR |  |  |  |  |  |  |  |  |  |  |  |  |  |  |  |
| PKM |  |  |  |  |  |  |  |  |  |  |  |  |  |  |  |
| PLEKHA1 |  |  |  |  |  |  |  |  |  |  |  |  |  |  |  |
| PLEKHA7 |  |  |  |  |  |  |  |  |  |  |  |  |  |  |  |
| PLS1 |  |  |  |  |  |  |  |  |  |  |  |  |  |  |  |
| PM20D1 |  |  |  |  |  |  |  |  |  |  |  |  |  |  |  |
| PNP |  |  |  |  |  |  |  |  |  |  |  |  |  |  |  |
| PPFIA2 |  |  |  |  |  |  |  |  |  |  |  |  |  |  |  |
| PPIA |  |  |  |  |  |  |  |  |  |  |  |  |  |  |  |
| PPL |  |  |  |  |  |  |  |  |  |  |  |  |  |  |  |
| PPP1R7 |  |  |  |  |  |  |  |  |  |  |  |  |  |  |  |
| PRDX5 |  |  |  |  |  |  |  |  |  |  |  |  |  |  |  |
| PRKAR2A |  |  |  |  |  |  |  |  |  |  |  |  |  |  |  |
| PRKCD |  |  |  |  |  |  |  |  |  |  |  |  |  |  |  |
| PRKCH |  |  |  |  |  |  |  |  |  |  |  |  |  |  |  |
| PROM1 |  |  |  |  |  |  |  |  |  |  |  |  |  |  |  |
| PRRC2A |  |  |  |  |  |  |  |  |  |  |  |  |  |  |  |
| PSA |  |  |  |  |  |  |  |  |  |  |  |  |  |  |  |
| PSAT1 |  |  |  |  |  |  |  |  |  |  |  |  |  |  |  |
| PSMA2 |  |  |  |  |  |  |  |  |  |  |  |  |  |  |  |
| PSMA4 |  |  |  |  |  |  |  |  |  |  |  |  |  |  |  |
| PSMB1 |  |  |  |  |  |  |  |  |  |  |  |  |  |  |  |
| PSMB2 |  |  |  |  |  |  |  |  |  |  |  |  |  |  |  |
| PSMD12 |  |  |  |  |  |  |  |  |  |  |  |  |  |  |  |
| PSMD8 |  |  |  |  |  |  |  |  |  |  |  |  |  |  |  |
| PTBP1 |  |  |  |  |  |  |  |  |  |  |  |  |  |  |  |
| PTGR1 |  |  |  |  |  |  |  |  |  |  |  |  |  |  |  |
| PTPN13 |  |  |  |  |  |  |  |  |  |  |  |  |  |  |  |
| PTPN23 |  |  |  |  |  |  |  |  |  |  |  |  |  |  |  |
| PTPN6 |  |  |  |  |  |  |  |  |  |  |  |  |  |  |  |
| PTTG1IP |  |  |  |  |  |  |  |  |  |  |  |  |  |  |  |
| PWWP3B |  |  |  |  |  |  |  |  |  |  |  |  |  |  |  |
| RAB10 |  |  |  |  |  |  |  |  |  |  |  |  |  |  |  |
| RAB15 |  |  |  |  |  |  |  |  |  |  |  |  |  |  |  |
| RAB19 |  |  |  |  |  |  |  |  |  |  |  |  |  |  |  |
| RAB21 |  |  |  |  |  |  |  |  |  |  |  |  |  |  |  |
| RAB27A |  |  |  |  |  |  |  |  |  |  |  |  |  |  |  |
| RAB2A |  |  |  |  |  |  |  |  |  |  |  |  |  |  |  |
| RAB33B |  |  |  |  |  |  |  |  |  |  |  |  |  |  |  |
| RAB3D |  |  |  |  |  |  |  |  |  |  |  |  |  |  |  |
| RAB3GAP1 |  |  |  |  |  |  |  |  |  |  |  |  |  |  |  |
| RAB7A |  |  |  |  |  |  |  |  |  |  |  |  |  |  |  |
| RAB7L1 |  |  |  |  |  |  |  |  |  |  |  |  |  |  |  |
| RAB8B |  |  |  |  |  |  |  |  |  |  |  |  |  |  |  |
| RACK1 |  |  |  |  |  |  |  |  |  |  |  |  |  |  |  |
| RAN |  |  |  |  |  |  |  |  |  |  |  |  |  |  |  |
| RAP2C |  |  |  |  |  |  |  |  |  |  |  |  |  |  |  |
| RAPGEF3 |  |  |  |  |  |  |  |  |  |  |  |  |  |  |  |
| RBL2 |  |  |  |  |  |  |  |  |  |  |  |  |  |  |  |
| RBM3 |  |  |  |  |  |  |  |  |  |  |  |  |  |  |  |
| RBP4 |  |  |  |  |  |  |  |  |  |  |  |  |  |  |  |
| RDH5 |  |  |  |  |  |  |  |  |  |  |  |  |  |  |  |
| REN |  |  |  |  |  |  |  |  |  |  |  |  |  |  |  |
| RFC1 |  |  |  |  |  |  |  |  |  |  |  |  |  |  |  |
| RHOA |  |  |  |  |  |  |  |  |  |  |  |  |  |  |  |
| RNASE1 |  |  |  |  |  |  |  |  |  |  |  |  |  |  |  |
| RNH1 |  |  |  |  |  |  |  |  |  |  |  |  |  |  |  |
| ROBO2 |  |  |  |  |  |  |  |  |  |  |  |  |  |  |  |
| RREB1 |  |  |  |  |  |  |  |  |  |  |  |  |  |  |  |
| RUSC2 |  |  |  |  |  |  |  |  |  |  |  |  |  |  |  |
| RYR1 |  |  |  |  |  |  |  |  |  |  |  |  |  |  |  |
| S100P |  |  |  |  |  |  |  |  |  |  |  |  |  |  |  |
| SAMM50 |  |  |  |  |  |  |  |  |  |  |  |  |  |  |  |
| SARG |  |  |  |  |  |  |  |  |  |  |  |  |  |  |  |
| SCEL |  |  |  |  |  |  |  |  |  |  |  |  |  |  |  |
| SCIN |  |  |  |  |  |  |  |  |  |  |  |  |  |  |  |
| SCN10A |  |  |  |  |  |  |  |  |  |  |  |  |  |  |  |
| SCPEP1 |  |  |  |  |  |  |  |  |  |  |  |  |  |  |  |
| SEC13 |  |  |  |  |  |  |  |  |  |  |  |  |  |  |  |
| SERPINA7 |  |  |  |  |  |  |  |  |  |  |  |  |  |  |  |
| SERPINB13 |  |  |  |  |  |  |  |  |  |  |  |  |  |  |  |
| SERPINB3 |  |  |  |  |  |  |  |  |  |  |  |  |  |  |  |
| SERPINF1 |  |  |  |  |  |  |  |  |  |  |  |  |  |  |  |
| SERPING1 |  |  |  |  |  |  |  |  |  |  |  |  |  |  |  |
| SFN |  |  |  |  |  |  |  |  |  |  |  |  |  |  |  |
| SH3BP4 |  |  |  |  |  |  |  |  |  |  |  |  |  |  |  |
| SHMT2 |  |  |  |  |  |  |  |  |  |  |  |  |  |  |  |
| SHROOM2 |  |  |  |  |  |  |  |  |  |  |  |  |  |  |  |
| SIAE |  |  |  |  |  |  |  |  |  |  |  |  |  |  |  |
| SLC12A1 |  |  |  |  |  |  |  |  |  |  |  |  |  |  |  |
| SLC15A2 |  |  |  |  |  |  |  |  |  |  |  |  |  |  |  |
| SLC1A1 |  |  |  |  |  |  |  |  |  |  |  |  |  |  |  |
| SLC22A12 |  |  |  |  |  |  |  |  |  |  |  |  |  |  |  |
| SLC22A2 |  |  |  |  |  |  |  |  |  |  |  |  |  |  |  |
| SLC22A4 |  |  |  |  |  |  |  |  |  |  |  |  |  |  |  |
| SLC25A1 |  |  |  |  |  |  |  |  |  |  |  |  |  |  |  |
| SLC39A5 |  |  |  |  |  |  |  |  |  |  |  |  |  |  |  |
| SLC4A4 |  |  |  |  |  |  |  |  |  |  |  |  |  |  |  |
| SLC5A10 |  |  |  |  |  |  |  |  |  |  |  |  |  |  |  |
| SLC5A2 |  |  |  |  |  |  |  |  |  |  |  |  |  |  |  |
| SLC5A4 |  |  |  |  |  |  |  |  |  |  |  |  |  |  |  |
| SLIT2 |  |  |  |  |  |  |  |  |  |  |  |  |  |  |  |
| SMC2 |  |  |  |  |  |  |  |  |  |  |  |  |  |  |  |
| SMO |  |  |  |  |  |  |  |  |  |  |  |  |  |  |  |
| SMPDL3B |  |  |  |  |  |  |  |  |  |  |  |  |  |  |  |
| SMURF1 |  |  |  |  |  |  |  |  |  |  |  |  |  |  |  |
| SNCG |  |  |  |  |  |  |  |  |  |  |  |  |  |  |  |
| SNF8 |  |  |  |  |  |  |  |  |  |  |  |  |  |  |  |
| SNX18 |  |  |  |  |  |  |  |  |  |  |  |  |  |  |  |
| SNX3 |  |  |  |  |  |  |  |  |  |  |  |  |  |  |  |
| SNX9 |  |  |  |  |  |  |  |  |  |  |  |  |  |  |  |
| SOGA1 |  |  |  |  |  |  |  |  |  |  |  |  |  |  |  |
| SORL1 |  |  |  |  |  |  |  |  |  |  |  |  |  |  |  |
| SPAST |  |  |  |  |  |  |  |  |  |  |  |  |  |  |  |
| SPATA31H1 |  |  |  |  |  |  |  |  |  |  |  |  |  |  |  |
| SPEN |  |  |  |  |  |  |  |  |  |  |  |  |  |  |  |
| SPON2 |  |  |  |  |  |  |  |  |  |  |  |  |  |  |  |
| SPTBN1 |  |  |  |  |  |  |  |  |  |  |  |  |  |  |  |
| STK11 |  |  |  |  |  |  |  |  |  |  |  |  |  |  |  |
| STK25 |  |  |  |  |  |  |  |  |  |  |  |  |  |  |  |
| STRIP1 |  |  |  |  |  |  |  |  |  |  |  |  |  |  |  |
| STXBP2 |  |  |  |  |  |  |  |  |  |  |  |  |  |  |  |
| STXBP4 |  |  |  |  |  |  |  |  |  |  |  |  |  |  |  |
| SUSD2 |  |  |  |  |  |  |  |  |  |  |  |  |  |  |  |
| SYNPO |  |  |  |  |  |  |  |  |  |  |  |  |  |  |  |
| TAOK1 |  |  |  |  |  |  |  |  |  |  |  |  |  |  |  |
| TASOR2 |  |  |  |  |  |  |  |  |  |  |  |  |  |  |  |
| TAX1BP3 |  |  |  |  |  |  |  |  |  |  |  |  |  |  |  |
| TBC1D10A |  |  |  |  |  |  |  |  |  |  |  |  |  |  |  |
| TCP1 |  |  |  |  |  |  |  |  |  |  |  |  |  |  |  |
| TECTA |  |  |  |  |  |  |  |  |  |  |  |  |  |  |  |
| TEKT3 |  |  |  |  |  |  |  |  |  |  |  |  |  |  |  |
| TF |  |  |  |  |  |  |  |  |  |  |  |  |  |  |  |
| TGM2 |  |  |  |  |  |  |  |  |  |  |  |  |  |  |  |
| TGM4 |  |  |  |  |  |  |  |  |  |  |  |  |  |  |  |
| THBS1 |  |  |  |  |  |  |  |  |  |  |  |  |  |  |  |
| TKFC |  |  |  |  |  |  |  |  |  |  |  |  |  |  |  |
| TLN1 |  |  |  |  |  |  |  |  |  |  |  |  |  |  |  |
| TMPRSS2 |  |  |  |  |  |  |  |  |  |  |  |  |  |  |  |
| TOLLIP |  |  |  |  |  |  |  |  |  |  |  |  |  |  |  |
| TOM1L2 |  |  |  |  |  |  |  |  |  |  |  |  |  |  |  |
| TSPAN3 |  |  |  |  |  |  |  |  |  |  |  |  |  |  |  |
| TSPAN6 |  |  |  |  |  |  |  |  |  |  |  |  |  |  |  |
| TTN |  |  |  |  |  |  |  |  |  |  |  |  |  |  |  |
| TUBB2A |  |  |  |  |  |  |  |  |  |  |  |  |  |  |  |
| TUBB6 |  |  |  |  |  |  |  |  |  |  |  |  |  |  |  |
| TUT4 |  |  |  |  |  |  |  |  |  |  |  |  |  |  |  |
| UACA |  |  |  |  |  |  |  |  |  |  |  |  |  |  |  |
| UBA1 |  |  |  |  |  |  |  |  |  |  |  |  |  |  |  |
| UBAC1 |  |  |  |  |  |  |  |  |  |  |  |  |  |  |  |
| UBL3 |  |  |  |  |  |  |  |  |  |  |  |  |  |  |  |
| UGP2 |  |  |  |  |  |  |  |  |  |  |  |  |  |  |  |
| UMOD |  |  |  |  |  |  |  |  |  |  |  |  |  |  |  |
| UPB1 |  |  |  |  |  |  |  |  |  |  |  |  |  |  |  |
| UTRN |  |  |  |  |  |  |  |  |  |  |  |  |  |  |  |
| VAMP7 |  |  |  |  |  |  |  |  |  |  |  |  |  |  |  |
| VASP |  |  |  |  |  |  |  |  |  |  |  |  |  |  |  |
| VAT1 |  |  |  |  |  |  |  |  |  |  |  |  |  |  |  |
| VCL |  |  |  |  |  |  |  |  |  |  |  |  |  |  |  |
| VDAC1 |  |  |  |  |  |  |  |  |  |  |  |  |  |  |  |
| VIL1 |  |  |  |  |  |  |  |  |  |  |  |  |  |  |  |
| VPS13C |  |  |  |  |  |  |  |  |  |  |  |  |  |  |  |
| VPS13D |  |  |  |  |  |  |  |  |  |  |  |  |  |  |  |
| VPS4B |  |  |  |  |  |  |  |  |  |  |  |  |  |  |  |
| VWF |  |  |  |  |  |  |  |  |  |  |  |  |  |  |  |
| WASL |  |  |  |  |  |  |  |  |  |  |  |  |  |  |  |
| WIZ |  |  |  |  |  |  |  |  |  |  |  |  |  |  |  |
| WNT5B |  |  |  |  |  |  |  |  |  |  |  |  |  |  |  |
| XPNPEP1 |  |  |  |  |  |  |  |  |  |  |  |  |  |  |  |
| XPNPEP2 |  |  |  |  |  |  |  |  |  |  |  |  |  |  |  |
| XPO2 |  |  |  |  |  |  |  |  |  |  |  |  |  |  |  |
| YWHAH |  |  |  |  |  |  |  |  |  |  |  |  |  |  |  |
| ZNF114 |  |  |  |  |  |  |  |  |  |  |  |  |  |  |  |
